# Heterogeneity of circulating epithelial cells in breast cancer at single-cell resolution: identifying tumor and hybrid cells

**DOI:** 10.1101/2021.11.24.469962

**Authors:** Maxim E. Menyailo, Viktoria R. Zainullina, Liubov A. Tashireva, Sofia Yu. Zolotareva, Tatiana S. Gerashchenko, Vladimir V. Alifanov, Olga E. Savelieva, Evgeniya S. Grigoryeva, Nataliya A. Tarabanovskaya, Nataliya O. Popova, Anna A. Khozyainova, Evgeny L. Choinzonov, Nadezhda V. Cherdyntseva, Vladimir M. Perelmuter, Evgeny V. Denisov

## Abstract

Circulating tumor cells and hybrid cells formed by the fusion of tumor cells with normal cells are leading players in metastasis and have prognostic relevance. Circulating tumor cells and hybrid cells are identified as CD45-negative and CD45-positive epithelial cells. However, such an approach is challenging because epithelial cells are observed in the blood of healthy individuals. In this study, we applied single-cell RNA sequencing to profile CD45-negative and CD45-positive circulating epithelial cells (CECs) in 20 breast cancer patients and one healthy donor. DNA ploidy analysis was used to identify the tumor and hybrid cells among CD45^─^ and CD45^+^ CECs in patients, respectively. Functional enrichment analysis was applied to characterize aneuploid and diploid cells. Diploid cells were also annotated to generate cell-type candidates and analyzed for copy-number aberrations (CNAs) to confirm or refute their tumor origin. CD45^─^ and CD45^+^ CECs were found in cancer patients (25.5 (range 0-404) and median 6.5 (0-147)) and the healthy donor (8 and 11 cells) and divided into three clusters. Two CD45^─^ CEC clusters were predominantly aneuploid (97% and 98%), but one cluster contained more diploid (59%) than aneuploid cells. CD45^+^ CECs were mostly diploid: only clusters 1 and 2 had aneuploid cells (16% and 2%). Diploid CD45^─^ and CD45^+^ CECs were annotated as different immune cells and surprisingly harbored many CNAs. Cancer-associated signaling pathways were found only in aneuploid cells of CD45^─^ CEC cluster 1 and diploid cells of CD45^+^ CEC cluster 1. Thus, our findings suggest that CECs in breast cancer patients are a highly heterogeneous population comprising aneuploid (tumor and hybrid) and diploid (normal) cells. DNA ploidy analysis is an effective instrument for identifying tumor and hybrid cells among CD45^─^ and CD45^+^ CECs, respectively.

## Introduction

Breast cancer (BC) is the most frequently diagnosed malignant neoplasm and the leading cause of cancer death in women ^1^. The majority of deaths from BC are related to distant metastases ^2^. Metastasis is a multistep process that includes cancer cell invasion, intravasation, circulation, extravasation, and the formation of micro- and macrometastases ^3^. Survival of circulating tumor cells (CTCs) in the bloodstream and their extravasation are critical steps of the metastatic cascade ^4^. CTCs are highly heterogeneous, and only a few of them can survive and form metastases ^5^. Many studies showed that breast CTCs possess specific stem and epithelial-mesenchymal transition (EMT) properties ^6-10^, while the molecular repertoire of CTCs varies from patient to patient ^6^. Deciphering CTC heterogeneity is thought to be a vital issue for identifying metastasis-initiating cells and therapeutic targets for the prevention of metastasis.

CTCs are usually identified as negative cells for CD45 leukocyte antigen and positive for epithelial markers such as epithelial cell adhesion molecule (EpCAM) and keratins (KRTs) ^10^. However, CD45-negative epithelial cells are also observed in the bloodstream of healthy subjects ^11,12^. Therefore, the detection of CTCs based on single epithelial and leukocyte markers is questioned. Different strategies for the enrichment, detection, and molecular characterization of CTCs have been proposed to address this problem in recent years ^13^; however, none of them can cover the complete diversity of CTCs.

Increasing evidence shows that metastases can also arise from tumor hybrid cells represented by fusions between tumor and normal cells ^14-16^. Tumor fusion cells with CD45 and epithelial markers demonstrate both increased metastatic potential and chemoresistance ^17,18^ and are frequently present in the blood of cancer patients ^19^. At present, the molecular landscape and composition of tumor hybrid cells remain unclear. Moreover, the detection of tumor hybrid cells is challenging. Most studies use parental cell markers (e.g., EpCAM/KRTs and CD45) to observe tumor fusion cells. However, atypical cells expressing epithelial and leukocyte markers are also present in healthy individuals ^20^.

Single-cell RNA sequencing (scRNA-seq) is an effective instrument for deciphering cell heterogeneity, identifying new cell populations, and analyzing their phenotypes. Previous scRNA-seq studies showed that breast CTCs coexpress epithelial and mesenchymal markers ^7^ and several specific transcripts ^21^ and can be in the state of estrogen responsiveness/increased proliferation or reduced proliferation/EMT ^22^ Tumor hybrid cells exhibit gene expression patterns distinct from parental cells but retain the expression of critical genes of each parental cell ^17^.

This study investigated the diversity of circulating epithelial cells (CECs) in patients with non-metastatic BC using 10x Genomics scRNA-seq. We focused mainly on CD45-negative CECs due to using a RosettSep negative selection by depleting white blood cells, red blood cells, and granulocytes. However, since depleted blood samples still contained white blood cells due to insufficiently good negative selection, we could also analyze CD45-positive CECs. Finally, an attempt was made to identify CTCs and tumor hybrid cells among CD45^⍰^ CECs and CD45^+^ CECs by analyzing DNA ploidy.

## Material and methods

### Cases

The study included 81 patients with non-metastatic invasive breast carcinoma of no special type (T^1-4^N^0-3^M^0^, all molecular subtypes) treated in the Cancer Research Institute, Tomsk National Research Medical Center from 2019 to 2021 and 1 cancer-free donor.

The 24.7% (20/81) of patients had CD45^⍰^EpCAM^+^/KRT7/8^+^ CTCs (8 cells per 1 ml of the blood) and were included in scRNA-seq. The clinical and pathological parameters of these patients are given in Supplementary Table 1. Supplementary Tables 2 and 3 demonstrates the number of CD45^⍰^and CD45^+^ CECs in patients. The proportion of aneuploid and diploid CD45^⍰^and CD45^+^ CECs among all cells after reclustering in each case is shown in Supplementary Tables 4 and 5.

The procedures followed in this study were in accordance with the Helsinki Declaration (1964, amended in 1975 and 1983). The study was approved by the Local Committee for Medical Ethics of the Cancer Research Institute, Tomsk NRMC (17 June 2016, the approval number is 8). All patients signed informed consent for voluntary participation.

### Blood collection and preparation

Venous blood samples (15 ml) were collected in EDTA tubes one to two days before surgical intervention and neoadjuvant chemotherapy and stored at room temperature (RT) for up to 4 hours.

CD45^⍰^cells were enriched using negative selection by depletion of CD45^+^, glycophorin A^+^, and CD66b^+^ cells (RosetteSep Human CD45 Depletion Cocktail, STEMCELL Technologies Inc., Canada). Blood samples were incubated with RosetteSep Cocktail (50 μl per 1 ml blood) at RT for 20 min, diluted at 1:1 ratio with 1X phosphate-buffered saline (PBS), and transferred equally into separate Falcon conical 15 ml tubes, containing lymphosep-1077 (Biowest, USA) density gradient media. After density gradient centrifugation (1200g for 20 min, break off), the peripheral blood mononuclear cell layer was isolated and washed RPMI-1640 medium with 5% fetal bovine serum (FBS) (300g for 10 min). Platelets were removed by two centrifugations at 110g for 10 min at RT with no brake. The final cell pellet was divided into two parts. One part was used to analyze CTCs by flow cytometry, another part was resuspended in 100 μl of RPMI-1640 medium with 20% FBS and 15% dimethylsulfoxide (DMSO), froze at −80^°^C, and stored not more than one month before scRNA-seq.

### Flow cytometry

CTCs were identified as CD45^⍰^EpCAM^+^/KRT7/8^+^ cells and detected by flow cytometry using the following antibodies and fluorescent dyes: anti-EpCAM-BV 650 (clone 9C4, mouse IgG2b, Sony Biotechnology, USA), anti-cytokeratin 7/8-AF647 (clone CAM5.2, Mouse IgG2a, BD Pharmingen, USA), anti-CD45-APC-Cy7 (clone HI30, mouse IgG1, Sony Biotechnology, USA), and 7[AAD (Sony Biotechnology, USA). For intracellular staining, cells were permeabilized using BD Cytofix/Cytoperm (BD Biosciences, San Jose, CA, USA). The isotype control antibodies at the same concentration were added to the control sample. MCF[7 and U937 cells were used as positive and negative controls, respectively. The phenotypic characteristics of CTCs were analyzed on a Novocyte 3000 (ACEA Biosciences, USA) (Supplementary Figure 1). Samples with more than 8 CTCs per ml of the blood were used in scRNA-seq.

### Single-cell RNA sequencing

Cell samples were thawed at 37^°^C for 2 min using a water bath. Then, 1 ml of warm RPMI-1640 medium with 10% FBS was added to cells. After centrifuging (300 g for 5 min), the supernatant was carefully collected, washed in 50 ul PBS with 0.04% BSA by pipetting with wide-bore pipette tips, and placed on the ice. Cell counting was carried out using 0.4% trypan blue (Thermo Fisher Scientific, USA) and the Goryaev chamber.

Single-cell cDNA libraries were prepared using the Single Cell 3′ Reagent Kit v3.1 and a 10x Genomics Chromium Controller. The number of cells in each channel of the Single-Cell Chip G varied from 3300 to 10000. The concentration of cDNA libraries was measured by the dsDNA High Sensitivity kit on a Qubit 4.0 fluorometer (Thermo Fisher Scientific, USA). The quality of cDNA libraries was assessed using High Sensitivity D1000 ScreenTape on a 4150 TapeStation (Agilent, USA). The ready cDNA libraries were pooled, denatured, and sequenced on NextSeq 500 and NextSeq 2000 (Illumina, USA) using pair-end reads (28 cycles for read 1, 91 cycles for read 2, and 8 cycles for i7 index).

### Bioinformatics and statistical analyzes

The Seurat software package version 4.0.4 ^23^ was used for quality control and analysis of the scRNA-seq data. The DoubletFinder ^24^ was used to detect cell doublets. The integrated data of 21 cases (20 patients and one donor) has been pre-processed by filtering cells with unique feature counts less than 200 and mitochondrial percent more than 25. Then, raw RNA UMI counts were normalized. The principal component analysis (PCA) was performed to reduce the dimensionality. The uniform manifold approximation and projection (UMAP) method, a non-linear dimensionality reduction technique, was used to visualize and explore the dataset.

CD45^−^ CECs were identified as cells with no *PTPRC* (CD45) gene and expression level of epithelial genes (*EPCAM, CDH1, KRT5, KRT7, KRT8, KRT17, KRT18*, and *KRT19*) more than 0. CD45^+^ CECs were detected as cells with the expression level of the *PTPRC, EPCAM, CDH1, KRT5, KRT7, KRT8, KRT17, KRT18*, and *KRT19* genes more than 0.

DEGs were identified by comparing each aneuploid and diploid cluster of CD45^⍰^and CD45^+^ CECs to the reference NK cells detected as *IL7R* and *KLRB1* positive cells based in the scRNA-seq data. The average fold change, percentage of expression in each specific cluster, percentage of expression in other clusters, and adjusted p-value were calculated. TOP10 upregulated specific DEGs were visualized using volcano plots.

DEGs with adjusted p < 0.05 were used for functional enrichment analysis by Enrichr ^25^. TOP5 specific signaling pathways together with cancer-associated signaling pathways (Rap1, PI3K-Akt, etc.), if presented, were given in figures and discussed.

CNAs and DNA ploidy were analyzed using inferCNV (https://github.com/broadinstitute/infercnv) and CopyKAT ^26^ instruments, respectively. The identification of cell types was performed by the SingleR package using Human Primary Cell Atlas as a reference dataset ^27^. Raw gene expression data of CD45^⍰^and CD45^+^ CECs were extracted using the Seurat. The reference was selected as cells with expression of *IL7R* and *KLRB1* and no expression of *KRT5, KRT7, KRT8, KRT17, KRT18, KRT19, EPCAM*, and *CDH1*. The inferCNV analysis included the following parameters: “denoise”, default Hidden Markov Model (HMM) settings, and a value of 0.1 for “cutoff”. Each CNA was annotated in a six-state model: complete loss, loss of one copy, neutral, the addition of one copy, and the addition of two copies and more than two copies. The following parameters were used in CopyKAT analysis: the minimal number of genes per chromosome for cell filtering – 5, minimal window sizes for segmentation – 25 genes per segment, segmentation parameter – 0.05, and the number of cores for parallel computing – 4. Euclidean distance was chosen as distance parameters for clustering to favor data with larger copy-number segments.

Statistical analysis was performed using IBM SPSS Statistics for Windows, version 23.0 (IBM Corp., USA). None of the quantitative variables in the comparison groups had a normal distribution according to the Kolmogorov-Smirnov test, and non-parametric tests were used. Mann-Whitney test was used to assess the correlation between the number of CECs and estrogen receptor status. Kruskal-Wallis test was used to determine the association of the number of CECs with molecular subtype, tumor size, tumor grade, and lymph node metastasis. In case of significant difference, pair-wise comparison between groups was done by Mann-Whitney test. Spearman’s linear correlation analysis was used to assess the relationship between the number of CECs and Ki-67 expression. Medians with the first and third quartiles were calculated for the variables in the comparison groups. P values < 0.05 were considered statistically significant.

## Results

### Detection of CECs

The following protocol was used to analyze CECs: (1) enrollment of patients with invasive breast carcinoma of no special type, (2) collection of peripheral blood samples and depletion of leukocytes (CD45), erythrocytes (glycophorin A), and granulocytes (CD66b), (3) flow cytometry analysis and selection of cases with CD45^⍰^EpCAM^+^/KRT7/8^+^ cells, and (4) scRNA-seq.

A total of 81 patients with non-metastatic BC were included. The study also included one cancer-free donor as a control. Flow cytometry analysis showed CD45^⍰^EpCAM^+^/KRT7/8^+^ cells only in 20 patients. Blood samples of 20 patients and one donor were used to enrich epithelial cells and to perform scRNA-seq.

In total, scRNA-seq identified 42225 cells. Data of 20 patients and one donor were consolidated into one file. The reclustering workflow was initiated with the corresponding parameters (threshold by UMIs and features and mitochondrial UMIs), resulting in 67.5% of removed barcodes. The remaining 13741 cells were divided into 16 clusters (Figure 1a).

**Fig. 1.**
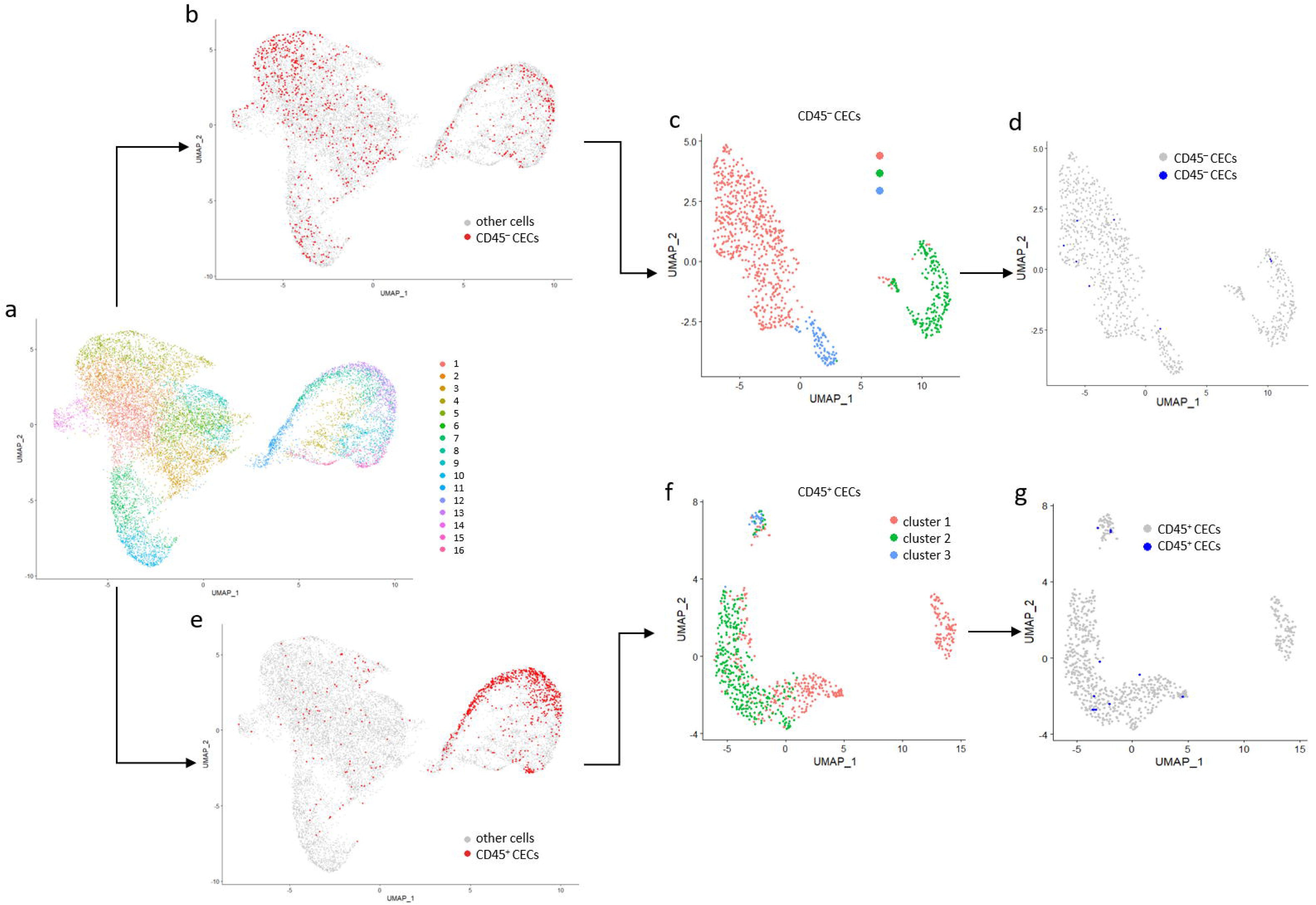
ScRNA-seq UMAP plots of blood samples of BC patients after depleting whole blood cells, red blood cells, and granulocytes. **a** Identified cell clusters. **b, c** CD45^⍰^ CECs and their clusters. **d** CD45^⍰^CECs identified in BC patients (gray color) and the healthy donor (blue color). **e, f** CD45^+^ CECs and their clusters. **g** CD45^+^ CECs identified in BC patients (gray color) and the healthy donor (blue color).

CD45^⍰^CECs were identified based on the expression of one or more epithelial markers: KRT 5, 7, 8, 17, 18, 19, EpCAM, and CDH1 (E-cadherin) (Figure 1b). The number of CD45^⍰^CECs varied from 0 to 404 in patients (median 25.5, Table 1). CD45^⍰^CECs were represented by 923 cells consisting of 3 clusters with 632, 203, and 88 cells, respectively (Figure 1c).

**Table 1.**
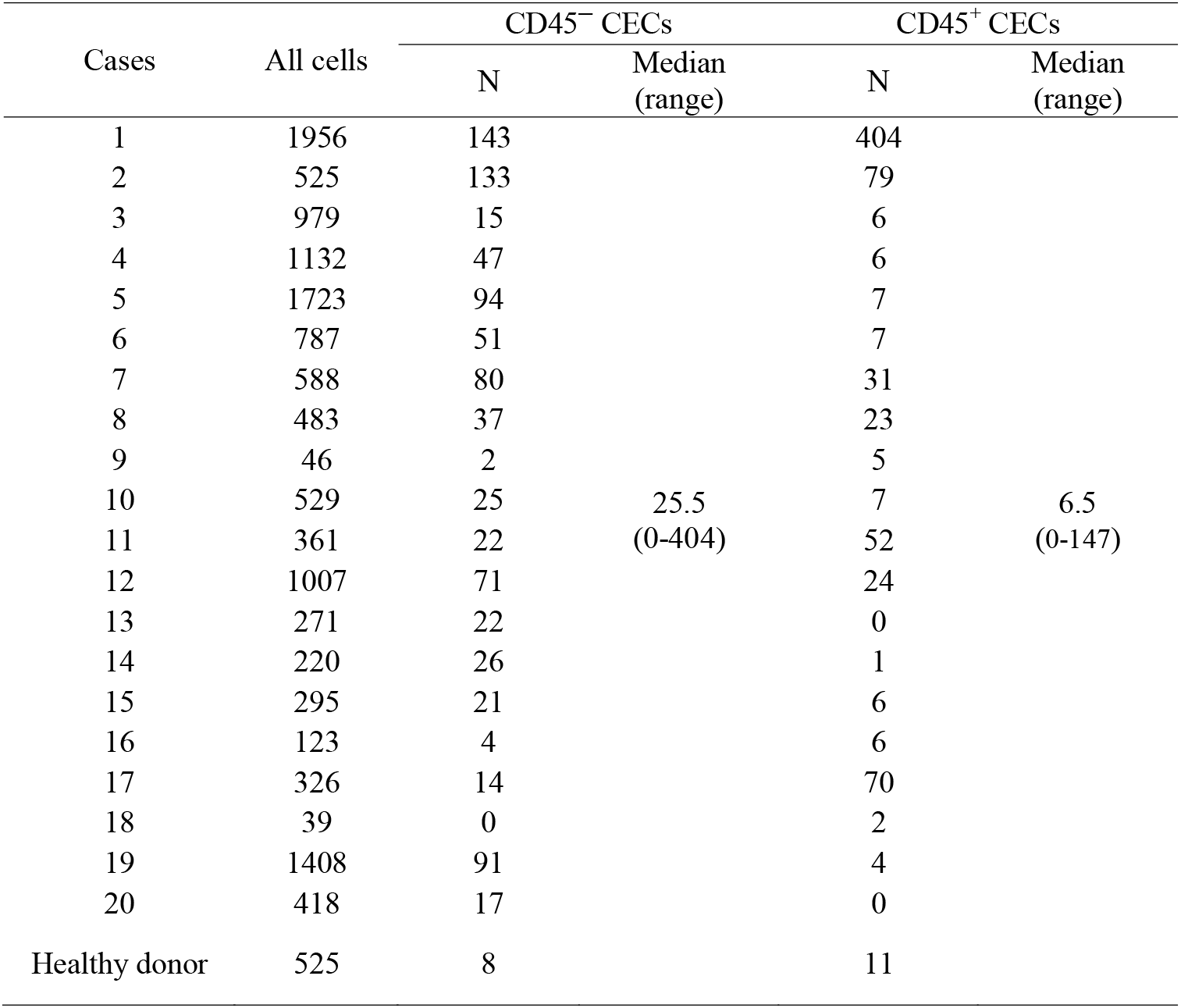
Number of cells detected after scRNA-seq

Unexpectedly, we found many CD45^+^ cells (n=2727), among which 751 cells had epithelial markers (KRT 5, 7, 8, 17, 18, 19, EpCAM, and CDH1) (Figure 1e). It may be due to the incomplete removal of CD45^+^ cells by the RosetteSep enrichment method ^28^. However, since this method non-selectively depletes CD45^+^ cells, the population composition of these cells probably remains unchanged. The number of CD45^+^ CECs ranged from 0 to 147 in patients (median 6.5, Table 1). After reclustering, CD45^+^ CECs were divided into 3 clusters with 361, 360, and 30 cells, respectively (Figure 1f).

CECs were also detected in the healthy donor. CD45^⍰^CECs were found in all three clusters with 5, 2, and 1 cells, respectively (Figure 1d). CD45^+^ CECs were also observed in three clusters and represented by 1, 9, and 1 cells, respectively (Figure 1g).

### Analysis of DNA ploidy in CD45-negative and CD45-positive CECs

Aneuploidy is a hallmark of cancer and is high in breast cancers ^29^. Here, we analyzed DNA ploidy to determine how many tumor cells can be presented in CD45^⍰^CEC clusters. Aneuploidy was primarily identified in CD45^⍰^CEC clusters 1 (97%, 579/597 cells) and 3 (98%, 61/62). Cluster 2 of CD45^⍰^CECs contained more diploid (59%, 115/195) than aneuploid cells (Figure 2a). No aneuploidy was detected in cluster 3 of CD45^+^ CECs, whereas clusters 1 and 2 contained only 16% (57/360) and 2% (7/351) aneuploid cells, respectively (Figure 2b). The number of diploid and aneuploid cells in each cluster is given in Supplementary Table 6. Note that several CD45^⍰^and CD45^+^ CECs were not assigned by the CopyKAT instrument as aneuploid and diploid that explain differences in the number of cells in clusters here and above.

**Fig. 2.**
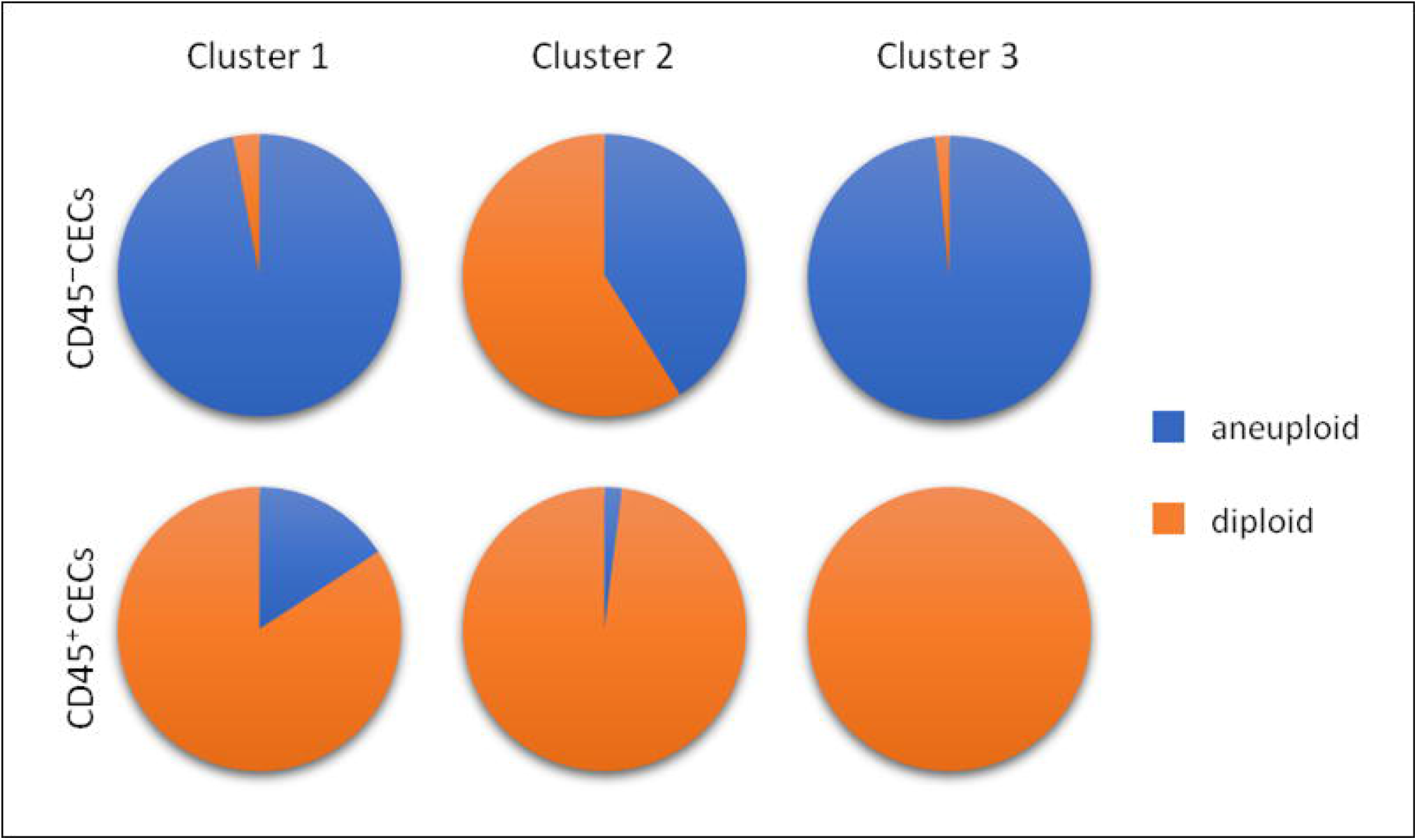
Percentage of aneuploid and diploid cells in CD45^⍰^ and CD45^+^ CEC clusters.

### The transcriptional landscape of aneuploid CD45-negative CECs

The TOP10 genes upregulated in aneuploid cells of CD45^⍰^CEC clusters are shown in Figure 3a-c. Aneuploid cells of CD45^⍰^CEC cluster 1 demonstrated overexpression of platelet genes (*PF4* and *GP1BB*), and several other genes, including *NRGN* (neurogranin) and *GNG11* (G protein subunit gamma 11) (Figure 3a). Aneuploid cells of cluster 2 had upregulation of the *KRT5* (keratin 5), *HLA-DRA* (major histocompatibility complex, class II, DR alpha), and *BCL11A* (BAF chromatin remodeling complex subunit) genes (Figure 3b). In cluster 3, aneuploid cells overexpressed different hemoglobin genes, as well as *CA1* (carbonic anhydrase 1) and *YBX3* (Y-box binding protein 3) genes (Figure 3c). The complete list of genes expressed in aneuploid CD45^⍰^CECs is shown in Supplementary Tables 7 and 8.

**Fig. 3.**
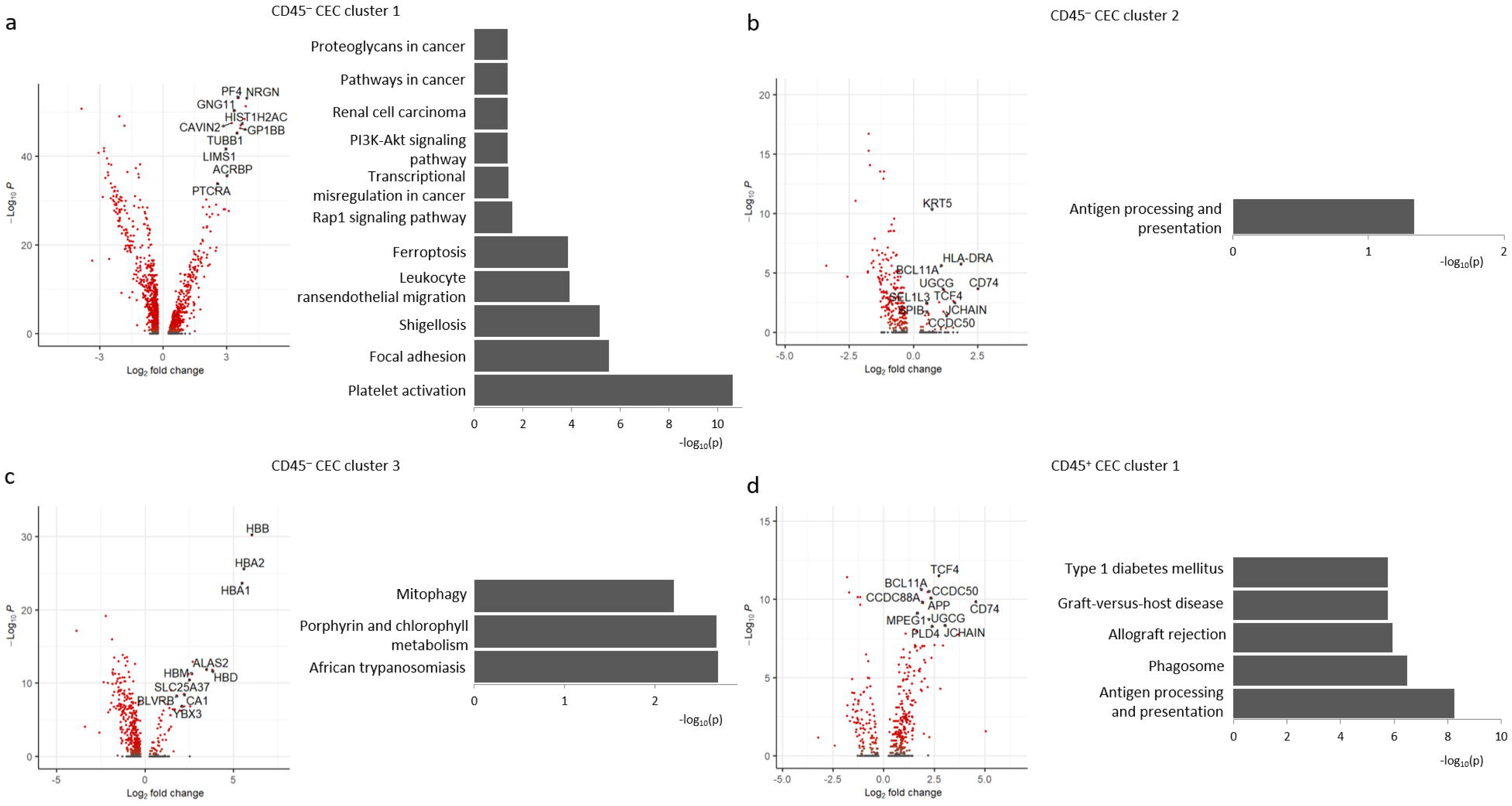
Differential gene expression analysis of aneuploid cells in CD45^⍰^ and CD45^+^ CEC clusters. **a, b, c** CD45^⍰^ CECs. **d** CD45^+^ CECs. Left panel – TOP10 specific DEGs (red color – adjusted p<0.05), right panel – TOP5 specific signaling pathways together with cancer-associated pathways.

Differentially expressed genes (DEGs) were used to reveal the enrichment of specific signaling pathways in aneuploid CD45^⍰^CECs (Figure 3a-c). In cluster 1, aneuploid cells had 33 signaling pathways that are mainly associated with platelet activation, leukocyte transendothelial migration, ferroptosis, and cancer (Rap1 signaling pathway, transcriptional misregulation in cancer, PI3K-Akt signaling pathway, renal cell carcinoma, pathways in cancer, and proteoglycans in cancer; Figure 3a). Aneuploid cells of cluster 2 had only one specific signaling pathway – antigen processing and presentation (Figure 3b). In cluster 3, aneuploid cells demonstrated three specific signaling pathways that are associated with African trypanosomiasis, porphyrin and chlorophyll metabolism, and mitophagy (Figure 3c). The complete list of signaling pathways enriched in aneuploid CD45^⍰^CECs is shown in Supplementary Tables 9 and 10.

### The transcriptional landscape of aneuploid CD45-positive CECs

Aneuploid cells of CD45^+^ CEC cluster 1 showed overexpression of the *TCF4* (transcription factor 4), *BCL11A*, and *CCDC50* (Coiled-coil domain containing 50) genes (Figure 3d). A total of 33 specific signaling pathways were identified in aneuploid cells of CD45^+^ CEC cluster 1 to be related mainly to antigen processing and presentation, phagosome, and allograft rejection (Figure 3d). The complete list of genes and signaling pathways enriched in aneuploid cells of CD45^+^ CEC cluster 1 is given in Supplementary Tables 11 and 12. In cluster 2, aneuploid cells had no significant DEGs, and cluster 3 did not contain aneuploid cells.

### Cellular composition of diploid CD45-negative and CD45-positive CECs

Diploid cells identified in CD45^⍰^and CD45^+^ CEC clusters may be non-tumor cells harboring epithelial features. In CD45^⍰^CECs, the SingleR instrument showed that diploid cells (3%, 18/597) of cluster 1 are predominantly represented by platelets, neutrophils, and B cells cluster 2 (59%, 115/195) – by T and NK cells, and cluster 3 (2%, 1/62) – by bone marrow-derived mononuclear cells (Figure 4a). In CD45^+^ CECs, diploid cells of cluster 1 (84%, 303/360) predominantly consisted of NK, T, and B cells as well as platelets, cluster 2 (98%, 344/351) was mainly represented by T and NK cells, and cluster 3 (100%, 29/29) – by stem and T cells (Figure 4a).

**Fig. 4.**
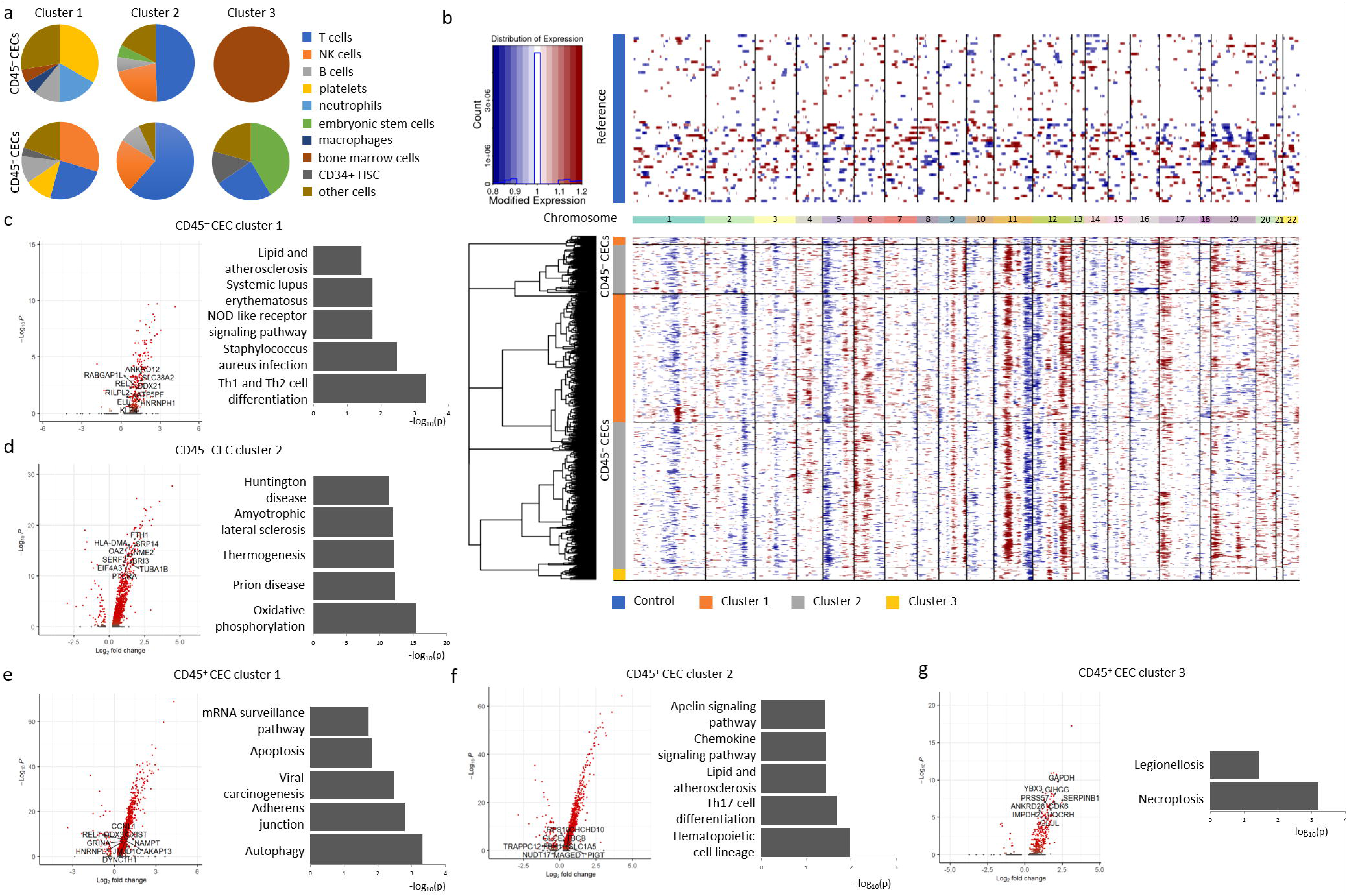
The molecular landscape of diploid cells identified in CD45^⍰^ and CD45^+^ CEC clusters. **a** CNA profile of diploid cells. **b** Prediction of cell types among diploid cells. **c, d** TOP10 specific DEGs (red color – adjusted p<0.05) and TOP5 specific signaling pathways in diploid cells of CD45^+^ CEC clusters. **e, f, g** TOP10 specific DEGs (red color – adjusted p<0.05) and TOP5 specific signaling pathways in diploid cells of CD45^⍰^ CEC clusters.

Also, we analyzed copy-number aberrations (CNAs) to exclude the presence of diploid tumor cells in CD45^⍰^and CD45^+^ CECs. Most diploid cells of CD45^⍰^CEC clusters 1 and 2 harbored gains in 12q23.3-12q24.33 and 17p13.2-17q11.2 and losses at 5p13.3-5q13.1 and 11q21-11q25 regions (Figure 4b, Supplementary Table 13). Cluster 3 contained only one diploid cell that was not enough to conduct CNA analysis. CNAs were also identified in diploid CD45^+^ CECs and represented by losses at 5p15.2-5q11.2 and 11q23.1-11q25 (Figure 4b, Supplementary Table 13). Additionally, most diploid cells of CD45^+^ CEC clusters 1 and 2 demonstrated losses in 11q21-11q25 and 12p11.23-12p13.31 and gains in 11p11.2-11q13.1 and 12q23.2-12q24.33

(Figure 4b, Supplementary Table 13). These findings were surprising given that SingleR analysis identified no epithelial (tumor) cells that could harbor CNAs. Moreover, it was strange that not a single cell was found to be without CNAs. However, CD45^⍰^CECs and CD45^+^ CECs are known to be present in normal physiological conditions ^11,12,20^ and most likely negative for chromosome aberrations.

### The transcriptional landscape of diploid CD45-negative CECs

DEGs were found only in diploid cells of CD45^⍰^CEC clusters 1 and 2. The *ANKRD12* (ankyrin repeat domain 12), *SLC38A2* (solute carrier family 38 member 2), and *RABGAP1L* (RAB GTPase activating protein 1 like) were among the TOP10 genes overexpressed in diploid cells of cluster 1 (Figure 4c). In cluster 2, diploid cells overexpressed the *FTH1* (ferritin heavy chain 1), *HLA-DMA* (major histocompatibility complex, class II, DM alpha), *SRP14* (signal recognition particle 14), and several other genes (Figure 4d). The complete list of genes expressed in diploid CD45^⍰^CECs is shown in Supplementary Tables 14 and 15.

Diploid cells of CD45^⍰^CEC cluster 1 had five specific signaling pathways that are mainly associated with Th1 and Th2 cell differentiation, Staphylococcus aureus infection, and NOD-like receptors (Figure 4c). In cluster 2, diploid cells demonstrated 28 specific signaling pathways that are involved mainly in oxidative phosphorylation, prion disease, and thermogenesis (Figure 4d). The complete list of signaling pathways enriched in diploid CD45^⍰^CEC clusters is given in Supplementary Tables 16 and 17.

### The transcriptional landscape of diploid CD45-positive CECs

The TOP10 genes upregulated in diploid cells of CD45^+^CEC clusters are shown in Figure 4e-g. Cluster 1 of diploid CD45^+^ CECs overexpressed the *CCNL1* (cyclin L1), *NAMPT* (nicotinamide phosphoribosyltransferase), *DDX3X* (DEAD-box helicase 3 X-linked), and other genes (Figure 4E). Diploid cells of cluster 2 demonstrated upregulation of the *RPS10* (ribosomal protein S10), *CHCHD10* (coiled-coil-helix-coiled-coil-helix domain containing 10), *TBCB* (tubulin folding cofactor B) and other genes (Figure 4f). In cluster 3, diploid cells had overexpression of the *GAPDH* (glyceraldehyde-3-phosphate dehydrogenase), *YBX3* (Y-box binding protein 3), and *GIHCG* (miR-200b/200a/429 inhibitor) and other genes (Figure 4g). The complete list of genes expressed in diploid CD45^⍰^CECs is shown in Supplementary Tables 18 and 19.

Cluster 1 of diploid CD45^+^ CECs had ten specific signaling pathways that are mainly related to autophagy, adherens junction, and viral carcinogenesis (Figure 4e). Cluster 2 showed seven specific signaling pathways that are associated with hematopoietic cell lineage, Th17 cell differentiation, and lipid and atherosclerosis (Figure 4f). Cluster 3 demonstrated only two specific signaling pathways that are involved in necroptosis and legionellosis (Figure 4g). The complete list of signaling pathways enriched in diploid CD45^+^ CEC clusters is given in Supplementary Tables 20 and 21.

### Association between the number of CECs and clinicopathological features of BC

The number of aneuploid and diploid CD45^⍰^CECs and CD45^+^ CECs identified in each cluster was correlated to molecular subtype, tumor size, grade, lymph node metastasis, estrogen receptor status, and Ki-67 expression. Significant differences were found only for the association of CECs with tumor grade. The number of CD45^+^ CECs, particularly diploid cells of clusters 1 and 2, was higher in grade 3 than in grades 1 and 2 (p<0.05; Supplementary Table 22).

## Discussion

Epithelial cells and atypical cells with epithelial and leukocyte features are observed in the blood of not only cancer patients where they are described as CTCs and tumor hybrid cells, respectively, but also in healthy subjects ^11,12,20^. These findings suggest that not all CECs detected in cancer patients can be CTCs and tumor hybrid cells.

This study first provides a thorough analysis of the diversity of CECs in patients with non-metastatic BC using scRNA-seq. Due to the depletion of CD45-positive cells in blood samples, we focused on CD45-negative CECs. Since depleted blood samples still contained white blood cells, we could also analyze CD45-positive CECs. In CD45^⍰^CECs and CD45^+^ CECs, we attempted to identify CTCs and tumor hybrid cells based on DNA ploidy analysis because aneuploidy is known to be high in breast tumors ^29^.

CD45^⍰^CECs consisted from three clusters. Clusters 1 and 2 were predominantly aneuploid, whereas cluster 3 contained more diploid than aneuploid cells. Aneuploid cells of cluster 1 had the most cancer signaling pathways and demonstrated platelet genes and platelet activation signaling. These findings can indicate an increased potential of aneuploid cells to adhere with thrombocytes known to protect CTCs from the immune system and increase their survival in the circulation ^30^. Aneuploid cells of clusters 2 and 3 had no cancer signaling pathways. Nevertheless, aneuploid cells of cluster 2 overexpressed the *BCL11A* gene that is a potential driver of the development and progression of triple-negative breast cancer ^31^. In cluster 3, aneuploid cells showed hemoglobin genes that may indicate their increased potential to adhesion with red blood cells and upregulation of the *CA1* and *YBX3* genes associated with tumorigenesis and progression in BC ^32,33^.

CD45^+^ CECs included three clusters, two of which contained more diploid than aneuploid cells, whereas the third cluster was completely diploid. No cancer signaling pathways were found in aneuploid CD45^+^ CECs. Only one cluster had enough aneuploid cells to identify antigen processing and presentation signaling pathway. These cells also overexpressed the *BCL11A* gene like aneuploid cells of CD45^⍰^CEC cluster 2. In addition, upregulation of the *TCF4* gene related to the enhancement of BC invasion ^34^ was found in aneuploid CD45^+^ CECs.

Given these results, we suggested that aneuploid cells identified in CD45^⍰^and CD45^+^ CEC clusters can represent different populations of CTCs and tumor hybrid cells, respectively. Only one population of aneuploid CD45^⍰^CECs demonstrated the most transcriptional cancer features that may indicate the presence of an aggressive subset of CTCs. Interestingly, no important molecular features were identified in aneuploid CD45^+^ CECs; however, tumor hybrid cells are known to have increased proliferation and migration, drug resistance, decreased apoptosis rate, and avoiding immune surveillance ^14,15,35^.

Diploid CD45^⍰^ and CD45^+^CECs included different normal cells (stem, bone marrow-derived mononuclear, B, T, and NK cells) and were enriched by Th1 and Th2 cell differentiation, oxidative phosphorylation hematopoietic cell lineage, necroptosis, and other signaling pathways. Surprisingly, diploid cells of CD45^+^ CEC cluster 1 had the viral carcinogenesis signaling pathway and overexpressed the *NAMPT* gene that promotes progression and metastasis of triple-negative breast cancer^36^. These findings can indicate the tumor nature of diploid cells of CD45^+^ CEC cluster 1 because not all breast cancer cells are aneuploid ^29^. We attempted to prove this suggestion by the analysis of CNAs based on the scRNA-seq data. However, the obtained results were surprising and puzzling because all diploid CD45^⍰^and CD45^+^CECs harbored CNAs. Probably, the scRNA-seq data is not the most appropriate resource to assess CNAs.

Some previous scRNA-seq studies identified CTCs and differentiated them from normal peripheral blood mononuclear cells but only in metastatic BC patients ^21,22^. It is well known that CTCs are significantly higher in advanced than in early cancers ^37^. However, these studies lack possibility to follow up patients with non-metastatic BC and identify CTCs which can be metastasis-initiating cells. In addition, CTCs were identified in these studies based only on positive epithelial and negative CD45 markers, whereas epithelial cells are also observed in the blood of healthy individuals as shown here and in other studies ^11,12^.

Overall, the findings reported here indicate that CD45-negative and CD45-positive CECs are highly heterogeneous in BC patients and represented by transcriptionally-distinct populations that include both aneuploid cells, most likely represented by CTCs and tumor hybrid cells, and normal diploid cells. The analysis of DNA ploidy appears to be an effective instrument for discriminating tumor and non-tumor cells among CECs.

The study has several limitations. The significant differences in the number of loaded cells and cells calculated after scRNA-seq indicate that many cells are lost and thus not analyzed. It is probably related both to the not absolute capture of cells in gel beads at the step of RNA library preparation and low cell viability after negative selection. In addition, the results of transcriptional profiling of CD45^+^ CECs should be interpreted with caution because negative selection used for the enrichment of epithelial cells removes most of the CD45-positive cells. And finally, our study does not focus on the heterogeneity of circulating mesenchymal cells and the identification of CTCs and tumor hybrid cells that have undergone epithelial-mesenchymal transition and lost epithelial features. Hypothetically, these cells can be detected using the current scRNA-seq data.

## Author contributions

Conception and design: M.E.M., E.V.D. and V.M.P. Enrolling of patients and collection of biological samples: N.A.T. and N.O.P. Collection and assembly of data: M.E.M., T.S.G., V.V.A., O.E.S. and E.S.G. Data analysis and interpretation: M.E.M., V.R.Z., L.A.T., S.Yu.Z.,

A.A.K. and E.V.D. Paper writing: M.E.M., V.R.Z., E.V.D. and L.A.T. Final approval of paper: M.E.M., E.V.D., E.L.C., N.V.C.

## Acknowledgments

We thank Konstantin Zaytsev, Konstantin Okonechnikov, and Mikhail Arbatsky for valuable comments in the bioinformatic processing of scRNA-seq data. Sequencing was carried out on The Core Facility “Medical Genomics” (Tomsk NRMC) and The Tomsk Regional Common Use Center.

## Data availability

The scRNA-seq data generated for this study are available via BioProject under the accession number PRJNA776403.

## References

1. Dyba T, Randi G, Bray F, et al. The European cancer burden in 2020: Incidence and mortality estimates for 40 countries and 25 major cancers. Eur J Cancer. 2021;157:308–347.

2. Dillekås H, Rogers MS, Straume O. Are 90% of deaths from cancer caused by metastases? Cancer medicine. 2019;8(12):5574–5576.

3. Valastyan S, Weinberg RA. Tumor metastasis: molecular insights and evolving paradigms. Cell. 2011;147(2):275–292.

4. Tayoun T, Faugeroux V, Oulhen M, Aberlenc A, Pawlikowska P, Farace F. CTC-Derived Models: A Window into the Seeding Capacity of Circulating Tumor Cells (CTCs). Cells. 2019;8(10).

5. Massagué J, Obenauf AC. Metastatic colonization by circulating tumour cells. Nature. 2016;529(7586):298–306.

6. Savelieva OE, Tashireva LA, Kaigorodova EV, et al. Heterogeneity of Stemlike Circulating Tumor Cells in Invasive Breast Cancer. Int J Mol Sci. 2020;21(8).

7. Cheng Y-H, Chen Y-C, Lin E, et al. Hydro-Seq enables contamination-free high-throughput single-cell RNA-sequencing for circulating tumor cells. Nature Communications. 2019;10(1):2163.

8. Tashireva LA, Savelieva OE, Grigoryeva ES, et al. Heterogeneous Manifestations of Epithelial-Mesenchymal Plasticity of Circulating Tumor Cells in Breast Cancer Patients. Int J Mol Sci. 2021;22(5).

9. Aktas B, Tewes M, Fehm T, Hauch S, Kimmig R, Kasimir-Bauer S. Stem cell and epithelial-mesenchymal transition markers are frequently overexpressed in circulating tumor cells of metastatic breast cancer patients. Breast Cancer Res. 2009;11(4):R46.

10. Yu M, Bardia A, Wittner BS, et al. Circulating breast tumor cells exhibit dynamic changes in epithelial and mesenchymal composition. Science. 2013;339(6119):580–584.

11. Molnar B, Ladanyi A, Tanko L, Sréter L, Tulassay Z. Circulating Tumor Cell Clusters in the Peripheral Blood of Colorectal Cancer Patients. Clinical Cancer Research. 2001;7(12):4080–4085.

12. Allard WJ, Matera J, Miller MC, et al. Tumor Cells Circulate in the Peripheral Blood of All Major Carcinomas but not in Healthy Subjects or Patients With Nonmalignant Diseases. Clin Cancer Res. 2004;10(20):6897–6904.

13. Bankó P, Lee SY, Nagygyörgy V, et al. Technologies for circulating tumor cell separation from whole blood. J Hematol Oncol. 2019;12(1):48.

14. Gast CE, Silk AD, Zarour L, et al. Cell fusion potentiates tumor heterogeneity and reveals circulating hybrid cells that correlate with stage and survival. Science advances. 2018;4(9):eaat7828.

15. Dietz MS, Sutton TL, Walker BS, et al. Relevance of circulating hybrid cells as a non-invasive biomarker for myriad solid tumors. Sci Rep. 2021;11(1):13630.

16. Tretyakova MS, Subbalakshmi AR, Menyailo ME, Jolly MK, Denisov EV. Tumor Hybrid Cells: Nature and Biological Significance. Frontiers in cell and developmental biology. 2022;10:814714.

17. Chitwood CA, Dietzsch C, Jacobs G, et al. Breast tumor cell hybrids form spontaneously in vivo and contribute to breast tumor metastases. APL bioengineering. 2018;2(3):031907.

18. Carloni V, Mazzocca A, Mello T, Galli A, Capaccioli S. Cell fusion promotes chemoresistance in metastatic colon carcinoma. Oncogene. 2013;32(21):2649–2660.

19. Aguirre LA, Montalbán-Hernández K, Avendaño-Ortiz J, et al. Tumor stem cells fuse with monocytes to form highly invasive tumor-hybrid cells. Oncoimmunology. 2020;9(1):1773204.

20. Lustberg MB, Balasubramanian P, Miller B, et al. Heterogeneous atypical cell populations are present in blood of metastatic breast cancer patients. Breast Cancer Res. 2014;16(2):R23.

21. Pauken CM, Kenney SR, Brayer KJ, Guo Y, Brown-Glaberman UA, Marchetti D. Heterogeneity of Circulating Tumor Cell Neoplastic Subpopulations Outlined by Single-Cell Transcriptomics. Cancers. 2021;13(19).

22. Brechbuhl HM, Vinod[Paul K, Gillen AE, et al. Analysis of circulating breast cancer cell heterogeneity and interactions with peripheral blood mononuclear cells. Mol Carcinog. 2020;59(10):1129–1139.

23. Hao Y, Hao S, Andersen-Nissen E, et al. Integrated analysis of multimodal single-cell data. Cell. 2021.

24. McGinnis CS, Murrow LM, Gartner ZJ. DoubletFinder: Doublet Detection in Single-Cell RNA Sequencing Data Using Artificial Nearest Neighbors. Cell Systems. 2019;8(4):329-337.e324.

25. Xie Z, Bailey A, Kuleshov MV, et al. Gene Set Knowledge Discovery with Enrichr. Current Protocols. 2021;1(3):e90.

26. Gao R, Bai S, Henderson YC, et al. Delineating copy number and clonal substructure in human tumors from single-cell transcriptomes. Nat Biotechnol. 2021;39(5):599–608.

27. Aran D, Looney AP, Liu L, et al. Reference-based analysis of lung single-cell sequencing reveals a transitional profibrotic macrophage. Nat Immunol. 2019;20(2):163–172.

28. Maertens Y, Humberg V, Erlmeier F, et al. Comparison of isolation platforms for detection of circulating renal cell carcinoma cells. Oncotarget. 2017;8(50):87710–87717.

29. Pfister K, Pipka JL, Chiang C, et al. Identification of Drivers of Aneuploidy in Breast Tumors. Cell reports. 2018;23(9):2758–2769.

30. Palumbo JS, Talmage KE, Massari JV, et al. Platelets and fibrin(ogen) increase metastatic potential by impeding natural killer cell-mediated elimination of tumor cells. Blood. 2005;105(1):178–185.

31. Khaled WT, Choon Lee S, Stingl J, et al. BCL11A is a triple-negative breast cancer gene with critical functions in stem and progenitor cells. Nature Communications. 2015;6(1):5987.

32. Zheng Y, Xu B, Zhao Y, et al. CA1 contributes to microcalcification and tumourigenesis in breast cancer. BMC Cancer. 2015;15(1):679.

33. Qin H-L, Wang X-J, Yang B-X, Du B, Yun X-L. Notoginsenoside R1 attenuates breast cancer progression by targeting CCND2 and YBX3. Chinese Medical Journal. 2021;134(5):546.

34. Ravindranath A, Yuen HF, Chan KK, et al. Wnt-β-catenin-Tcf-4 signalling-modulated invasiveness is dependent on osteopontin expression in breast cancer. British journal of cancer. 2011;105(4):542–551.

35. Tretyakova MS, Subbalakshmi AR, Menyailo ME, Jolly MK, Denisov E. Tumor Hybrid Cells: Nature and Biological Significance. 2021.

36. Zhang H, Zhang N, Liu Y, et al. Epigenetic regulation of NAMPT by NAMPT-AS drives metastatic progression in triple-negative breast cancer. Cancer research. 2019;79(13):3347–3359.

37. Jin L, Zhao W, Zhang J, et al. Evaluation of the diagnostic value of circulating tumor cells with CytoSorter(®) CTC capture system in patients with breast cancer. Cancer medicine. 2020;9(5):1638–1647.

