## Supplementary Table 1 for "Heterogeneity of circulating epithelial cells in breast cancer at single-cell resolution: identifying tumor and hybrid cells"

Supplementary Table 1. Clinicopathological characteristics of breast cancer patients

| Cases | Age at diagnosis (yr) | Grade | TNM | Focality | Molecular subtype | ER | PR | HER2 | Ki-67 (%) | Neoadjuvant chemotherapy |
|---|---|---|---|---|---|---|---|---|---|---|
| 1 | 37 | 3 | T2N2M0 | UF | TN | - | + | 0 | 80 | Yes |
| 2 | 50 | 3 | T2N0M0 | MF | Lum B HER2- | + | + | 1+ | 38 | No |
| 3 | 53 | 2 | T1N0M0 | MF | TN | - | - | 0 | 87 | No |
| 4 | 64 | 2 | T2N1M0 | UF | Lum B HER2- | + | + | 1+ | 30 | No |
| 5 | 64 | 2 | T2N0M0 | UF | Lum B HER2+ | + | + | 3+ | 21 | Yes |
| 6 | 56 | 1 | T4N3M0 | UF | Lum B HER2- | + | - | 1+ | 42 | No |
| 7 | 44 | 2 | T2N1M0 | UF | Lum B HER2- | + | + | 0 | 20 | No |
| 8 | 58 | 3 | T4N1M0 | UF | TN | - | - | 2+ (FISH-) | 52 | Yes |
| 9 | 48 | 3 | T2N2M0 | UF | TN | - | - | 0 | 36 | Yes |
| 10 | 58 | 2 | T2N1M0 | UF | Lum B HER2- | + | + | 1+ | 43 | No |
| 11 | 71 | 3 | T2N1M0 | UF | Lum B HER2- | + | 0 | 0 | 85 | No |
| 12 | 76 | 1 | T2NxM0 | UF | Lum A HER2- | + | + | 1+ | 7 | No |
| 13 | 45 | 2 | T2N0M0 | UF | Lum A HER2- | + | + | 1+ | 14 | No |
| 14 | 60 | 2 | T3N1M0 | UF | Lum B HER2- | + | + | 0 | 32 | ND |
| 15 | 62 | 1 | T2N1M0 | UF | Lum B HER2- | + | + | 0 | 44 | Yes |
| 16 | 36 | 2 | T2NxM0 | MF | Lum B HER2- | + | + | 0 | 35 | Yes |
| 17 | 45 | 2 | T2N0M0 | UF | TN | - | - | 0 | 77 | Yes |
| 18 | 68 | 3 | T2NxMx | UF | Lum B HER2- | + | 0 | - | 33 | Yes |
| 19 | 60 | 1 | T2N0M0 | UF | Lum B HER2- | + | + | 2+ (FISH-) | 38 | No |
| 20 | 55 | 2 | T4N1M0 | UF | HER2+ | - | - | 3+ | 18 | Yes |

UF, unifocal; MF, multifocal; Lum, luminal; TN, triple-negative; yr, years; ER, estrogen receptors; PR, progesterone receptors; TNM, tumor-node-metastasis classification; "+", presence; "–", absence; FISH, fluorescence *in situ* hybridization
