## Supplementary Table 2 for "Heterogeneity of circulating epithelial cells in breast cancer at single-cell resolution: identifying tumor and hybrid cells"

Supplementary Table 2. Number of CD45$^{-}$ CECs in breast cancer patients

| Cases | total number of cells | CD45$^{-}$ CECs | | | | | | | | | |
|---|---|---|---|---|---|---|---|---|---|---|---|
| | | total CD45$^{-}$ CECs | Cluster 1 | | | Cluster 2 | | | Cluster 3 | | |
| | | | total | Aneuploid | Diploid | total | Aneuploid | Diploid | total | Aneuploid | Diploid |
| 1 | 1956 | 143 | 62 | 50 | 6 | 80 | 10 | 69 | 1 | 0 | 1 |
| 2 | 525 | 133 | 58 | 53 | 3 | 19 | 4 | 13 | 56 | 34 | 0 |
| 3 | 979 | 15 | 8 | 7 | 0 | 4 | 4 | 0 | 3 | 3 | 0 |
| 4 | 1132 | 47 | 43 | 42 | 0 | 3 | 2 | 1 | 1 | 0 | 0 |
| 5 | 1723 | 94 | 61 | 59 | 0 | 31 | 27 | 1 | 2 | 2 | 0 |
| 6 | 787 | 51 | 47 | 47 | 0 | 1 | 1 | 0 | 3 | 3 | 0 |
| 7 | 588 | 80 | 65 | 63 | 0 | 10 | 0 | 10 | 5 | 5 | 0 |
| 8 | 483 | 37 | 32 | 29 | 0 | 2 | 1 | 1 | 3 | 3 | 0 |
| 9 | 46 | 2 | 1 | 1 | 0 | 1 | 1 | 0 | 0 | 0 | 0 |
| 10 | 529 | 25 | 23 | 22 | 0 | 1 | 0 | 1 | 1 | 1 | 0 |
| 11 | 361 | 22 | 14 | 14 | 0 | 7 | 1 | 6 | 1 | 0 | 0 |
| 12 | 1007 | 71 | 49 | 45 | 1 | 17 | 11 | 6 | 5 | 5 | 0 |
| 13 | 271 | 22 | 20 | 20 | 0 | 2 | 0 | 2 | 0 | 0 | 0 |
| 14 | 220 | 26 | 20 | 19 | 0 | 6 | 6 | 0 | 0 | 0 | 0 |
| 15 | 295 | 21 | 18 | 14 | 0 | 3 | 2 | 1 | 0 | 0 | 0 |
| 16 | 123 | 4 | 2 | 2 | 0 | 2 | 2 | 0 | 0 | 0 | 0 |
| 17 | 326 | 14 | 10 | 2 | 8 | 4 | 1 | 3 | 0 | 0 | 0 |
| 18 | 39 | 0 | 0 | 0 | 0 | 0 | 0 | 0 | 0 | 0 | 0 |
| 19 | 1408 | 91 | 80 | 76 | 0 | 7 | 6 | 1 | 4 | 4 | 0 |
| 20 | 418 | 17 | 14 | 14 | 0 | 1 | 1 | 0 | 2 | 1 | 0 |
