## Supplementary Table 3 for "Heterogeneity of circulating epithelial cells in breast cancer at single-cell resolution: identifying tumor and hybrid cells"

Supplementary Table 3. Number of CD45$^+$ CECs in breast cancer patients

| Cases | total number of cells | CD45$^+$ CECs | | | | | | | | | |
|---|---|---|---|---|---|---|---|---|---|---|---|
| | | total CD45$^+$ CECs | Cluster 1 | | | Cluster 2 | | | Cluster 3 | | |
| | | | total | Aneuploid | Diploid | total | Aneuploid | Diploid | total | Aneuploid | Diploid |
| 1 | 1956 | 404 | 211 | 26 | 185 | 184 | 1 | 183 | 9 | 0 | 9 |
| 2 | 525 | 79 | 36 | 1 | 35 | 40 | 0 | 40 | 3 | 0 | 3 |
| 3 | 979 | 6 | 3 | 3 | 0 | 3 | 0 | 3 | 0 | 0 | 0 |
| 4 | 1132 | 6 | 1 | 1 | 0 | 3 | 0 | 3 | 2 | 0 | 2 |
| 5 | 1723 | 7 | 6 | 6 | 0 | 1 | 1 | 0 | 0 | 0 | 0 |
| 6 | 787 | 7 | 0 | 0 | 0 | 4 | 0 | 4 | 3 | 0 | 3 |
| 7 | 588 | 31 | 0 | 0 | 0 | 30 | 0 | 30 | 1 | 0 | 1 |
| 8 | 483 | 23 | 2 | 1 | 1 | 18 | 0 | 18 | 3 | 0 | 3 |
| 9 | 46 | 5 | 3 | 3 | 0 | 2 | 0 | 2 | 0 | 0 | 0 |
| 10 | 529 | 7 | 0 | 0 | 0 | 6 | 0 | 6 | 1 | 0 | 1 |
| 11 | 361 | 52 | 18 | 1 | 17 | 31 | 0 | 31 | 3 | 0 | 3 |
| 12 | 1007 | 24 | 11 | 6 | 5 | 11 | 2 | 9 | 2 | 0 | 2 |
| 13 | 271 | 0 | 0 | 0 | 0 | 0 | 0 | 0 | 0 | 0 | 0 |
| 14 | 220 | 1 | 0 | 0 | 0 | 0 | 0 | 0 | 1 | 0 | 1 |
| 15 | 295 | 6 | 3 | 3 | 0 | 3 | 2 | 1 | 0 | 0 | 0 |
| 16 | 123 | 6 | 2 | 0 | 2 | 4 | 0 | 4 | 0 | 0 | 0 |
| 17 | 326 | 70 | 61 | 3 | 58 | 9 | 0 | 9 | 0 | 0 | 0 |
| 18 | 39 | 2 | 1 | 1 | 0 | 1 | 0 | 1 | 0 | 0 | 0 |
| 19 | 1408 | 4 | 2 | 2 | 0 | 1 | 1 | 0 | 1 | 0 | 1 |
| 20 | 418 | 0 | 0 | 0 | 0 | 0 | 0 | 0 | 0 | 0 | 0 |
