## Supplementary Table 4 for "Heterogeneity of circulating epithelial cells in breast cancer at single-cell resolution: identifying tumor and hybrid cells"

Supplementary Table 4. The percentage of CD45⁻CECs in breast cancer patients

| Cases | total number of cells | CD45⁻ CECs | | | | | | | | | |
|---|---|---|---|---|---|---|---|---|---|---|---|
| | | total CD45⁻ CECs | Cluster 1 | | | Cluster 2 | | | Cluster 3 | | |
| | | | total | Aneuploid | Diploid | total | Aneuploid | Diploid | total | Aneuploid | Diploid |
| 1 | 100 | 7.31 | 3.17 | 2.56 | 0.31 | 4.09 | 0.51 | 3.53 | 0.05 | 0.00 | 0.05 |
| 2 | 100 | 25.33 | 11.05 | 10.10 | 0.57 | 3.62 | 0.76 | 2.48 | 10.67 | 6.48 | 0.00 |
| 3 | 100 | 1.53 | 0.82 | 0.72 | 0.00 | 0.41 | 0.41 | 0.00 | 0.31 | 0.31 | 0.00 |
| 4 | 100 | 4.15 | 3.80 | 3.71 | 0.00 | 0.27 | 0.18 | 0.09 | 0.09 | 0.00 | 0.00 |
| 5 | 100 | 5.46 | 3.54 | 3.42 | 0.00 | 1.80 | 1.57 | 0.06 | 0.12 | 0.12 | 0.00 |
| 6 | 100 | 6.48 | 5.97 | 5.97 | 0.00 | 0.13 | 0.13 | 0.00 | 0.38 | 0.38 | 0.00 |
| 7 | 100 | 13.61 | 11.05 | 10.71 | 0.00 | 1.70 | 0.00 | 1.70 | 0.85 | 0.85 | 0.00 |
| 8 | 100 | 7.66 | 6.63 | 6.00 | 0.00 | 0.41 | 0.21 | 0.21 | 0.62 | 0.62 | 0.00 |
| 9 | 100 | 4.35 | 2.17 | 2.17 | 0.00 | 2.17 | 2.17 | 0.00 | 0.00 | 0.00 | 0.00 |
| 10 | 100 | 4.73 | 4.35 | 4.16 | 0.00 | 0.19 | 0.00 | 0.19 | 0.19 | 0.19 | 0.00 |
| 11 | 100 | 6.09 | 3.88 | 3.88 | 0.00 | 1.94 | 0.28 | 1.66 | 0.28 | 0.00 | 0.00 |
| 12 | 100 | 7.05 | 4.87 | 4.47 | 0.10 | 1.69 | 1.09 | 0.60 | 0.50 | 0.50 | 0.00 |
| 13 | 100 | 8.12 | 7.38 | 7.38 | 0.00 | 0.74 | 0.00 | 0.74 | 0.00 | 0.00 | 0.00 |
| 14 | 100 | 11.82 | 9.09 | 8.64 | 0.00 | 2.73 | 2.73 | 0.00 | 0.00 | 0.00 | 0.00 |
| 15 | 100 | 7.12 | 6.10 | 4.75 | 0.00 | 1.02 | 0.68 | 0.34 | 0.00 | 0.00 | 0.00 |
| 16 | 100 | 3.25 | 1.63 | 1.63 | 0.00 | 1.63 | 1.63 | 0.00 | 0.00 | 0.00 | 0.00 |
| 17 | 100 | 4.29 | 3.07 | 0.61 | 2.45 | 1.23 | 0.31 | 0.92 | 0.00 | 0.00 | 0.00 |
| 18 | 100 | 0.00 | 0.00 | 0.00 | 0.00 | 0.00 | 0.00 | 0.00 | 0.00 | 0.00 | 0.00 |
| 19 | 100 | 6.46 | 5.68 | 5.40 | 0.00 | 0.50 | 0.43 | 0.07 | 0.28 | 0.28 | 0.00 |
| 20 | 100 | 4.07 | 3.35 | 3.35 | 0.00 | 0.24 | 0.24 | 0.00 | 0.48 | 0.24 | 0.00 |

The proportion of aneuploid and diploid CD45⁻ CECs among all cells after reclustering in each case
