## Supplementary Table 5 for "Heterogeneity of circulating epithelial cells in breast cancer at single-cell resolution: identifying tumor and hybrid cells"

Supplementary Table 5. The percentage of CD45$^{+}$ CECs CECs in breast cancer patients (%)

| Cases | total number of cells | CD45$^{+}$ CECs | | | | | | | | | |
|---|---|---|---|---|---|---|---|---|---|---|---|
| | | total CD45$^{+}$ CECs | Cluster 1 | | | Cluster 2 | | | Cluster 3 | | |
| | | | total | Aneuploid | Diploid | total | Aneuploid | Diploid | total | Aneuploid | Diploid |
| 1 | 100 | 20.65 | 10.79 | 1.33 | 9.46 | 9.41 | 0.05 | 9.36 | 0.46 | 0.00 | 0.46 |
| 2 | 100 | 15.05 | 6.86 | 0.19 | 6.67 | 7.62 | 0.00 | 7.62 | 0.57 | 0.00 | 0.57 |
| 3 | 100 | 0.61 | 0.31 | 0.31 | 0.00 | 0.31 | 0.00 | 0.31 | 0.00 | 0.00 | 0.00 |
| 4 | 100 | 0.53 | 0.09 | 0.09 | 0.00 | 0.27 | 0.00 | 0.27 | 0.18 | 0.00 | 0.18 |
| 5 | 100 | 0.41 | 0.35 | 0.35 | 0.00 | 0.06 | 0.06 | 0.00 | 0.00 | 0.06 | 0.00 |
| 6 | 100 | 0.89 | 0.00 | 0.00 | 0.00 | 0.51 | 0.00 | 0.51 | 0.38 | 0.00 | 0.38 |
| 7 | 100 | 5.27 | 0.00 | 0.00 | 0.00 | 5.10 | 0.00 | 5.10 | 0.17 | 0.00 | 0.17 |
| 8 | 100 | 4.76 | 0.41 | 0.21 | 0.21 | 3.73 | 0.00 | 3.73 | 0.62 | 0.00 | 0.62 |
| 9 | 100 | 10.87 | 6.52 | 6.52 | 0.00 | 4.35 | 0.00 | 4.35 | 0.00 | 0.00 | 0.00 |
| 10 | 100 | 1.32 | 0.00 | 0.00 | 0.00 | 1.13 | 0.00 | 1.13 | 0.19 | 0.00 | 0.19 |
| 11 | 100 | 14.40 | 4.99 | 0.28 | 4.71 | 8.59 | 0.00 | 8.59 | 0.83 | 0.00 | 0.83 |
| 12 | 100 | 2.38 | 1.09 | 0.60 | 0.50 | 1.09 | 0.20 | 0.89 | 0.20 | 0.00 | 0.20 |
| 13 | 100 | 0.00 | 0.00 | 0.00 | 0.00 | 0.00 | 0.00 | 0.00 | 0.00 | 0.00 | 0.00 |
| 14 | 100 | 0.45 | 0.00 | 0.00 | 0.00 | 0.00 | 0.00 | 0.00 | 0.45 | 0.00 | 0.45 |
| 15 | 100 | 2.03 | 1.02 | 1.02 | 0.00 | 1.02 | 0.68 | 0.34 | 0.00 | 0.00 | 0.00 |
| 16 | 100 | 4.88 | 1.63 | 0.00 | 1.63 | 3.25 | 0.00 | 3.25 | 0.00 | 0.00 | 0.00 |
| 17 | 100 | 21.47 | 18.71 | 0.92 | 17.79 | 2.76 | 0.00 | 2.76 | 0.00 | 0.00 | 0.00 |
| 18 | 100 | 5.13 | 2.56 | 2.56 | 0.00 | 2.56 | 0.00 | 2.56 | 0.00 | 0.00 | 0.00 |
| 19 | 100 | 0.28 | 0.14 | 0.14 | 0.00 | 0.07 | 0.07 | 0.00 | 0.07 | 0.00 | 0.07 |
| 20 | 100 | 0.00 | 0.00 | 0.00 | 0.00 | 0.00 | 0.00 | 0.00 | 0.00 | 0.00 | 0.00 |

The proportion of aneuploid and diploid CD45$^{+}$ CECs among all cells after reclustering in each case
