## Supplementary Table 6 for "Heterogeneity of circulating epithelial cells in breast cancer at single-cell resolution: identifying tumor and hybrid cells"

Supplementary Table 6. The percentage of diploid and aneuploid cells in $CD45^{-}$ and $CD45^{+}$ CEC clusters

| Cells | $CD45^{-}$ CECs | | | $CD45^{+}$ CECs | | |
|---|---|---|---|---|---|---|
| | Cluster 1 | Cluster 2 | Cluster 3 | Cluster 1 | Cluster 2 | Cluster 3 |
| diploid | 3.02 (18/597) | 58.97 (115/195) | 1.61 (1/62) | 84.17 303/360 | 98.01 344/351 | 100.00 29/29 |
| aneuploid | 96.98 (579/597) | 41.03 (80/195) | 98.39 (61/62) | 15.83 57/360 | 1.99 7/351 | 0.00 0/29 |
