## Supplementary Table 7 for "Heterogeneity of circulating epithelial cells in breast cancer at single-cell resolution: identifying tumor and hybrid cells"

Supplementary Table 7. Upregulated genes in aneuploid $CD45^{-}$ CEC clusters

| Genes | Cluster 1 | | Genes | Cluster 2 | | Genes | Cluster 3 | |
|---|---|---|---|---|---|---|---|---|
| | LogFC | Adjusted P-value | | LogFC | Adjusted P-value | | LogFC | Adjusted P-value |
| *PF4* | 3.55907635243169 | 7.12687755562388E-54 | *KRT5* | 0.711494906650088 | 4.76567105471982E-11 | *HBB* | 6.0310023830796 | 6.1512135554132E-31 |
| *NRGN* | 3.9461137335058 | 1.03257233713827E-53 | *HLA-DRA* | 1.8430917894333 | 1.97849779635367E-06 | *HBA2* | 5.5709512663504 | 2.94254724097695E-26 |
| *PPBP* | 3.91160400586983 | 5.12511370746463E-52 | *BCL11A* | 1.10253816171244 | 2.37327542993918E-06 | *HBA1* | 5.49840551791622 | 1.99329664495992E-24 |
| *GNG11* | 3.3813553509026 | 6.12230172626577E-51 | *CD74* | 2.50459575853138 | 0.000238959192272488 | *SNCA* | 2.70149831794954 | 1.12513069457018E-13 |
| *RGS18* | 3.84180755490312 | 3.93803745337272E-49 | *UGCG* | 1.19463675493907 | 0.000317930746791621 | *ALAS2* | 3.46586744361897 | 1.47190798475692E-12 |
| *CAVIN2* | 3.24517089590332 | 3.15068176014312E-48 | *TCF4* | 1.55377006899908 | 0.00242936300872088 | *HBD* | 3.79377651555706 | 1.47190798475692E-12 |
| *HIST1H2AC* | 3.66386572679067 | 1.05651357985605E-47 | *PPBP* | 1.00710105637514 | 0.00292165217818974 | *AHSP* | 2.45727994778225 | 5.4255278063083E-12 |
| *GP1BB* | 3.64175246274741 | 4.99406537256889E-47 | *JCHAIN* | 1.59814114229495 | 0.00294612093507634 | *HBM* | 2.66171969970519 | 5.4255278063083E-12 |
| *TUBB1* | 3.51093012688524 | 4.64469758969202E-46 | *SEL1L3* | 0.53605290024021 | 0.0040190357826524 | *SLC25A37* | 2.53059799154315 | 3.40865106129183E-11 |
| *LIMS1* | 2.98381734081338 | 1.96284785346431E-42 | *DAB2* | 0.39231742277876 | 0.0181049319386528 | *FKBP8* | 1.4716615249165 | 1.03950450363951E-09 |
| *ACRBP* | 3.00854625977392 | 2.88551331451206E-36 | *CCDC50* | 1.16675772236363 | 0.0183574328182493 | *CA1* | 2.1969834756118 | 2.90692019485686E-09 |
| *PTCRA* | 2.5885026863951 | 1.20770604420691E-34 | *SPIB* | 0.586782936345525 | 0.0245862074579508 | *BLVRB* | 1.80690099853246 | 5.28580135989305E-09 |
| *GP9* | 2.47794347218218 | 2.86212355597438E-33 | *CST3* | 1.35969600120956 | 0.0299642683040604 | *KRT18* | 0.988125899711708 | 2.43111877843464E-08 |
| *PRKAR2B* | 2.03334002514034 | 5.69269012397517E-31 | *RGS18* | 0.592379972283639 | 0.0356386523151597 | *SELENBP1* | 1.27871602806606 | 7.38648238338404E-08 |
| *RUFY1* | 2.50397576179517 | 8.16347623397833E-30 | *RNASE6* | 0.459431618637297 | 0.0379918479951279 | *YBX3* | 2.05613179424418 | 1.33360102804704E-07 |
| *MMD* | 2.17625470224244 | 2.20975103294572E-29 | | | | *UBB* | 2.54056384708825 | 1.4430930704255E-07 |
| *DAB2* | 1.88327639369832 | 3.35191665482421E-29 | | | | *SLC25A39* | 2.2071076928855 | 1.5629831235959E-07 |
| *TSC22D1* | 2.87083057224391 | 8.43390579457991E-29 | | | | *MPP1* | 1.3801999734065 | 2.43739113654074E-07 |
| *RGS10* | 2.92413672631063 | 9.63239494446067E-29 | | | | *BSG* | 1.68274520316832 | 3.93205902115367E-07 |
| *OAZ1* | 3.10324002156866 | 2.47524301318034E-28 | | | | *OAZ1* | 1.56714183377073 | 4.27193905959223E-07 |
| *TMEM40* | 2.32299557184386 | 1.00183320699806E-27 | | | | *STRADB* | 1.04723635610711 | 5.31425772216371E-07 |
| *MAP3K7CL* | 2.05490144841295 | 1.33214354151997E-27 | | | | *GYPC* | 2.03538636482595 | 7.65158893462396E-07 |
| *TUBA4A* | 2.59807703623041 | 1.7689882576445E-27 | | | | *SLC4A1* | 0.976153258045632 | 1.84814586674706E-06 |
| *NT5C3A* | 2.20596532345243 | 4.8988393371062E-27 | | | | *MKRN1* | 1.4349322211455 | 2.76200783451662E-06 |
| *C2orf88* | 2.07404572025356 | 6.84103447879683E-25 | | | | *BNIP3L* | 1.44096472769358 | 2.77786468027959E-06 |

| | | |
|---|---|---|
| *MPP1* | 2.06162485449836 | 1.06570853353321E-24 |
| *RAB32* | 1.89974492736924 | 1.23131208999182E-24 |
| *ODC1* | 2.23284251078379 | 8.26935949286108E-24 |
| *PLEKHO1* | 1.8596948604084 | 6.58641188231497E-22 |
| *CMTM5* | 1.87490590850592 | 1.49498514403822E-21 |
| *HIST1H2BJ* | 1.78193255782953 | 4.63165189400448E-21 |
| *TAGLN2* | 2.523937300038 | 4.2109098311395E-20 |
| *GSTO1* | 2.20988925888667 | 4.83543136178458E-20 |
| *PTPN18* | 1.69449654255795 | 2.10926217886774E-19 |
| *H3F3A* | 2.47896063144133 | 2.50087017914704E-19 |
| *PLA2G12A* | 1.44711286583531 | 2.8453910964698E-19 |
| *GPX4* | 1.91905748906773 | 5.16205095761182E-19 |
| *F13A1* | 1.47673163251337 | 9.42718183029499E-19 |
| *MTURN* | 1.53722667265013 | 1.66955552183237E-18 |
| *TLN1* | 2.0776915446289 | 1.72954668311792E-18 |
| *VCL* | 1.62776452078339 | 3.70772784871163E-18 |
| *CLU* | 1.82346550139447 | 6.05314163533734E-18 |
| *HIST1H3H* | 2.06172927335073 | 7.28856274366846E-18 |
| *FERMT3* | 1.66334842812187 | 1.38179480671741E-17 |
| *AC127502.2* | 1.78224034918829 | 1.58655348395446E-17 |
| *RBX1* | 1.40612744200471 | 1.96985770432901E-17 |
| *SPARC* | 1.50657584904448 | 2.19036834985176E-17 |
| *GRAP2* | 1.65778873618055 | 2.43296721866752E-17 |
| *NCOA4* | 1.74844029418587 | 5.50107842230093E-17 |
| *TRIM58* | 1.32215459540449 | 7.55249382434951E-17 |
| *MAX* | 1.78774883074935 | 1.29536290236378E-16 |
| *RAP1B* | 2.2067505872809 | 2.3912133242037E-16 |
| *TSPAN33* | 1.26137231458902 | 3.67821422230231E-16 |
| *TPM4* | 1.6573348841845 | 4.87588215941774E-16 |

| | | |
|---|---|---|
| *PDZK1IP1* | 0.810729648838261 | 5.21702511599888E-06 |
| *FAM210B* | 0.871031335952171 | 0.0000135728520494557 |
| *DCAF12* | 0.914752713381489 | 0.0000144850682652632 |
| *TRIM58* | 0.775156810670314 | 0.0000428196817499134 |
| *GLRX5* | 1.47629432570089 | 0.0000945755339111815 |
| *NCOA4* | 0.881911884164231 | 0.000105605606411941 |
| *DMTN* | 0.712041812191014 | 0.000149663976401336 |
| *LGALS3* | 1.37815323422173 | 0.000175106699522584 |
| *GUK1* | 1.34030958832229 | 0.000306889185792455 |
| *ADIPOR1* | 1.12137781885962 | 0.00109041141484005 |
| *GMPR* | 0.862148629342801 | 0.00135216013910317 |
| *BAG1* | 0.730531783297528 | 0.00330456675113301 |
| *FBXO7* | 1.17909331671591 | 0.00542826262898899 |
| *SNX3* | 0.988870901040367 | 0.00583850240083024 |
| *HAGH* | 0.745634620616376 | 0.00873235343027938 |
| *EPB42* | 0.495527417139212 | 0.0122217261045839 |
| *NFE2* | 0.426814667055197 | 0.0122217261045839 |
| *FECH* | 0.775156810670314 | 0.012630992128137 |
| *IFIT1B* | 0.727474145188908 | 0.0303923497214864 |
| *BCL2L1* | 0.803284981102587 | 0.0378351082867348 |

| | | |
|---|---|---|
| *FTH1* | 2.11029284156021 | 5.2701658083294E-16 |
| *OST4* | 1.84412832898652 | 6.00067070307234E-16 |
| *PDLIM1* | 1.50870402225165 | 8.02680265978423E-16 |
| *AP003068.2* | 1.24646463466444 | 9.51029638175986E-16 |
| *CLDN5* | 1.41526399979597 | 1.31707537423469E-15 |
| *TMEM140* | 1.43080769530489 | 1.3442874982639E-15 |
| *CALM3* | 1.49074836141285 | 2.00160544163124E-15 |
| *ESAM* | 1.48442602430337 | 3.30420632216123E-15 |
| *PDZK1IP1* | 1.27379157384309 | 3.54484304599325E-15 |
| *CA2* | 1.50378067238673 | 4.4709056942367E-15 |
| *CTSA* | 1.60713688104869 | 4.97349422963023E-15 |
| *YWHAH* | 1.59420024253325 | 1.08178869244686E-14 |
| *MARCH2* | 1.36132369050296 | 1.45935096119335E-14 |
| *DAPP1* | 1.38277680016928 | 1.61323193965934E-14 |
| *KRT8* | 0.804147743913778 | 3.1695527070307E-14 |
| *TREML1* | 1.18689217530286 | 3.98057188051159E-14 |
| *CTTN* | 1.01116930771811 | 4.68439514622261E-14 |
| *CCL5* | 1.05362676192379 | 4.73372934629071E-14 |
| *RAB27B* | 1.42459697586391 | 5.29315913196734E-14 |
| *BEX3* | 1.25966370822879 | 5.35682375644029E-14 |
| *ITGA2B* | 1.01979715512082 | 5.68994914520865E-14 |
| *PGRMC1* | 1.44445908432953 | 6.82927465918906E-14 |
| *DYNLL1* | 1.30078214856791 | 8.12876788862728E-14 |
| *CLEC1B* | 1.2610100519904 | 1.00788626975826E-13 |
| *GNAS* | 1.94815778037429 | 1.44405051882214E-13 |
| *AC147651.1* | 1.29962009047779 | 1.53172998534957E-13 |
| *MPIG6B* | 1.17428296139494 | 1.70992340090791E-13 |
| *KIF2A* | 1.56413171552101 | 2.09470699790698E-13 |
| *RAB11A* | 1.41971500865449 | 2.12577485805971E-13 |

| | | |
|---|---|---|
| *ETFA* | 1.30244986657812 | 2.16585004454981E-13 |
| *LYL1* | 1.51251605336141 | 4.36538938459498E-13 |
| *CARD19* | 1.30727010173674 | 8.18415514391553E-13 |
| *TALDO1* | 1.3277303809539 | 9.11497628249603E-13 |
| *ARHGAP6* | 1.01856776003927 | 9.84355021329617E-13 |
| *LAMTOR1* | 1.38926879126303 | 1.61359776924531E-12 |
| *SH3BGRL2* | 0.823988656403572 | 1.71602724927549E-12 |
| *NFE2* | 1.184524235746 | 2.06317794790089E-12 |
| *TGFB1* | 1.57812029604698 | 4.84595021265627E-12 |
| *LCN2* | 0.973597000903011 | 5.14421786696122E-12 |
| *LMNA* | 1.19776581583935 | 8.39668185815715E-12 |
| *LGALSL* | 1.11457085958212 | 1.02555491358492E-11 |
| *SMIM5* | 1.08595637454301 | 1.04830861267285E-11 |
| *MTPN* | 1.19424126909776 | 1.22487007750645E-11 |
| *EIF2AK1* | 1.28963626329021 | 1.77863740014503E-11 |
| *GMPR* | 1.11063225928696 | 1.81469308438894E-11 |
| *PDGFA* | 1.01487327648153 | 1.84388020296456E-11 |
| *ARHGAP18* | 1.0654168461894 | 2.13783168831732E-11 |
| *SMOX* | 1.16156257003011 | 3.35214173474172E-11 |
| *NORAD* | 1.2992609230515 | 4.4872890338359E-11 |
| *NAP1L1* | 1.68269106373664 | 5.15535636432948E-11 |
| *NEXN* | 1.25762870405721 | 1.42913098396322E-10 |
| *AC090409.1* | 0.873789402925528 | 1.76357178508155E-10 |
| *GAS2L1* | 0.837995514397451 | 1.76357178508155E-10 |
| *PRDX6* | 1.14872902321632 | 1.76701064050345E-10 |
| *ENKUR* | 1.08616036111683 | 2.46072831670253E-10 |
| *THBS1* | 0.602299817154167 | 2.95009884836041E-10 |
| *TAL1* | 0.893311296600617 | 4.72473609399464E-10 |
| *PDE5A* | 0.750358424093105 | 4.91569653025464E-10 |

| | | |
|---|---|---|
| *RIOK3* | 1.22152705037583 | 5.81052638231803E-10 |
| *C19orf33* | 1.14928230991184 | 7.84659519666956E-10 |
| *SOD2* | 0.96988582170822 | 1.10834300249665E-09 |
| *KRT18* | 0.819759942948385 | 1.34929306786963E-09 |
| *SNAP23* | 1.10140616095339 | 2.68192953688605E-09 |
| *HIST2H2BE* | 0.726462307746944 | 4.30096694937186E-09 |
| *H2AFJ* | 1.28235358734652 | 5.02084917208786E-09 |
| *SLC40A1* | 1.0033776376437 | 5.19052919651212E-09 |
| *MYL12A* | 1.67546348201606 | 5.61055219250606E-09 |
| *STOM* | 1.2025002796573 | 6.63857533065138E-09 |
| *NCK2* | 1.19912216629374 | 8.26175159688293E-09 |
| *FRMD3* | 0.738459840846537 | 1.57978104277397E-08 |
| *CNST* | 1.13794649887785 | 1.80924198757919E-08 |
| *GFI1B* | 0.724955576496789 | 1.85554289150076E-08 |
| *GUCY1B1* | 0.7006313747393 | 2.17861403467902E-08 |
| *TIMP1* | 1.18641655632223 | 2.17952217944669E-08 |
| *MYLK* | 0.945212398512647 | 2.40143924798858E-08 |
| *TLK1* | 1.06321281075299 | 3.61053255185842E-08 |
| *LEPROT* | 0.820264011658621 | 4.16211381370304E-08 |
| *SNCA* | 0.848980029058268 | 5.83987431386629E-08 |
| *DNM3* | 0.642759424425951 | 7.76290147174563E-08 |
| *ARPC1B* | 1.44606654415141 | 9.0857921296277E-08 |
| *R3HDM4* | 1.0594785786027 | 1.11086254612873E-07 |
| *FAM110A* | 1.07941352867624 | 1.33726748188324E-07 |
| *RPA1* | 0.803369171287819 | 1.95160862144604E-07 |
| *AL162424.1* | 0.757745511614458 | 1.98280029639153E-07 |
| *CDKN2D* | 1.19586558182668 | 2.11840346748896E-07 |
| *RASGRP2* | 1.48098496035464 | 2.12020325300432E-07 |
| *TPM1* | 0.93615028479535 | 2.29460236884597E-07 |

| | | |
|---|---|---|
| *FKBP1A* | 0.871158154362067 | 2.34825282256698E-07 |
| *NUTF2* | 0.820264011658621 | 2.57904746685812E-07 |
| *SAT1* | 1.45035919949379 | 3.91397883899945E-07 |
| *TNNC2* | 0.637963757824302 | 4.29001427266137E-07 |
| *PTGS1* | 0.834301030839709 | 4.85344689871345E-07 |
| *SSX2IP* | 0.854338234069463 | 5.45406947005725E-07 |
| *ILK* | 1.15397292269093 | 5.63992590171456E-07 |
| *CYTOR* | 1.01938170215328 | 6.89961178305401E-07 |
| *ASAH1* | 0.834455901221576 | 7.39260168248742E-07 |
| *ERV3-1* | 1.00044620584625 | 7.8153736412171E-07 |
| *CDKN1A* | 0.705941160698895 | 7.95211943015374E-07 |
| *CST3* | 1.01095822647815 | 1.02108951637354E-06 |
| *HEXIM2* | 0.658630817068833 | 1.07097823454899E-06 |
| *ARPC5* | 1.04419385015175 | 1.24412631192351E-06 |
| *CTDSPL* | 0.690818432701874 | 0.0000012836547605572 |
| *LINC00989* | 0.805202096246461 | 1.48981360504413E-06 |
| *VIM-AS1* | 0.963889347762725 | 1.52534488804184E-06 |
| *TUBA1C* | 0.884189906632068 | 1.62962747652816E-06 |
| *SPINT2* | 0.69772232185604 | 1.63329260104431E-06 |
| *ANO6* | 0.73360329919867 | 1.79279391429583E-06 |
| *WBP2* | 0.989821904426163 | 1.90823975725051E-06 |
| *FAXDC2* | 0.70292927016679 | 2.47394814664581E-06 |
| *CMIP* | 0.908642950356607 | 2.62229796154768E-06 |
| *TRAPPC5* | 0.885144151864834 | 3.66527407625141E-06 |
| *SH3BGRL3* | 1.75389252941571 | 3.78206956000138E-06 |
| *DMTN* | 0.757569543687803 | 3.92898904316727E-06 |
| *SNN* | 0.691748420811407 | 4.31852520724729E-06 |
| *RIPOR2* | 1.4422340542105 | 5.32178818395387E-06 |
| *ROCK2* | 0.57913678453485 | 0.0000064420997151397 |

| | | |
|---|---|---|
| *PTGIR* | 0.543679650717948 | 7.46565411571993E-06 |
| *AP001189.1* | 0.749007279864574 | 8.12771294270693E-06 |
| *SUPT4H1* | 0.808422399014104 | 8.37241522676105E-06 |
| *ACTN1* | 0.793964346320413 | 9.42977806488878E-06 |
| *LY6G6F* | 0.516067419510089 | 0.0000115980219639088 |
| *AP1S2* | 0.739595274892989 | 0.0000118340483393438 |
| *DNAJB6* | 1.09960949984801 | 0.000011965037575651 |
| *CD9* | 0.717487365856959 | 0.000012956488128676 |
| *CAPZA2* | 0.998337908755231 | 0.0000131040690153835 |
| *EMC3* | 0.941809814291807 | 0.0000148170737002279 |
| *SCN1B* | 0.652303197096822 | 0.0000149175733920177 |
| *TACC3* | 0.79207600175646 | 0.0000153000541990321 |
| *HIST1H1C* | 1.11862120131936 | 0.0000163674680319671 |
| *BMP6* | 0.493236711129203 | 0.0000179682889411388 |
| *ADIPOR1* | 0.894191794059067 | 0.0000185694703602368 |
| *INAFM2* | 0.666287426532523 | 0.0000202543927310481 |
| *CORO1C* | 0.484358559782751 | 0.0000207787311350305 |
| *SWI5* | 0.533386895710115 | 0.0000207787311350316 |
| *BEND2* | 0.502060561516209 | 0.0000207787311350329 |
| *AKIRIN2* | 1.00937309810444 | 0.0000213152845184775 |
| *TNFSF4* | 0.707444751289283 | 0.0000227079516331009 |
| *ITGB3* | 0.672160615442009 | 0.0000253997518030025 |
| *GNAZ* | 0.503818873788568 | 0.0000277622566788861 |
| *MEIS1* | 0.623480361931068 | 0.0000277622566788861 |
| *ITM2B* | 1.33031004414803 | 0.0000283023288227518 |
| *YWHAE* | 0.703650077097042 | 0.0000406756002156681 |
| *NT5M* | 0.619136333161623 | 0.000049284277495991 |
| *SELP* | 0.459205082554412 | 0.0000493843915570765 |
| *TMEM91* | 0.717487365856959 | 0.0000516735393283004 |

| | | |
|---|---|---|
| *PYGL* | 0.54538799318031 | 0.0000657505858012929 |
| *SMIM3* | 0.751838870647736 | 0.0000739478514318835 |
| *SENCR* | 0.553899493035827 | 0.0000758346219696475 |
| *MOB1B* | 0.62278013587048 | 0.0000957105737927562 |
| *ANKRD9* | 0.529939594091646 | 0.00010079349624042 |
| *SEC14L1* | 0.777951710966868 | 0.000102685984286293 |
| *CXCL5* | 0.568738480457691 | 0.000116845304402643 |
| *SPNS1* | 0.883257631936694 | 0.000125673102185676 |
| *AL034397.3* | 0.701420963661513 | 0.000130557156247324 |
| *ABCC3* | 0.448289305075229 | 0.000133815819414058 |
| *KIFC3* | 0.475425435012743 | 0.000154121350757993 |
| *PBX1* | 0.439129305789753 | 0.000154121350758002 |
| *TBXA2R* | 0.480791944762558 | 0.000177458828186984 |
| *MFAP3L* | 0.578631285216123 | 0.000184801958616558 |
| *FTL* | 1.45984212464424 | 0.000214182898623048 |
| *RSU1* | 0.708603594260065 | 0.000224107354339673 |
| *RNF11* | 0.670114473213283 | 0.000230514362187955 |
| *PCP2* | 0.46463215493874 | 0.000270448918281741 |
| *STXBP2* | 0.774739272757815 | 0.000328302947872689 |
| *CLIC1* | 1.25547029319557 | 0.000343960304439724 |
| *AL731557.1* | 0.457391512182481 | 0.000357673024399085 |
| *MLH3* | 0.78927884120644 | 0.000398947600428833 |
| *PEAR1* | 0.450114346483555 | 0.000411160466606843 |
| *CDK2AP1* | 0.461016376001235 | 0.000472520335072583 |
| *MINDY1* | 0.521285128781108 | 0.000542893060604532 |
| *EGFL7* | 0.507329082432389 | 0.000542893060604564 |
| *MKRN1* | 0.676704540586957 | 0.000575662795078363 |
| *TSPOAP1-AS1* | 0.594242380394984 | 0.000592194064506708 |
| *PF4V1* | 0.786636541858143 | 0.000622129873458491 |

| | | |
|---|---|---|
| *HIST1H2BG* | 0.451937082090373 | 0.000623581691272142 |
| *ITGB5* | 0.448289305075229 | 0.000623581691272142 |
| *PKM* | 0.969430305720363 | 0.000662536815324545 |
| *MCUR1* | 0.692073725789423 | 0.0007296232269127 |
| *NDUFA6* | 0.906059789342605 | 0.000735415902480204 |
| *STK24* | 0.630608125997503 | 0.000737866409743236 |
| *ARHGAP21* | 0.591036390095893 | 0.000796658342409964 |
| *GTPBP2* | 0.523020180151856 | 0.000943514654649791 |
| *PSTPIP2* | 0.751838870647736 | 0.00097781682867036 |
| *EGLN3* | 0.461016376001235 | 0.00108261803311588 |
| *AQP10* | 0.509080989193982 | 0.00108261803311595 |
| *LINC00853* | 0.455575659153721 | 0.00108261803311595 |
| *SQSTM1* | 0.902450744080552 | 0.00118066630257728 |
| *LYPLAL1* | 0.686743842425852 | 0.00122952085578321 |
| *TRAPPC3L* | 0.470038888465684 | 0.00142427741821877 |
| *PPP1R14A* | 0.611125790421912 | 0.00145659932265591 |
| *RAP2B* | 0.827787723881035 | 0.00148428872178491 |
| *SERPINB1* | 0.762356742561736 | 0.00166455177342081 |
| *CCND3* | 0.87469561843327 | 0.00175819710652378 |
| *UBXN11* | 0.70292927016679 | 0.00186771817364746 |
| *ASAP2* | 0.422493304197891 | 0.00187185855935683 |
| *GP6* | 0.413167511925798 | 0.00187185855935683 |
| *SEPTIN11* | 0.587823259500707 | 0.00190632263570678 |
| *PIP4K2A* | 0.765044409447004 | 0.00239339777909452 |
| *MYL9* | 0.905043709302539 | 0.00243307534806865 |
| *TMEM50A* | 0.759469311772114 | 0.00256372082542916 |
| *HIST1H2AE* | 0.548696928210885 | 0.002582440825572 |
| *ALOX12* | 0.530174899233803 | 0.00258561007879686 |
| *VDAC3* | 0.714161667503864 | 0.00260103497247284 |

| | | |
|---|---|---|
| *MGAT4B* | 0.68186765463896 | 0.00295611676293932 |
| *LAT* | 0.944179550437112 | 0.00303200461235569 |
| *NPTN* | 0.507913287761647 | 0.00305134583482809 |
| *AIG1* | 0.388634500181855 | 0.00322350121144541 |
| *PTP4A2* | 0.810732559701305 | 0.00370743452460038 |
| *TPST2* | 0.795366246188658 | 0.00382640338953853 |
| *AC123912.4* | 0.484358559782751 | 0.00422390880532124 |
| *INKA1* | 0.526484035455427 | 0.00422390880532175 |
| *PNMA1* | 0.652303197096822 | 0.00434102874245945 |
| *ENDOD1* | 0.506186543042179 | 0.00634469215291296 |
| *HEMGN* | 0.695594760008117 | 0.00667862584886059 |
| *GABARAPL2* | 0.653640883832334 | 0.00675875824659786 |
| *RYBP* | 0.577201741856103 | 0.00682983786332116 |
| *CAV2* | 0.453757517714742 | 0.00826711928160127 |
| *ELOVL7* | 0.49482107483659 | 0.00905035939209929 |
| *CD68* | 0.478584622496063 | 0.00946131117948693 |
| *PIP4P2* | 0.625167067159308 | 0.00947593003211796 |
| *PTPN12* | 0.653590742957282 | 0.00978632866246476 |
| *RAB31* | 0.444410345455489 | 0.0103050781566836 |
| *SVIP* | 0.788833078129938 | 0.0107420952298616 |
| *PPP3R1* | 0.499958273288631 | 0.0107446307383192 |
| *PRKCD* | 0.386729951996495 | 0.0107967487456844 |
| *CD36* | 0.353961922527076 | 0.0107967487456851 |
| *KIAA0513* | 0.35005746807116 | 0.012334210946669 |
| *HIST1H2AG* | 0.593535788846948 | 0.0126733449778839 |
| *CYB5R1* | 0.504457729125428 | 0.0127330240469221 |
| *YWHAZ* | 0.796896145633076 | 0.0130284401446308 |
| *TAX1BP3* | 0.480034255186211 | 0.0130923969927892 |
| *FHL1* | 0.35979886254936 | 0.0140873711721927 |

| | | |
|---|---|---|
| *HACD4* | 0.639181069878274 | 0.0148041458585889 |
| *IGF2BP3* | 0.531873186705829 | 0.0151752536835243 |
| *INF2* | 0.437178776224413 | 0.0153079644001385 |
| *SAV1* | 0.476511188622513 | 0.0159277568577391 |
| *TST* | 0.469439139947012 | 0.0164394130840464 |
| *CABP5* | 0.405663229513697 | 0.0183641333795086 |
| *PECAM1* | 0.363677077360287 | 0.0183641333795108 |
| *CD99* | 0.770309474146462 | 0.0197032182692137 |
| *MT-ND2* | 1.1379795175735 | 0.0197159552394237 |
| *ASAP1* | 0.645027866293647 | 0.0203266226179892 |
| *MYL4* | 0.647539202539286 | 0.0209601160712601 |
| *MISP3* | 0.48530277610239 | 0.0209787609244846 |
| *USF2* | 0.636564549654671 | 0.0210993581080451 |
| *RTN3* | 0.672202754534527 | 0.0216183323627642 |
| *RDH11* | 0.59284934688658 | 0.0223815646155652 |
| *ZNF185* | 0.428088040153557 | 0.022381787088813 |
| *P2RX1* | 0.361739273118795 | 0.0239177232939854 |
| *CD226* | 0.679783933518929 | 0.0250953315719864 |
| *UBA7* | 0.428088040153557 | 0.025146314513048 |
| *ARG2* | 0.429910775760374 | 0.0272866100512861 |
| *CMPK1* | 0.666350971582455 | 0.028310934223278 |
| *TMEM158* | 0.478273797321773 | 0.0310943712999144 |
| *TMEM219* | 0.590902652432679 | 0.032367161573009 |
| *LDLRAP1* | 0.599061175819843 | 0.0339038851812145 |
| *INKA2* | 0.400009290046512 | 0.0354913581719949 |
| *SYTL4* | 0.371402370469515 | 0.0354913581719971 |
| *LTBP1* | 0.318435488897934 | 0.0354913581719992 |
| *BCL2L1* | 0.519422027181462 | 0.0356728208584909 |
| *TMBIM1* | 0.698777626896323 | 0.0367095701640852 |

| | | |
|---|---|---|
| *CLCN3* | 0.525373405719909 | 0.0396453567211721 |
| *PTK2* | 0.381001175279428 | 0.0404638589393004 |
| *MAGED2* | 0.451605846859693 | 0.0473071204201932 |
