## Supplementary Table 8 for "Heterogeneity of circulating epithelial cells in breast cancer at single-cell resolution: identifying tumor and hybrid cells"

Supplementary Table 8. Specific upregulated genes in aneuploid CD45$^{-}$ CEC clusters

| Genes | Cluster 1 | | Genes | Cluster 2 | | Genes | Cluster 3 | |
|---|---|---|---|---|---|---|---|---|
| | LogFC | Adjusted P-value | | LogFC | Adjusted P-value | | LogFC | Adjusted P-value |
| *PF4* | 3.55907635243169 | 7.12687755562388E-54 | *KRT5* | 0.711494906650088 | 4.76567105471982E-11 | *HBB* | 6.0310023830796 | 6.1512135554132E-31 |
| *NRGN* | 3.9461137335058 | 1.03257233713827E-53 | *HLA-DRA* | 1.8430917894333 | 1.97849779635367E-06 | *HBA2* | 5.5709512663504 | 2.94254724097695E-26 |
| *GNG11* | 3.3813553509026 | 6.12230172626577E-51 | *BCL11A* | 1.10253816171244 | 2.37327542993918E-06 | *HBA1* | 5.49840551791622 | 1.99329664495992E-24 |
| *CAVIN2* | 3.24517089590332 | 3.15068176014312E-48 | *CD74* | 2.50459575853138 | 0.000238959192272488 | *ALAS2* | 3.46586744361897 | 1.47190798475692E-12 |
| *HIST1H2AC* | 3.66386572679067 | 1.05651357985605E-47 | *UGCG* | 1.19463675493907 | 0.000317930746791621 | *HBD* | 3.79377651555706 | 1.47190798475692E-12 |
| *GP1BB* | 3.64175246274741 | 4.99406537256889E-47 | *TCF4* | 1.55377006899908 | 0.00242936300872088 | *HBM* | 2.66171969970519 | 5.4255278063083E-12 |
| *TUBB1* | 3.51093012688524 | 4.64469758969202E-46 | *JCHAIN* | 1.59814114229495 | 0.00294612093507634 | *SLC25A37* | 2.53059799154315 | 3.40865106129183E-11 |
| *LIMS1* | 2.98381734081338 | 1.96284785346431E-42 | *SEL1L3* | 0.53605290024021 | 0.0040190357826524 | *CA1* | 2.1969834756118 | 2.90692019485686E-09 |
| *ACRBP* | 3.00854625977392 | 2.88551331451206E-36 | *CCDC50* | 1.16675772236363 | 0.0183574328182493 | *BLVRB* | 1.80690099853246 | 5.28580135989305E-09 |
| *PTCRA* | 2.5885026863951 | 1.20770604420691E-34 | *SPIB* | 0.586782936345525 | 0.0245862074579508 | *YBX3* | 2.05613179424418 | 1.33360102804704E-07 |
| *GP9* | 2.47794347218218 | 2.86212355597438E-33 | *RNASE6* | 0.459431618637297 | 0.0379918479951279 | *UBB* | 2.54056384708825 | 1.4430930704255E-07 |
| *PRKAR2B* | 2.03334002514034 | 5.69269012397517E-31 | | | | *SLC25A39* | 2.2071076928855 | 1.5629831235959E-07 |
| *RUFY1* | 2.50397576179517 | 8.16347623397833E-30 | | | | *BSG* | 1.68274520316832 | 3.93205902115367E-07 |
| *MMD* | 2.17625470224244 | 2.20975103294572E-29 | | | | *STRADB* | 1.04723635610711 | 5.31425772216371E-07 |
| *TSC22D1* | 2.87083057224391 | 8.43390579457991E-29 | | | | *GYPC* | 2.03538636482595 | 7.65158893462396E-07 |
| *RGS10* | 2.92413672631063 | 9.63239494446067E-29 | | | | *SLC4A1* | 0.976153258045632 | 1.84814586674706E-06 |
| *TMEM40* | 2.32299557184386 | 1.00183320699806E-27 | | | | *BNIP3L* | 1.44096472769358 | 2.77786468027959E-06 |
| *MAP3K7CL* | 2.05490144841295 | 1.33214354151997E-27 | | | | *FAM210B* | 0.871031335952171 | 0.0000135728520494557 |
| *TUBA4A* | 2.59807703623041 | 1.7689882576445E-27 | | | | *DCAF12* | 0.914752713381489 | 0.0000144850682652632 |
| *NT5C3A* | 2.20596532345243 | 4.8988393371062E-27 | | | | *GLRX5* | 1.47629432570089 | 0.0000945755339111815 |
| *C2orf88* | 2.07404572025356 | 6.84103447879683E-25 | | | | *LGALS3* | 1.37815323422173 | 0.000175106699522584 |
| *RAB32* | 1.89974492736924 | 1.23131208999182E-24 | | | | *GUK1* | 1.34030958832229 | 0.000306889185792455 |
| *ODC1* | 2.23284251078379 | 8.26935949286108E-24 | | | | *BAG1* | 0.730531783297528 | 0.00330456675113301 |
| *PLEKHO1* | 1.8596948604084 | 6.58641188231497E-22 | | | | *FBXO7* | 1.17909331671591 | 0.00542826262898899 |
| *CMTM5* | 1.87490590850592 | 1.49498514403822E-21 | | | | *SNX3* | 0.988870901040367 | 0.00583850240083024 |
| *HIST1H2BJ* | 1.78193255782953 | 4.63165189400448E-21 | | | | *HAGH* | 0.745634620616376 | 0.00873235343027938 |
| *TAGLN2* | 2.523937300038 | 4.2109098311395E-20 | | | | *EPB42* | 0.495527417139212 | 0.0122217261045839 |
| *GSTO1* | 2.20988925888667 | 4.83543136178458E-20 | | | | *FECH* | 0.775156810670314 | 0.012630992128137 |
| *PTPN18* | 1.69449654255795 | 2.10926217886774E-19 | | | | *IFIT1B* | 0.727474145188908 | 0.0303923497214864 |
| *H3F3A* | 2.47896063144133 | 2.50087017914704E-19 | | | | | | |
| *PLA2G12A* | 1.44711286583531 | 2.8453910964698E-19 | | | | | | |
| *GPX4* | 1.91905748906773 | 5.16205095761182E-19 | | | | | | |
| *F13A1* | 1.47673163251337 | 9.42718183029499E-19 | | | | | | |
| *MTURN* | 1.53722667265013 | 1.66955552183237E-18 | | | | | | |
