## Supplementary Table 10 for "Heterogeneity of circulating epithelial cells in breast cancer at single-cell resolution: identifying tumor and hybrid cells"

Supplementary Table 10. Specific signaling pathways enriched in aneuploid CD45⁻ CEC clusters

| Cluster 1 | | |
|---|---|---|
| Term | Adjusted P-value | Genes |
| Platelet activation | 2.3658483228682E-11 | *PTGIR;GUCY1B1;GP1BB;ROCK2;ITGB3;ITGA2B;SNAP23;RASGRP2;GP6;MYL12A;MYLK;PTGS1;RAP1B;GP9;TBXA2R;P2RX1;GNAS;TLN1;FERMT3* |
| Focal adhesion | 2.94720483638925E-06 | *ITGB5;CAV2;ROCK2;ACTN1;ITGB3;ITGA2B;PDGFA;ILK;THBS1;MYL12A;PTK2;MYLK;RAP1B;CCND3;TLN1;MYL9;VCL* |
| Shigellosis | 7.07650252242854E-06 | *GABARAPL2;ROCK2;ACTN1;ARPC1B;PRKCD;ILK;ARPC5;MYL12A;PTK2;SEPTIN11;RBX1;CTTN;CCL5;TLN1;MYL9;SQSTM1;VCL;BCL2L1* |
| Leukocyte transendothelial migration | 0.000123412832738766 | *RAP1B;CLDN5;ROCK2;ACTN1;PECAM1;ESAM;MYL9;VCL;MYL12A;CD99;PTK2* |
| Ferroptosis | 0.000143741222663461 | *GPX4;FTH1;NCOA4;SLC40A1;VDAC3;SAT1;FTL* |
| Bacterial invasion of epithelial cells | 0.000143741222663461 | *DNM3;CTTN;CAV2;ARPC1B;ILK;ARPC5;SEPTIN11;VCL;PTK2* |
| Regulation of actin cytoskeleton | 0.000376998095621831 | *ITGB5;ROCK2;ACTN1;ARPC1B;ITGB3;ITGA2B;PDGFA;ARPC5;MYL12A;PTK2;MYLK;PIP4K2A;MYL9;VCL* |
| Vascular smooth muscle contraction | 0.00160280740573598 | *PPP1R14A;PLA2G12A;PTGIR;GUCY1B1;ROCK2;PRKCD;GNAS;CALM3;MYL9;MYLK* |
| ECM-receptor interaction | 0.0021091696184099 | *GP9;ITGB5;GP1BB;ITGB3;ITGA2B;CD36;GP6;THBS1* |
| Dilated cardiomyopathy | 0.00350937633562694 | *TGFB1;ITGB5;TPM4;ITGB3;ITGA2B;TPM1;LMNA;GNAS* |
| Chemokine signaling pathway | 0.00570790178232303 | *RAP1B;ROCK2;CCL5;PRKCD;PPBP;RASGRP2;GNG11;PTK2;CXCL5;PF4V1;PF4* |
| Tight junction | 0.00714075305820634 | *CLDN5;TUBA1C;CTTN;ROCK2;ACTN1;ARPC1B;ARPC5;MYL9;TUBA4A;MYL12A* |
| Fluid shear stress and atherosclerosis | 0.00714075305820634 | *CAV2;GSTO1;ITGB3;ITGA2B;PECAM1;PDGFA;CALM3;SQSTM1;PTK2* |

| | | |
|---|---|---|
| Hypertrophic cardiomyopathy | 0.00988751277802386 | *TGFB1;ITGB5;TPM4;ITGB3;ITGA2B;TPM1;LMNA* |
| Endocytosis | 0.0118751047514207 | *RUFY1;DNM3;DAB2;RAB31;CAV2;ARPC1B;CAPZA2;ASAP1;ARPC5;ASAP2;LDLRAP1;RAB11A* |
| Cell cycle | 0.0124219034058814 | *YWHAE;CDKN2D;CCND3;CDKN1A;TGFB1;YWHAZ;RBX1;YWHAH* |
| Rap1 signaling pathway | 0.0268016044882464 | *RAP1B;ITGB3;ITGA2B;GNAS;PDGFA;CALM3;TLN1;RASGRP2;THBS1;LAT* |
| Lipid and atherosclerosis | 0.0301382151955936 | *RAP1B;SELP;PPP3R1;ROCK2;CCL5;CALM3;CD36;SOD2;PTK2;BCL2L1* |
| Gap junction | 0.0333529492999705 | *TUBA1C;GUCY1B1;TUBB1;GNAS;PDGFA;TUBA4A* |
| Oxytocin signaling pathway | 0.0372999958527224 | *PPP3R1;CDKN1A;GUCY1B1;ROCK2;GNAS;CALM3;MYL9;MYLK* |
| Transcriptional misregulation in cancer | 0.0401792267025728 | *PTCRA;LYL1;CDKN1A;MEIS1;MAX;PDGFA;PBX1;PTK2;BCL2L1* |
| Pathogenic Escherichia coli infection | 0.0417891338908558 | *CLDN5;TUBA1C;CTTN;ROCK2;ARPC1B;TUBB1;NCK2;ARPC5;TUBA4A* |
| Fc gamma R-mediated phagocytosis | 0.0417891338908558 | *ARPC1B;PRKCD;ASAP1;ARPC5;ASAP2;LAT* |
| PI3K-Akt signaling pathway | 0.0417891338908558 | *YWHAE;CDKN1A;ITGB5;ITGB3;ITGA2B;PDGFA;GNG11;YWHAZ;THBS1;PTK2;CCND3;BCL2L1;YWHAH* |
| Hippo signaling pathway | 0.0417891338908558 | *MOB1B;YWHAE;CCND3;TGFB1;YWHAZ;BMP6;SAV1;YWHAH* |
| Renal cell carcinoma | 0.0417891338908558 | *RAP1B;CDKN1A;EGLN3;TGFB1;RBX1* |
| Hematopoietic cell lineage | 0.0417891338908558 | *GP9;GP1BB;ITGB3;ITGA2B;CD9;CD36* |
| Pathways in cancer | 0.0417891338908558 | *CDKN1A;EGLN3;TGFB1;GSTO1;MAX;ROCK2;NCOA4;ITGA2B;PDGFA;RASGRP2;GNG11;PTK2;RBX1;CCND3;GNAS;CALM3;BCL2L1* |
| cGMP-PKG | 0.0417891338908 | *PPP3R1;GUCY1B1;ROCK2;VDAC3;CALM3;PDE5A;MYL9;MYLK* |

| | | |
|---|---|---|
| signaling pathway | 558 | |
| Proteoglycans in cancer | 0.0417891338908558 | *CDKN1A;TGFB1;CTTN;ITGB5;CAV2;ROCK2;ITGB3;THBS1;PTK2* |
| Amoebiasis | 0.0417891338908558 | *ARG2;TGFB1;ACTN1;GNAS;VCL;PTK2* |
| Pancreatic secretion | 0.0417891338908558 | *RAP1B;PLA2G12A;CA2;GNAS;RAB27B;RAB11A* |
| Parkinson disease | 0.0471479958417193 | *TUBA1C;NDUFA6;UBA7;TUBB1;VDAC3;GNAS;CALM3;TUBA4A;BCL2L1;SNCA* |
| | | Cluster 2 |
| Antigen processing and presentation | 0.04576678699363543 | *CD74;HLA-DRA* |
| | | Cluster 3 |
| African trypanosomiasis | 0.00199893220474141 | *HBB;HBA2;HBA1* |
| Porphyrin and chlorophyll metabolism | 0.00209671706165243 | *ALAS2;FECH;BLVRB* |
| Mitophagy | 0.00614008897198106 | *BNIP3L;UBB;BCL2L1* |
