## Supplementary Table 11 for "Heterogeneity of circulating epithelial cells in breast cancer at single-cell resolution: identifying tumor and hybrid cells"

Supplementary Table 11. Upregulated genes in aneuploid cells of $CD45^+$ CEC cluster 1

| Genes | LogFC | Adjusted P-value |
|---|---|---|
| *TCF4* | 2.7155466105313 | 3.33383282410324E-12 |
| *BCL11A* | 1.87397597313578 | 2.48126254379445E-11 |
| *CCDC50* | 2.17995502474521 | 3.68229012184876E-11 |
| *APP* | 2.32432034115058 | 8.17895439953916E-11 |
| *CD74* | 4.53470251366221 | 1.34279891644327E-10 |
| *CCDC88A* | 1.94283783128059 | 1.79237021476123E-10 |
| *MPEG1* | 1.68224660137512 | 7.42047546955445E-10 |
| *UGCG* | 2.22424321702439 | 2.13964757630882E-09 |
| *JCHAIN* | 3.01632889654711 | 4.38232907019852E-09 |
| *PLD4* | 2.40892293614982 | 5.06822026150439E-09 |
| *MT-ND2* | 1.49270129481321 | 8.50335287373588E-09 |
| *MT-ND1* | 1.60760978846218 | 1.21263193310508E-08 |
| *SCN9A* | 1.08597322310985 | 1.5464737344375E-08 |
| *HLA-DRA* | 3.68097062673331 | 1.72486906053853E-08 |
| *MZB1* | 1.54855889216854 | 8.66743514987941E-08 |
| *ITM2C* | 2.50369092862307 | 8.69243909053794E-08 |
| *HLA-DRB1* | 2.92474510555789 | 9.08809420562747E-08 |
| *HLA-DPB1* | 1.90155221748821 | 9.44153492981926E-08 |
| *GRN* | 1.99683272092132 | 1.00688479039002E-07 |
| *HERPUD1* | 1.9129601636663 | 1.00765114008556E-07 |
| *CYBB* | 1.58874798228424 | 1.01975307711638E-07 |
| *CYB561A3* | 1.54855889216854 | 1.19394028680932E-07 |
| *IRF8* | 1.81696491623143 | 1.30741125418777E-07 |
| *TXNDC5* | 0.974464907892862 | 1.59283241193237E-07 |
| *PTPRE* | 1.64570698546267 | 1.97570861747211E-07 |
| *COBLL1* | 1.28192404762831 | 2.47527871692412E-07 |
| *SEL1L3* | 0.987288948250446 | 4.94925165716899E-07 |
| *DAB2* | 0.86754970397635 | 4.9492516571691E-07 |
| *CST3* | 2.35940280030603 | 5.47358070566832E-07 |
| *SULF2* | 1.27436669645023 | 5.95011744154015E-07 |
| *IRF4* | 1.99683272092132 | 7.52541386548574E-07 |
| *NPC2* | 2.03747470541866 | 0.0000010558824027562 |
| *CORO1C* | 0.736965594166206 | 1.50694202746816E-06 |
| *TSPAN13* | 1.12130629622213 | 1.50694202746817E-06 |
| *SERPINF1* | 1.25986712675511 | 1.89443651667126E-06 |
| *IRF7* | 1.16929991692608 | 2.41447938709945E-06 |
| *CD2AP* | 1.3175826246114 | 2.42083463166665E-06 |
| *PSAP* | 1.79716037608495 | 2.87303225986675E-06 |
| *MT-ND5* | 1.72691192950228 | 0.000003558210518694 |
| *MT-ND4* | 1.49343246001082 | 3.93505785281081E-06 |
| *LILRA4* | 1.78129397062593 | 4.20179461557245E-06 |
| *SMPD3* | 1.23319917629303 | 4.50012425450818E-06 |
| *KRT5* | 0.767022828022386 | 4.50012425450825E-06 |
| *ELMSAN1* | 1.03275229517626 | 5.26639619827063E-06 |
| *CLN8* | 1.21117286996303 | 7.80599722667471E-06 |
| *CTSB* | 1.50733622944203 | 8.62077307041631E-06 |
| *HSP90B1* | 2.03032037623064 | 0.0000119998720641588 |
| *SPIB* | 1.16745149253371 | 0.0000127282222471901 |
| *LGMN* | 0.974464907892862 | 0.0000131912670169393 |
| *FLNB* | 1.09784732339814 | 0.0000131912670169394 |
| *NGLY1* | 0.956278622570682 | 0.0000172494693830457 |
| *SOX4* | 1.87484219654271 | 0.0000176094480489777 |
| *RRBP1* | 1.39039049278971 | 0.0000289436175719827 |
| *CLEC4C* | 0.839535327806754 | 0.0000379863742067757 |
| *CD36* | 0.721698837512896 | 0.0000379863742067759 |
| *ALOX5AP* | 2.17219980743188 | 0.0000430165788596537 |
| *HLA-DPA1* | 2.80059258744084 | 0.0000519548425230399 |

| | | |
|---|---|---|
| *SELENOS* | 1.83929144393484 | 0.0000590272905236241 |
| *MAP1A* | 0.990573730449635 | 0.0000692039628995203 |
| *MAN2B1* | 1.27313572791062 | 0.0000698668508286453 |
| *MT-CO1* | 1.87528394697051 | 0.0000709637235629876 |
| *HLA-DQB1* | 1.30651722005393 | 0.000103137443201692 |
| *AFF3* | 0.767022828022386 | 0.000107542057401624 |
| *CXCR3* | 1.13289427049735 | 0.000107542057401624 |
| *TGFBI* | 1.2000470405739 | 0.000169306862743007 |
| *ZEB2* | 1.13755000681985 | 0.000170677305379202 |
| *DERL3* | 1.13701037896958 | 0.000178016151534634 |
| *PPP1R14B* | 1.18106555904751 | 0.000229379800542707 |
| *CD4* | 1.48183277836773 | 0.000294299101396376 |
| *IL3RA* | 1.396635706505 | 0.000307202020180566 |
| *STX7* | 0.882296289345537 | 0.000324421284349755 |
| *XIST* | 1.56293619439116 | 0.000411753172079099 |
| *PHACTR1* | 0.917804474756046 | 0.000473225026115447 |
| *ARID3A* | 0.831584206688478 | 0.000512926716111969 |
| *MAPKAPK2* | 1.34242972170231 | 0.000560950871375145 |
| *SEC61B* | 1.41598632867239 | 0.000590241498022053 |
| *LINC02812* | 0.610053481683987 | 0.000821415797411332 |
| *WDFY4* | 0.658963082164933 | 0.000821415797411332 |
| *GAS6* | 0.921997487998727 | 0.000821415797411349 |
| *PFKFB2* | 0.736965594166206 | 0.000821415797411349 |
| *FCHSD2* | 0.939499545855365 | 0.00117210188600097 |
| *SCT* | 1.02087819488797 | 0.00148514764648592 |
| *MT-ATP6* | 1.26884014134294 | 0.00163503731037237 |
| *JAML* | 0.981369603533513 | 0.00215779265909269 |
| *TMBIM6* | 1.22104787786458 | 0.0021602125880483 |
| *ATG101* | 0.610053481683987 | 0.0022192231203221 |
| *LYN* | 1.07243837767349 | 0.0022727785812643 |
| *RGS2* | 1.40288143697291 | 0.00304903144122583 |
| *RAB11FIP1* | 1.11547721741994 | 0.00317305235524494 |
| *NIBAN3* | 1.07929488111329 | 0.00320559015757982 |
| *RUNX2* | 1.03275229517626 | 0.00326212349022374 |
| *GNAS* | 1.25267162535091 | 0.00333591504743566 |
| *CD68* | 0.959995952740946 | 0.00335899756083926 |
| *SFT2D2* | 1.41728371638653 | 0.00359310430690834 |
| *DUSP5* | 0.845523397646352 | 0.00362168746900521 |
| *SCAF11* | 1.11622881683307 | 0.00398331538082454 |
| *ANKRD12* | 1.00254637211571 | 0.00422730558245808 |
| *TRAF4* | 1.12628397237652 | 0.00444983608105309 |
| *SF1* | 1.07844648647497 | 0.00445852764441158 |
| *ITPR2* | 0.822420328486691 | 0.00470618049525059 |
| *SFPQ* | 1.05664211615662 | 0.00487850423385656 |
| *CTSZ* | 1.18947779886371 | 0.00520659991913554 |
| *SCARB2* | 0.942837831280588 | 0.00540270709613944 |
| *SCAMP4* | 0.721698837512896 | 0.00591099536672562 |
| *PECAM1* | 0.489038080722621 | 0.00591099536672566 |
| *AC007381.1* | 0.57650092197296 | 0.00591099536672571 |
| *CBFA2T3* | 0.542149417182183 | 0.00591099536672571 |
| *EPHA2* | 0.626541604472555 | 0.00591099536672571 |
| *TNFRSF21* | 0.70626879694329 | 0.00591099536672571 |
| *LILRB4* | 0.873004134068459 | 0.0060174308389875 |
| *PLAC8* | 1.29183951649339 | 0.00638311436976682 |
| *MTDH* | 1.13344376394938 | 0.00638465107595745 |
| *DDX21* | 0.310332193738098 | 0.00642429716036056 |
| *ERP29* | 1.31916887550335 | 0.00675964424519174 |
| *MED13L* | 1.10260034818375 | 0.00684389637828128 |
| *CSF2RB* | 0.934904971778115 | 0.00726959295088663 |
| *MDM4* | 1.15100365104908 | 0.00856865261387948 |

| | | |
|---|---|---|
| *MT-CO3* | 1.49276348737718 | 0.00941138474343373 |
| *SYK* | 0.809889135589159 | 0.00983248358088755 |
| *NCF1* | 0.888308264143148 | 0.0101870316922239 |
| *PLP2* | 1.26390263999027 | 0.0109673034545636 |
| *MTRNR2L8* | 1.00625597480794 | 0.0136340274297955 |
| *LRRC26* | 0.690671941892271 | 0.0155321501323986 |
| *SLC9A7* | 0.57650092197296 | 0.0155321501323986 |
| *TLR9* | 0.542149417182183 | 0.0155321501323987 |
| *LHFPL2* | 0.610053481683987 | 0.0155321501323988 |
| *HNRNPU* | 1.04745879399128 | 0.0165757316998324 |
| *MT-CO2* | 1.44538983922401 | 0.0179997638759394 |
| *HLA-DMA* | 1.05823326558546 | 0.0215114428376331 |
| *OFD1* | 1.00505322772629 | 0.022492870400027 |
| *OGT* | 1.02901312667047 | 0.0225451752182263 |
| *TM9SF2* | 0.993926989745906 | 0.0243308792715839 |
| *ADPGK* | 0.760634500059839 | 0.0257792443189949 |
| *CTSS* | 1.43233432275988 | 0.0262295224320107 |
| *CYTH4* | 0.835840844249012 | 0.0267738234946188 |
| *IGKC* | 5.03886736432653 | 0.027136957368775 |
| *ANKRD11* | 1.39241785753399 | 0.0276236946846469 |
| *ST3GAL2* | 0.620817110471657 | 0.0290673015381141 |
| *TACC1* | 0.863249566542725 | 0.0300012936057018 |
| *CUX1* | 0.731371577692983 | 0.0311586917259528 |
| *PNISR* | 1.20641766128543 | 0.033074652452135 |
| *SLC35E1* | 0.588027175353988 | 0.0338588200556696 |
| *ZMYM2* | 1.04192061044985 | 0.03603091355085 |
| *HNRNPH1* | 0.832321215665369 | 0.0382529793891622 |
| *GZMB* | 1.98405411620001 | 0.0390631420025266 |
| *CHD9* | 0.888475542830899 | 0.0394748128659458 |
| *SNX9* | 0.760226440365168 | 0.0400828256069684 |
| *SHTN1* | 0.452512204697507 | 0.0402906708727327 |
| *CDH1* | 0.524661990453342 | 0.040290670872733 |
| *CYP46A1* | 0.810966175609983 | 0.040290670872733 |
| *MCOLN2* | 0.593374740537356 | 0.040290670872733 |
| *NECTIN1* | 0.524661990453342 | 0.040290670872733 |
| *ST14* | 0.658963082164933 | 0.040290670872733 |
| *STAT2* | 0.542149417182183 | 0.040290670872733 |
| *COL26A1* | 0.506959988719883 | 0.0402906708727333 |
| *PRXL2A* | 0.626541604472555 | 0.0402906708727333 |
| *SEMA7A* | 0.559427408614019 | 0.0402906708727333 |
| *PHEX* | 0.674904626033955 | 0.0402906708727336 |
| *SPINT2* | 1.10066853190586 | 0.0404487734206491 |
| *ZDHHC17* | 0.760226440365168 | 0.0499787481142625 |
