## Supplementary Table 12 for "Heterogeneity of circulating epithelial cells in breast cancer at single-cell resolution: identifying tumor and hybrid cells"

Supplementary Table 12. Signaling pathways enriched in aneuploid cells of CD45$^{+}$ CEC cluster 1

| Term | Adjusted P-value | Genes |
|---|---|---|
| Antigen processing and presentation | 5.56747226901495E-09 | *CD74;HLA-DMA;CD4;HLA-DPB1;HLA-DRA;CTSS;HLA-DRB1;CTSB;HLA-DPA1;HLA-DQB1;LGMN* |
| Phagosome | 3.28711140329495E-07 | *HLA-DMA;NCF1;STX7;HLA-DPB1;CYBB;HLA-DRA;CD36;SEC61B;HLA-DRB1;CTSS;HLA-DPA1;HLA-DQB1* |
| Allograft rejection | 1.17395764144599E-06 | *HLA-DMA;HLA-DPB1;GZMB;HLA-DRA;HLA-DRB1;HLA-DPA1;HLA-DQB1* |
| Graft-versus-host disease | 1.73926197525878E-06 | *HLA-DMA;HLA-DPB1;GZMB;HLA-DRA;HLA-DRB1;HLA-DPA1;HLA-DQB1* |
| Type I diabetes mellitus | 1.73926197525878E-06 | *HLA-DMA;HLA-DPB1;GZMB;HLA-DRA;HLA-DRB1;HLA-DPA1;HLA-DQB1* |
| Hematopoietic cell lineage | 0.00000312413436307 34 | *HLA-DMA;CD4;IL3RA;HLA-DPB1;HLA-DRA;CD36;HLA-DRB1;HLA-DPA1;HLA-DQB1* |
| Asthma | 0.0000039771922619735 | *HLA-DMA;HLA-DPB1;HLA-DRA;HLA-DRB1;HLA-DPA1;HLA-DQB1* |
| Leishmaniasis | 0.0000043225950675283 | *HLA-DMA;NCF1;HLA-DPB1;CYBB;HLA-DRA;HLA-DRB1;HLA-DPA1;HLA-DQB1* |
| Autoimmune thyroid disease | 0.0000043225950675283 | *HLA-DMA;HLA-DPB1;GZMB;HLA-DRA;HLA-DRB1;HLA-DPA1;HLA-DQB1* |
| Cell adhesion molecules | 5.98536464148725E-06 | *HLA-DMA;CD4;CDH1;PECAM1;HLA-DPB1;HLA-DRA;HLA-DRB1;HLA-DPA1;HLA-DQB1;NECTIN1* |
| Lysosome | 0.0000154072010033558 | *SCARB2;NPC2;PSAP;CTSZ;MAN2B1;CD68;CTSS;LGMN;CTSB* |
| Tuberculosis | 0.0000300315877813976 | *CD74;HLA-DMA;SYK;TLR9;HLA-DPB1;HLA-DRA;HLA-DRB1;CTSS;HLA-DPA1;HLA-DQB1* |
| Intestinal immune network for IgA production | 0.0000313332795625093 | *HLA-DMA;HLA-DPB1;HLA-DRA;HLA-DRB1;HLA-DPA1;HLA-DQB1* |
| Th17 cell differentiation | 0.0000313332795625093 | *HLA-DMA;CD4;IRF4;HLA-DPB1;HLA-DRA;HLA-DRB1;HLA-DPA1;HLA-DQB1* |
| Epstein-Barr virus infection | 0.000067376118243074 | *LYN;HLA-DMA;SYK;STAT2;IRF7;HLA-DPB1;HLA-DRA;HLA-DRB1;HLA-DPA1;HLA-DQB1* |
| Viral myocarditis | 0.00009771375738932 | *HLA-DMA;HLA-DPB1;HLA-DRA;HLA-DRB1;HLA-DPA1;HLA-DQB1* |
| Th1 and Th2 cell differentiation | 0.000100200826463222 | *HLA-DMA;CD4;HLA-DPB1;HLA-DRA;HLA-DRB1;HLA-DPA1;HLA-DQB1* |
| Protein processing in endoplasmic reticulum | 0.000102574776121652 | *SELENOS;DERL3;NGLY1;ERP29;RRBP1;SEC61B;HSP90B1;TXNDC5;HERPUD1* |
| Inflammatory bowel | 0.000131362573390027 | *HLA-DMA;HLA-DPB1;HLA-DRA;HLA-DRB1;HLA-DPA1;HLA-DQB1* |

| | | |
|---|---|---|
| disease | | |
| Influenza A | 0.000699361217202784 | *HLA-DMA;STAT2;IRF7;HLA-DPB1;HLA-DRA;HLA-DRB1;HLA-DPA1;HLA-DQB1* |
| Rheumatoid arthritis | 0.000912401324412658 | *HLA-DMA;HLA-DPB1;HLA-DRA;HLA-DRB1;HLA-DPA1;HLA-DQB1* |
| Staphylococcus aureus infection | 0.000980078108872886 | *HLA-DMA;HLA-DPB1;HLA-DRA;HLA-DRB1;HLA-DPA1;HLA-DQB1* |
| Apoptosis | 0.0012041393815796 | *CTSZ;IL3RA;ITPR2;GZMB;CSF2RB;CTSS;CTSB* |
| Toxoplasmosis | 0.00221223514197386 | *HLA-DMA;HLA-DPB1;HLA-DRA;HLA-DRB1;HLA-DPA1;HLA-DQB1* |
| Herpes simplex virus 1 infection | 0.00513415615329036 | *CD74;HLA-DMA;SYK;STAT2;IRF7;TLR9;HLA-DPB1;HLA-DRA;HLA-DRB1;HLA-DPA1;HLA-DQB1;NECTIN1* |
| Systemic lupus erythematosus | 0.00551632235991588 | *HLA-DMA;HLA-DPB1;HLA-DRA;HLA-DRB1;HLA-DPA1;HLA-DQB1* |
| Human T-cell leukemia virus 1 infection | 0.0134775587185687 | *HLA-DMA;CD4;HLA-DPB1;HLA-DRA;HLA-DRB1;HLA-DPA1;HLA-DQB1* |
| Osteoclast differentiation | 0.0238785930908273 | *NCF1;SYK;STAT2;LILRB4;LILRA4* |
| B cell receptor signaling pathway | 0.0262469431786747 | *LYN;SYK;LILRB4;LILRA4* |
| Kaposi sarcoma-associated herpesvirus infection | 0.02895928647686 | *LYN;SYK;STAT2;MAPKAPK2;IRF7;ITPR2* |
| Sphingolipid metabolism | 0.0426572108642896 | *SMPD3;UGCG;PSAP* |
| Malaria | 0.0437074194417158 | *PECAM1;TLR9;CD36* |
| Lipid and atherosclerosis | 0.043991560847709 | *LYN;NCF1;IRF7;CYBB;CD36;HSP90B1* |
