## Supplementary Table 13 for "Heterogeneity of circulating epithelial cells in breast cancer at single-cell resolution: identifying tumor and hybrid cells"

Supplementary Table 13. The spectrum of CNAs in diploid cells of CD45$^{-}$ CEC and CD45$^{+}$ CEC clusters

| CEC type, clusters | CNA | State | Chr | Start | End |
|---|---|---|---|---|---|
| CD45$^{-}$ CECs, cluster 1 | chr1-region_46 | 2 | chr1 | 112709994 | 151909637 |
| CD45$^{-}$ CECs, cluster 1 | chr3-region_50 | 4 | chr3 | 187721377 | 197956610 |
| CD45$^{-}$ CECs, cluster 1 | chr5-region_53 | 2 | chr5 | 31400494 | 78360507 |
| CD45$^{-}$ CECs, cluster 1 | chr6-region_55 | 4 | chr6 | 291630 | 15663058 |
| CD45$^{-}$ CECs, cluster 1 | chr6-region_57 | 4 | chr6 | 38675925 | 100881372 |
| CD45$^{-}$ CECs, cluster 1 | chr9-region_62 | 4 | chr9 | 98069275 | 123104866 |
| CD45$^{-}$ CECs, cluster 1 | chr9-region_64 | 4 | chr9 | 136862119 | 137870016 |
| CD45$^{-}$ CECs, cluster 1 | chr11-region_67 | 4 | chr11 | 59208510 | 64284704 |
| CD45$^{-}$ CECs, cluster 1 | chr11-region_69 | 2 | chr11 | 93661682 | 134253370 |
| CD45$^{-}$ CECs, cluster 1 | chr12-region_71 | 4 | chr12 | 104117086 | 133032952 |
| CD45$^{-}$ CECs, cluster 1 | chr14-region_74 | 4 | chr14 | 103333544 | 105488947 |
| CD45$^{-}$ CECs, cluster 1 | chr17-region_78 | 4 | chr17 | 4004445 | 31382116 |
| CD45$^{-}$ CECs, cluster 1 | chr21-region_83 | 4 | chr21 | 5079294 | 36294274 |
| CD45$^{-}$ CECs, cluster 2 | chr2-region_132 | 2 | chr2 | 189560590 | 203306026 |
| CD45$^{-}$ CECs, cluster 2 | chr5-region_137 | 2 | chr5 | 9035033 | 69131069 |
| CD45$^{-}$ CECs, cluster 2 | chr6-region_139 | 4 | chr6 | 291630 | 13287843 |
| CD45$^{-}$ CECs, cluster 2 | chr9-region_144 | 4 | chr9 | 99216425 | 120842951 |
| CD45$^{-}$ CECs, cluster 2 | chr9-region_146 | 4 | chr9 | 136440096 | 137870016 |
| CD45$^{-}$ CECs, cluster 2 | chr11-region_149 | 4 | chr11 | 47465933 | 64917211 |
| CD45$^{-}$ CECs, cluster 2 | chr11-region_151 | 2 | chr11 | 95066919 | 134253370 |
| CD45$^{-}$ CECs, cluster 2 | chr12-region_153 | 4 | chr12 | 104117086 | 133032952 |
| CD45$^{-}$ CECs, cluster 2 | chr17-region_159 | 4 | chr17 | 3662895 | 20319026 |
| CD45$^{-}$ CECs, cluster 2 | chr19-region_163 | 4 | chr19 | 531760 | 5720572 |

| | | | | | |
|---|---|---|---|---|---|
| CD45$^+$ CECs, cluster 1 | chr2-region_3 | 2 | chr2 | 188974373 | 203226378 |
| CD45$^+$ CECs, cluster 1 | chr5-region_8 | 2 | chr5 | 14581792 | 68301821 |
| CD45$^+$ CECs, cluster 1 | chr5-region_10 | 4 | chr5 | 139341587 | 151224093 |
| CD45$^+$ CECs, cluster 1 | chr6-region_12 | 4 | chr6 | 291630 | 16295549 |
| CD45$^+$ CECs, cluster 1 | chr6-region_14 | 4 | chr6 | 42879616 | 87399749 |
| CD45$^+$ CECs, cluster 1 | chr9-region_19 | 4 | chr9 | 98056732 | 114646422 |
| CD45$^+$ CECs, cluster 1 | chr9-region_21 | 4 | chr9 | 136862119 | 137870016 |
| CD45$^+$ CECs, cluster 1 | chr11-region_23 | 2 | chr11 | 167784 | 5243657 |
| CD45$^+$ CECs, cluster 1 | chr11-region_25 | 4 | chr11 | 46617527 | 64321811 |
| CD45$^+$ CECs, cluster 1 | chr11-region_27 | 2 | chr11 | 93784227 | 134253370 |
| CD45$^+$ CECs, cluster 1 | chr12-region_29 | 2 | chr12 | 8032716 | 27014434 |
| CD45$^+$ CECs, cluster 1 | chr12-region_31 | 4 | chr12 | 101696947 | 133032952 |
| CD45$^+$ CECs, cluster 1 | chr16-region_36 | 4 | chr16 | 85899162 | 89968060 |
| CD45$^+$ CECs, cluster 1 | chr17-region_38 | 4 | chr17 | 4433940 | 29294148 |
| CD45$^+$ CECs, cluster 2 | chr2-region_88 | 2 | chr2 | 187464230 | 207167267 |
| CD45$^+$ CECs, cluster 2 | chr5-region_93 | 2 | chr5 | 14704800 | 69178245 |
| CD45$^+$ CECs, cluster 2 | chr6-region_95 | 4 | chr6 | 291630 | 15663058 |
| CD45$^+$ CECs, cluster 2 | chr6-region_97 | 4 | chr6 | 38675925 | 81752774 |
| CD45$^+$ CECs, cluster 2 | chr9-region_102 | 4 | chr9 | 98056732 | 120894896 |
| CD45$^+$ CECs, cluster 2 | chr9-region_104 | 4 | chr9 | 136712572 | 137870016 |
| CD45$^+$ CECs, cluster 2 | chr11-region_106 | 2 | chr11 | 167784 | 4093210 |
| CD45$^+$ CECs, cluster 2 | chr11-region_108 | 4 | chr11 | 46617527 | 64778786 |
| CD45$^+$ CECs, cluster 2 | chr11-region_110 | 2 | chr11 | 94415570 | 134253370 |
| CD45$^+$ CECs, cluster 2 | chr12-region_112 | 2 | chr12 | 7919230 | 27325959 |

| | | | | | |
|---|---|---|---|---|---|
| CD45$^+$ CECs, cluster 2 | chr12-region_114 | 4 | chr12 | 102012840 | 133032952 |
| CD45$^+$ CECs, cluster 2 | chr16-region_119 | 4 | chr16 | 87696485 | 89968060 |
| CD45$^+$ CECs, cluster 2 | chr17-region_121 | 4 | chr17 | 3923870 | 18107970 |
| CD45$^+$ CECs, cluster 2 | chr19-region_125 | 4 | chr19 | 1609290 | 5153598 |
| CD45$^+$ CECs, cluster 2 | chr2-region_132 | 2 | chr2 | 189560590 | 203306026 |
| CD45$^+$ CECs, cluster 2 | chr5-region_137 | 2 | chr5 | 9035033 | 69131069 |
| CD45$^+$ CECs, cluster 2 | chr6-region_139 | 4 | chr6 | 291630 | 13287843 |
| CD45$^+$ CECs, cluster 2 | chr9-region_144 | 4 | chr9 | 99216425 | 120842951 |
| CD45$^+$ CECs, cluster 2 | chr9-region_146 | 4 | chr9 | 136440096 | 137870016 |
| CD45$^+$ CECs, cluster 2 | chr11-region_149 | 4 | chr11 | 47465933 | 64917211 |
| CD45$^+$ CECs, cluster 2 | chr11-region_151 | 2 | chr11 | 95066919 | 134253370 |
| CD45$^+$ CECs, cluster 2 | chr12-region_153 | 4 | chr12 | 104117086 | 133032952 |
| CD45$^+$ CECs, cluster 2 | chr17-region_159 | 4 | chr17 | 3662895 | 20319026 |
| CD45$^+$ CECs, cluster 2 | chr19-region_163 | 4 | chr19 | 531760 | 5720572 |
| CD45$^+$ CECs, cluster 3 | chr5-region_173 | 2 | chr5 | 5420664 | 60522120 |
| CD45$^+$ CECs, cluster 3 | chr5-region_175 | 4 | chr5 | 177511577 | 181272307 |
| CD45$^+$ CECs, cluster 3 | chr11-region_182 | 4 | chr11 | 18468336 | 61829318 |
| CD45$^+$ CECs, cluster 3 | chr11-region_184 | 2 | chr11 | 111878935 | 134253370 |
| CD45$^+$ CECs, cluster 3 | chr12-region_186 | 4 | chr12 | 51281038 | 57750219 |
| CD45$^+$ CECs, cluster 3 | chr12-region_188 | 4 | chr12 | 111443485 | 122896127 |
| CD45$^+$ CECs, cluster 3 | chr17-region_195 | 4 | chr17 | 5432777 | 16777881 |
