## Supplementary Table 14 for "Heterogeneity of circulating epithelial cells in breast cancer at single-cell resolution: identifying tumor and hybrid cells"

Supplementary Table 14. Upregulated genes in diploid cells of CD45$^{-}$ CEC clusters

| Genes | Cluster 1 | | Genes | Cluster 2 | |
|---|---|---|---|---|---|
| | LogFC | Adjusted P-value | | LogFC | Adjusted P-value |
| *CST3* | 2.78951155585969 | 1.9239977529654E-10 | *CD74* | 4.4382802916372 | 2.22407761672012E-28 |
| *CCDC88A* | 2.11547721741994 | 2.12464129123277E-10 | *GNAS* | 1.92136020636745 | 5.46503741402967E-26 |
| *CD74* | 4.15862816992776 | 3.41822668981688E-10 | *HLA-DRA* | 3.55601271437792 | 2.24594189501257E-25 |
| *HLA-DPA1* | 2.47217073046798 | 1.65935192916629E-09 | *PPP1R14B* | 2.94660030541049 | 2.92749432066336E-24 |
| *CCDC50* | 2.59319998491472 | 2.70668327496194E-09 | *TCF4* | 2.46769532443141 | 6.30753125011233E-24 |
| *TCF4* | 2.58940840575235 | 5.1255019155193E-09 | *CST3* | 2.5624771515785 | 1.03841778494416E-23 |
| *SNHG5* | 1.48471102708566 | 6.22275393197281E-09 | *ITM2C* | 2.64154602908752 | 1.39690480443144E-23 |
| *SEC61B* | 2.36845795858981 | 8.39317368509339E-09 | *NPC2* | 2.53969828761834 | 1.61771208457373E-22 |
| *APP* | 1.90977598665209 | 1.34129921832772E-08 | *HLA-DRB1* | 3.05865752526824 | 7.74083300212633E-22 |
| *ITM2C* | 2.1782129727679 | 2.34958831560998E-08 | *HLA-DPA1* | 2.8444653197445 | 1.31640913467306E-21 |
| *HLA-DRA* | 3.05618948012519 | 4.0528140929865E-08 | *PLD4* | 2.3445109520184 | 1.7719009265542E-21 |
| *GAS6* | 1.22239242133645 | 4.36403133419496E-08 | *SEC61B* | 2.19479967008134 | 8.0556634892717E-21 |
| *TXNDC5* | 1.3536369546147 | 4.36403133419496E-08 | *JCHAIN* | 2.99766455894723 | 1.10870657266469E-20 |
| *AFF3* | 1.22239242133645 | 4.36403133419506E-08 | *SERPINF1* | 2.33761359207173 | 1.18635520745152E-20 |
| *PLD4* | 2.23095443483987 | 7.00897388019666E-08 | *HLA-DPB1* | 2.27324615655631 | 2.37755411677512E-20 |
| *JCHAIN* | 2.78205348369474 | 9.43299249964079E-08 | *GRN* | 1.89264913398116 | 2.96460128384625E-20 |
| *PPP1R14B* | 2.59610305832636 | 1.29949941335842E-07 | *TAGLN2* | 1.80950669643555 | 3.36743369037209E-20 |
| *IRF8* | 2.33522491750082 | 3.07362293542686E-07 | *IRF7* | 2.1868618506567 | 4.82233978852478E-20 |
| *GNAS* | 1.97617818000839 | 3.43937935272568E-07 | *APP* | 1.79397288374604 | 4.97605330561333E-20 |
| *SCT* | 1.83106916079358 | 3.50476633846863E-07 | *PSAP* | 1.74745634464258 | 8.05479970695975E-20 |
| *MAPKAPK2* | 1.67060448860033 | 3.72793959611719E-07 | *CCDC50* | 2.12862259347686 | 2.87774114174187E-19 |
| *TAGLN2* | 1.8184954796628 | 4.73400659924974E-07 | *UGCG* | 2.1276647876054 | 3.06144227012981E-19 |
| *NPC2* | 2.25843517126198 | 5.07111612166532E-07 | *PTMS* | 1.29235497950402 | 7.11674140596626E-19 |
| *BCL11A* | 2.3670159844159 | 5.29390724470722E-07 | *LILRA4* | 2.02773432277827 | 9.72048385779859E-19 |
| *SEL1L3* | 1.32192809488736 | 5.64512125186823E-07 | *IRF8* | 1.99213788222703 | 1.69648540857171E-18 |
| *ATG101* | 1.07800251200127 | 5.64512125186825E-07 | *C12orf75* | 2.24246610371166 | 3.44989473280743E-18 |
| *HLA-DPB1* | 1.83333398863844 | 7.02686279634413E-07 | *ALOX5AP* | 2.13991143573013 | 5.82829462437266E-18 |
| *TRAF4* | 1.93261316027013 | 7.18455312878862E-07 | *FTH1* | 1.77551519970231 | 5.97376094878555E-18 |
| *C12orf75* | 1.96533686799208 | 1.72109745045749E-06 | *RPLP0* | 1.7146256243844 | 3.56936047910317E-17 |
| *SERPINF1* | 1.85244281158614 | 3.82236296098015E-06 | *CCDC88A* | 1.79242858263866 | 3.99855370270661E-17 |
| *MT-CO1* | 2.0688139594038 | 3.87330625689037E-06 | *TXNDC5* | 1.16573720447888 | 7.73251594481327E-17 |
| *REPIN1* | 1.12997678711505 | 4.23659921746477E-06 | *HERPUD1* | 1.97220012864679 | 8.38474166328494E-17 |
| *PSAP* | 1.83516482015431 | 4.62735054870871E-06 | *PPIA* | 1.60269510011617 | 1.04342167386526E-16 |
| *LILRA4* | 1.76755391399963 | 4.76875132085007E-06 | *CYB561A3* | 1.57614794550461 | 1.47461786383313E-16 |
| *PABPC1* | 1.51323672526694 | 5.99285744705566E-06 | *STMN1* | 1.53492528277809 | 2.48181749733593E-16 |

| | | | | | |
|---|---|---|---|---|---|
| *BCL7A* | 0.959358015502654 | 0.0000063157950053872 | *SPIB* | 1.26690996294655 | 2.84401667170926E-16 |
| *GRN* | 1.48987273220143 | 6.47804736515545E-06 | *IGKC* | 3.18343141591013 | 5.0190951048343E-16 |
| *CTSB* | 1.37512103405296 | 6.48280371648268E-06 | *CYBA* | 2.00391505000338 | 5.76028622067019E-16 |
| *PTPRE* | 2.09480364915519 | 7.41054300794285E-06 | *SOX4* | 1.81604733688447 | 5.7627946177674E-16 |
| *PPIA* | 1.66801824044871 | 0.0000077936683346965 | *HLA-DMA* | 1.45637059065105 | 6.03631617153421E-16 |
| *PLAC8* | 1.88045718775574 | 7.80891685377363E-06 | *MT-CO1* | 1.64716607164538 | 8.84703934879552E-16 |
| *HSP90B1* | 1.45703411905623 | 8.13055726715328E-06 | *SRP14* | 1.60121300594501 | 1.01739510885295E-15 |
| *RAB11FIP1* | 2.32493058304889 | 0.0000100363969073827 | *GAS6* | 1.14886338591448 | 1.17068838894947E-15 |
| *HERPUD1* | 2.36869256222328 | 0.0000102602764041613 | *TXN* | 2.05729614633141 | 1.51318025673601E-15 |
| *UGCG* | 2.10960065108327 | 0.000014382742571551 | *PLAC8* | 1.81051410954092 | 1.52758918655248E-15 |
| *CD68* | 1.28854192639282 | 0.0000148704625794382 | *ATP5F1E* | 1.57486596059351 | 1.7015296727292E-15 |
| *CTSZ* | 1.48160511621783 | 0.0000161996001686301 | *BCL11A* | 1.72707484464456 | 2.86004860201427E-15 |
| *DERL3* | 1.27774864631881 | 0.0000180676807895319 | *FCGRT* | 1.37669233544532 | 3.07106280216144E-15 |
| *HLA-DQB1* | 1.81122014885963 | 0.0000217269738937075 | *MZB1* | 1.54813356933501 | 3.71431479527532E-15 |
| *GRB2* | 1.40178140257658 | 0.0000221525226405447 | *CXXC5* | 1.4051467497572 | 4.19028141687546E-15 |
| *SNHG29* | 1.2735805104014 | 0.0000245552467524272 | *MT-ND1* | 1.30712141288594 | 5.97557410543073E-15 |
| *SLC15A4* | 1.56704059272389 | 0.000027239728709362 | *SCT* | 1.58526994772091 | 5.99807708026994E-15 |
| *MAN2B1* | 1.43020981090309 | 0.0000284298881839256 | *OAZ1* | 1.32690320608869 | 6.32816365063845E-15 |
| *CSF2RB* | 1.34994247105696 | 0.0000407766782465548 | *CXCR3* | 1.35418229389199 | 6.72167835832808E-15 |
| *DUSP5* | 1.58940840575235 | 0.0000471591865227499 | *SERF2* | 1.35156975643384 | 8.99382470262576E-15 |
| *KRT5* | 0.959358015502654 | 0.00006250573596841 | *EIF4A1* | 1.40394667063 | 1.22616226044644E-14 |
| *TSPAN13* | 0.959358015502654 | 0.000062505735968411 | *H3F3A* | 1.14330822939038 | 1.25382475106315E-14 |
| *TGFBI* | 1.51986747249927 | 0.0000648385010766247 | *IL3RA* | 1.34680276352639 | 1.3495311488326E-14 |
| *IGKC* | 2.03189058714696 | 0.0000685980457077348 | *TSPAN13* | 1.18241594562551 | 1.58190890651125E-14 |
| *CYB561A3* | 1.60189934993555 | 0.000070012693603295 | *NME2* | 1.52577290457171 | 1.6178686993265E-14 |
| *IRF7* | 1.89197117577278 | 0.0000759163804359829 | *HLA-DQB1* | 1.51497067640245 | 1.78253669090857E-14 |
| *HNRNPK* | 1.19709098297359 | 0.0000768832457946207 | *LRRC26* | 1.12029423371771 | 3.68026926464034E-14 |
| *MT-ND4* | 1.81062263589151 | 0.0000787723870147998 | *ERP29* | 1.64794929627725 | 3.81903085149415E-14 |
| *ATP5F1E* | 1.60451529814256 | 0.000084668725103289 | *CTSB* | 1.28922548794958 | 4.48109637165443E-14 |
| *RRBP1* | 1.67628585643231 | 0.0000866748844305913 | *PARK7* | 1.60702215375313 | 6.26086777043363E-14 |
| *MT-ND1* | 1.74784060063879 | 0.0000907110634983819 | *SPCS1* | 1.69977929467298 | 7.04418479362216E-14 |
| *MT-ND5* | 1.92283213947754 | 0.000102918926167278 | *HLA-DQA1* | 1.54002851969795 | 7.80583884108017E-14 |
| *RPL6* | 1.74527931663711 | 0.000108321899027756 | *SMPD3* | 1.18241594562551 | 8.46674795442416E-14 |
| *SIVA1* | 1.24837148791728 | 0.000111732953415427 | *VEGFB* | 1.50292196544367 | 8.96048404038357E-14 |
| *UBE2J1* | 1.63612490465938 | 0.000119477421801248 | *MAN2B1* | 1.19825141287665 | 9.39290134683521E-14 |
| *CD2AP* | 1.55604980880592 | 0.000122410797674635 | *TRAF4* | 1.48638659771737 | 1.75666630993375E-13 |
| *GABARAP* | 1.76585512659005 | 0.000127546959611234 | *MAPKAPK2* | 1.58000193981942 | 1.78639972913088E-13 |
| *EIF4G2* | 1.56139402958275 | 0.000133245036269593 | *SSR4* | 1.57542713857435 | 1.81752048055561E-13 |

| | | | | | |
|---|---|---|---|---|---|
| *TXN* | 1.80240212908655 | 0.000136149363376346 | *DERL3* | 1.32371915041335 | 1.8692182025954E-13 |
| *IRF4* | 1.78924898509895 | 0.000138210215285898 | *SEL1L3* | 1.07337318633022 | 1.92677426760944E-13 |
| *STMN1* | 1.67997561558138 | 0.000211600365556099 | *IRF4* | 1.75149831858378 | 2.56708385536965E-13 |
| *RPS11* | 1.70739247878286 | 0.000225592179819291 | *NCF1* | 1.26311015972816 | 2.69907645120304E-13 |
| *NACA* | 1.55815747176791 | 0.000289758017385205 | *GABARAP* | 1.54836895766004 | 2.75132581871506E-13 |
| *ANKRD12* | 1.48987273220143 | 0.000336814532639003 | *IGFLR1* | 1.28267242610668 | 3.20816839003347E-13 |
| *RNASET2* | 1.39676332780995 | 0.000406821570361085 | *RNASET2* | 1.4957971070991 | 3.36386409437849E-13 |
| *HLA-DRB1* | 2.14349159358953 | 0.000407564955593587 | *BRI3* | 1.34482239939815 | 4.39319818561038E-13 |
| *MT-CO2* | 1.56988493646774 | 0.000431576870559488 | *TPM2* | 1.68058994228207 | 4.81936194663658E-13 |
| *SLC7A5* | 0.915636638073336 | 0.000434105640214975 | *MAP1A* | 0.952662605140814 | 6.15459706138637E-13 |
| *MAP1A* | 1.12997678711505 | 0.000482914302112962 | *TUBA1B* | 1.62000756839228 | 6.40247850929198E-13 |
| *SSR4* | 1.67504656071269 | 0.000534998698196731 | *PTCRA* | 1.29859367048377 | 6.50557113124882E-13 |
| *LRRC26* | 0.874469117916141 | 0.000556761365591199 | *SNHG29* | 1.61841525934098 | 7.15222639586493E-13 |
| *SMPD3* | 1.11547721741994 | 0.000556761365591199 | *RPS8* | 1.74224953444735 | 7.2815931443857E-13 |
| *TLR7* | 0.959358015502654 | 0.000556761365591199 | *EIF4A3* | 1.05882200056237 | 7.8236524882568E-13 |
| *CIB2* | 0.830074998557688 | 0.0005567613655912 | *ATG101* | 1.03715299841747 | 9.66973726493626E-13 |
| *KCTD12* | 0.874469117916141 | 0.0005567613655912 | *SELENOS* | 1.48882176839882 | 1.08527808315209E-12 |
| *PACSIN1* | 0.736965594166206 | 0.0005567613655912 | *GZMB* | 2.23683971852514 | 1.46972842842841E-12 |
| *SCN9A* | 0.959358015502654 | 0.000556761365591202 | *EEF1A1* | 1.49989099808382 | 1.74477012097583E-12 |
| *PTMS* | 1.09345091108994 | 0.000570731173028285 | *UNC93B1* | 1.18225093418293 | 2.60175890737595E-12 |
| *ERP29* | 1.4036846944633 | 0.000582562675480432 | *RPL15* | 1.36807138028004 | 3.01900150707116E-12 |
| *RBM3* | 1.20937735630142 | 0.000612525922453347 | *SPINT2* | 1.13869456819619 | 3.22452502347751E-12 |
| *MPEG1* | 1.45108024920438 | 0.000661502691605603 | *MYL6* | 1.18077167818857 | 4.66358493755851E-12 |
| *VAMP8* | 1.58283575182886 | 0.000705861411928665 | *CD68* | 1.20835599355725 | 4.7188691041161E-12 |
| *FCER1G* | 1.55698692805997 | 0.000776107405715852 | *RACK1* | 1.43047605125834 | 5.09360719936936E-12 |
| *YWHAZ* | 1.24238932990216 | 0.000778735547486349 | *PLP2* | 1.47139039939104 | 6.52535964365244E-12 |
| *TPM2* | 1.85244281158614 | 0.000799706499520279 | *VIM* | 1.74122998312995 | 8.61662749362373E-12 |
| *SNX3* | 1.23095443483987 | 0.000845023361980021 | *RNF130* | 1.19705564402171 | 1.15400478144965E-11 |
| *NIBAN3* | 1.08690806522317 | 0.000883920959045502 | *CD4* | 1.43904071211313 | 1.35588504798336E-11 |
| *RPLP0* | 1.58940840575235 | 0.000962274585786854 | *HSP90B1* | 1.5339476630917 | 1.44951748399663E-11 |
| *ANKRD11* | 1.62503231548307 | 0.000988844522538187 | *MPEG1* | 1.33299696400136 | 1.47163789653911E-11 |
| *ALOX5AP* | 1.56864984558555 | 0.000990082921419879 | *EEF2* | 1.30498258783176 | 1.70407691191264E-11 |
| *MZB1* | 1.23577145113765 | 0.00118137933191047 | *LGMN* | 0.902702798645085 | 2.16460783269161E-11 |
| *H3F3B* | 1.31346651613917 | 0.00120531535355464 | *RPL6* | 1.70142166822871 | 2.17832860698366E-11 |
| *RPS8* | 1.63698310441158 | 0.00122733396696521 | *OSTC* | 1.10394623963023 | 2.71955799775869E-11 |
| *CYBB* | 1.26748031086499 | 0.00126762524447661 | *PTPRE* | 1.48110385523798 | 2.9882564192203E-11 |
| *SLC38A2* | 1.51457317282976 | 0.00130597598921201 | *CNPY3* | 1.20662510547813 | 3.93755472561891E-11 |
| *IL3RA* | 1.31956882201344 | 0.00155499706835713 | *ARPC1B* | 1.17362478428899 | 4.39754545010632E-11 |

| | | | | | |
|---|---|---|---|---|---|
| *ATP6V0B* | 1.01845276302507 | 0.00165681508870603 | *CTSZ* | 1.19598658503178 | 4.41658533104673E-11 |
| *PARK7* | 1.25986712675511 | 0.00171477011377282 | *DAB2* | 1.0124909441832 | 4.60099273277718E-11 |
| *FCGRT* | 1.10147035942606 | 0.00180130906331174 | *WDFY4* | 0.746966986323705 | 4.60099273277718E-11 |
| *VEGFB* | 1.53051471669878 | 0.00212232335398105 | *CLEC4C* | 0.791134569599274 | 4.60099273277724E-11 |
| *MT-ND2* | 1.60980362650342 | 0.00239354831908197 | *GSTP1* | 1.87856386910692 | 5.99957196437911E-11 |
| *COBLL1* | 1.40883616011053 | 0.00242255599662484 | *ACTB* | 1.11636041000392 | 6.64961101582148E-11 |
| *SELENOS* | 1.67628585643231 | 0.00250683123731124 | *CAT* | 1.09072111833437 | 7.39950703912558E-11 |
| *SERBP1* | 1.15669988014645 | 0.00264732256105001 | *MT-ND4* | 1.25094274811888 | 7.98096119660015E-11 |
| *MLF2* | 0.953371720436923 | 0.00265736018564436 | *NIBAN3* | 1.07809960760591 | 8.79630684176999E-11 |
| *IGFLR1* | 0.913843356250285 | 0.00273988727023816 | *SCN9A* | 0.87560913776281 | 9.69442010916447E-11 |
| *RASSF2* | 1.01750205785664 | 0.00287386350352088 | *BCL7A* | 0.942412508447056 | 9.69442010916461E-11 |
| *HLA-DQA1* | 1.02932057367022 | 0.00294277827881661 | *PRXL2A* | 0.841010476238843 | 9.69442010916461E-11 |
| *CD4* | 1.26748031086499 | 0.00332613075733401 | *RASD1* | 1.35418229389199 | 9.69442010916461E-11 |
| *H3F3A* | 1.44124737860169 | 0.00340508606290889 | *RNASE6* | 0.895976935456772 | 9.69442010916461E-11 |
| *SH2B3* | 1.30812229536233 | 0.00375785369506744 | *KRT5* | 0.98105843634654 | 9.69442010916475E-11 |
| *CNPY3* | 1.14194942878113 | 0.00407899662513602 | *HINT1* | 1.33708760869318 | 1.01256826214091E-10 |
| *GGA2* | 0.977973693670001 | 0.00423707958191816 | *HIGD1A* | 0.883830787183846 | 1.04346949139381E-10 |
| *CLN8* | 0.937331709172655 | 0.00431336826671852 | *DPYSL2* | 0.993431620988148 | 1.04844897296513E-10 |
| *FLNB* | 0.874469117916141 | 0.00452600588132334 | *VAMP8* | 1.38987148541828 | 1.0487248826834E-10 |
| *IDH3A* | 0.736965594166206 | 0.00452600588132334 | *SEC61G* | 1.29906520109357 | 1.83415550564713E-10 |
| *NEK8* | 0.917537839808027 | 0.00452600588132334 | *IGHM* | 1.40860804799986 | 1.88096468021389E-10 |
| *CD302* | 0.736965594166206 | 0.00452600588132335 | *FLNB* | 0.769219793170833 | 2.0252852680208E-10 |
| *CD36* | 0.68805599368526 | 0.00452600588132335 | *AFF3* | 0.798366138830349 | 2.02528526802086E-10 |
| *FCER1A* | 0.830074998557688 | 0.00452600588132335 | *HMGA1* | 1.16063712107687 | 2.80211977439362E-10 |
| *SHTN1* | 0.68805599368526 | 0.00452600588132335 | *MS4A6A* | 1.1381081919195 | 2.91001420307214E-10 |
| *TNFRSF21* | 0.874469117916141 | 0.00452600588132335 | *CLN8* | 1.02122689162389 | 3.35873010710826E-10 |
| *PIK3CG* | 1.07800251200127 | 0.0045260058813234 | *SNHG7* | 1.14849596071426 | 3.54724346520094E-10 |
| *RABGAP1L* | 1.16655140201421 | 0.00455966459409859 | *NACA* | 1.64719769686411 | 3.83648097606162E-10 |
| *LILRB4* | 0.977973693670001 | 0.004661962646164 | *HLA-DMB* | 0.853582831432811 | 3.95525360547372E-10 |
| *STRBP* | 0.977973693670001 | 0.00466196264616405 | *PPIB* | 1.38569177691999 | 4.01972181358396E-10 |
| *COX6A1* | 1.15583495503608 | 0.00498700752153172 | *MYBL2* | 0.948925815405731 | 4.19601764030209E-10 |
| *ETF1* | 0.934904971778115 | 0.00513191322322734 | *RPS6KA4* | 0.924569762843344 | 4.75021489516549E-10 |
| *SPIB* | 1.26748031086499 | 0.00518122220354735 | *C12orf45* | 0.999531820524566 | 4.75896830316982E-10 |
| *RELT* | 0.808048692227689 | 0.00537389818813835 | *FCER1G* | 1.60934233312111 | 6.00764846629927E-10 |
| *EIF4A1* | 1.81255008446263 | 0.00581002923302713 | *CFL1* | 1.18241594562551 | 6.14705349565107E-10 |
| *HSP90AB1* | 1.3197748183072 | 0.0058554488287873 | *TMED2* | 0.92742632852321 | 6.61326317754961E-10 |
| *PPP1R9B* | 0.714939287836207 | 0.00607324306484568 | *SFT2D2* | 1.20507465391473 | 7.64223369607927E-10 |
| *MT-CO3* | 1.49983451658072 | 0.00616615214531233 | *CYBB* | 1.2332602319816 | 8.18790282712318E-10 |

| | | | | | |
|---|---|---|---|---|---|
| *PRKCB* | 1.4211034999782 | 0.00633382418698923 | *TNFRSF21* | 0.805561640234553 | 8.62312197337228E-10 |
| *RPS16* | 1.15034767726699 | 0.00648844587009821 | *REPIN1* | 0.839842774806509 | 8.9382627752981E-10 |
| *CYBA* | 1.90019594303451 | 0.00972305374074932 | *UBC* | 1.1974390845664 | 1.07365701703103E-09 |
| *NGLY1* | 1.03428113457196 | 0.0104195434340597 | *CAPG* | 1.03560107523209 | 1.0868853537085E-09 |
| *SPINT2* | 1.03428113457196 | 0.0104195434340598 | *PHACTR1* | 0.981153291209711 | 1.16079411765672E-09 |
| *DDX21* | 1.09667962196581 | 0.0113126623985457 | *GNG5* | 1.0911478880582 | 1.20963841336371E-09 |
| *DCK* | 1.10828171601573 | 0.0123327977766215 | *MT-CO2* | 1.1866658424911 | 1.23184284016117E-09 |
| *RPS23* | 1.46969871551418 | 0.0143101005476855 | *NGLY1* | 0.943678585791049 | 1.26481801735545E-09 |
| *RACK1* | 1.12850389771618 | 0.0144056743276608 | *ARPC3* | 1.27482281249604 | 1.30308955571849E-09 |
| *NCF1* | 1.13623577758673 | 0.0153674744136695 | *RUNX2* | 0.864201980434913 | 1.3861837375421E-09 |
| *RPL37A* | 1.43572873449991 | 0.0154031761507342 | *ATP5F1B* | 1.07499814806487 | 1.45274465860476E-09 |
| *COPE* | 1.03199223191685 | 0.0168892812993696 | *P4HB* | 1.12173620789162 | 1.57176019216452E-09 |
| *SPCS1* | 1.41187022020016 | 0.0186825423025193 | *SNHG5* | 1.17085397445631 | 1.7356458464322E-09 |
| *RILPL2* | 0.852442811586142 | 0.0189786056602095 | *RPL22* | 1.22465012036563 | 1.73616056397756E-09 |
| *CXXC5* | 1.27774864631881 | 0.0193129610817072 | *PECAM1* | 0.81272143180742 | 1.75814291626932E-09 |
| *SLC25A3* | 1.24238932990216 | 0.0202545268336772 | *MIF4GD* | 0.754422791242752 | 1.75814291626935E-09 |
| *UBA52* | 1.23251663861016 | 0.0214201333750928 | *ST14* | 0.776561768512001 | 1.75814291626935E-09 |
| *ATP5PF* | 1.01471424048502 | 0.0219586369326247 | *PHB* | 1.15669988014645 | 1.92499083255158E-09 |
| *HNRNPH1* | 0.950290134984287 | 0.0221320681660857 | *LAPTM4A* | 0.937690098204653 | 2.18563695479673E-09 |
| *HNRNPA2B1* | 1.24148510233204 | 0.0222464399122058 | *SIVA1* | 1.1335213928897 | 2.19746661378274E-09 |
| *SOX4* | 1.83281400483721 | 0.0242352229477666 | *YBX1* | 1.24776124824059 | 2.27120181851476E-09 |
| *SFT2D2* | 1.21922514225776 | 0.0254523932527769 | *RNASEK* | 1.06708961654597 | 2.28770351344425E-09 |
| *STX7* | 0.948020471568373 | 0.0264749105935649 | *JAML* | 0.905204437976413 | 2.47046569791084E-09 |
| *RNF130* | 1.19454878841113 | 0.0271226692260783 | *RPS24* | 1.47460158876292 | 2.78633424227132E-09 |
| *RPL35* | 1.29722765425904 | 0.0278119362108857 | *RPL37A* | 1.3242865820985 | 2.90189249999476E-09 |
| *OFD1* | 1.62362412109026 | 0.0281913712053726 | *GPX4* | 0.883987372091026 | 2.94158543396009E-09 |
| *PHB* | 1.04006981476191 | 0.0282134595760467 | *DUSP5* | 1.00284836230903 | 3.11260464510067E-09 |
| *HMGN2* | 1.32351434577931 | 0.0282963648286052 | *RPL10A* | 1.26987388509347 | 3.20468285810699E-09 |
| *TMA7* | 1.09541956507868 | 0.029665457686203 | *MT-CO3* | 1.25425730782347 | 3.27894079699219E-09 |
| *PPIB* | 1.15052416451914 | 0.0317859612961813 | *SNX3* | 1.14035188605897 | 3.32270727657247E-09 |
| *PRXL2A* | 0.874469117916141 | 0.0339718098052036 | *RPL4* | 1.26351644162055 | 3.41537069893626E-09 |
| *CBFA2T3* | 0.68805599368526 | 0.0339718098052038 | *TGFBI* | 1.05519920549583 | 3.41915936961715E-09 |
| *CLEC4C* | 0.53051471669878 | 0.0339718098052038 | *LILRB4* | 0.873950629126773 | 3.53120955768621E-09 |
| *CXCR3* | 0.917537839808027 | 0.0339718098052039 | *FCHSD2* | 0.867193262951464 | 3.65993006167142E-09 |
| *ELL* | 0.53051471669878 | 0.0339718098052039 | *ATP5F1A* | 1.03747470541866 | 3.8383997016163E-09 |
| *EPHB1* | 0.874469117916141 | 0.0339718098052039 | *MT-ND2* | 1.02849271452495 | 5.44587346575598E-09 |
| *KLF4* | 0.736965594166206 | 0.0339718098052039 | *UBE2J1* | 1.19050962767185 | 5.92139826229959E-09 |
| *MYCL* | 0.53051471669878 | 0.0339718098052039 | *RGS2* | 1.61524776519355 | 6.18093741096257E-09 |

| | | |
|---|---|---|
| *RNASE6* | 0.830074998557688 | 0.0339718098052039 |
| *FABP5* | 0.68805599368526 | 0.033971809805204 |
| *LGMN* | 0.736965594166206 | 0.0339718098052041 |
| *CBX4* | 1.17018310951007 | 0.0347964569357991 |
| *GSTP1* | 2.27152714392769 | 0.0355007083103987 |
| *UBC* | 1.30495501628365 | 0.0364362345804425 |
| *SGSM3* | 0.644334616255942 | 0.0395329244251275 |
| *COX6B1* | 0.913843356250285 | 0.0410450830161437 |
| *EIF2AK4* | 0.873816462378709 | 0.0459665938136045 |

| | | |
|---|---|---|
| *RPS28* | 1.48346177925581 | 6.69717862052221E-09 |
| *GRAMD1B* | 0.873950629126773 | 6.84447511995499E-09 |
| *FABP5* | 0.754422791242752 | 7.14252046520974E-09 |
| *CALM2* | 1.11594876344175 | 7.35687362416934E-09 |
| *ATP6V0B* | 1.13220871275035 | 7.43538074763432E-09 |
| *SULF2* | 0.818984169908844 | 7.54104815754724E-09 |
| *CERS6* | 0.987305518191307 | 7.81484255781189E-09 |
| *RBM3* | 1.06850160103929 | 9.46647192575457E-09 |
| *TYROBP* | 1.36504188181143 | 1.00724924505836E-08 |
| *RPS23* | 1.49225951831497 | 1.10061395695599E-08 |
| *CHCHD2* | 1.25289562892886 | 1.11148715587652E-08 |
| *SLC25A6* | 1.17376515021674 | 1.1697339626593E-08 |
| *EEF1G* | 1.14571683059762 | 1.18212977880469E-08 |
| *GSN* | 0.80723216561925 | 1.19020265838771E-08 |
| *TMA7* | 1.1024278129242 | 1.30302569962599E-08 |
| *COMT* | 0.83161054110941 | 1.32602785472725E-08 |
| *RPN2* | 1.00041261119712 | 1.33812567710253E-08 |
| *SSR3* | 0.917959464722517 | 1.38347404450247E-08 |
| *LYN* | 1.09000031140411 | 1.39475247945991E-08 |
| *BLNK* | 0.724365557386573 | 1.42369773887861E-08 |
| *LINC02812* | 0.739472449776781 | 1.42369773887861E-08 |
| *ALDH2* | 0.739472449776781 | 1.42369773887863E-08 |
| *GNG7* | 0.646363045385299 | 1.42369773887863E-08 |
| *RPL29* | 1.35941223983441 | 1.81654477311093E-08 |
| *COX6B1* | 1.09210765860533 | 1.84279200987557E-08 |
| *COBLL1* | 1.03192942617298 | 1.88980845659992E-08 |
| *SYK* | 0.918143493683911 | 2.07722341150563E-08 |
| *PLEK* | 1.4602460104404 | 2.16001191700199E-08 |
| *RPS5* | 1.36121244287967 | 2.16188095791049E-08 |
| *RPS3A* | 1.47495265131513 | 2.19533623871502E-08 |
| *RPS9* | 1.25404562260654 | 2.23338009955547E-08 |
| *SCARB2* | 0.880484759303855 | 2.26948658951239E-08 |
| *RPS2* | 1.33792993923751 | 2.29934053642872E-08 |
| *EIF2AK4* | 0.618583211825003 | 2.6007285386172E-08 |
| *OPN3* | 0.693668760163656 | 2.81750096063245E-08 |
| *IDH3A* | 0.605721060887954 | 2.81750096063248E-08 |
| *SLC15A4* | 1.23132554610646 | 2.83766901872312E-08 |
| *ZNF706* | 0.921948405887341 | 3.1755868380086E-08 |

| | | |
|---|---|---|
| *RASSF2* | 0.790695125477421 | 3.31196417922256E-08 |
| *IRF2BP2* | 1.1213004141892 | 3.48943912727326E-08 |
| *AP1S2* | 0.866334430676341 | 3.50871308607093E-08 |
| *TM9SF2* | 0.860020656744139 | 3.5957310315458E-08 |
| *RPL22L1* | 0.975965068158083 | 3.65388460532006E-08 |
| *A1BG* | 0.815033201970829 | 3.79365625310086E-08 |
| *RPS11* | 1.24228023730572 | 3.9494003405985E-08 |
| *RPS7* | 1.27046670613126 | 4.08879568480242E-08 |
| *H3F3B* | 1.09578927020906 | 4.31809435608299E-08 |
| *SLC7A5* | 0.852255558027454 | 4.59410011636933E-08 |
| *DSTN* | 1.15647155142224 | 4.73158322361328E-08 |
| *CD2AP* | 0.978332082701999 | 5.02885038342584E-08 |
| *SUB1* | 1.10036486822329 | 5.09954182106301E-08 |
| *CBFA2T3* | 0.693668760163656 | 5.53685997063633E-08 |
| *CORO1C* | 0.662304589254321 | 5.53685997063649E-08 |
| *RPL7A* | 1.292578255066 | 5.68817323770694E-08 |
| *SSR1* | 0.899942539434023 | 5.93216216581487E-08 |
| *POMP* | 0.880160229993997 | 6.12490481356117E-08 |
| *THEMIS2* | 0.802482247302581 | 6.42697770649438E-08 |
| *PSMB3* | 0.897495362162801 | 6.94703124801417E-08 |
| *SLC25A3* | 1.13053904741721 | 7.22512097839027E-08 |
| *LAMTOR1* | 0.868755466721747 | 7.32473367960297E-08 |
| *BCAP31* | 0.787603358870317 | 7.32919432910085E-08 |
| *RPS4X* | 1.38395162026137 | 8.1402187249116E-08 |
| *HMGN1* | 1.05018575716822 | 9.27906400163624E-08 |
| *RPS13* | 1.39069289869731 | 1.01938024759886E-07 |
| *EPHB1* | 0.605721060887954 | 1.08065536217977E-07 |
| *LINC00996* | 0.693668760163656 | 1.08065536217977E-07 |
| *CDCA7L* | 0.580774703757723 | 1.08065536217979E-07 |
| *EPHA2* | 0.670209787339667 | 1.08065536217979E-07 |
| *GPM6B* | 0.746966986323705 | 1.08065536217979E-07 |
| *DNASE1L3* | 0.678071905112638 | 1.0806553621798E-07 |
| *NHP2* | 0.891316787233678 | 1.16829908020205E-07 |
| *COX6A1* | 1.02316834668863 | 1.17825826468078E-07 |
| *MEF2C* | 0.969645147918637 | 1.20643713665735E-07 |
| *CBX4* | 0.803062925531748 | 1.25480080126301E-07 |
| *FKBP1A* | 0.799808113004256 | 1.26141669628754E-07 |
| *ATP13A2* | 0.747193486840834 | 1.27819072841625E-07 |

| | | |
|---|---|---|
| *RPLP1* | 1.24997637723346 | 1.55391357483511E-07 |
| *ELMSAN1* | 0.718771540913349 | 1.71611366427654E-07 |
| *TACC1* | 0.756103893256091 | 1.8970353679769E-07 |
| *CD36* | 0.701404408943261 | 2.0950965048098E-07 |
| *PFKFB2* | 0.731938777091373 | 2.0950965048098E-07 |
| *MYL12A* | 1.13880972125056 | 2.2967124949443E-07 |
| *RPL18A* | 1.25882692755426 | 2.83894863600655E-07 |
| *HMGN2* | 1.05782509044032 | 3.11445844539943E-07 |
| *RPS19* | 1.22133336822162 | 3.49271038775866E-07 |
| *SEC31A* | 0.733271110608465 | 3.50224099388243E-07 |
| *ATP6V0C* | 0.895363935642625 | 3.60424161961227E-07 |
| *CANX* | 0.910703648719718 | 3.76829484185142E-07 |
| *SH2B3* | 0.942412508447056 | 3.82552505219215E-07 |
| *PABPC1* | 1.12619034863523 | 3.83580600833114E-07 |
| *PIK3CG* | 0.580774703757723 | 4.03534794946622E-07 |
| *CIB2* | 0.622115499138622 | 4.03534794946634E-07 |
| *RPL37* | 1.31663543656796 | 4.29287234125076E-07 |
| *ARHGAP27* | 0.724940679993706 | 5.1959624446377E-07 |
| *FERMT3* | 0.88990530159353 | 5.32653937875148E-07 |
| *TMEM258* | 0.988805413931508 | 5.53913371755574E-07 |
| *SLC25A5* | 1.18461341086368 | 5.63410393404694E-07 |
| *OFD1* | 1.1028771806403 | 5.71536084640918E-07 |
| *CCDC69* | 0.966954692709848 | 6.18416045692974E-07 |
| *CD164* | 1.01206689233841 | 6.62627908745424E-07 |
| *SCAF11* | 0.915271060514143 | 7.21139039829417E-07 |
| *TUBB* | 0.957564733679603 | 7.5670430204948E-07 |
| *NDUFA4* | 1.09625930187579 | 8.17471698147661E-07 |
| *ATP6V1G1* | 0.984232684141683 | 8.25443087534721E-07 |
| *PAXX* | 1.01693684921439 | 8.2937622261964E-07 |
| *CSF2RB* | 0.796774052914623 | 8.48790764104139E-07 |
| *RRBP1* | 1.06036375335765 | 8.53446942389633E-07 |
| *GUK1* | 0.876218283950368 | 1.04789025010251E-06 |
| *SELENOH* | 0.949467265981007 | 1.11232315772746E-06 |
| *MYL12B* | 0.966660054693734 | 1.21780739616288E-06 |
| *ATP5MG* | 1.02812032814268 | 1.44343468806654E-06 |
| *PACSIN1* | 0.555389385337809 | 1.46883900656974E-06 |
| *RPL10* | 1.17871364149308 | 1.48224596863384E-06 |
| *CUX1* | 0.796774052914623 | 1.54217889941215E-06 |

| | | |
|---|---|---|
| *ATP5MPL* | 0.9865996986856 | 1.59563485662517E-06 |
| *BAG1* | 0.842010611489219 | 1.71870151899936E-06 |
| *SMIM3* | 0.754535462182002 | 1.74924104029797E-06 |
| *GAPT* | 0.732840391082683 | 0.0000017559839695679 |
| *RPL35A* | 1.02907930014497 | 2.03026234991696E-06 |
| *RBIS* | 0.813601690494977 | 2.03312910327776E-06 |
| *ST3GAL2* | 0.656045598782639 | 2.09978067168883E-06 |
| *RPL35* | 1.13107407247095 | 2.20037125756864E-06 |
| *UQCRQ* | 0.773522067852871 | 2.20293678058282E-06 |
| *ATP5MD* | 0.970728419791936 | 2.35666901869898E-06 |
| *TIMM13* | 0.782598822972033 | 2.67340502851788E-06 |
| *HM13* | 0.740778647277406 | 2.69293039532769E-06 |
| *SUMO2* | 0.958697592034546 | 2.69538424858986E-06 |
| *FCER1A* | 0.65435583613883 | 2.77661523151958E-06 |
| *SCAMP4* | 0.685891409571937 | 2.77661523151958E-06 |
| *SLC9A7* | 0.512061953673708 | 2.77661523151962E-06 |
| *PRKCD* | 0.572362463941523 | 2.77661523151966E-06 |
| *TLR7* | 0.52083216330144 | 2.77661523151966E-06 |
| *BTF3* | 1.07938991718202 | 2.81900294339509E-06 |
| *CTSS* | 1.04696843948965 | 2.82779138066142E-06 |
| *N4BP2* | 0.809145977583592 | 0.0000029170949710413 |
| *NADK* | 0.69674523458335 | 2.92480836908036E-06 |
| *MDH2* | 0.871138905296047 | 0.0000029260319363471 |
| *DAD1* | 0.755816475177465 | 2.96537317788133E-06 |
| *RPL11* | 1.23601492342595 | 2.99672058618439E-06 |
| *RPL36A* | 1.04468314971921 | 3.07945656542273E-06 |
| *CTSC* | 1.03485875721165 | 3.18608259517729E-06 |
| *FTL* | 0.891123279750202 | 3.22450978614625E-06 |
| *SNRNP25* | 0.695751072347463 | 0.0000034936800890224 |
| *RHOA* | 0.970935861528226 | 3.61716271780158E-06 |
| *LAIR1* | 0.671642453833657 | 3.77231052756005E-06 |
| *ERCC1* | 0.782598822972033 | 3.90513866683938E-06 |
| *SRSF9* | 0.908681651134506 | 3.93148682024191E-06 |
| *SEPTIN11* | 0.665377423303944 | 3.97049169427611E-06 |
| *OST4* | 0.835695110116772 | 4.10947939379859E-06 |
| *ST6GALNAC4* | 0.616299419989882 | 4.15105440409488E-06 |
| *RPL19* | 1.25018966833415 | 4.36905479736536E-06 |
| *UVRAG* | 0.528641086512205 | 4.38843356855194E-06 |

| | | |
|---|---|---|
| *RAB11FIP1* | 1.19479967008134 | 4.44001307201767E-06 |
| *HNRNPA1* | 0.863891534274932 | 4.64901551617267E-06 |
| *CLTC* | 0.668266147493038 | 4.65072870453364E-06 |
| *COX5A* | 1.01181851581358 | 4.72543892160147E-06 |
| *SHD* | 0.630243380022023 | 5.21756735973875E-06 |
| *LAMP5* | 0.538214241529677 | 5.21756735973883E-06 |
| *SH3TC1* | 0.546827371834385 | 5.21756735973883E-06 |
| *SHTN1* | 0.646363045385299 | 5.21756735973883E-06 |
| *COX7A2L* | 0.834903724994959 | 5.21775389696256E-06 |
| *ATP5MC3* | 1.14293043016074 | 5.83528791357662E-06 |
| *PFN1* | 0.716268089687977 | 6.09841857081998E-06 |
| *MLF2* | 0.775712437386793 | 6.15609340644526E-06 |
| *ARID3A* | 0.783535333904555 | 6.33323492408723E-06 |
| *RPL3* | 1.18181005762631 | 6.47900447865477E-06 |
| *MT-ND5* | 0.800068315916115 | 6.51466364308009E-06 |
| *COMMD6* | 1.21448271577989 | 6.65639310918054E-06 |
| *ILF2* | 0.727570360720489 | 7.26105315922452E-06 |
| *ABHD15* | 0.656045598782639 | 0.0000073864138382357 |
| *IFNAR2* | 0.640278282924322 | 7.65019904326305E-06 |
| *USF2* | 0.922945563357538 | 8.09517778069008E-06 |
| *NUDT1* | 0.666843748869488 | 8.54177039234191E-06 |
| *BRK1* | 0.838881298569063 | 8.68786701176781E-06 |
| *PRDX6* | 0.761840262805235 | 8.89352072643227E-06 |
| *JTB* | 0.883402242686149 | 8.95634909426632E-06 |
| *RPS26* | 1.00017013247618 | 9.07885088859235E-06 |
| *COPE* | 0.899870413697681 | 9.27378452009442E-06 |
| *RPL8* | 1.04059290238596 | 9.54254698358342E-06 |
| *LIME1* | 0.984232684141683 | 9.67405829647339E-06 |
| *RPL5* | 1.15908081559597 | 9.67551779064162E-06 |
| *AC007381.1* | 0.597453444904353 | 9.74743193332961E-06 |
| *CMTM3* | 0.741081702638438 | 0.0000100917449894056 |
| *SEC63* | 0.647507587326241 | 0.0000101004900350752 |
| *HMGB1* | 0.973021292185059 | 0.0000103164223801978 |
| *MT-ND3* | 0.892324793995732 | 0.0000106738293169937 |
| *EEF1B2* | 1.19697297898379 | 0.0000107998573932155 |
| *ZDHHC17* | 0.666843748869488 | 0.0000108041541257282 |
| *PIM3* | 0.959939770839739 | 0.0000109127786663481 |
| *NDUFB8* | 0.836134045152549 | 0.0000120264604298451 |

| | | |
|---|---|---|
| *PPM1G* | 0.781205587672167 | 0.0000127357713600351 |
| *PAIP1* | 0.776339832500351 | 0.0000130196442952144 |
| *UBB* | 0.867068597145304 | 0.0000134747029043915 |
| *GGA2* | 0.687072494403263 | 0.000013521791946854 |
| *FAU* | 1.03975225354419 | 0.0000140959667324238 |
| *SNX9* | 0.754750838018916 | 0.0000146396258869254 |
| *GNAI2* | 0.915271060514143 | 0.0000147382879440573 |
| *KRTCAP2* | 0.843370148301834 | 0.0000148659507802851 |
| *UBA52* | 1.06714632142728 | 0.0000152430957591346 |
| *CHD9* | 0.822980557619139 | 0.000015427979122044 |
| *RPL27* | 0.984833431616901 | 0.0000158384689640314 |
| *HSP90AB1* | 1.07405320285939 | 0.0000161807617219803 |
| *GNA15* | 0.668266147493038 | 0.0000163905066111699 |
| *HNRNPA2B1* | 0.887848147717341 | 0.0000181020446620231 |
| *CD302* | 0.503238103286702 | 0.0000181067198505624 |
| *SLC3A2* | 0.780455940972582 | 0.0000183457921553261 |
| *SYNGR2* | 0.644003772511377 | 0.0000191139618338498 |
| *RPL12* | 1.15558822119044 | 0.0000193555936435802 |
| *ACTG1* | 1.1458900696004 | 0.0000196342532051214 |
| *SCAMP2* | 0.625023573174862 | 0.0000249057067524275 |
| *EIF4G2* | 0.833990048561071 | 0.0000251132992594605 |
| *SLC35E1* | 0.687072494403263 | 0.0000252454318669229 |
| *SCAMP5* | 0.663865103241938 | 0.0000256689477018665 |
| *SUMO3* | 0.768693933846298 | 0.0000259588140180131 |
| *DBNL* | 0.690405124762286 | 0.0000266866902396787 |
| *PTMA* | 0.72769204039339 | 0.0000268142872427801 |
| *C15orf39* | 0.575427138574354 | 0.0000269627066356744 |
| *TMBIM6* | 0.895857610776923 | 0.0000272846698795654 |
| *GNB1* | 0.708728926345673 | 0.0000280497594778484 |
| *REEP5* | 0.69770992538552 | 0.000029217599485671 |
| *TMEM59* | 0.822212880784717 | 0.0000304644257843308 |
| *RPS12* | 1.10557750513893 | 0.000032171106457484 |
| *PPM1J* | 0.52954938040255 | 0.0000334482378514031 |
| *RPL41* | 0.905859219910155 | 0.0000339973351907344 |
| *ETF1* | 0.589260807916945 | 0.0000356436939729832 |
| *MYDGF* | 0.74298123555392 | 0.000036103562644007 |
| *H2AFY* | 0.826493221147489 | 0.0000408020972100244 |
| *TPT1* | 0.986000692991493 | 0.0000432708182995778 |

| | | |
|---|---|---|
| *RPS21* | 1.02708489893616 | 0.0000436566847800543 |
| *TMEM219* | 0.748419747042914 | 0.0000445084582882374 |
| *SNU13* | 0.826493221147489 | 0.0000453501191094317 |
| *UCP2* | 0.957096554204169 | 0.0000460607781907646 |
| *RABAC1* | 0.685891409571937 | 0.0000473618178931326 |
| *LAPTM5* | 0.867022499497292 | 0.0000484559119507806 |
| *ODC1* | 0.775515199702312 | 0.0000500990780811401 |
| *RBX1* | 0.71230080200392 | 0.0000501359429986622 |
| *STRBP* | 0.583694754557955 | 0.0000540190435925438 |
| *EEF1D* | 0.883628743203278 | 0.0000576346682597132 |
| *PYCARD* | 0.701298720870185 | 0.000061115634970048 |
| *SLC39A6* | 0.49435995194025 | 0.0000614534110250038 |
| *TMED3* | 0.49435995194025 | 0.0000614534110250038 |
| *PLVAP* | 0.580774703757723 | 0.0000614534110250056 |
| *ANAPC5* | 0.659478545175558 | 0.0000650107016952994 |
| *TXNL4A* | 0.72119827830789 | 0.0000679420096286192 |
| *TPP1* | 0.690405124762286 | 0.0000733625680070911 |
| *ATP5ME* | 0.91114420664933 | 0.0000740835004409063 |
| *ATP5F1C* | 0.73448977466028 | 0.0000783164745152264 |
| *CCDC186* | 0.97618041184695 | 0.0000795557683648599 |
| *ARF1* | 0.826380514999946 | 0.0000797998506190824 |
| *EIF3L* | 0.80966878789585 | 0.0000803596632625006 |
| *RPS6* | 1.07869736814518 | 0.0000840800723628394 |
| *NDUFB1* | 0.873097013906976 | 0.000085550201746689 |
| *SERTAD2* | 0.602641667955982 | 0.0000874092304076189 |
| *GPR183* | 1.52429082852616 | 0.0000890621024914746 |
| *ASPH* | 0.583694754557955 | 0.0000944422227749817 |
| *ATP5MF* | 0.874608192967185 | 0.000101061878358654 |
| *RPS16* | 1.00115047457523 | 0.000102543405872028 |
| *STX7* | 0.470868980468318 | 0.000105238798996262 |
| *RPL17* | 1.0795499481348 | 0.000106434637518203 |
| *MT-CYB* | 0.799437049462887 | 0.00010922609950962 |
| *RPL24* | 0.960680092547181 | 0.000110376283678946 |
| *MT-ATP6* | 0.894243319460283 | 0.000111494329511171 |
| *SMARCB1* | 0.607512116413942 | 0.000111632299682828 |
| *CDH1* | 0.485426827170242 | 0.000112308463334255 |
| *WAKMAR2* | 0.503238103286702 | 0.000112308463334255 |
| *COX4I1* | 0.899081978181185 | 0.000112537148711295 |

| | | |
|---|---|---|
| *RPL13* | 1.08767410648014 | 0.000113322316388053 |
| *DCK* | 0.732840391082683 | 0.000118546482196202 |
| *COX8A* | 0.865948666145974 | 0.000121540963026257 |
| *ARPC2* | 0.825671874674471 | 0.000121648599903976 |
| *MGAT1* | 0.726039541377389 | 0.000128164771978273 |
| *RAC1* | 0.794261740497613 | 0.00013046020030829 |
| *ITGAE* | 0.591915261362923 | 0.000140581846403923 |
| *TMEM109* | 0.775137085418693 | 0.000151538423446561 |
| *CDK2AP2* | 0.741940705367531 | 0.000161709629641924 |
| *CDYL* | 0.78094908575294 | 0.000165838308886355 |
| *NME4* | 0.583694754557955 | 0.000167494068938887 |
| *TRAM1* | 0.868039666637161 | 0.000173167488156786 |
| *OS9* | 0.706610742084843 | 0.000181780389606322 |
| *NDUFS6* | 0.708152391911247 | 0.000185269880212576 |
| *AHCY* | 0.49003564734371 | 0.000186685411527538 |
| *GPAA1* | 0.49003564734371 | 0.000189648708453548 |
| *BUD23* | 0.628573731941771 | 0.000191734874304281 |
| *CIITA* | 0.449130697947252 | 0.000204186175211687 |
| *NOTCH4* | 0.49435995194025 | 0.000204186175211687 |
| *PDXP* | 0.485426827170242 | 0.000204186175211687 |
| *MTRNR2L1* | 0.877317480225171 | 0.000204261236756932 |
| *ZMYM2* | 0.734873215204966 | 0.000206017083557851 |
| *NUCB2* | 0.903116949657227 | 0.00020835828394107 |
| *CD37* | 0.980612368462986 | 0.000211338083944093 |
| *RPL32* | 1.01988781596927 | 0.000214683431890415 |
| *LYPLA1* | 0.620796381350052 | 0.000214726804640474 |
| *MXD4* | 0.755661687876111 | 0.000219078891589448 |
| *HDGF* | 0.636309380721376 | 0.000220334202878227 |
| *GTF2I* | 0.72119827830789 | 0.000237976226613757 |
| *EID1* | 0.821961255242806 | 0.000245591401204609 |
| *PHB2* | 0.729418785112858 | 0.000252714565461567 |
| *PLEKHO1* | 0.67355373038728 | 0.000265977814679789 |
| *GDI2* | 0.681669914121252 | 0.000267319408310033 |
| *DBI* | 0.777781806674257 | 0.000267762026399549 |
| *DPP7* | 0.68045193596458 | 0.000272453701159543 |
| *TMEM179B* | 0.557020470916737 | 0.000279819921076123 |
| *KTN1* | 0.83819114893535 | 0.000283411256262045 |
| *TCF3* | 0.405146749757197 | 0.000286792501225955 |

| | | |
|---|---|---|
| *ZFAT* | 0.558748397427724 | 0.000289178395601989 |
| *FAM118A* | 0.567111870362351 | 0.00031300421745248 |
| *APPL1* | 0.690501481084973 | 0.000313443753347842 |
| *SPCS2* | 0.69744967553528 | 0.000316247332541142 |
| *ANKRD11* | 1.02236781302845 | 0.000317798103737131 |
| *TMEM14B* | 0.689526057989345 | 0.000318416045428607 |
| *NDUFA13* | 0.667447892530281 | 0.000329842350047971 |
| *RPL18* | 1.00386149089062 | 0.000356863877351347 |
| *SEPTIN2* | 0.718024030512746 | 0.000366327266016835 |
| *UBE2E2* | 0.421296489750526 | 0.000369351995572057 |
| *GNA12* | 0.430634354329862 | 0.000369351995572062 |
| *MARCH1* | 0.476438043942987 | 0.000369351995572062 |
| *NRP1* | 0.46739290433998 | 0.000369351995572062 |
| *PEA15* | 0.476438043942987 | 0.000369351995572062 |
| *SEMA7A* | 0.46739290433998 | 0.000369351995572062 |
| *ASIP* | 0.430634354329862 | 0.000369351995572068 |
| *CYP46A1* | 0.597453444904353 | 0.000369351995572068 |
| *EGLN3* | 0.439912167917873 | 0.000369351995572068 |
| *PTPRS* | 0.411897791748276 | 0.000369351995572068 |
| *KCTD5* | 0.605114759117782 | 0.000375618412638527 |
| *TSPAN3* | 0.591915261362923 | 0.000381164179873375 |
| *RPN1* | 0.490768537906023 | 0.000422502742720861 |
| *ZSCAN16-AS1* | 0.584053143589953 | 0.00042591701540226 |
| *SRP72* | 0.646904495960576 | 0.000440226437844035 |
| *SAMHD1* | 0.876218283950368 | 0.00044873130160754 |
| *QKI* | 0.694153455293661 | 0.000454309929672047 |
| *CSNK2B* | 0.597209561032436 | 0.000468939925187021 |
| *CDC40* | 0.685891409571937 | 0.000470687475206416 |
| *PDIA6* | 0.694153455293661 | 0.000480517381415285 |
| *PTGDS* | 2.92209745746294 | 0.000482430634082981 |
| *BST2* | 0.73818899474059 | 0.000482875104686934 |
| *RPL28* | 0.87758486879806 | 0.000484151923120192 |
| *RPS14* | 0.981679692781977 | 0.000496512469334119 |
| *NDUFC2* | 0.807462879931033 | 0.000498513475355073 |
| *NDUFA11* | 0.683837750803962 | 0.00051726854053826 |
| *EIF3H* | 0.777781806674257 | 0.000540197734568906 |
| *RPL26* | 1.03848016340277 | 0.000540430996188681 |
| *IRF2BPL* | 0.575427138574354 | 0.000557074552374462 |

| | | |
|---|---|---|
| *RPL9* | 1.02529718878874 | 0.000564812703749449 |
| *TP53I13* | 0.622942156983547 | 0.000611223442461063 |
| *S100A6* | 0.985300161098183 | 0.000621019237993317 |
| *ETV6* | 0.570220190263604 | 0.000624234970320418 |
| *KIF5B* | 0.726216353074514 | 0.000624991880705733 |
| *PRKCB* | 0.855962440169785 | 0.000630298127081932 |
| *SELL* | 0.859451059431658 | 0.000642755245650859 |
| *SCRN1* | 0.439912167917873 | 0.000664824727399452 |
| *PHEX* | 0.476438043942987 | 0.000664824727399462 |
| *PLXNB2* | 0.421296489750526 | 0.000664824727399462 |
| *MYCBP* | 0.46739290433998 | 0.000664824727399472 |
| *NOP10* | 0.55988143372321 | 0.000668071076643031 |
| *YWHAE* | 0.690405124762286 | 0.000727356000021976 |
| *RPS18* | 0.843723798467557 | 0.000742215185858445 |
| *PEBP1* | 0.734873215204966 | 0.000748398619505658 |
| *ADI1* | 0.59448920502675 | 0.000773254307284593 |
| *RPS27A* | 1.00544413869013 | 0.00079392656928536 |
| *NDUFA1* | 0.771148005031463 | 0.000827571616867779 |
| *EIF3I* | 0.553847220529403 | 0.000851613077047422 |
| *CREB3L2* | 0.653315806027067 | 0.000891102654353539 |
| *YWHAZ* | 0.676448771574591 | 0.000907756455862318 |
| *C4orf48* | 0.578394121709304 | 0.000972134170554677 |
| *MTDH* | 0.905575740266685 | 0.00100728834497931 |
| *LSP1* | 0.904005163925033 | 0.00103707106041462 |
| *TAX1BP3* | 0.541874578863328 | 0.00104436580226771 |
| *SNRPN* | 0.761840262805235 | 0.00106073292665325 |
| *N4BP2L1* | 0.694412335284945 | 0.00106404120964815 |
| *ELOB* | 0.477667089913291 | 0.00107356111072379 |
| *CYC1* | 0.630909392978787 | 0.00113800925993139 |
| *CDKN2D* | 0.919545715614469 | 0.00116545639455445 |
| *BID* | 0.630595729526983 | 0.00116664513140834 |
| *SELENOF* | 0.631672970600658 | 0.0011907948486802 |
| *ATF5* | 0.392914688380704 | 0.0011909084402811 |
| *FAM160A1* | 0.383328639551506 | 0.00119090844028112 |
| *LTK* | 0.373678469517786 | 0.00119090844028112 |
| *PROC* | 0.458290697232728 | 0.00119090844028112 |
| *TLR9* | 0.421296489750526 | 0.00119090844028112 |
| *UNC119* | 0.599734765822222 | 0.00120517077033293 |

| | | |
|---|---|---|
| *PGK1* | 0.713544705764454 | 0.00120713464285296 |
| *TTC3* | 0.773111375140256 | 0.00121972668188 |
| *ATP5PB* | 0.645264685955857 | 0.0012294439570825 |
| *CYTH4* | 0.615247765193547 | 0.00123004246472403 |
| *SLC20A1* | 0.625023573174862 | 0.00131566379270491 |
| *M6PR* | 0.804364879451792 | 0.0013429385217105 |
| *DDT* | 0.639356256280241 | 0.00140052239090898 |
| *LSM10* | 0.568199192389116 | 0.00142197864839116 |
| *FIS1* | 0.563294583423156 | 0.00143520639024328 |
| *AP3S1* | 0.58696604539119 | 0.00144216479051111 |
| *ST13* | 0.755816475177465 | 0.00145937980578328 |
| *PRDX1* | 1.83770227732054 | 0.00146264336252701 |
| *ATP5F1D* | 0.777270303374841 | 0.0014700476533903 |
| *MAP2K3* | 0.625635879206924 | 0.00148851704232147 |
| *PLAAT3* | 0.60176304731927 | 0.00160739861849617 |
| *RPSA* | 0.897331142168629 | 0.00162289914933716 |
| *SH3BGRL3* | 0.804364879451792 | 0.00164644899963377 |
| *GLRX* | 0.52265259892581 | 0.00168048938260116 |
| *RAP1GDS1* | 0.516187935199678 | 0.0017288869258467 |
| *C1QBP* | 0.906520015444427 | 0.00177916861252784 |
| *LMAN1* | 0.678275546166104 | 0.00178492365521229 |
| *MPG* | 0.520179507764009 | 0.0018788606680617 |
| *HDLBP* | 0.49003564734371 | 0.00192715229138113 |
| *ADD1* | 0.516507858704901 | 0.00197906085387271 |
| *PPP1R9B* | 0.463400520840243 | 0.00201873474328299 |
| *SERP1* | 0.699874509498843 | 0.00202932532652916 |
| *UQCR10* | 0.663502343611943 | 0.00203582930501645 |
| *SPCS3* | 0.782160247184831 | 0.00204846054099119 |
| *TRIM8* | 0.586753556247144 | 0.00205889725222206 |
| *COX7A2* | 0.817582814732232 | 0.00210180940819694 |
| *CHML* | 0.46739290433998 | 0.00212327228480498 |
| *SIRPB1* | 0.421296489750526 | 0.00212327228480498 |
| *NSMCE4A* | 0.363963314684575 | 0.00212327228480501 |
| *TMEM170B* | 0.411897791748276 | 0.00212327228480501 |
| *MCOLN2* | 0.411897791748276 | 0.00212327228480507 |
| *MIIP* | 0.494492864100359 | 0.00213840132597343 |
| *SFT2D1* | 0.557020470916737 | 0.00214101746932159 |
| *HNRNPC* | 0.779066676915832 | 0.00217924476812881 |

| | | |
|---|---|---|
| *LAP3* | 0.584053143589953 | 0.00219802980947135 |
| *PSMA6* | 0.664978723552646 | 0.00222150439525442 |
| *NUDT22* | 0.527784923943208 | 0.00224540304984896 |
| *MRPS21* | 0.627539171093644 | 0.00225546445220641 |
| *CHMP4B* | 0.591915261362923 | 0.00226828398804246 |
| *NRDC* | 0.643577381185416 | 0.00229653856616388 |
| *SEPTIN6* | 0.761840262805235 | 0.00230481859470479 |
| *NIN* | 0.60176304731927 | 0.00230910430045294 |
| *GTF2A2* | 0.654925058888724 | 0.00245825908322267 |
| *CCNI* | 0.737057161875347 | 0.00247389267821302 |
| *PTPN2* | 0.653924923638348 | 0.00251062986796516 |
| *HNRNPD* | 0.666024475391435 | 0.00254846329572093 |
| *POLE4* | 0.402297876118095 | 0.00260973646533821 |
| *SF3B5* | 0.671074798882053 | 0.00267254765060798 |
| *PARP10* | 0.498805856971442 | 0.00272897200076126 |
| *NEDD8* | 0.6266806795236 | 0.00285782613193751 |
| *RRP7A* | 0.550336157611524 | 0.00303422885910885 |
| *SUSD1* | 0.610634458709512 | 0.00311041179064451 |
| *KDELR2* | 0.617450353470061 | 0.00317264154733112 |
| *PCBP2* | 0.692642752446209 | 0.0031870958089033 |
| *MIS18BP1* | 0.667864114596222 | 0.00319995687553862 |
| *SINHCAF* | 0.578976205655425 | 0.00321656841578403 |
| *PGD* | 0.528641086512205 | 0.00330853741583278 |
| *EIF3D* | 0.627129953946981 | 0.00344217598357094 |
| *RPL36AL* | 0.787666250173141 | 0.00356552381096429 |
| *TYMP* | 0.448474181421297 | 0.00364719181228268 |
| *LSM4* | 0.568199192389116 | 0.00365517302573252 |
| *RPS19BP1* | 0.626185163447669 | 0.00365847607764818 |
| *AP2S1* | 0.631190410096363 | 0.00369928320728379 |
| *ALCAM* | 0.383328639551506 | 0.00376825789656184 |
| *FLT3* | 0.392914688380704 | 0.00376825789656184 |
| *HEXB* | 0.392914688380704 | 0.00376825789656184 |
| *HHIP-AS1* | 0.383328639551506 | 0.00376825789656184 |
| *NIPSNAP2* | 0.344334507935642 | 0.00376825789656184 |
| *PLEKHD1* | 0.40243746249921 | 0.00376825789656184 |
| *PNOC* | 0.512061953673708 | 0.00376825789656184 |
| *NECTIN1* | 0.411897791748276 | 0.0037682578965619 |
| *TMED10* | 0.698414882663602 | 0.0038037054966776 |

| | | |
|---|---|---|
| *SND1* | 0.532358416682468 | 0.0039009659453201 |
| *SLC66A2* | 0.552169082570167 | 0.00394939834340399 |
| *EIF3F* | 0.704389990621336 | 0.00408765488047344 |
| *H1FX* | 0.838075848632321 | 0.00410756469729321 |
| *PNISR* | 0.898999509011265 | 0.00426570654736681 |
| *TMEM14C* | 0.532358416682468 | 0.00428377592617449 |
| *SNRPD2* | 0.746733370415027 | 0.00430813707153562 |
| *SFPQ* | 0.827131723056955 | 0.00435704010148048 |
| *PTBP3* | 0.779762170802498 | 0.00441507132806053 |
| *NDUFB11* | 0.681221802596951 | 0.0044672818925556 |
| *ZDHHC4* | 0.532358416682468 | 0.00456751028540493 |
| *UQCR11* | 0.765713271952442 | 0.00466839408185965 |
| *MAP4K4* | 0.603578178888503 | 0.0050138214578066 |
| *CBX6* | 0.656487262659007 | 0.00518623112780234 |
| *MARCH9* | 0.507267435719638 | 0.00521566287923685 |
| *SERBP1* | 0.790819329777001 | 0.00538512336364489 |
| *RPL21* | 0.891460187479337 | 0.00568297168917082 |
| *MZT2B* | 0.707392478782859 | 0.00568787400561387 |
| *CHAF1A* | 0.454411737612988 | 0.00585472894246556 |
| *PLIN3* | 0.349193310951386 | 0.0058575117404417 |
| *IER3IP1* | 0.485828002973232 | 0.0059139345348157 |
| *CSF2RA* | 0.498805856971442 | 0.00592231231735616 |
| *RPL34* | 1.01754683850784 | 0.00604105526824268 |
| *DNAJC4* | 0.632557245860269 | 0.00622585886768697 |
| *PET100* | 0.656045598782639 | 0.00623276488160448 |
| *LMO4* | 0.586522002592705 | 0.00625557537080864 |
| *EIF1* | 0.611369273772738 | 0.00632326255002776 |
| *CIRBP* | 0.776339832500351 | 0.00660052400206554 |
| *DGKZ* | 0.666262602823005 | 0.00662064501551529 |
| *HSD17B4* | 0.344334507935642 | 0.00665786109360267 |
| *FMNL3* | 0.354182293891989 | 0.00665786109360272 |
| *NEK8* | 0.421296489750526 | 0.00665786109360272 |
| *COL26A1* | 0.363963314684575 | 0.00665786109360277 |
| *QDPR* | 0.334419039070559 | 0.00665786109360277 |
| *CHAMP1* | 0.383328639551506 | 0.00665786109360287 |
| *TMEM8B* | 0.383328639551506 | 0.00665786109360287 |
| *DDX17* | 0.775015651552969 | 0.00671966398457958 |
| *PDIA4* | 0.634648246351456 | 0.00675485302280418 |

| | | |
|---|---|---|
| *CHMP3* | 0.446484821841359 | 0.0068381272748098 |
| *TM9SF3* | 0.648498789729291 | 0.00708063174125514 |
| *PGLS* | 0.632760313640306 | 0.00786282323761098 |
| *APBB1IP* | 0.625635879206924 | 0.00793518523069079 |
| *NENF* | 0.523328176105511 | 0.00846172493998114 |
| *GRASP* | 0.679378102613262 | 0.00853838804911176 |
| *RALBP1* | 0.494736192662395 | 0.00864835535721476 |
| *TMX1* | 0.465447260025011 | 0.00867212492734939 |
| *CD99* | 0.548846539471037 | 0.00884742097097719 |
| *ULK1* | 0.591915261362923 | 0.00903705908950931 |
| *TALDO1* | 0.6463630453853 | 0.00971184534358157 |
| *BTG2* | 0.899730619924129 | 0.00981601257233356 |
| *KDELR1* | 0.560206401635585 | 0.0104376002204795 |
| *COX14* | 0.529666820676202 | 0.0106360036422169 |
| *ACADVL* | 0.617450353470061 | 0.0107555480539786 |
| *UBE2V1* | 0.556811460407007 | 0.0108992931235544 |
| *RPL39* | 1.05150282174397 | 0.0110347143581468 |
| *ANP32B* | 0.74304266735111 | 0.0111341604246646 |
| *APEX1* | 0.625635879206924 | 0.011308896875878 |
| *SEC11C* | 0.719135811850997 | 0.0113654165466762 |
| *MLEC* | 0.510615159261075 | 0.0115769331656075 |
| *SRSF7* | 0.884175805729288 | 0.0116788831196111 |
| *FBH1* | 0.314381285834014 | 0.0117122380442032 |
| *HDAC9* | 0.344334507935642 | 0.0117122380442034 |
| *HHEX* | 0.421296489750526 | 0.0117122380442035 |
| *TM9SF1* | 0.373678469517786 | 0.0117122380442035 |
| *ABCD4* | 0.324434950497937 | 0.0117122380442036 |
| *MGST2* | 0.334419039070559 | 0.0117122380442036 |
| *SLC15A3* | 0.373678469517786 | 0.0117122380442036 |
| *TBC1D9* | 0.334419039070559 | 0.0117122380442036 |
| *ADA* | 0.354182293891989 | 0.0117122380442037 |
| *ERN1* | 0.609837169360185 | 0.0117954285375936 |
| *XRCC5* | 0.596591748572396 | 0.0118927595506774 |
| *SGSM3* | 0.441705449740924 | 0.0119291934949819 |
| *GLRX5* | 0.621662604756975 | 0.0123041455426651 |
| *LAMTOR5* | 0.615450775823712 | 0.0123414615929487 |
| *PTRHD1* | 0.621662604756975 | 0.0123583676036943 |
| *GRSF1* | 0.531410522403369 | 0.0125340945049304 |

| | | |
|---|---|---|
| *CCDC85B* | 0.546715841932827 | 0.0125383976041087 |
| *HMGN3* | 0.570955642727277 | 0.0125686758370775 |
| *C11orf58* | 0.761840262805235 | 0.0125721346149565 |
| *CPNE3* | 0.502753041488105 | 0.0128805995805252 |
| *MARCKSL1* | 0.632760313640306 | 0.0134367573014009 |
| *SLC2A1* | 0.498805856971442 | 0.0137604217183825 |
| *METAP2* | 0.606690614043321 | 0.013983026370556 |
| *NAA38* | 0.534925282778093 | 0.0140554776676692 |
| *C17orf49* | 0.61053093953027 | 0.0141046425924901 |
| *RBBP4* | 0.544386891280337 | 0.0142943893611334 |
| *APRT* | 0.719612027415938 | 0.0143922350560857 |
| *HNRNPM* | 0.66039499924569 | 0.0144460426030778 |
| *MEF2D* | 0.606865602828895 | 0.0147107927637054 |
| *PRELID1* | 0.690581579635414 | 0.0148096745575919 |
| *RNF11* | 0.486205820191808 | 0.0148455540227751 |
| *SUPT4H1* | 0.473859499841211 | 0.0148936577845817 |
| *SNHG8* | 0.684048102199084 | 0.0152171554449714 |
| *OIP5-AS1* | 0.494736192662394 | 0.0152321128377017 |
| *RPL23* | 0.800461556832543 | 0.0152331341289965 |
| *TOMM6* | 0.722156561332754 | 0.0152857127154788 |
| *HNRNPAB* | 0.675979526151398 | 0.015617798090272 |
| *NOP56* | 0.67355373038728 | 0.0160356502437669 |
| *SET* | 0.812824191524041 | 0.0163143312509548 |
| *RAD23A* | 0.526212014312093 | 0.016844721009523 |
| *PDIA3* | 0.654199539617266 | 0.0170369557193703 |
| *MRPL36* | 0.545416799263032 | 0.0170455252233575 |
| *PMEPA1* | 0.472333645610251 | 0.0173322506628959 |
| *RPS15A* | 0.90719704971721 | 0.0176203315740083 |
| *TBC1D1* | 0.454411737612988 | 0.0177498932228153 |
| *GRINA* | 0.427104391617253 | 0.0180899844298747 |
| *ANXA5* | 0.477110785872123 | 0.0184451291935261 |
| *MAGED1* | 0.417885861587874 | 0.0184843216191932 |
| *CLTA* | 0.621662604756975 | 0.0198821071548484 |
| *ING3* | 0.432716666513669 | 0.0201178143372384 |
| *KCTD12* | 0.344334507935642 | 0.0205167465314083 |
| *NREP* | 0.314381285834014 | 0.0205167465314085 |
| *ZDHHC24* | 0.344334507935642 | 0.0205167465314085 |
| *LCNL1* | 0.485426827170242 | 0.0205167465314087 |

| | | |
|---|---|---|
| *PPP1R14B-AS1* | 0.383328639551506 | 0.0205167465314087 |
| *VASH2* | 0.392914688380704 | 0.0205167465314087 |
| *MFSD2A* | 0.383328639551506 | 0.020516746531409 |
| *NUDT5* | 0.334419039070559 | 0.020516746531409 |
| *MRPS10* | 0.314381285834014 | 0.0205167465314093 |
| *MOB1A* | 0.756426766618871 | 0.0206291108708661 |
| *LSM3* | 0.544853224490878 | 0.0209695003417798 |
| *GAPDH* | 1.25779775746765 | 0.0210167396380959 |
| *KIAA2013* | 0.441705449740924 | 0.0212462921252708 |
| *RCC2* | 0.318233611329621 | 0.0216100829624079 |
| *CD63* | 0.397324539105558 | 0.0217191540616998 |
| *HADHA* | 0.619587679847478 | 0.0218495856988548 |
| *UBE2E3* | 0.629004580098143 | 0.0221973487027357 |
| *UBE2D2* | 0.655711911943892 | 0.0233492544963442 |
| *ZNF22* | 0.486562261216694 | 0.0246271759445494 |
| *PPP1R14A* | 0.663865103241938 | 0.0250276379626006 |
| *BORCS7* | 0.502981532942635 | 0.0255117102486306 |
| *PSMB7* | 0.482222972775837 | 0.0257175729832928 |
| *SNHG9* | 0.723943655046707 | 0.0262410817137734 |
| *RNF5* | 0.464454352180665 | 0.0272591955290224 |
| *LILRB1* | 0.368176414318958 | 0.0280143479319923 |
| *RFLNB* | 0.557020470916737 | 0.0285146066813738 |
| *MAN1A1* | 0.469232741588094 | 0.0289426679738446 |
| *ATAD2B* | 0.411343015721102 | 0.0294608513596793 |
| *SMC6* | 0.454411737612988 | 0.0300988501088835 |
| *P2RY14* | 0.472333645610251 | 0.0305798567587001 |
| *ZRANB2* | 0.602738230657459 | 0.030713216218879 |
| *UBE2L3* | 0.570022655514081 | 0.0309510213631836 |
| *GIT2* | 0.445366598009981 | 0.0314274169318949 |
| *CAPN15* | 0.584053143589953 | 0.0317870986256772 |
| *CYB5R3* | 0.441705449740924 | 0.0324592663904339 |
| *PHPT1* | 0.53051471669878 | 0.0327843902788606 |
| *GRB2* | 0.701968806827837 | 0.0331019133978335 |
| *PRDX5* | 0.596254197087261 | 0.0334076006781254 |
| *TRMT112* | 0.591915261362923 | 0.0336275061564474 |
| *NR3C1* | 0.669917773364196 | 0.034067464934923 |
| *GABARAPL2* | 0.671642453833657 | 0.0343636509109744 |
| *RANBP1* | 0.674783528536012 | 0.0344030050200205 |

| | | |
|---|---|---|
| *CSNK1E* | 0.443392736652835 | 0.0345901012544281 |
| *CCAR1* | 0.55069259863641 | 0.0346495375812308 |
| *ICAM1* | 0.51520029522211 | 0.0353101549467892 |
| *ISCU* | 0.59428616582567 | 0.0356313848196343 |
| *LHFPL2* | 0.334419039070559 | 0.0357926407590754 |
| *AL138756.1* | 0.304257068560307 | 0.0357926407590762 |
| *MRPL14* | 0.334419039070559 | 0.0357926407590762 |
| *AC011893.1* | 0.314381285834014 | 0.0357926407590764 |
| *ARHGAP24* | 0.363963314684575 | 0.0357926407590764 |
| *MILR1* | 0.314381285834014 | 0.0357926407590767 |
| *VTI1B* | 0.432716666513669 | 0.0367688987309711 |
| *DOCK2* | 0.520832163301441 | 0.0374509746324188 |
| *SEC62* | 0.661263308053658 | 0.0390415163162465 |
| *SF1* | 0.765398082218425 | 0.0404013143793718 |
| *SRSF6* | 0.61053093953027 | 0.0413055746403114 |
| *UBL5* | 0.701036983743036 | 0.0414487744819469 |
| *RTF2* | 0.466384379279064 | 0.0415392375301151 |
| *MED13L* | 0.624336739055301 | 0.0422135269610863 |
| *ANXA11* | 0.614799152036262 | 0.0425093603071409 |
| *XRCC6* | 0.618387472724114 | 0.0425271738146234 |
| *RPL30* | 0.780360350776033 | 0.0429422244146017 |
| *PSMA2* | 0.6202256573863 | 0.0433339935246298 |
| *PHYKPL* | 0.456985681276815 | 0.0439774985328428 |
| *MAP3K8* | 0.596254197087261 | 0.0466410242237592 |
| *DYNLRB1* | 0.517914679919146 | 0.0469633807259523 |
| *ENO1* | 1.02743037354279 | 0.047430621322176 |
| *HNRNPK* | 0.630050390249694 | 0.0476947287828833 |
| *ATP5MC2* | 0.751033507848653 | 0.04888039829417 |
