## Supplementary Table 15 for "Heterogeneity of circulating epithelial cells in breast cancer at single-cell resolution: identifying tumor and hybrid cells"

Supplementary Table 15. Specific upregulated genes in diploid cells of CD45$^{-}$ CEC clusters

| Cluster 1 | | Genes | Cluster 2 | |
|---|---|---|---|---|
| LogFC | Adjusted P-value | | LogFC | Adjusted P-value |
| *ANKRD12* | 1.48987273220143 | *FTH1* | 1.77551519970231 | 5.97376094878555E-18 |
| *SLC38A2* | 1.51457317282976 | *HLA-DMA* | 1.45637059065105 | 6.03631617153421E-16 |
| *RABGAP1L* | 1.16655140201421 | *SRP14* | 1.60121300594501 | 1.01739510885295E-15 |
| *RELT* | 0.808048692227689 | *OAZ1* | 1.32690320608869 | 6.32816365063845E-15 |
| *DDX21* | 1.09667962196581 | *SERF2* | 1.35156975643384 | 8.99382470262576E-15 |
| *RILPL2* | 0.852442811586142 | *NME2* | 1.52577290457171 | 1.6178686993265E-14 |
| *ATP5PF* | 1.01471424048502 | *BRI3* | 1.34482239939815 | 4.39319818561038E-13 |
| *HNRNPH1* | 0.950290134984287 | *TUBA1B* | 1.62000756839228 | 6.40247850929198E-13 |
| *ELL* | 0.53051471669878 | *PTCRA* | 1.29859367048377 | 6.50557113124882E-13 |
| *KLF4* | 0.736965594166206 | *EIF4A3* | 1.05882200056237 | 7.8236524882568E-13 |
| *MYCL* | 0.53051471669878 | *GZMB* | 2.23683971852514 | 1.46972842842841E-12 |
| | | *EEF1A1* | 1.49989099808382 | 1.74477012097583E-12 |
| | | *UNC93B1* | 1.18225093418293 | 2.60175890737595E-12 |
| | | *RPL15* | 1.36807138028004 | 3.01900150707116E-12 |
| | | *MYL6* | 1.18077167818857 | 4.66358493755851E-12 |
| | | *PLP2* | 1.47139039939104 | 6.52535964365244E-12 |
| | | *VIM* | 1.74122998312995 | 8.61662749362373E-12 |
| | | *EEF2* | 1.30498258783176 | 1.70407691191264E-11 |
| | | *OSTC* | 1.10394623963023 | 2.71955799775869E-11 |
| | | *ARPC1B* | 1.17362478428899 | 4.39754545010632E-11 |
| | | *DAB2* | 1.0124909441832 | 4.60099273277718E-11 |
| | | *WDFY4* | 0.746966986323705 | 4.60099273277718E-11 |
| | | *ACTB* | 1.11636041000392 | 6.64961101582148E-11 |
| | | *CAT* | 1.09072111833437 | 7.39950703912558E-11 |
| | | *RASD1* | 1.35418229389199 | 9.69442010916461E-11 |
| | | *HINT1* | 1.33708760869318 | 1.01256826214091E-10 |
| | | *HIGD1A* | 0.883830787183846 | 1.04346949139381E-10 |
| | | *DPYSL2* | 0.993431620988148 | 1.04844897296513E-10 |
| | | *SEC61G* | 1.29906520109357 | 1.83415550564713E-10 |
| | | *IGHM* | 1.40860804799986 | 1.88096468021389E-10 |
| | | *HMGA1* | 1.16063712107687 | 2.80211977439362E-10 |
| | | *MS4A6A* | 1.1381081919195 | 2.91001420307214E-10 |
| | | *SNHG7* | 1.14849596071426 | 3.54724346520094E-10 |
| | | *HLA-DMB* | 0.853582831432811 | 3.95525360547372E-10 |
| | | *MYBL2* | 0.948925815405731 | 4.19601764030209E-10 |
| | | *RPS6KA4* | 0.924569762843344 | 4.75021489516549E-10 |
| | | *C12orf45* | 0.999531820524566 | 4.75896830316982E-10 |
| | | *CFL1* | 1.18241594562551 | 6.14705349565107E-10 |
| | | *TMED2* | 0.92742632852321 | 6.61326317754961E-10 |
| | | *CAPG* | 1.03560107523209 | 1.0868853537085E-09 |
| | | *PHACTR1* | 0.981153291209711 | 1.16079411765672E-09 |
| | | *GNG5* | 1.0911478880582 | 1.20963841336371E-09 |
| | | *ARPC3* | 1.27482281249604 | 1.30308955571849E-09 |
| | | *RUNX2* | 0.864201980434913 | 1.3861837375421E-09 |
| | | *ATP5F1B* | 1.07499814806487 | 1.45274465860476E-09 |
| | | *P4HB* | 1.12173620789162 | 1.57176019216452E-09 |
| | | *RPL22* | 1.22465012036563 | 1.73616056397756E-09 |
| | | *PECAM1* | 0.81272143180742 | 1.75814291626932E-09 |
| | | *MIF4GD* | 0.754422791242752 | 1.75814291626935E-09 |
| | | *ST14* | 0.776561768512001 | 1.75814291626935E-09 |
| | | *LAPTM4A* | 0.937690098204653 | 2.18563695479673E-09 |
| | | *YBX1* | 1.24776124824059 | 2.27120181851476E-09 |
| | | *RNASEK* | 1.06708961654597 | 2.28770351344425E-09 |
| | | *JAML* | 0.905204437976413 | 2.47046569791084E-09 |
| | | *RPS24* | 1.47460158876292 | 2.78633424227132E-09 |
| | | *GPX4* | 0.883987372091026 | 2.94158543396009E-09 |
| | | *RPL10A* | 1.26987388509347 | 3.20468285810699E-09 |

| | | |
|---|---|---|
| *RPL4* | 1.26351644162055 | 3.41537069893626E-09 |
| *FCHSD2* | 0.867193262951464 | 3.65993006167142E-09 |
| *ATP5F1A* | 1.03747470541866 | 3.8383997016163E-09 |
| *RGS2* | 1.61524776519355 | 6.18093741096257E-09 |
| *RPS28* | 1.48346177925581 | 6.69717862052221E-09 |
| *GRAMD1B* | 0.873950629126773 | 6.84447511995499E-09 |
| *CALM2* | 1.11594876344175 | 7.35687362416934E-09 |
| *SULF2* | 0.818984169908844 | 7.54104815754724E-09 |
| *CERS6* | 0.987305518191307 | 7.81484255781189E-09 |
| *TYROBP* | 1.36504188181143 | 1.00724924505836E-08 |
| *CHCHD2* | 1.25289562892886 | 1.11148715587652E-08 |
| *SLC25A6* | 1.17376515021674 | 1.1697339626593E-08 |
| *EEF1G* | 1.14571683059762 | 1.18212977880469E-08 |
| *GSN* | 0.80723216561925 | 1.19020265838771E-08 |
| *COMT* | 0.83161054110941 | 1.32602785472725E-08 |
| *RPN2* | 1.00041261119712 | 1.33812567710253E-08 |
| *SSR3* | 0.917959464722517 | 1.38347404450247E-08 |
| *LYN* | 1.09000031140411 | 1.39475247945991E-08 |
| *BLNK* | 0.724365557386573 | 1.42369773887861E-08 |
| *LINC02812* | 0.739472449776781 | 1.42369773887861E-08 |
| *ALDH2* | 0.739472449776781 | 1.42369773887863E-08 |
| *GNG7* | 0.646363045385299 | 1.42369773887863E-08 |
| *RPL29* | 1.35941223983441 | 1.81654477311093E-08 |
| *SYK* | 0.918143493683911 | 2.07722341150563E-08 |
| *PLEK* | 1.4602460104404 | 2.16001191700199E-08 |
| *RPS5* | 1.36121244287967 | 2.16188095791049E-08 |
| *RPS3A* | 1.47495265131513 | 2.19533623871502E-08 |
| *RPS9* | 1.25404562260654 | 2.23338009955547E-08 |
| *SCARB2* | 0.880484759303855 | 2.26948658951239E-08 |
| *RPS2* | 1.33792993923751 | 2.29934053642872E-08 |
| *OPN3* | 0.693668760163656 | 2.81750096063245E-08 |
| *ZNF706* | 0.921948405887341 | 3.1755868380086E-08 |
| *IRF2BP2* | 1.1213004141892 | 3.48943912727326E-08 |
| *AP1S2* | 0.866334430676341 | 3.50871308607093E-08 |
| *TM9SF2* | 0.860020656744139 | 3.5957310315458E-08 |
| *RPL22L1* | 0.975965068158083 | 3.65388460532006E-08 |
| *A1BG* | 0.815033201970829 | 3.79365625310086E-08 |
| *RPS7* | 1.27046670613126 | 4.08879568480242E-08 |
| *DSTN* | 1.15647155142224 | 4.73158322361328E-08 |
| *SUB1* | 1.10036486822329 | 5.09954182106301E-08 |
| *CORO1C* | 0.662304589254321 | 5.53685997063649E-08 |
| *RPL7A* | 1.292578255066 | 5.68817323770694E-08 |
| *SSR1* | 0.899942539434023 | 5.93216216581487E-08 |
| *POMP* | 0.880160229993997 | 6.12490481356117E-08 |
| *THEMIS2* | 0.802482247302581 | 6.42697770649438E-08 |
| *PSMB3* | 0.897495362162801 | 6.94703124801417E-08 |
| *LAMTOR1* | 0.868755466721747 | 7.32473367960297E-08 |
| *BCAP31* | 0.787603358870317 | 7.32919432910085E-08 |
| *RPS4X* | 1.38395162026137 | 8.1402187249116E-08 |
| *HMGN1* | 1.05018575716822 | 9.27906400163624E-08 |
| *RPS13* | 1.39069289869731 | 1.01938024759886E-07 |
| *LINC00996* | 0.693668760163656 | 1.08065536217977E-07 |
| *CDCA7L* | 0.580774703757723 | 1.08065536217979E-07 |
| *EPHA2* | 0.670209787339667 | 1.08065536217979E-07 |
| *GPM6B* | 0.746966986323705 | 1.08065536217979E-07 |
| *DNASE1L3* | 0.678071905112638 | 1.0806553621798E-07 |
| *NHP2* | 0.891316787233678 | 1.16829908020205E-07 |
| *MEF2C* | 0.969645147918637 | 1.20643713665735E-07 |
| *FKBP1A* | 0.799808113004256 | 1.26141669628754E-07 |
| *ATP13A2* | 0.747193486840834 | 1.27819072841625E-07 |
