## Supplementary Table 16 for "Heterogeneity of circulating epithelial cells in breast cancer at single-cell resolution: identifying tumor and hybrid cells"

Supplementary Table 16. Signaling pathways enriched in diploid CD45$^{-}$ CEC clusters

| Cluster 1 | | |
|---|---|---|
| Term | Adjusted P-value | Genes |
| Antigen processing and presentation | 5.993995555870497E-8 | *CD74;CD4;HSP90AB1;HLA-DPB1;HLA-DRA;HLA-DQA1;HLA-DRB1;CTSB;HLA-DPA1;HLA-DQB1;LGMN* |
| Asthma | 5.993995555870497E-8 | *FCER1G;HLA-DPB1;FCER1A;HLA-DRA;HLA-DQA1;HLA-DRB1;HLA-DPA1;HLA-DQB1* |
| Phagosome | 2.60524495380653E-7 | *ATP6V0B;NCF1;STX7;CYBB;CYBA;HLA-DPB1;HLA-DRA;CD36;SEC61B;HLA-DQA1;HLA-DRB1;HLA-DPA1;HLA-DQB1* |
| Leishmaniasis | 2.60524495380653E-7 | *NCF1;PRKCB;HLA-DPB1;CYBB;HLA-DRA;CYBA;HLA-DQA1;HLA-DRB1;HLA-DPA1;HLA-DQB1* |
| Protein processing in endoplasmic reticulum | 7.778297705178803E-7 | *HSP90AB1;SSR4;DERL3;NGLY1;RRBP1;EIF2AK4;HSP90B1;UBE2J1;HERPUD1;SELENOS;ERP29;SEC61B;TXNDC5* |
| Hematopoietic cell lineage | 2.3610742460519625E-5 | *CD4;IL3RA;HLA-DPB1;HLA-DRA;CD36;HLA-DQA1;HLA-DRB1;HLA-DPA1;HLA-DQB1* |
| Th17 cell differentiation | 3.9132448693656764E-5 | *CD4;HSP90AB1;IRF4;HLA-DPB1;HLA-DRA;HLA-DQA1;HLA-DRB1;HLA-DPA1;HLA-DQB1* |
| Allograft rejection | 5.222723885303661E-5 | *HLA-DPB1;HLA-DRA;HLA-DQA1;HLA-DRB1;HLA-DPA1;HLA-DQB1* |
| Viral myocarditis | 5.254214006180385E-5 | *HLA-DPB1;HLA-DRA;HLA-DQA1;EIF4G2;HLA-DRB1;HLA-DPA1;HLA-DQB1* |
| Graft-versus-host disease | 7.375983509281822E-5 | *HLA-DPB1;HLA-DRA;HLA-DQA1;HLA-DRB1;HLA-DPA1;HLA-DQB1* |
| Coronavirus disease | 7.375983509281822E-5 | *RPS16;PRKCB;RPS8;RPLP0;RPL37A;RPL35;CYBB;TLR7;RPS11;UBA52;RPL6;RPS23* |
| Type I diabetes mellitus | 7.375983509281822E-5 | *HLA-DPB1;HLA-DRA;HLA-DQA1;HLA-DRB1;HLA-DPA1;HLA-DQB1* |
| Lysosome | 9.354106175351658E-5 | *GGA2;ATP6V0B;NPC2;PSAP;CTSZ;MAN2B1;CD68;LGMN;CTSB* |

| | | |
|---|---|---|
| Intestinal immune network for IgA production | 1.220841474830904E-4 | *HLA-DPB1;HLA-DRA;HLA-DQA1;HLA-DRB1;HLA-DPA1;HLA-DQB1* |
| Autoimmune thyroid disease | 2.0450322093584777E-4 | *HLA-DPB1;HLA-DRA;HLA-DQA1;HLA-DRB1;HLA-DPA1;HLA-DQB1* |
| Ribosome | 4.133166069079217E-4 | *RPS16;RPS8;RPLP0;RPL37A;RPL35;RPS11;UBA52;RPL6;RPS23* |
| Th1 and Th2 cell differentiation | 4.8188373045463336E-4 | *CD4;HLA-DPB1;HLA-DRA;HLA-DQA1;HLA-DRB1;HLA-DPA1;HLA-DQB1* |
| Rheumatoid arthritis | 4.880352754940554E-4 | *ATP6V0B;HLA-DPB1;HLA-DRA;HLA-DQA1;HLA-DRB1;HLA-DPA1;HLA-DQB1* |
| Inflammatory bowel disease | 5.258814262599895E-4 | *HLA-DPB1;HLA-DRA;HLA-DQA1;HLA-DRB1;HLA-DPA1;HLA-DQB1* |
| Influenza A | 6.419310662889179E-4 | *PRKCB;IRF7;HLA-DPB1;HLA-DRA;TLR7;HLA-DQA1;HLA-DRB1;HLA-DPA1;HLA-DQB1* |
| Tuberculosis | 8.678375855852277E-4 | *CD74;ATP6V0B;FCER1G;HLA-DPB1;HLA-DRA;HLA-DQA1;HLA-DRB1;HLA-DPA1;HLA-DQB1* |
| Toxoplasmosis | 0.0013017250888193068 | *HLA-DPB1;HLA-DRA;HLA-DQA1;HLA-DRB1;PIK3CG;HLA-DPA1;HLA-DQB1* |
| Diabetic cardiomyopathy | 0.001886387528385928 | *ATP5PF;NCF1;PRKCB;CYBB;CYBA;CD36;COX6A1;ATP5F1E;COX6B1* |
| Parkinson disease | 0.001886387528385928 | *ATP5PF;UBC;GNAS;TXN;PARK7;COX6A1;UBA52;ATP5F1E;UBE2J1;COX6B1* |
| Staphylococcus aureus infection | 0.0032820442779997743 | *HLA-DPB1;HLA-DRA;HLA-DQA1;HLA-DRB1;HLA-DPA1;HLA-DQB1* |
| Cell adhesion molecules | 0.006010205594515829 | *CD4;HLA-DPB1;HLA-DRA;HLA-DQA1;HLA-DRB1;HLA-DPA1;HLA-DQB1* |
| Pathways of neurodegeneration | 0.008109247709738261 | *APP;ATP5PF;PRKCB;CYBB;PARK7;COX6A1;GABARAP;ATP5F1E;UBE2J1;COX6B1;ATG101;UBC;UBA52* |
| NOD-like receptor signaling pathway | 0.017604968161403268 | *HSP90AB1;IRF7;CYBB;CYBA;TXN;GABARAP;CTSB* |
| Systemic lupus erythematosus | 0.017604968161403268 | *HLA-DPB1;HLA-DRA;HLA-DQA1;HLA-DRB1;HLA-DPA1;HLA-DQB1* |

| | | |
|---|---|---|
| Fluid shear stress and atherosclerosis | 0.019688253343894718 | *HSP90AB1;NCF1;GSTP1;CYBA;TXN;HSP90B1* |
| Epstein-Barr virus infection | 0.029501424060740532 | *HLA-DPB1;IRF7;HLA-DRA;HLA-DQA1;HLA-DRB1;HLA-DPA1;HLA-DQB1* |
| Fc epsilon RI signaling pathway | 0.03194071532324141 | *FCER1G;ALOX5AP;FCER1A;GRB2* |
| Lipid and atherosclerosis | 0.03875441024932082 | *HSP90AB1;NCF1;IRF7;CYBB;CYBA;CD36;HSP90B1* |
| Human T-cell leukemia virus 1 infection | 0.04147770364806648 | *CD4;HLA-DPB1;HLA-DRA;HLA-DQA1;HLA-DRB1;HLA-DPA1;HLA-DQB1* |
| | | Cluster 2 |
| Ribosome | 3.41165779812286E-50 | *RPL4;RPL5;RPL30;RPL3;RPL32;RPL34;RPLP1;RPLP0;MRPS10;MRPL36;RPL10A;RPL8;RPL9;RPL6;RPS4X;RPS14;RPL7A;RPS16;RPL18A;RPS19;RPL36AL;RPS18;RPL35;RPL37;RPS11;RPL39;RPS13;RPS12;RPS9;RPL21;RPS7;RPS8;RPL23;RPS5;RPL22;RPS6;MRPS21;RPSA;RPS3A;RPL37A;RPL24;RPL27;RPL26;RPL29;RPL28;UBA52;RPL10;RPL12;RPL11;RPL36A;MRPL14;RPS15A;RPL13;RPL15;RPS2;RPL18;RPS27A;RPL17;RPL19;RPL41;RPL35A;RPS26;RPS28;RPS27;RPL22L1;FAU;RPS21;RPS24;RPS23* |
| Coronavirus disease | 1.5268145182191912E-39 | *RPL4;RPL5;NRP1;RPL30;RPL3;RPL32;RPL34;RPLP1;RPLP0;RPL8;RPL10A;RPL9;RPL6;RPS4X;RPS14;RPL7A;RPS16;RPL18A;RPS19;RPL36AL;RPS18;RPL35;RPL37;RPS11;RPL39;RPS13;RPS12;IFNAR2;RPS9;RPL21;SYK;RPS7;PRKCB;RPS8;RPL23;RPS5;RPL22;RPS6;CYBB;RPSA;RPS3A;RPL37A;RPL24;RPL27;TLR7;RPL26;RPL29;RPL28;UBA52;RPL10;RPL12;RPL11;RPL36A;RPS15A;RPL13;RPL15;RPS2;RPL18;RPS27A;RPL17;RPL19;RPL41;RPL35A;RPS26;RPS28;RPS27;RPL22L1;FAU;RPS21;RPS24;RPS23* |

| | | |
|---|---|---|
| Parkinson disease | 7.49832086611806E-18 | *NDUFA13;NDUFA11;COX4I1;PARK7;COX6A1;UBE2L3;TUBA1B;KIF5B;COX8A;TUBB;NDUFC2;COX6B1;ERN1;PSMA6;COX7A2L;PSMA2;NDUFS6;SLC25A5;UBA52;SLC25A6;NDUFB8;NDUFB11;ATP5MC2;ATP5MC3;NDUFB1;UQCR11;COX7A2;TXN;UQCR10;COX5A;UBE2J1;GNAI2;PSMB7;ATP5F1A;ATP5F1B;UBB;PSMB3;UBC;CYC1;RPS27A;ATP5PB;NDUFA4;NDUFA1;ATP5F1C;ATP5F1D;ATP5F1E;UQCRQ;GNAS;CALM2* |
| Protein processing in endoplasmic reticulum | 1.6212437618232593E-16 | *TRAM1;HSP90AB1;RPN2;UBE2D2;DERL3;RPN1;NGLY1;RRBP1;RNF5;HSP90B1;UBE2J1;HERPUD1;LMAN1;OS9;SEC61G;BAG1;SSR1;MAN1A1;SEC61B;SEC62;SEC63;SEC31A;TXNDC5;PDIA3;BCAP31;SSR4;SSR3;RAD23A;EIF2AK4;PDIA6;PDIA4;RBX1;ERN1;DNAJC3;SELENOS;DAD1;CANX;ERP29;P4HB* |
| Oxidative phosphorylation | 4.534447170210254E-16 | *NDUFA13;NDUFB8;NDUFA11;NDUFB11;COX4I1;ATP5MC2;ATP5MC3;NDUFB1;COX7A2;UQCR11;UQCR10;COX6A1;COX5A;ATP5F1A;ATP5F1B;CYC1;ATP5MG;ATP5MF;ATP5ME;COX8A;ATP6V1G1;ATP6V0B;ATP5PB;NDUFA4;NDUFA1;NDUFC2;ATP5F1C;ATP5F1D;ATP5F1E;COX6B1;COX7A2L;UQCRQ;NDUFS6;ATP6V0C* |
| Diabetic cardiomyopathy | 5.360164082214383E-14 | *NDUFA13;NDUFB8;NCF1;NDUFA11;NDUFB11;COX4I1;ATP5MC2;SLC2A1;ATP5MC3;NDUFB1;UQCR11;COX7A2;UQCR10;COX6A1;COX5A;ATP5F1A;ATP5F1B;CD36;RAC1;CYC1;COX8A;PRKCB;ATP5PB;NDUFA4;PRKCD;NDUFA1;CYBB;NDUFC2;CYBA;ATP5F1C;ATP5F1D;ATP5F1E;COX6B1;COX7A2L;UQCRQ;NDUFS6;SLC25A5;GAPDH;SLC25A6* |
| Prion disease | 5.498528850715846E-13 | *NDUFA13;NDUFB8;NCF1;NDUFA11;NDUFB11;COX4I1;ATP5MC2;ATP5MC3;NDUFB1;UQCR11;COX7A2;UQCR10;COX6A1;COX5A;PSMB7;TUBA1B;ATP5F1A;ATP5F1B;PSMB3;KIF5B;CREB3L2;RAC1;CYC1;COX8A;ATP5PB;NDUFA4;TUBB;PRKCD;NDUFA1;CYBB;NDUFC2;CYBA;ATP5F1C;ATP5F1D;ATP5F1E;COX6B1;PSMA6;COX7A2L;PSMA2;UQCRQ;NDUFS6;CSNK2B;SLC25A5;SLC25A6* |

| | | |
|---|---|---|
| Thermogenesis | 7.813523227544385E-13 | *NDUFA13;NDUFB8;SMARCB1;NDUFA11;NDUFB11;COX4I1;ATP5MC2;ATP5MC3;NDUFB1;UQCR11;COX7A2;UQCR10;COX6A1;COX5A;ACTB;ACTG1;ATP5F1A;ATP5F1B;CREB3L2;COX14;CYC1;ATP5MG;ATP5MF;ATP5ME;COX8A;MAP2K3;ATP5PB;NDUFA4;RPS6;NDUFA1;NDUFC2;ATP5F1C;ATP5F1D;ATP5F1E;COX6B1;COX7A2L;UQCRQ;NDUFS6;GNAS;GRB2* |
| Amyotrophic lateral sclerosis | 1.100349524211426E-12 | *NDUFA13;NDUFA11;COX4I1;COX6A1;ACTB;ACTG1;TUBA1B;KIF5B;RAC1;COX8A;MAP2K3;GABARAPL2;TUBB;ANXA11;NDUFC2;COX6B1;ERN1;PSMA6;COX7A2L;PSMA2;NDUFS6;CAT;ULK1;PFN1;SRSF7;NDUFB8;NDUFB11;ATP5MC2;ATP5MC3;NDUFB1;UQCR11;COX7A2;UQCR10;COX5A;GABARAP;PSMB7;ATP5F1A;ATP5F1B;PSMB3;ATG101;CYC1;BID;HNRNPA1;ATP5PB;NDUFA4;NDUFA1;ATP5F1C;ATP5F1D;ATP5F1E;UQCRQ;HNRNPA2B1* |
| Pathways of neurodegeneration | 2.152918179861841E-12 | *APP;NDUFA13;NDUFA11;COX4I1;PARK7;COX6A1;UBE2L3;TUBA1B;KIF5B;RAC1;MAP2K3;COX8A;GABARAPL2;PRKCB;TUBB;NDUFC2;CYBB;CSNK1E;COX6B1;ERN1;PSMA6;COX7A2L;PSMA2;NDUFS6;CSNK2B;CAT;ULK1;SLC25A5;UBA52;SLC25A6;NDUFB8;NDUFB11;ATP5MC2;NDUFB1;ATP5MC3;UQCR11;COX7A2;UQCR10;COX5A;GABARAP;UBE2J1;PSMB7;ATP5F1A;ATP5F1B;UBB;PSMB3;ATG101;UBC;CYC1;BID;RPS27A;NDUFA4;ATP5PB;NDUFA1;ATP5F1C;ATP5F1D;ATP5F1E;UQCRQ;CALM2* |
| Huntington disease | 5.22234502001218E-12 | *NDUFA13;NDUFB8;NDUFA11;NDUFB11;COX4I1;CLTC;ATP5MC2;CLTA;ATP5MC3;NDUFB1;UQCR11;COX7A2;UQCR10;COX6A1;COX5A;PSMB7;TUBA1B;ATP5F1A;ATP5F1B;PSMB3;ATG101;KIF5B;CREB3L2;AP2S1;CYC1;POLR2L;COX8A;ATP5PB;NDUFA4;TUBB;NDUFA1;NDUFC2;ATP5F1C;ATP5F1D;ATP5F1E;COX6B1;ERN1;PSMA6;COX7A2L;PSMA2;UQCRQ;NDUFS6;ULK1;SLC25A5;SLC25A6* |

| | | |
|---|---|---|
| Alzheimer disease | 2.7142692560457604E-10 | *APP;NDUFA13;NDUFB8;NDUFA11;NDUFB11;COX4I1;ATP5MC2;ATP5MC3;NDUFB1;UQCR11;COX7A2;UQCR10;COX6A1;COX5A;PSMB7;TUBA1B;ATP5F1A;ATP5F1B;PSMB3;ATG101;KIF5B;CYC1;BID;COX8A;ATP5PB;NDUFA4;TUBB;NDUFA1;CYBB;NDUFC2;CSNK1E;ATP5F1C;ATP5F1D;ATP5F1E;COX6B1;ERN1;PSMA6;COX7A2L;PSMA2;UQCRQ;NDUFS6;CSNK2B;ULK1;SLC25A5;CALM2;GAPDH;SLC25A6* |
| Lysosome | 1.3289166780436293E-8 | *SCARB2;CD164;CD63;ATP6V0B;HEXB;CLTC;CTSZ;M6PR;LAPTM5;CLTA;CTSS;GGA2;LAPTM4A;NPC2;PSAP;AP1S2;MAN2B1;TPP1;AP3S1;CD68;ATP6V0C;CTSC;LGMN;CTSB* |
| Phagosome | 1.9267426100090187E-8 | *NCF1;ACTB;CTSS;ACTG1;TUBA1B;HLA-DMA;HLA-DMB;SEC61G;CD36;RAC1;SEC61B;HLA-DQA1;HLA-DPA1;ATP6V1G1;ATP6V0B;TUBB;STX7;M6PR;CYBB;CYBA;CANX;HLA-DPB1;HLA-DRA;ATP6V0C;HLA-DRB1;HLA-DQB1* |
| Protein export | 2.1008948053295677E-7 | *SPCS3;SPCS2;SPCS1;SRP72;SEC61G;SEC61B;SRP14;SEC62;SEC11C;SEC63* |
| Antigen processing and presentation | 3.286117547933189E-7 | *PDIA3;CD74;CIITA;HSP90AB1;CTSS;HLA-DMA;CD4;HLA-DMB;CANX;HLA-DPB1;HLA-DRA;HLA-DQA1;HLA-DRB1;CTSB;HLA-DPA1;HLA-DQB1;LGMN* |
| Viral myocarditis | 2.127905561635449E-6 | *ACTB;ACTG1;ICAM1;HLA-DMA;HLA-DMB;HLA-DPB1;HLA-DRA;RAC1;BID;HLA-DQA1;EIF4G2;HLA-DRB1;HLA-DPA1;HLA-DQB1* |
| Non-alcoholic fatty liver disease | 2.127905561635449E-6 | *COX8A;NDUFA13;NDUFB8;NDUFA11;NDUFB11;NDUFA4;COX4I1;NDUFA1;NDUFC2;NDUFB1;COX7A2;UQCR11;UQCR10;COX6A1;COX5A;COX6B1;ERN1;COX7A2L;UQCRQ;NDUFS6;CYC1;RAC1;BID* |
| Asthma | 4.73559348312127E-6 | *HLA-DMA;HLA-DMB;FCER1G;HLA-DPB1;FCER1A;HLA-DRA;HLA-DQA1;HLA-DRB1;HLA-DPA1;HLA-DQB1* |
| Bacterial invasion of epithelial cells | 8.501999549236407E-6 | *SEPTIN2;ARPC1B;CLTC;CLTA;SEPTIN6;ACTB;SEPTIN11;RHOA;ACTG1;CD2AP;ARPC2;CDH1;ARPC3;ELMO1;RAC1* |

| | | |
|---|---|---|
| Salmonella infection | 9.405434914538123E-6 | *ARF1;HSP90AB1;ARPC1B;BRK1;TXN;ACTB;MYL12A;PIK3CG;MYL12B;ACTG1;HSP90B1;PYCARD;TUBA1B;CYTH4;KIF5B;SNX9;FLNB;RAC1;MAP2K3;TUBB;M6PR;RHOA;ARPC2;ARPC3;TLR9;ELMO1;DYNLRB1;PFN1;GAPDH* |
| Vibrio cholerae infection | 9.846417754474804E-6 | *ARF1;ATP6V0B;ATP6V1G1;KDELR1;SEC61G;GNAS;KDELR2;SEC61B;ATP6V0C;ACTB;ACTG1;PDIA4* |
| Leishmaniasis | 4.142940152262681E-5 | *MARCKSL1;NCF1;PRKCB;CYBB;CYBA;EEF1A1;HLA-DMA;HLA-DMB;HLA-DPB1;HLA-DRA;HLA-DQA1;HLA-DRB1;HLA-DPA1;HLA-DQB1* |
| Shigellosis | 6.137515379037002E-5 | *ARF1;ARPC1B;UBE2D2;ACTB;GABARAP;MYL12A;SEPTIN11;MYL12B;ACTG1;PYCARD;CYTH4;UBB;UBC;RAC1;RPS27A;GABARAPL2;SEPTIN2;PRKCD;SEPTIN6;RHOA;RBX1;ARPC2;ARPC3;ELMO1;UBE2V1;PFN1;UBA52* |
| Influenza A | 1.281248736697152E-4 | *IFNAR2;CIITA;PRKCB;ACTB;ACTG1;ICAM1;PYCARD;DNAJC3;HLA-DMA;HLA-DMB;IRF7;HLA-DPB1;HLA-DRA;TLR7;SLC25A5;BID;HLA-DQA1;HLA-DRB1;HLA-DPA1;HLA-DQB1;SLC25A6* |
| Cardiac muscle contraction | 1.600235828341895E-4 | *COX8A;COX4I1;TPM2;COX7A2;UQCR11;UQCR10;COX6A1;COX5A;COX6B1;COX7A2L;ASPH;SLC9A7;UQCRQ;CYC1* |
| Allograft rejection | 2.198744505211406E-4 | *HLA-DMA;HLA-DMB;HLA-DPB1;GZMB;HLA-DRA;HLA-DQA1;HLA-DRB1;HLA-DPA1;HLA-DQB1* |
| Graft-versus-host disease | 4.989158426466526E-4 | *HLA-DMA;HLA-DMB;HLA-DPB1;GZMB;HLA-DRA;HLA-DQA1;HLA-DRB1;HLA-DPA1;HLA-DQB1* |
| Type I diabetes mellitus | 5.867508523584223E-4 | *HLA-DMA;HLA-DMB;HLA-DPB1;GZMB;HLA-DRA;HLA-DQA1;HLA-DRB1;HLA-DPA1;HLA-DQB1* |
| Hematopoietic cell lineage | 6.120189663979133E-4 | *FLT3;CSF2RA;HLA-DMA;CD4;HLA-DMB;IL3RA;HLA-DPB1;HLA-DRA;CD37;CD36;HLA-DQA1;HLA-DRB1;HLA-DPA1;HLA-DQB1* |

| | | |
|---|---|---|
| Tuberculosis | 6.473797395613023E-4 | *CD74;CIITA;ATP6V0B;FCER1G;SYK;LSP1;CTSS;RHOA;HLA-DMA;HLA-DMB;TLR9;HLA-DPB1;HLA-DRA;BID;CALM2;ATP6V0C;HLA-DQA1;HLA-DRB1;HLA-DPA1;HLA-DQB1* |
| Fluid shear stress and atherosclerosis | 6.549900916377609E-4 | *MEF2C;HSP90AB1;NCF1;GSTP1;MGST2;CYBA;TXN;ACTB;RHOA;ACTG1;ICAM1;HSP90B1;SUMO3;SUMO2;PECAM1;RAC1;CALM2* |
| Leukocyte transendothelial migration | 0.0025640674006215365 | *NCF1;PRKCB;CYBB;CYBA;ACTB;MYL12A;RHOA;ACTG1;ICAM1;MYL12B;GNAI2;PECAM1;RAC1;CD99* |
| Autoimmune thyroid disease | 0.0027032855375961123 | *HLA-DMA;HLA-DMB;HLA-DPB1;GZMB;HLA-DRA;HLA-DQA1;HLA-DRB1;HLA-DPA1;HLA-DQB1* |
| Cell adhesion molecules | 0.003883941420326298 | *ICAM1;HLA-DMA;CD4;ALCAM;HLA-DMB;SELL;CDH1;PECAM1;HLA-DPB1;HLA-DRA;CD99;HLA-DQA1;HLA-DRB1;HLA-DPA1;HLA-DQB1;NECTIN1* |
| Rheumatoid arthritis | 0.003883941420326298 | *ATP6V0B;ATP6V1G1;HLA-DMA;HLA-DMB;HLA-DPB1;HLA-DRA;ATP6V0C;HLA-DQA1;HLA-DRB1;ICAM1;HLA-DPA1;HLA-DQB1* |
| Kaposi sarcoma-associated herpesvirus infection | 0.003883941420326298 | *LYN;IFNAR2;GABARAPL2;SYK;GABARAP;PIK3CG;ICAM1;UBB;GNG5;GNG7;MAPKAPK2;UBC;GNB1;IRF7;RAC1;BID;RPS27A;UBA52;CALM2* |
| Platelet activation | 0.005271130436579866 | *LYN;FCER1G;SYK;ACTB;MYL12A;PIK3CG;RHOA;ACTG1;MYL12B;GNAI2;APBB1IP;VAMP8;GNAS;FERMT3* |
| Fc gamma R-mediated phagocytosis | 0.005372149707833894 | *LYN;MARCKSL1;GSN;NCF1;ARPC2;SYK;PRKCB;ARPC3;ARPC1B;CFL1;PRKCD;RAC1* |
| Intestinal immune network for IgA production | 0.005578936829613176 | *HLA-DMA;HLA-DMB;HLA-DPB1;HLA-DRA;HLA-DQA1;HLA-DRB1;HLA-DPA1;HLA-DQB1* |
| Epstein-Barr virus infection | 0.0061159943111627705 | *LYN;IFNAR2;MAP2K3;PDIA3;SYK;ICAM1;HLA-DMA;HLA-DMB;HLA-DPB1;BLNK;IRF7;HLA-DRA;RAC1;VIM;BID;HLA-DQA1;HLA-DRB1;HLA-DPA1;HLA-DQB1* |

| | | |
|---|---|---|
| Human T-cell leukemia virus 1 infection | 0.00639966696053165 | *NRP1;RANBP1;SLC2A1;ICAM1;ANAPC11;HLA-DMA;CD4;HLA-DMB;CREB3L2;CANX;HLA-DPB1;HLA-DRA;ANAPC5;TCF3;SLC25A5;HLA-DQA1;HLA-DRB1;HLA-DPA1;HLA-DQB1;SLC25A6* |
| Spliceosome | 0.010706171787007022 | *SF3B5;EIF4A3;SNU13;LSM4;CDC40;LSM3;HNRNPM;HNRNPK;SNRPD2;HNRNPC;TXNL4A;SRSF6;HNRNPA1;SRSF7;SRSF9* |
| Mitophagy | 0.013282131408728245 | *GABARAPL2;FIS1;UBB;CSNK2B;UBC;ULK1;RPS27A;UBA52;GABARAP* |
| Toxoplasmosis | 0.0163136469490332 | *MAP2K3;CIITA;HLA-DMA;HLA-DMB;HLA-DPB1;HLA-DRA;HLA-DQA1;HLA-DRB1;PIK3CG;HLA-DPA1;HLA-DQB1;GNAI2* |
| Retrograde endocannabinoid signaling | 0.02347161798284537 | *NDUFA13;NDUFB8;NDUFA11;PRKCB;NDUFB11;NDUFA4;NDUFA1;NDUFC2;NDUFB1;GNAI2;GNG5;NDUFS6;GNG7;GNB1* |
| Pathogenic Escherichia coli infection | 0.023981787659981792 | *ARF1;TMED10;ARPC1B;TUBB;BRK1;ACTB;RHOA;ACTG1;PYCARD;TUBA1B;CYTH4;ARPC2;ARPC3;GNA12;TMBIM6;RAC1;GAPDH* |
| Th17 cell differentiation | 0.03087514922025053 | *HLA-DMA;CD4;HSP90AB1;HLA-DMB;IRF4;HLA-DPB1;HLA-DRA;HLA-DQA1;HLA-DRB1;HLA-DPA1;HLA-DQB1* |
| Various types of N-glycan biosynthesis | 0.03087514922025053 | *RPN2;DAD1;HEXB;RPN1;MAN1A1;MGAT1* |
| Inflammatory bowel disease | 0.03198979336559227 | *HLA-DMA;HLA-DMB;HLA-DPB1;HLA-DRA;HLA-DQA1;HLA-DRB1;HLA-DPA1;HLA-DQB1* |
| Ubiquitin mediated proteolysis | 0.033677165130778626 | *UBE2D2;UBE2E3;UBE2E2;UBE2L3;RBX1;ANAPC11;UBE2J1;UBB;UBC;ANAPC5;ELOB;RPS27A;UBA52* |
| B cell receptor signaling pathway | 0.036654496555207174 | *LYN;SYK;PRKCB;BLNK;LILRB1;GRB2;RAC1;LILRB4;LILRA4* |
| Ferroptosis | 0.036654496555207174 | *GPX4;FTH1;PCBP2;CYBB;SLC3A2;FTL* |
| Osteoclast differentiation | 0.037543183745167276 | *IFNAR2;TYROBP;NCF1;SYK;BLNK;CYBA;GRB2;LILRB1;RAC1;LILRB4;SIRPB1;LILRA4* |
| Fc epsilon RI signaling pathway | 0.03817763426546309 | *MAP2K3;LYN;FCER1G;SYK;ALOX5AP;FCER1A;GRB2;RAC1* |
| Human immunodeficiency virus 1 infection | 0.0414022356608123 | *MAP2K3;PDIA3;PRKCB;SAMHD1;GNAI2;RBX1;BST2;CD4;GNG5;GNG7;CFL1;GNB1;AP1S2;ELOB;RAC1;BID;CALM2* |
| Glutathione metabolism | 0.04719953333025442 | *GPX4;GSTP1;ODC1;MGST2;LAP3;PGD;PRDX6* |
