## Supplementary Table 18 for "Heterogeneity of circulating epithelial cells in breast cancer at single-cell resolution: identifying tumor and hybrid cells"

Supplementary Table 18. Upregulated genes in diploid cells of $CD45^+$ CEC clusters

| Genes | Cluster 1 | | Genes | Cluster 2 | | Genes | Cluster 3 | |
|---|---|---|---|---|---|---|---|---|
| | LogFC | Adjusted P-value | | LogFC | Adjusted P-value | | LogFC | Adjusted P-value |
| *CD74* | 4.29373205295416 | 1.45539423384385E-69 | *CD74* | 4.27651028719235 | 4.95341904464289E-65 | *SNHG29* | 3.1114665036251 | 5.96364437215464E-18 |
| *HLA-DRA* | 3.55642143498279 | 2.06591328642268E-60 | *HLA-DRA* | 3.60242125639387 | 3.74926734715927E-58 | *NPM1* | 1.92622708070612 | 1.18675695795195E-11 |
| *CST3* | 2.71798450963031 | 2.67611953962921E-50 | *ITM2C* | 2.78355411235368 | 1.54388856909492E-57 | *AIF1* | 1.76342450383376 | 1.25452437002452E-11 |
| *HLA-DRB1* | 2.98359235789038 | 9.65853304035756E-49 | *HLA-DPA1* | 2.93551857994256 | 9.98990876918739E-54 | *RPLP0* | 2.10867738531459 | 1.86591718885703E-11 |
| *HLA-DPA1* | 2.75087756667122 | 8.96096061010688E-46 | *TCF4* | 2.77614480385524 | 5.2583744865145E-52 | *HSP90AB1* | 2.19314581088231 | 6.26551326204804E-11 |
| *SEC61B* | 2.34113088917794 | 9.60742443742708E-43 | *PPP1R14B* | 2.97437270661948 | 7.29527505128806E-52 | *GAPDH* | 2.19186755432299 | 1.92698334140158E-10 |
| *CYBA* | 2.04742090460962 | 4.00261439688184E-42 | *NPC2* | 2.63117705570398 | 1.06014647877394E-51 | *YBX1* | 1.72492584468282 | 3.75857716979063E-09 |
| *NPC2* | 2.5631742432393 | 5.79337543117612E-42 | *HLA-DPB1* | 2.52497202303037 | 1.94621180832509E-50 | *YBX3* | 1.20391518208752 | 5.10869890823066E-09 |
| *ATP5F1E* | 1.62816967685483 | 1.37683821837928E-41 | *BCL11A* | 2.03382541162842 | 4.89919864814281E-50 | *STMN1* | 1.38967912974313 | 6.67283417070089E-09 |
| *TCF4* | 2.93407943153368 | 1.4167357802119E-41 | *HLA-DRB1* | 3.13763247421875 | 1.14092302898686E-49 | *BTF3* | 1.78177079631442 | 7.01245026427551E-09 |
| *HLA-DPB1* | 2.36968239190926 | 2.35485712245474E-41 | *SEC61B* | 2.4003631739611 | 1.40434878667233E-49 | *RPL15* | 1.82132489740539 | 8.34665216711862E-09 |
| *MT-CO1* | 1.83330567000968 | 4.31057443578051E-41 | *JCHAIN* | 3.18856009559444 | 2.33802945442548E-48 | *RACK1* | 1.95055641006786 | 1.01724740912691E-08 |
| *GRN* | 2.05738513352265 | 2.2181757921601E-39 | *CCDC50* | 2.49601643855511 | 8.7749572360115E-48 | *GIHCG* | 1.93118764160785 | 1.47469152994057E-08 |
| *JCHAIN* | 3.16905560060795 | 2.56081865876742E-39 | *UGCG* | 2.3857559786505 | 2.42787499251813E-47 | *HNRNPA1* | 1.74177609891724 | 1.58980148478435E-08 |
| *UGCG* | 2.55991362510964 | 8.13403132611675E-39 | *PLD4* | 2.38269942021716 | 1.76117884589083E-45 | *EEF1G* | 1.67340046538874 | 3.94265266173658E-08 |
| *ITM2C* | 2.72405227536304 | 1.19775950166577E-38 | *SERPINF1* | 2.32635482776404 | 3.36454563354095E-45 | *HMGA1* | 1.28812218632313 | 3.95220957089635E-08 |
| *MPEG1* | 1.89902426532427 | 2.41108709574682E-38 | *CST3* | 2.67578295474775 | 6.61307727245744E-45 | *SERPINB1* | 2.51419394536368 | 5.15533788404645E-08 |
| *HLA-DQB1* | 1.89740379445366 | 2.47647835786758E-38 | *IGKC* | 2.35769831900128 | 3.3547153759287E-44 | *CAT* | 1.3037898745663 | 5.60465350281564E-08 |
| *GNAS* | 1.87612527867852 | 4.89900588724212E-38 | *C12orf75* | 2.40525423922678 | 3.58026153785531E-44 | *CDK6* | 1.91332505373316 | 6.07050932474477E-08 |
| *CCDC88A* | 2.10626522771883 | 6.57886931375001E-38 | *HERPUD1* | 2.12403461401113 | 7.81247628335867E-43 | *ACTG1* | 1.87367745362424 | 9.3753163568532E-08 |
| *PPP1R14B* | 2.82663482447282 | 8.35290168854407E-38 | *IRF8* | 2.1717877454595 | 9.4002325030363E-43 | *SOX4* | 2.45477933935514 | 1.14961755413138E-07 |
| *CCDC50* | 2.66440852851252 | 1.18420254959162E-37 | *IRF7* | 2.20999481620423 | 4.1840369073758E-41 | *RPS24* | 1.98372945499653 | 1.58033879140983E-07 |
| *RPS8* | 1.66776113359692 | 1.9507247960484E-37 | *LILRA4* | 2.04519273274219 | 5.37954240262608E-41 | *ZFAS1* | 1.66446042103553 | 1.65450055873727E-07 |
| *PLD4* | 2.38462043283638 | 1.79323996522963E-36 | *CCDC88A* | 1.92625166001869 | 1.08876752593105E-40 | *PRSS57* | 1.47754470316051 | 1.78181497466538E-07 |
| *MT-ND1* | 1.62423741872111 | 1.436798416665E-35 | *APP* | 2.00686951820489 | 1.08876879533311E-40 | *EEF1B2* | 1.70584767998545 | 2.8556933376525E-07 |
| *PSAP* | 1.94911883099782 | 1.48397130432801E-35 | *GNAS* | 1.84855667757131 | 1.75835584065425E-39 | *UQCRH* | 1.47788629071766 | 2.89842425051721E-07 |
| *BCL11A* | 2.195840581696 | 2.16353069538012E-35 | *ALOX5AP* | 2.28792964616274 | 4.41115426104413E-39 | *SLC25A3* | 1.57942431717973 | 2.97562939853724E-07 |
| *PTPRE* | 2.00754327462819 | 2.64295923124342E-35 | *SPCS1* | 1.90583069960819 | 3.98337103696719E-38 | *ANP32B* | 1.48911354614092 | 3.33477512395441E-07 |
| *CYBB* | 1.81787429771956 | 4.10032811079976E-35 | *FTH1* | 1.83213302505727 | 1.80251330694048E-37 | *NME2* | 1.58621348284237 | 4.32109759209803E-07 |
| *FTH1* | 2.06662715538549 | 5.0832343783065E-35 | *PLAC8* | 2.10545115516367 | 3.39367502545819E-37 | *ATP5MC2* | 1.49917706372529 | 4.40403969798746E-07 |
| *HERPUD1* | 2.36345718073989 | 9.7889825229903E-35 | *CYB561A3* | 1.77644467571401 | 2.86229849537658E-36 | *H2AFY* | 1.57037917757672 | 5.22238058057849E-07 |
| *RPL6* | 1.61779145214118 | 1.36284659344626E-34 | *CYBA* | 2.04095867511886 | 3.50419559185563E-36 | *HMGB1* | 1.68392624362399 | 6.21924958232921E-07 |
| *C12orf75* | 2.34303303871091 | 1.90396482358255E-34 | *GRN* | 1.89785327127609 | 3.66849809001415E-36 | *YWHAE* | 1.18514537806768 | 7.40172177000244E-07 |
| *RPS11* | 1.56245307599564 | 3.26072711594422E-34 | *ATP5F1E* | 1.61228298538179 | 2.40314978433789E-35 | *H3F3A* | 1.51148548852315 | 1.01364536735186E-06 |
| *APP* | 2.18812596618516 | 3.98376353612844E-34 | *ERP29* | 1.788859726557 | 2.43654596728593E-35 | *RPL4* | 1.49469457840422 | 0.0000012443155647556 |

| | | | | | | | | |
|---|---|---|---|---|---|---|---|---|
| *IRF8* | 2.37872525864302 | 4.10347415408458E-34 | *PSAP* | 1.63841386318432 | 1.16782866836948E-34 | *TKT* | 1.14600828858603 | 1.95733999195378E-06 |
| *SERPINF1* | 2.1124130852078 | 1.48965844884573E-33 | *PTPRE* | 1.71124189295105 | 3.85128196762149E-34 | *EEF2* | 1.47804729680464 | 2.14552600328874E-06 |
| *PABPC1* | 1.37271834378008 | 1.61566024042927E-33 | *MAPKAPK2* | 1.67909242299739 | 3.77567035243775E-33 | *ANKRD28* | 1.16438681790088 | 3.32343409792001E-06 |
| *RPS24* | 1.40009160183893 | 2.32920146319268E-33 | *MPEG1* | 1.56089772705329 | 4.17433033371546E-33 | *COX6B1* | 1.35507037950999 | 3.51972099220222E-06 |
| *IRF7* | 2.17246510032993 | 4.40076873554365E-33 | *TXN* | 2.06148394329484 | 2.02211616745951E-32 | *RPLP1* | 1.6943173292704 | 0.0000048850013067018 |
| *HLA-DMA* | 1.58056245654738 | 1.5452248333366E-32 | *GZMB* | 2.32469547146756 | 3.37324059989067E-32 | *HINT1* | 1.33753572021748 | 0.0000049970899255723 |
| *PLAC8* | 2.06749448242307 | 2.34395046885293E-32 | *TSPAN13* | 1.39710248534414 | 4.71648829968494E-32 | *TXN* | 1.25954405094122 | 5.55742240175415E-06 |
| *PPIA* | 1.50747718062191 | 2.9480225096947E-32 | *MZB1* | 1.65499674737993 | 1.22079548012409E-31 | *CLIC1* | 1.55866575701293 | 5.85393761054742E-06 |
| *RPLP0* | 1.5214777440877 | 3.37419823038537E-32 | *SELENOS* | 1.66787361661611 | 2.20494549615071E-31 | *RPL7* | 1.32892258991277 | 5.87154873331416E-06 |
| *SRP14* | 1.61469979192726 | 9.62146723558954E-32 | *HSP90B1* | 1.65420601288992 | 4.12184111852398E-31 | *RPS4X* | 1.87292400374057 | 5.91777821236664E-06 |
| *RPL15* | 1.30926209876785 | 1.23207127685975E-31 | *RNASET2* | 1.63136038311269 | 4.54647353926297E-31 | *RPS8* | 1.79180472798465 | 6.40799049480657E-06 |
| *LILRA4* | 1.99309652805842 | 1.78570788068362E-31 | *TYROBP* | 1.56760273535795 | 1.01759607019496E-30 | *RPL8* | 1.71230233186672 | 0.0000068001061719181 |
| *EEF1A1* | 1.36207419231035 | 4.57670491464137E-31 | *HLA-DMA* | 1.53742715099875 | 5.91904525784923E-30 | *RNF130* | 1.20446425727622 | 7.72010489739395E-06 |
| *ALOX5AP* | 2.19262027892836 | 5.21588279661305E-31 | *DERL3* | 1.38866097845588 | 7.91599410010985E-30 | *PPIA* | 1.3949997460423 | 7.80133308360163E-06 |
| *IGKC* | 2.64898681276625 | 6.96129600098354E-31 | *TPM2* | 2.00014977368722 | 1.17098239660896E-29 | *SLC25A6* | 1.38322141982414 | 0.0000106587458591063 |
| *UBE2J1* | 1.56287592262874 | 8.50340618192732E-31 | *IL3RA* | 1.4744245017381 | 1.46884688778059E-29 | *IMPDH2* | 1.12066544047156 | 0.0000111375314440676 |
| *RACK1* | 1.26470828131449 | 9.43902011721939E-31 | *SMPD3* | 1.34852230489908 | 1.48249527443151E-29 | *GLUL* | 1.37779045601008 | 0.0000130840642868637 |
| *NCF1* | 1.69164931168155 | 1.81569085323094E-30 | *VAMP8* | 1.62350646125211 | 4.25284512862185E-29 | *RPS5* | 1.5522028484447 | 0.0000137814870003772 |
| *TXN* | 2.01063322021052 | 3.185198757644E-30 | *SFT2D2* | 1.3952994569968 | 9.14789507773349E-29 | *LAPTM4B* | 0.869939459435627 | 0.0000153704554213148 |
| *GABARAP* | 1.66539352885958 | 4.41469422256629E-30 | *NCF1* | 1.43886331651516 | 1.61281420470195E-28 | *ENO1* | 1.44268798038799 | 0.0000172020623921878 |
| *TAGLN2* | 1.70607590035149 | 4.7080792473931E-30 | *SPIB* | 1.37188790138309 | 2.13250011157954E-28 | *GAS5* | 1.51986747249927 | 0.000023148180965188 |
| *CYB561A3* | 1.75414870010918 | 2.52977337766278E-29 | *VEGFB* | 1.59867700199018 | 2.42537802901375E-28 | *RPL5* | 1.74421333939613 | 0.0000263584999935807 |
| *RPS9* | 1.20567978299683 | 2.82803860388876E-29 | *IRF4* | 1.83532650288908 | 3.29703825557698E-28 | *APEX1* | 1.2768615471593 | 0.000027892369777772 |
| *UBC* | 1.37561544565172 | 2.90995378986024E-29 | *RPLP0* | 1.57754060729993 | 4.54700933699129E-28 | *RPL6* | 1.65027464365018 | 0.0000345929689959982 |
| *RPL37A* | 1.34687833884482 | 3.28447322681571E-29 | *CYBB* | 1.43886331651516 | 9.29658576623089E-28 | *RPL7A* | 1.68099052595801 | 0.0000463155887008235 |
| *EIF4A1* | 1.4418136586321 | 4.27475022282771E-29 | *PPIA* | 1.46334060701142 | 9.72842873012465E-28 | *SLC25A5* | 1.35314846135605 | 0.0000487890848254178 |
| *ERP29* | 1.67748840147564 | 4.65953189061706E-29 | *SSR4* | 1.57054135191233 | 1.01682088358544E-27 | *ATP5F1A* | 0.916459304457297 | 0.0000489733015297135 |
| *SFT2D2* | 1.65279983345195 | 5.40458845286932E-29 | *SCT* | 1.72780873753818 | 1.17218311872392E-27 | *RPL22L1* | 1.02303096865534 | 0.0000542917306286314 |

| | | | | | | | | |
|---|---|---|---|---|---|---|---|---|
| *H3F3A* | 1.24490714915147 | 5.42816991433493E-29 | *UBE2J1* | 1.36646377290176 | 1.23111578203742E-27 | *GATA2* | 1.0741890089293 | 0.0000549542388645328 |
| *MT-CO2* | 1.32282160944979 | 6.2393136795427E-29 | *TAGLN2* | 1.66035495239112 | 2.0926412027134E-27 | *ATP5F1C* | 0.991919621896117 | 0.0000658776189721527 |
| *PLP2* | 1.51700781836003 | 6.39445026546164E-29 | *CTSZ* | 1.34026415389677 | 2.12095466112643E-27 | *EIF3H* | 1.19609567762822 | 0.00006885628741647 |
| *HSP90B1* | 1.9141123811699 | 6.49860559259517E-29 | *RPS8* | 1.68662855737547 | 2.79050675342529E-27 | *TALDO1* | 1.200912693926 | 0.0000705611510646686 |
| *FCER1G* | 1.72993871475345 | 7.5616149718772E-29 | *FCER1G* | 1.77195289372577 | 6.56082208076727E-27 | *RPL29* | 1.60923812124595 | 0.0000755515981204137 |
| *TYROBP* | 1.74506621504146 | 7.91039439663765E-29 | *CTSB* | 1.33965746968224 | 1.36008960185001E-26 | *SPINT2* | 0.826218082006309 | 0.0000778581726566068 |
| *MT-ND4* | 1.34339777010986 | 8.33248303609318E-29 | *SRP14* | 1.52166602398532 | 2.0055423184956E-26 | *RPS2* | 1.62293556824009 | 0.0000814718513096095 |
| *RPS28* | 1.41327789808378 | 9.16240996847065E-29 | *CD2AP* | 1.25635368635175 | 2.81018488523158E-26 | *CDK2AP1* | 0.869939459435627 | 0.0000817996899260839 |
| *TGFBI* | 1.43151159807842 | 4.37288320586981E-28 | *RPS11* | 1.35516623646102 | 3.19783050705639E-26 | *NDUFA11* | 0.96085342381575 | 0.0000893753005240895 |
| *NACA* | 1.46435976363625 | 4.86305791777761E-28 | *TRAF4* | 1.52616450220582 | 3.96408897070951E-26 | *SUMO2* | 1.18947779886371 | 0.000109511062333688 |
| *CTSB* | 1.48119575146688 | 5.50079898236011E-28 | *PTCRA* | 1.46594666717956 | 1.49107127619703E-25 | *RPS23* | 1.63547207368603 | 0.00011601675386554 |
| *SSR4* | 1.65718138754807 | 8.6538744171928E-28 | *TGFBI* | 1.25347405457556 | 1.93673532646004E-25 | *EIF4A1* | 1.43830953987741 | 0.000118078524895016 |
| *IL3RA* | 1.51031316578602 | 2.43834069624127E-27 | *PARK7* | 1.63727224730276 | 3.23471929617723E-25 | *RPL12* | 1.56099159908274 | 0.000135891664228421 |
| *RNASET2* | 1.6004363342532 | 2.72930306857609E-27 | *OAZ1* | 1.29349911491472 | 3.75449857581942E-25 | *GNG5* | 0.901352412067088 | 0.000161999517421888 |
| *CTSS* | 2.09630490814509 | 4.20437367727047E-27 | *CLEC4C* | 1.0635832057372 | 6.05913425049268E-25 | *RPL10A* | 1.41650544625549 | 0.000165792342639952 |
| *SERF2* | 1.26187736812414 | 4.39627575537713E-27 | *BRI3* | 1.39708282621473 | 1.26738458035483E-24 | *SLC40A1* | 1.12256664220626 | 0.000168059620060165 |
| *IRF4* | 2.08558107354709 | 5.42680144372263E-27 | *LRRC26* | 1.25497365707571 | 1.52887042721912E-24 | *ATP5F1B* | 1.02688329415095 | 0.000191368132902053 |
| *MT-ND2* | 1.40315589576192 | 6.30873492481382E-27 | *TXNDC5* | 1.19028408907689 | 1.52887042721925E-24 | *RPL22* | 1.44522404625443 | 0.000217875189998684 |
| *MAN2B1* | 1.40659159659706 | 6.77229392269268E-27 | *SERF2* | 1.34209556644469 | 1.54623166606736E-24 | *SNHG5* | 1.1401392716542 | 0.000235613817127627 |
| *CD4* | 1.51929231320667 | 1.10201701523477E-26 | *PLP2* | 1.49618847602976 | 2.00367449286524E-24 | *RPS7* | 1.51187154553399 | 0.000243119001806647 |
| *CD68* | 1.29721050636683 | 1.27079992046547E-26 | *RPL6* | 1.66695338479681 | 3.46501570616452E-24 | *ATP5MPL* | 0.930896687681104 | 0.000251359815098196 |
| *TSPAN13* | 1.40068220630177 | 1.61313352582628E-26 | *CXCR3* | 1.35345460044131 | 3.80759892214795E-24 | *SNHG6* | 1.35581288855412 | 0.000257186247069394 |
| *CXXC5* | 1.49383186389957 | 1.68459863996244E-26 | *GABARAP* | 1.60396041836459 | 5.34996830862164E-24 | *GPX4* | 1.05613392724886 | 0.000276634842910739 |
| *GZMB* | 2.20504431432198 | 1.70838201993138E-26 | *PPIB* | 1.44754491459468 | 7.95500841456243E-24 | *DUT* | 0.951842182520782 | 0.000332112162989415 |
| *SPIB* | 1.38943573872502 | 1.76202485980828E-26 | *H3F3A* | 1.12866938159387 | 1.76990755983111E-23 | *CTNNBL1* | 1.12256664220626 | 0.000339600297801101 |
| *VAMP8* | 1.54496964786171 | 1.90047604813076E-26 | *STMN1* | 1.26173923463521 | 1.82538248381263E-23 | *ST13* | 1.26872347771562 | 0.000385866882824582 |
| *MZB1* | 1.69164931168155 | 1.90378630914373E-26 | *CNPY3* | 1.28260358550957 | 3.3000435842007E-23 | *LMO2* | 0.814444346843923 | 0.00041508720622197 |
| *VIM* | 1.91896286509932 | 2.71341451898261E-26 | *HLA-DQB1* | 1.64602828365804 | 4.77338162267378E-23 | *SMIM24* | 0.726981505593584 | 0.00041508720622197 |
| *TPM2* | 1.88831555193562 | 3.97284614386775E-26 | *SOX4* | 1.86743989379322 | 7.14993419269039E-23 | *PBX1* | 0.633872101202103 | 0.000415087206221973 |
| *MAPKAPK2* | 1.73828063628945 | 6.09292613181233E-26 | *VIM* | 1.55573511600685 | 8.17270887251255E-23 | *RPL31* | 1.18026985950754 | 0.00042216218968662 |
| *RPS2* | 1.23597522080943 | 7.00001522238135E-26 | *MAN2B1* | 1.24597585812373 | 8.25700526242024E-23 | *RPL35A* | 1.45246714230457 | 0.000444660080839766 |
| *H3F3B* | 1.24815171565074 | 9.38447433599691E-26 | *MS4A6A* | 1.46310092681136 | 9.16294929970974E-23 | *RPS6* | 1.57149304354351 | 0.000472547267630339 |
| *SELENOS* | 1.7080269780447 | 1.11436865667599E-25 | *TM9SF2* | 1.04998228380787 | 1.16094927261975E-22 | *RBIS* | 0.881452854629383 | 0.000555581995317406 |

| | | | | | | | | |
|---|---|---|---|---|---|---|---|---|
| *RPS7* | 1.12939010760947 | 1.23374108840977E-25 | *CXXC5* | 1.33409878109105 | 1.22275369408322E-22 | *EIF3E* | 1.17616398567724 | 0.000662320461845458 |
| *PHACTR1* | 1.32043495284734 | 1.30428648917332E-25 | *GAS6* | 1.12832409697554 | 1.29456177584281E-22 | *TMEM256* | 0.670572205237922 | 0.000691166161469004 |
| *ACTB* | 1.36810197262124 | 1.36084238874821E-25 | *NME2* | 1.48840242927064 | 1.39759203684277E-22 | *GTF2I* | 1.01682962948702 | 0.000707117969377993 |
| *OAZ1* | 1.46127450833467 | 1.76182342692928E-25 | *RNASE6* | 1.06959027218507 | 1.98528836589301E-22 | *NACA* | 1.37939970887173 | 0.000751128081780765 |
| *PARK7* | 1.52675083498942 | 1.77037563106921E-25 | *CD4* | 1.45191946934061 | 3.0393262796981E-22 | *MT-CO1* | 1.3852762752084 | 0.000765979476020856 |
| *SPCS1* | 1.79805285680682 | 2.5278031371232E-25 | *ARPC3* | 1.34966516302326 | 3.37186845019818E-22 | *UQCRB* | 1.07692397665054 | 0.000816597626411055 |
| *CFL1* | 1.29921305686501 | 3.34238394246608E-25 | *EEF1A1* | 1.3635203232834 | 3.75479269275062E-22 | *RPS11* | 1.20153981631835 | 0.000835171937011579 |
| *MYL6* | 1.2791689652949 | 3.88631570304607E-25 | *SLC15A4* | 1.36326889026826 | 3.98065132161056E-22 | *LEPROT* | 0.867993815120658 | 0.000835807450365672 |
| *IRF2BP2* | 1.46406474508811 | 4.1319026225394E-25 | *HLA-DQA1* | 1.62456971362842 | 4.58721821792149E-22 | *EEF1A1* | 1.57779507359507 | 0.000931530528933108 |
| *RNF130* | 1.34197371049064 | 5.1269315027862E-25 | *RNF130* | 1.22935414774209 | 5.5901389942116E-22 | *SNHG7* | 1.21952837209334 | 0.000975739974985742 |
| *MS4A6A* | 1.52921557654716 | 5.45153293103037E-25 | *IGFLR1* | 1.22228822177403 | 6.80816914535757E-22 | *GNAS* | 1.18576046869345 | 0.0010030412166019 |
| *RPL37* | 1.24204415842026 | 9.06288854131367E-25 | *MT-CO1* | 1.37294369473545 | 7.72058083462244E-22 | *ATP5MG* | 1.01021588777312 | 0.00102171346786963 |
| *DPYSL2* | 1.24448609961404 | 1.04805569188283E-24 | *HINT1* | 1.37974955347125 | 9.43904973449582E-22 | *CYTOR* | 1.08072888885397 | 0.00113113038366498 |
| *TXNDC5* | 1.3342548445628 | 1.18038410556125E-24 | *RPL15* | 1.28883075712099 | 1.09727358628099E-21 | *RPL37A* | 1.38749717657033 | 0.00121799967361988 |
| *PLEK* | 1.81925769541522 | 1.21828848504325E-24 | *SEC61G* | 1.31768561979318 | 1.10538052582083E-21 | *HSBP1* | 0.568777072980218 | 0.00123130113732357 |
| *ARPC3* | 1.36713403007656 | 1.42635943702537E-24 | *MYL6* | 1.25516361659287 | 1.23788996581607E-21 | *RPL24* | 1.28627912292139 | 0.00143340195582213 |
| *VEGFB* | 1.46369200382757 | 1.51658525587162E-24 | *CD68* | 1.20038373624729 | 1.45672065577338E-21 | *SNRPD2* | 0.954463748526651 | 0.00143543436932569 |
| *CTSZ* | 1.47370262447881 | 2.45957491793428E-24 | *MT-ND1* | 1.11248952354781 | 1.50742553667186E-21 | *LDHB* | 1.34041718199238 | 0.00160332704556095 |
| *SCT* | 1.63717482468308 | 3.76200255011424E-24 | *OSTC* | 1.13460624795485 | 1.65147741389769E-21 | *IL18* | 0.798737345584202 | 0.00182444488232825 |
| *RNASEK* | 1.15603026554526 | 3.77719730261236E-24 | *EIF4A1* | 1.17997219405324 | 2.37188892226262E-21 | *HSPD1* | 1.087424836086 | 0.00198493352817093 |
| *RPS23* | 1.28451645480041 | 3.79105875465591E-24 | *DAB2* | 1.0494686762643 | 2.4380142969056E-21 | *RPL18A* | 1.46548499036581 | 0.00200552578238534 |
| *MT-CO3* | 1.39444958439113 | 6.82650306649002E-24 | *PHACTR1* | 1.21125227964639 | 3.50986109472715E-21 | *ETV6* | 0.826218082006309 | 0.00200727592673703 |
| *EEF2* | 1.33027033612425 | 7.81869314657405E-24 | *COBLL1* | 1.22288573474214 | 4.23970053779519E-21 | *CNRIP1* | 0.601450623509725 | 0.00201607788311072 |
| *LYN* | 1.39505033822244 | 8.05556349266568E-24 | *CCDC186* | 1.13828069255072 | 8.36345761467012E-21 | *UROD* | 0.499571009490512 | 0.00201607788311072 |
| *DERL3* | 1.4164397212094 | 8.103159829381E-24 | *CAT* | 1.1022884456634 | 9.06742823975188E-21 | *FHL1* | 0.869939459435627 | 0.00201607788311073 |
| *SEL1L3* | 1.22646783376657 | 9.1087118027481E-24 | *SPINT2* | 1.07111048321811 | 1.14785648472972E-20 | *DIPK1B* | 0.601450623509725 | 0.00201607788311074 |
| *SMPD3* | 1.43454565816806 | 9.1087118027481E-24 | *SEL1L3* | 1.06157527912095 | 1.23166770389837E-20 | *ENSA* | 0.730490291370723 | 0.00267561625860648 |
| *RPS3A* | 1.3482327409833 | 9.21328699250779E-24 | *RPS28* | 1.44918866383537 | 1.48241759567808E-20 | *SH3BGRL* | 1.19541371352151 | 0.00283400317917603 |
| *RPS13* | 1.23998532673716 | 9.27935396738498E-24 | *NACA* | 1.55968727387601 | 1.55415820248609E-20 | *SNHG8* | 1.4045208216246 | 0.00283810187829936 |
| *HLA-DQA1* | 1.64508396258286 | 2.57650762068709E-23 | *RPL37A* | 1.23209603157097 | 1.81933508183296E-20 | *HADHA* | 0.847529712560942 | 0.00292404687466986 |
| *RPLP1* | 1.14867255816602 | 2.82166424424629E-23 | *ATG101* | 1.01252709787616 | 1.83507588780289E-20 | *EIF3K* | 0.883485991273575 | 0.00294905816162944 |
| *RPL22* | 1.15667163331613 | 3.63291097553206E-23 | *CTSS* | 1.27166810782842 | 1.86077908552899E-20 | *TUBA1B* | 1.22781219804252 | 0.00317088373013558 |
| *CAPG* | 1.09464759028232 | 4.51771561481447E-23 | *FCGRT* | 1.26071780826788 | 2.12925623405662E-20 | *HMGN1* | 1.11239218026262 | 0.00340288903315792 |
| *CANX* | 1.28905487503594 | 4.90642913892177E-23 | *NIBAN3* | 1.06322906875365 | 2.22509257861351E-20 | *RPS3A* | 1.45811802095759 | 0.00365072207768968 |
| *SULF2* | 1.14206747444178 | 5.98175461542903E-23 | *UNC93B1* | 1.15937048794819 | 2.48996211151394E-20 | *RPS3* | 1.41054740517028 | 0.00459766330424437 |
| *SNX3* | 1.25654735619006 | 6.51969494593386E-23 | *MAP1A* | 1.07134519181112 | 2.76465386896406E-20 | *FTH1* | 1.55140994101013 | 0.00638404557330925 |
| *RPL4* | 1.19466897332152 | 1.00372207377457E-22 | *EEF2* | 1.22504325610812 | 3.62462384711742E-20 | *SEPTIN6* | 1.04337141693952 | 0.00650355536472553 |
| *CLEC4C* | 1.03759678665765 | 1.07290621894171E-22 | *AFF3* | 0.92246339952898 | 4.04412699516682E-20 | *RPS9* | 1.23553089079557 | 0.00711109818403852 |

| | | | | | | | | |
|---|---|---|---|---|---|---|---|---|
| *BRI3* | 1.4575440054841 | 1.64176917307794E-22 | *PRXL2A* | 1.02906246560246 | 4.04412699516694E-20 | *RPL36A* | 1.23888594051447 | 0.00751319623038448 |
| *RNASE6* | 1.08325550364935 | 1.73761876674725E-22 | *DPYSL2* | 1.11497636858974 | 5.2398284668785E-20 | *RPL26* | 1.35150078987005 | 0.00816721330419789 |
| *FCGRT* | 1.22717542744926 | 1.77001185463804E-22 | *CLN8* | 1.06740877725195 | 6.00040140440165E-20 | *RPSA* | 1.27528874640174 | 0.00846409656151151 |
| *SNHG5* | 1.31731807981541 | 2.00355051005637E-22 | *IRF2BP2* | 1.23592749408203 | 7.23738126180223E-20 | *NME4* | 0.674581550220066 | 0.00853017570824315 |
| *AFF3* | 1.0697089718114 | 2.80411194082907E-22 | *BCL7A* | 1.0413407953809 | 8.82838912248285E-20 | *BZW2* | 0.643554654599442 | 0.00853681206393322 |
| *UBA52* | 1.10518014101722 | 3.12820370765788E-22 | *CANX* | 1.08096972433992 | 1.04176939545239E-19 | *COX5A* | 0.861824047880451 | 0.00862464957912216 |
| *RPS26* | 0.918060427161245 | 3.7910491852908E-22 | *RNASEK* | 1.10006583693403 | 1.10533829479735E-19 | *SPN* | 0.784114736782715 | 0.00916094838473162 |
| *RPL11* | 1.07035967067218 | 4.24066437523689E-22 | *PLEK* | 1.56707803760342 | 1.13690729550079E-19 | *FTL* | 1.17337560112814 | 0.00936416263686532 |
| *LRRC26* | 1.14699497272723 | 4.50928644457519E-22 | *SNX3* | 1.20047095822011 | 1.28489347287423E-19 | *FCER1A* | 0.869939459435627 | 0.00940556689698747 |
| *RAB11FIP1* | 1.74728458160076 | 5.2922644245981E-22 | *SULF2* | 1.04956077591902 | 1.43035544169727E-19 | *MBOAT7* | 0.46394709975979 | 0.0094055668969875 |
| *TPT1* | 1.04418335447154 | 6.5455334854982E-22 | *PTMS* | 1.26079275151825 | 1.59410908504124E-19 | *HBD* | 1.35147237050138 | 0.00940556689698754 |
| *CORO1C* | 1.09221637122097 | 7.2262820807524E-22 | *CAPG* | 1.06917267897662 | 1.79594187075804E-19 | *CYTL1* | 1.22948184612277 | 0.00940556689698757 |
| *CD2AP* | 1.3538143860552 | 9.20208886012987E-22 | *EIF4A3* | 1.0461359789614 | 2.06377978877469E-19 | *EREG* | 0.756728848987636 | 0.00940556689698757 |
| *NIBAN3* | 1.26727023513969 | 9.35412015830876E-22 | *CTSC* | 1.25301540092653 | 2.13312076520071E-19 | *MYB* | 0.665580960929441 | 0.00940556689698757 |
| *COBLL1* | 1.44661849046863 | 1.05405044280678E-21 | *IGHM* | 1.65761744753906 | 2.34317732490158E-19 | *SNHG19* | 0.534336427651188 | 0.00940556689698757 |
| *TRAF4* | 1.60265205979703 | 1.31695476513114E-21 | *SCN9A* | 0.957439537771954 | 2.79879191393646E-19 | *SYPL1* | 0.749349318622038 | 0.00969237271459255 |
| *RPS16* | 1.03926093294137 | 1.42917374114587E-21 | *MEF2C* | 1.1588829392615 | 3.28529427915347E-19 | *RPS13* | 1.28492048320837 | 0.00979916399182853 |
| *RGS2* | 2.07687437819619 | 1.43992706008609E-21 | *H3F3B* | 1.06822243956745 | 4.72999031026648E-19 | *RPL3* | 1.34960169036779 | 0.00995391193884734 |
| *SLC15A4* | 1.46101786952925 | 1.63379952183342E-21 | *DUSP5* | 1.19376063189588 | 6.43081386266815E-19 | *CLTA* | 0.971741739958487 | 0.0103087208084467 |
| *TM9SF2* | 1.19930231838177 | 1.69585013025879E-21 | *COMMD6* | 1.35324653443019 | 6.48938659731124E-19 | *NDUFA4* | 0.84245872301352 | 0.0103922067626886 |
| *RUNX2* | 1.22311244882148 | 1.82864663456901E-21 | *RACK1* | 1.26563896404082 | 7.06146563427772E-19 | *TUBB* | 1.08843796466758 | 0.0111372212411769 |
| *CNPY3* | 1.25312501788792 | 1.84522731329213E-21 | *RPLP1* | 1.18737933297971 | 9.69972669521264E-19 | *PCBP2* | 0.917948922597783 | 0.0131720662787702 |
| *FAU* | 0.984636181962014 | 2.01186680049396E-21 | *UBC* | 1.22636582823798 | 1.2670952495522E-18 | *PRDX1* | 0.897282747758041 | 0.0138917695061974 |
| *ATP5MG* | 1.15282323142949 | 2.46668757565698E-21 | *ACTB* | 1.02881158393226 | 1.44129808175219E-18 | *RPL18* | 1.20063033850071 | 0.0139196723166541 |
| *SOX4* | 1.88941258487355 | 2.59680266899927E-21 | *CFL1* | 1.17240812604996 | 1.73449253877626E-18 | *SERBP1* | 0.933606611071715 | 0.0145048579364362 |
| *RPL29* | 1.09655539777865 | 2.66493753243919E-21 | *RPS23* | 1.37084625808249 | 1.76082329501061E-18 | *PARP1* | 0.763024255519115 | 0.0152788831988103 |
| *GAS6* | 1.06288770595539 | 2.91424414232196E-21 | *RPL22* | 1.12558395553203 | 2.15326466067582E-18 | *RPL14* | 1.25148538821357 | 0.0155607375286757 |
| *TMBIM6* | 1.20812831985404 | 3.27294956545844E-21 | *TACC1* | 0.845837328475573 | 2.47836174225332E-18 | *RPL10* | 1.29765334876435 | 0.0159613442000173 |
| *RASSF2* | 1.063474637525 | 3.68604777077865E-21 | *CBX4* | 0.989415786450484 | 3.82939833728649E-18 | *RPL17* | 1.33781776284473 | 0.0166690216862241 |
| *JAML* | 1.09464759028232 | 4.12619363477353E-21 | *RPS6KA4* | 0.947866382754482 | 3.95140087909729E-18 | *LAMTOR1* | 0.833180909425509 | 0.017367871659167 |
| *ARPC1B* | 1.31793331351601 | 6.77925967982234E-21 | *SNHG29* | 1.32816985307972 | 4.41055531241933E-18 | *RPL32* | 1.29454005381932 | 0.0180035062991432 |
| *CAT* | 1.17250655767963 | 7.37278654385621E-21 | *LYN* | 1.10574955007619 | 5.06380146629002E-18 | *LYL1* | 1.01238372445583 | 0.018327963990973 |
| *MEF2C* | 1.13128389495312 | 9.32440406430156E-21 | *NGLY1* | 1.00168908226063 | 5.17455832234085E-18 | *RPL37* | 1.30763535451759 | 0.0185221608689477 |
| *MAP1A* | 1.21053314116345 | 9.41583491915403E-21 | *SSR3* | 1.05927707864947 | 5.94906110740535E-18 | *RPS18* | 1.29464320385222 | 0.0191796480943087 |
| *SYK* | 1.10970942027959 | 9.46969415302692E-21 | *SCARB2* | 0.986456294224603 | 6.1793912021874E-18 | *SNRPF* | 0.84245872301352 | 0.0201432043635464 |
| *DAB2* | 1.10997284202513 | 1.14121540782616E-20 | *SIVA1* | 1.12625267441813 | 7.46031729839495E-18 | *XRCC5* | 0.804490872814499 | 0.0205593758904285 |
| *FLNB* | 1.02595269567647 | 1.14121540782616E-20 | *TNFRSF21* | 0.891147859062771 | 7.92846622430387E-18 | *RPS15* | 1.2456541995228 | 0.0239150706679405 |
| *RPL17* | 1.05484258000659 | 1.59757032270699E-20 | *LILRB4* | 1.03954897279095 | 7.96283299208347E-18 | *TPI1* | 1.11191939800675 | 0.0240508519291889 |

| | | | | | | | | |
|---|---|---|---|---|---|---|---|---|
| *RPL27* | 1.07137458104881 | 1.66984078236963E-20 | *OFD1* | 1.27116621356666 | 8.36455760936282E-18 | *HLA-DRA* | 1.98327413582611 | 0.0249461496509083 |
| *SH2B3* | 1.22733964064629 | 1.74482624292184E-20 | *RUNX2* | 1.02236781302845 | 8.60484465693709E-18 | *ITGA4* | 0.787532512509926 | 0.0250018549786163 |
| *NGLY1* | 1.12884640738782 | 1.74687216398636E-20 | *CHCHD2* | 1.270637912358 | 8.81794102826634E-18 | *PTMA* | 1.28763091836451 | 0.02526134766305 |
| *GNG5* | 1.2028754232436 | 1.91217929876583E-20 | *ARPC1B* | 1.25598648559615 | 9.50871557221637E-18 | *NDUFS5* | 0.820432416683521 | 0.0269790314122695 |
| *TACC1* | 1.05221232396192 | 2.02164649902518E-20 | *TUBA1B* | 1.53611139565005 | 1.08695908491932E-17 | *RTN4* | 0.749349318622038 | 0.0273159983460714 |
| *CCDC69* | 1.2054585842576 | 2.43188834173725E-20 | *CORO1C* | 0.940057459543718 | 1.13777155507518E-17 | *HNRNPAB* | 0.818211176766843 | 0.0293333182567933 |
| *SSR1* | 1.16301645816881 | 3.1222456288202E-20 | *MYL12A* | 1.1564358925388 | 1.25829840787994E-17 | *RPL39* | 1.38800404196043 | 0.0309910773578713 |
| *MT-ND5* | 1.13204649763777 | 3.19059049572417E-20 | *RPS13* | 1.27800484018566 | 1.5229307422612E-17 | *SRSF9* | 0.816463514480575 | 0.0310891651338393 |
| *SEC61G* | 1.32601035858338 | 3.22362328442165E-20 | *PHB* | 1.09328959895907 | 1.65303849441082E-17 | *EIF3M* | 0.791343407286666 | 0.0317251793502064 |
| *RHOA* | 1.17214138776288 | 3.55241061316767E-20 | *RPS24* | 1.30847483519544 | 1.68043993067543E-17 | *ESD* | 0.699718550897438 | 0.0319230117854859 |
| *DDX5* | 0.980321748219219 | 3.6806982485885E-20 | *FCHSD2* | 1.00085758434334 | 1.7344128920323E-17 | *ATP5MC3* | 0.887869182703467 | 0.0320142417888681 |
| *FCHSD2* | 1.16315988290134 | 3.7067945611347E-20 | *RPL37* | 1.27505712503016 | 1.77688642478511E-17 | *RPL11* | 1.2375755281691 | 0.0333325896133559 |
| *EEF1G* | 1.10727415142446 | 4.06303337173472E-20 | *SCAF11* | 0.97543852953674 | 1.82012141366629E-17 | *VIM* | 1.64871695551555 | 0.0337335690309339 |
| *ATG101* | 1.07649813747404 | 4.34492996869372E-20 | *RPS3A* | 1.38491784983221 | 2.26568358293015E-17 | *RPS12* | 1.29121092782735 | 0.0351988458533426 |
| *SCARB2* | 1.17934136675669 | 4.40272043377274E-20 | *SYK* | 0.987394746941119 | 2.64459696537908E-17 | *RPL19* | 1.29397244523834 | 0.0352276989402303 |
| *UNC93B1* | 1.12884640738782 | 4.80920539922102E-20 | *TPT1* | 1.03968694556278 | 3.31722569172848E-17 | *NDUFB1* | 0.743249119085621 | 0.0361887946484694 |
| *YBX1* | 1.17200541059938 | 5.13403063641427E-20 | *WDFY4* | 0.86375409223052 | 3.32296460214181E-17 | *RPS15A* | 1.21371453026445 | 0.0395513121757371 |
| *IGHM* | 2.19934546053099 | 5.19388628045312E-20 | *LGMN* | 0.926882070795985 | 4.73157081362175E-17 | *MZT2B* | 0.892582202439356 | 0.0396956317453986 |
| *CLN8* | 1.19218058926959 | 5.97437170308269E-20 | *GAPT* | 0.987394746941119 | 5.2458048506581E-17 | *ALDH1A1* | 0.814444346843923 | 0.0422880476340244 |
| *PTMS* | 1.22665280652672 | 6.14452512711059E-20 | *IDH3A* | 0.838177845524504 | 6.72452843741582E-17 | *LINC02573* | 0.568283759574526 | 0.0422880476340244 |
| *RPL23* | 1.1518058429867 | 6.62274061774779E-20 | *SSR1* | 0.968520923667412 | 7.40461495616637E-17 | *MAP7D3* | 0.46394709975979 | 0.0422880476340244 |
| *RPL21* | 0.980229363537846 | 6.73437667515528E-20 | *DCK* | 0.972957363717313 | 8.02043417756369E-17 | *PTOV1* | 0.427421223734676 | 0.0422880476340244 |
| *CBX4* | 1.13623577758673 | 6.77276760349062E-20 | *HLA-DMB* | 0.918031153213719 | 8.22660754588414E-17 | *RAB13* | 0.46394709975979 | 0.0422880476340244 |
| *HLA-DMB* | 1.04313874033646 | 6.77830165172791E-20 | *RPL4* | 1.17638907183804 | 9.07173108000363E-17 | *NOA1* | 0.46394709975979 | 0.0422880476340246 |
| *ATP6V0B* | 1.25294906873947 | 7.05778799032717E-20 | *RPL27* | 0.974129279778539 | 1.03437217980195E-16 | *LGALS9* | 0.512310121321189 | 0.0439212674317261 |
| *CHCHD2* | 1.29806713981889 | 1.57790123016678E-19 | *RRBP1* | 1.25644737088965 | 1.05363181084311E-16 | *EIF3D* | 0.927946622078802 | 0.0450088144922118 |
| *RPL7A* | 1.08242330655082 | 1.67603312228925E-19 | *RPS5* | 1.21657984566045 | 1.18674543144548E-16 | *RPL35* | 1.1064668419717 | 0.0457855423926204 |
| *EIF4A3* | 1.13574012131244 | 2.63557637662487E-19 | *RGS2* | 1.8745188468725 | 2.11878359045851E-16 | *PEBP1* | 0.716927840929661 | 0.0477909998657806 |
| *RPS4X* | 1.13545790517192 | 2.98871291490784E-19 | *RASSF2* | 0.918031153213719 | 2.56319662167694E-16 | *CHCHD2* | 0.798737345584202 | 0.0493987699420424 |
| *RPL18A* | 1.02426785911465 | 3.7574836192815E-19 | *RHOA* | 1.01650757925678 | 2.6258444360735E-16 | *PRKACB* | 0.886852842371973 | 0.0496847601928101 |
| *BCL7A* | 1.04914764932909 | 3.8007741509962E-19 | *RPS9* | 1.10873283332809 | 2.76768731396442E-16 | | | |
| *RRBP1* | 1.47835739100933 | 3.95561221516553E-19 | *RAB11FIP1* | 1.29654277646745 | 3.60052405954863E-16 | | | |
| *STMN1* | 1.23121749845853 | 4.16168811942609E-19 | *CBFA2T3* | 0.893407366244896 | 3.79235827419767E-16 | | | |
| *OSTC* | 1.15104714977455 | 4.23460520811638E-19 | *GSTP1* | 1.15241535429077 | 3.98154017182408E-16 | | | |
| *RPN2* | 1.15331813556829 | 5.60236690347824E-19 | *YBX1* | 1.15047106116454 | 4.73675180989484E-16 | | | |
| *SUB1* | 1.2537210497249 | 6.44200667668383E-19 | *KRT5* | 0.997901534116002 | 5.33104465613E-16 | | | |
| *TUBA1B* | 1.52262891905127 | 6.859354885664E-19 | *ALDH2* | 0.893407366244896 | 5.33104465613031E-16 | | | |
| *RPL35* | 1.05669649625856 | 7.02345009013731E-19 | *CD164* | 1.11395136318898 | 5.62565392295268E-16 | | | |

| | | | | | |
|---|---|---|---|---|---|
| *EIF4G2* | 1.21446339751881 | 7.16187916536439E-19 | *ATP6V0C* | 0.888884807554202 | 5.85815311757279E-16 |
| *RPS19* | 1.03850340162205 | 8.29102541913441E-19 | *OPN3* | 0.76849209972015 | 7.48087864407387E-16 |
| *ATP6V0C* | 1.050402166835 | 9.42440513749099E-19 | *CUX1* | 0.961910851086641 | 8.96897958763529E-16 |
| *CUX1* | 1.13467448415389 | 1.03103009931294E-18 | *UCP2* | 1.06406294834504 | 1.07501410635025E-15 |
| *ATP5F1A* | 1.09541580609765 | 1.24580354277E-18 | *CCDC69* | 1.02236781302845 | 1.10811552028083E-15 |
| *DCK* | 1.07504841283297 | 1.30254277711797E-18 | *RPL10A* | 1.1759211769432 | 1.26969714746127E-15 |
| *HNRNPA2B1* | 1.2165991418628 | 1.55278352798273E-18 | *DNASE1L3* | 0.84985965057214 | 1.46547171779118E-15 |
| *RPL13* | 0.974959623084271 | 1.63176007528837E-18 | *REPIN1* | 0.902648120514334 | 1.61515413895787E-15 |
| *RPS6KA4* | 0.996192992121733 | 2.11914979970519E-18 | *TRAM1* | 1.10785571209374 | 1.93678925935331E-15 |
| *DUSP5* | 1.20647492458909 | 2.5869391721902E-18 | *CERS6* | 0.947866382754482 | 2.23922747372268E-15 |
| *PTCRA* | 1.28049840122975 | 2.78966929772678E-18 | *SAMHD1* | 1.09270968462667 | 2.4809567240017E-15 |
| *IGFLR1* | 1.16852120972845 | 2.99927809609671E-18 | *GNG5* | 1.06558834162758 | 2.69505509656151E-15 |
| *CD36* | 1.11876979026229 | 3.1000344635475E-18 | *RPS7* | 1.16434220145639 | 3.21112229204925E-15 |
| *WDFY4* | 0.911644125579153 | 3.1000344635475E-18 | *SNHG5* | 1.23707096877625 | 3.25483440316628E-15 |
| *TNFRSF21* | 0.941701359435333 | 3.10003446354759E-18 | *CIB2* | 0.888884807554202 | 3.96728202263914E-15 |
| *ARPC2* | 1.11955262985005 | 3.3229619137365E-18 | *EEF1G* | 1.10240640060062 | 4.01001938840093E-15 |
| *POMP* | 1.04646070820753 | 4.14197664926078E-18 | *LAPTM4A* | 1.01301784484587 | 4.0824861476415E-15 |
| *CALM2* | 1.20470159515013 | 4.20318845076854E-18 | *HIGD1A* | 0.918031153213719 | 4.4073678110217E-15 |
| *NME2* | 1.32623384713291 | 5.14030588361494E-18 | *CALM2* | 1.10019771335185 | 4.41731849178733E-15 |
| *IDH3A* | 0.886109033472016 | 7.04528748334781E-18 | *ATP6V1G1* | 1.06234949851997 | 5.53176742893588E-15 |
| *SCN9A* | 1.04914764932909 | 7.04528748334781E-18 | *COX6A1* | 1.02593266544986 | 5.61492865599525E-15 |
| *CXCR3* | 1.10776519807348 | 7.04528748334801E-18 | *FAU* | 1.01708282272054 | 6.86167844253989E-15 |
| *CSF2RB* | 1.10112192444287 | 7.21200319187417E-18 | *HMGN2* | 1.11655642658103 | 7.82343420255738E-15 |
| *LCP1* | 1.02264786641978 | 8.0152219729698E-18 | *ATP5F1A* | 1.00817143731275 | 8.29127522732132E-15 |
| *PRKCB* | 1.13318313367516 | 8.39997599298245E-18 | *RPS4X* | 1.20054328463717 | 9.80038207320973E-15 |
| *PPIB* | 1.47781311207223 | 8.48814960459646E-18 | *RPL7A* | 1.15124765959325 | 1.03572249516186E-14 |
| *HMGN2* | 1.2528393074125 | 9.80728435107349E-18 | *CD36* | 0.778306389547106 | 1.05819268370219E-14 |
| *SIVA1* | 1.14413481340296 | 1.07286847409593E-17 | *LINC00996* | 0.877515993475005 | 1.05819268370219E-14 |
| *THEMIS2* | 1.01185458634578 | 1.08440979788895E-17 | *ATP6V0B* | 1.12577166917666 | 1.25149619807924E-14 |
| *RPL10A* | 1.0960381712407 | 1.12318420890166E-17 | *SNHG7* | 1.07246791174396 | 1.37638625018682E-14 |
| *YWHAZ* | 1.1001266190975 | 1.21900751641069E-17 | *BLNK* | 0.816909228770853 | 1.4628198621476E-14 |
| *ARID3A* | 1.03171591707634 | 1.63144402656769E-17 | *MYL12B* | 1.09932240960598 | 1.82277648050463E-14 |
| *HINT1* | 1.21287028209532 | 1.64155972606178E-17 | *TMBIM6* | 1.07856301969872 | 1.92338690647954E-14 |
| *OFD1* | 1.4036846944633 | 1.83448793211058E-17 | *POMP* | 0.981357088582548 | 1.93252192393749E-14 |
| *RPS6* | 0.976511128513209 | 2.22626339360431E-17 | *JAML* | 0.887565872486668 | 2.20801559370998E-14 |
| *PIK3CG* | 0.838975057880921 | 2.36823067929322E-17 | *MT-ND4* | 0.865563733319584 | 2.27716074987331E-14 |
| *PRXL2A* | 1.01892086189723 | 2.36823067929322E-17 | *GGA2* | 0.818495479662805 | 2.7046261673732E-14 |
| *KRT5* | 1.12314823077287 | 2.36823067929329E-17 | *SERP1* | 0.897084214768807 | 3.03409624450324E-14 |

| | | | | | |
|---|---|---|---|---|---|
| *CERS6* | 1.05961837226337 | 2.9815309079235E-17 | *TMA7* | 1.00820309329565 | 3.10670141153201E-14 |
| *MYL12A* | 1.1374787247359 | 3.18025180159727E-17 | *RPL41* | 0.835120928942427 | 3.39954805387072E-14 |
| *SNHG29* | 1.25739786437333 | 3.34803559833606E-17 | *HMGN1* | 1.08994079981253 | 3.81233779161674E-14 |
| *SSR3* | 1.10934243124783 | 3.36040290713035E-17 | *RPL35* | 1.04548101229973 | 4.00765234806187E-14 |
| *RPL36A* | 0.961988631248858 | 4.41393341395913E-17 | *C12orf45* | 0.975028713737008 | 4.21194471943833E-14 |
| *RPL5* | 0.976206132984777 | 4.80491417822391E-17 | *SLC25A5* | 1.10133836657678 | 4.76661320388553E-14 |
| *GAPT* | 1.04177956642568 | 5.11360282503165E-17 | *SLC25A6* | 1.08130777864667 | 6.93238013236175E-14 |
| *OPN3* | 0.875767089250556 | 5.24913608259393E-17 | *PFKFB2* | 0.73864217197359 | 9.88004770091499E-14 |
| *ALDH2* | 0.883530489370239 | 5.24913608259409E-17 | *RPL10* | 1.05987026925339 | 1.04956088743004E-13 |
| *LILRB4* | 0.956386831461855 | 5.98391723435678E-17 | *HNRNPA1* | 0.817146031392226 | 1.0586926648798E-13 |
| *GRAMD1B* | 0.966020890908236 | 6.08701610260483E-17 | *PAXX* | 1.01614123866357 | 1.06207220956157E-13 |
| *RPL34* | 0.923349679829126 | 6.23702438187966E-17 | *GNAI2* | 0.949118830997816 | 1.07716907297014E-13 |
| *ZEB2* | 1.28810900964812 | 6.2548029848806E-17 | *ARID3A* | 0.844030571769942 | 1.07990315376919E-13 |
| *HM13* | 0.960688219709953 | 6.76412839524721E-17 | *NUCB2* | 1.03268608788647 | 1.2923405169845E-13 |
| *ATP5F1B* | 1.00426038519028 | 6.84847066687379E-17 | *RPS2* | 1.19987415123359 | 1.39188824789104E-13 |
| *MT-ND3* | 1.0151146357366 | 7.37474873073822E-17 | *RPL22L1* | 1.06945705319602 | 1.40567594295046E-13 |
| *ANKRD11* | 1.58117157415866 | 7.92949146946234E-17 | *RPL23* | 0.87157794229861 | 1.53695599539767E-13 |
| *RBM3* | 1.13365588859126 | 7.9540104742522E-17 | *ATP5MG* | 1.06964427637909 | 1.60373766963921E-13 |
| *GNAI2* | 1.20711857183102 | 1.18149505790546E-16 | *RPN2* | 0.992031734657495 | 1.62890581668498E-13 |
| *LGMN* | 0.901484259358931 | 1.70145095241186E-16 | *UBA52* | 1.01810858232522 | 1.66178772220461E-13 |
| *RPL9* | 1.01949474352439 | 1.75567226092435E-16 | *RBM3* | 0.964111649200059 | 1.72896067693826E-13 |
| *ACTR2* | 1.13643440512955 | 1.77236588476668E-16 | *FTL* | 0.882891817459567 | 1.8131709781271E-13 |
| *ATP5MPL* | 1.05221232396192 | 1.78603590317459E-16 | *SNX9* | 0.888018584147279 | 2.2101837951266E-13 |
| *MYL12B* | 1.09867460402412 | 1.80278000094285E-16 | *GSN* | 0.853361725655992 | 2.40060729124963E-13 |
| *CTSC* | 1.21911913384132 | 1.8074983379059E-16 | *LAMP5* | 0.797736919496007 | 2.51606267601777E-13 |
| *PHB* | 1.10112192444287 | 1.94710725988075E-16 | *NDUFA4* | 1.01153254104489 | 2.60037118084293E-13 |
| *MTDH* | 1.24361316289585 | 1.98306675919634E-16 | *RPS12* | 1.00751583462181 | 3.03130639237802E-13 |
| *YPEL5* | 1.14473670010313 | 2.34034616449647E-16 | *GPR183* | 1.78102748668591 | 3.05462108550222E-13 |
| *RPL26* | 0.927459164413489 | 2.56311033887084E-16 | *GNG7* | 0.728553354350006 | 3.42592593369317E-13 |
| *SPINT2* | 1.01460083408028 | 3.1224563479458E-16 | *MIF4GD* | 0.773407590134266 | 3.42592593369337E-13 |
| *RPS21* | 0.928149057150693 | 3.36651987706731E-16 | *RPL35A* | 0.964577887681249 | 3.44733420547867E-13 |
| *RPS14* | 0.88730419475767 | 3.38360771430852E-16 | *AP1S2* | 0.962055777105717 | 3.89737308814939E-13 |
| *RPL3* | 0.975231671316947 | 3.68793644020358E-16 | *FLNB* | 0.73864217197359 | 4.65814048860995E-13 |
| *SELL* | 1.11975609858192 | 4.00679912290835E-16 | *SPCS3* | 0.94690121166685 | 4.93356213539771E-13 |
| *TRAM1* | 1.2054585842576 | 4.15702776949146E-16 | *GUK1* | 1.04510865116594 | 4.93952600318428E-13 |
| *SAMHD1* | 1.28714542944982 | 4.20610162137444E-16 | *CDKN2D* | 0.974065188128047 | 5.57028798248051E-13 |
| *GRB2* | 1.0869777421226 | 4.22274088945328E-16 | *ATP5MF* | 0.940723739073821 | 5.94310124573239E-13 |
| *COX6B1* | 1.09010292348271 | 4.25089349718914E-16 | *SUB1* | 1.02707391250901 | 6.3105988769921E-13 |

| | | | | | |
|---|---|---|---|---|---|
| *CCDC186* | 1.31976221091821 | 4.47695261032503E-16 | *LAP3* | 0.793640290976529 | 6.42507488073072E-13 |
| *RPL19* | 1.02834731900471 | 4.89055414149024E-16 | *BTG2* | 1.14224631015698 | 6.67011245513296E-13 |
| *AP1S2* | 1.09092615733698 | 5.02237414777281E-16 | *C11orf58* | 0.92283213947754 | 6.90554927311631E-13 |
| *PSMB3* | 0.92176834285286 | 5.25166236475361E-16 | *RPS16* | 0.919354119873037 | 9.11789726520769E-13 |
| *SNHG7* | 1.15445898178858 | 5.41200261616765E-16 | *RPL21* | 0.906387699355434 | 9.46619941721151E-13 |
| *TMED2* | 0.981251451938625 | 6.08235973459718E-16 | *COX6B1* | 1.04408465622012 | 9.73171015845631E-13 |
| *P4HB* | 1.11369059794592 | 6.19091184675287E-16 | *RPS19* | 1.09752876562828 | 1.00783333988229E-12 |
| *MT-ATP6* | 0.965628372035907 | 9.3834878177368E-16 | *ZEB2* | 0.749744647167161 | 1.01211209425254E-12 |
| *ELMSAN1* | 0.972501758435766 | 1.04103332047363E-15 | *P4HB* | 0.944831212557434 | 1.05244923826358E-12 |
| *RPL28* | 0.787642702203915 | 1.14451123743778E-15 | *RPL29* | 1.16056656340925 | 1.07917723371597E-12 |
| *GSTP1* | 1.19372467561022 | 1.15553615188434E-15 | *PIK3CG* | 0.703018262242869 | 1.56982464261279E-12 |
| *TMA7* | 1.00310984913356 | 1.16075610625255E-15 | *SELL* | 0.929526792051548 | 1.68367429900881E-12 |
| *RPL41* | 0.714365067202669 | 1.21874914294549E-15 | *DSTN* | 1.13159688278321 | 1.91269872229263E-12 |
| *COMMD6* | 1.27600542020537 | 1.44703953124963E-15 | *PNISR* | 1.0199882183594 | 1.96377851756298E-12 |
| *GNB1* | 0.889983706910484 | 1.6292898724541E-15 | *TMEM258* | 0.997551492857246 | 2.06805106928907E-12 |
| *CD302* | 0.870568182576237 | 1.68098315696008E-15 | *PACSIN1* | 0.703018262242869 | 2.11965352112838E-12 |
| *PTBP3* | 1.12969107663964 | 1.7277750206086E-15 | *EPHA2* | 0.700439718141092 | 2.11965352112851E-12 |
| *RPL12* | 0.930527548847158 | 1.76218099204714E-15 | *GPM6B* | 0.821662758741488 | 2.11965352112851E-12 |
| *HNRNPK* | 1.07104058520501 | 1.79277278321624E-15 | *PABPC1* | 1.02587840992713 | 2.28933809511334E-12 |
| *USF2* | 0.997013521102127 | 1.81808562932778E-15 | *LCP1* | 0.645631329156247 | 2.31402846237733E-12 |
| *RPL39* | 1.00354543644633 | 1.96144361687531E-15 | *ANKRD11* | 1.09461988493704 | 2.6471139865834E-12 |
| *CD164* | 1.22364405665439 | 2.23830772547529E-15 | *USF2* | 0.888512066293664 | 2.71267695913323E-12 |
| *PRKCD* | 0.784732013096934 | 2.4435873107582E-15 | *BRK1* | 0.962978559267693 | 2.89069390743192E-12 |
| *COX4I1* | 1.02119838014592 | 3.01293725261448E-15 | *ATP5MPL* | 0.955675927598851 | 2.90925524275073E-12 |
| *HIGD1A* | 0.932522320467121 | 3.24922431305585E-15 | *STX7* | 0.615731865075289 | 3.05838019821399E-12 |
| *LINC02812* | 0.844288857777388 | 3.54463840954028E-15 | *GRAMD1B* | 0.848625145847192 | 3.42024511324655E-12 |
| *REPIN1* | 0.924624269077068 | 3.6731606963546E-15 | *B4GALT1* | 0.83145981224016 | 4.49259658466994E-12 |
| *C12orf45* | 1.00772538798936 | 3.87079340135967E-15 | *RPS21* | 0.962084572360201 | 4.49662103726667E-12 |
| *SFPQ* | 1.13021483596319 | 4.46225508296344E-15 | *PTBP3* | 0.944831212557434 | 4.53642621127815E-12 |
| *SPCS3* | 1.13599696572133 | 5.05122052237131E-15 | *SELENOH* | 0.918031153213719 | 5.42369451468598E-12 |
| *PFKFB2* | 0.825604294512462 | 5.13105151658045E-15 | *EEF1B2* | 1.05206770012932 | 5.42780812072654E-12 |
| *GGA2* | 0.887084552503632 | 5.40632977739786E-15 | *RPL3* | 1.07654132204451 | 5.65270386665405E-12 |
| *SCAF11* | 1.13081096417182 | 5.75699513220914E-15 | *THEMIS2* | 0.850524333855442 | 6.03949734152448E-12 |
| *CBFA2T3* | 0.838975057880921 | 7.41210842351361E-15 | *CSF2RB* | 0.850714885075156 | 6.14897627599254E-12 |
| *RAC1* | 1.0713062700073 | 8.47742455868215E-15 | *OST4* | 0.960250981264761 | 6.41330360751841E-12 |
| *BAG1* | 0.983066162921154 | 9.3730574908278E-15 | *SCAMP5* | 0.74153349784792 | 7.13446939464632E-12 |
| *ST14* | 0.770846486841984 | 1.06853442579938E-14 | *SUMO2* | 0.934389841813387 | 7.35295171454541E-12 |
| *RPS5* | 1.07069089923195 | 1.08369914790128E-14 | *BAG1* | 0.869832064956387 | 7.49460434877848E-12 |

| | | | | | |
|---|---|---|---|---|---|
| *QKI* | 1.06644447281012 | 1.12220431765846E-14 | *RPL9* | 0.9851568739906 | 9.1045576270366E-12 |
| *SLC25A5* | 1.02956866341783 | 1.12873372723359E-14 | *RASD1* | 1.1717877454595 | 9.32477003660953E-12 |
| *RPL18* | 0.867486807799165 | 1.13286938242623E-14 | *MYBL2* | 0.819287951553584 | 9.3247700366098E-12 |
| *LAPTM4A* | 1.03972137977872 | 1.28476202773098E-14 | *SHTN1* | 0.775859069128363 | 9.3247700366098E-12 |
| *HMGA1* | 0.88803032159841 | 1.50677402761004E-14 | *DBNL* | 0.754365142243089 | 1.00076509901358E-11 |
| *EPHA2* | 0.825604294512462 | 1.537293811379E-14 | *RPL24* | 0.896798516135632 | 1.04896467521452E-11 |
| *STX7* | 0.844397840494216 | 1.57532544344266E-14 | *LINC02812* | 0.761087318498399 | 1.24914103056732E-11 |
| *B4GALT1* | 1.08357649487125 | 2.44701918882556E-14 | *HMGA1* | 0.787401710561236 | 1.35586089132704E-11 |
| *BTG2* | 1.29552142695057 | 2.76192539702187E-14 | *CD37* | 0.929376126807728 | 1.39608332091205E-11 |
| *RPL35A* | 0.8638866954764 | 2.81496823660367E-14 | *NHP2* | 0.857866926833746 | 1.42340824155067E-11 |
| *NIN* | 0.951313044410728 | 2.92764803594803E-14 | *RPL5* | 1.01065983551855 | 1.42486759647787E-11 |
| *RPS12* | 0.937928896466028 | 2.9800022131793E-14 | *ELMSAN1* | 0.780103244980392 | 1.4323100901424E-11 |
| *RPN1* | 0.748820180511985 | 3.05717305052958E-14 | *RBIS* | 0.908115684348636 | 1.47568663473019E-11 |
| *SLC25A6* | 1.09364034307341 | 3.50345339329193E-14 | *RPS27* | 0.779463324843142 | 1.51164149436408E-11 |
| *RPLP2* | 0.685940651678339 | 3.66963346487246E-14 | *RPL11* | 1.10467791326522 | 1.59515115237426E-11 |
| *IFNAR2* | 0.889617819249154 | 3.73162285489653E-14 | *LIME1* | 0.988420481105117 | 1.63798784113361E-11 |
| *GPR183* | 2.0443098322229 | 3.7439779854393E-14 | *SH2B3* | 0.962675068268561 | 1.65640442657367E-11 |
| *CD37* | 1.04827948633999 | 3.90342740374201E-14 | *RPL19* | 1.12875505616907 | 1.76863565309892E-11 |
| *NDUFA4* | 1.01185458634578 | 3.92681134049365E-14 | *ZNF706* | 0.895504584197111 | 1.88147281889521E-11 |
| *NDUFB1* | 0.928683715389024 | 3.93782284572185E-14 | *TMED2* | 0.829593188546135 | 1.94107484214892E-11 |
| *ARF1* | 1.00158625089195 | 3.94252249826635E-14 | *PTMA* | 0.653730670035933 | 2.04810928808684E-11 |
| *HNRNPD* | 0.865633845494735 | 3.98080445549627E-14 | *A1BG* | 0.845099630759868 | 2.12058643636797E-11 |
| *SLC7A5* | 0.958657345353413 | 4.33957713001723E-14 | *ST14* | 0.748660927798581 | 2.23305161771717E-11 |
| *MED13L* | 1.0496256893992 | 4.50910477643167E-14 | *TMEM109* | 0.841919825174135 | 2.62774039198118E-11 |
| *PECAM1* | 0.759641032134103 | 4.52367519651899E-14 | *ATP5MD* | 0.870202628123103 | 2.73859090140545E-11 |
| *CYP46A1* | 0.931751699027728 | 4.52367519651912E-14 | *HSP90AB1* | 0.932482309511259 | 2.81976610376874E-11 |
| *MIF4GD* | 0.75400549746467 | 4.52367519651912E-14 | *MTDH* | 1.03794061694892 | 3.19676529858276E-11 |
| *ZNF706* | 0.969296712935446 | 4.62128829636183E-14 | *RPL26* | 0.970686333567389 | 3.83443162246205E-11 |
| *PCBP2* | 0.977973693670001 | 4.79866531571369E-14 | *EPHB1* | 0.669132268090459 | 3.97198242835721E-11 |
| *HMGB1* | 0.980364124596529 | 5.42276968909791E-14 | *CD302* | 0.645197607854526 | 3.97198242835733E-11 |
| *NADK* | 0.854122317062069 | 5.53849164975927E-14 | *RPL36A* | 0.925325880522628 | 4.21580039825004E-11 |
| *RBM39* | 0.829512782871063 | 5.69094659210469E-14 | *ATP13A2* | 0.7390610121684 | 4.44695655802215E-11 |
| *NCKAP1L* | 0.933449374731932 | 5.81290205760947E-14 | *COX8A* | 0.886780219229279 | 4.4613241110086E-11 |
| *COX8A* | 1.00526190903535 | 6.40378437940286E-14 | *RPL18A* | 1.06934473957664 | 5.12164977832031E-11 |
| *TPP1* | 0.89225649404214 | 6.65714505418325E-14 | *ERCC1* | 0.838517783094618 | 5.61599008855929E-11 |
| *SLC20A1* | 1.04437350722085 | 6.81857597899221E-14 | *HMGB1* | 0.830124064064291 | 6.12379698204557E-11 |
| *SNX9* | 0.949118830997816 | 7.71499105497751E-14 | *OGT* | 0.782803772519496 | 7.48477574761618E-11 |
| *PAXX* | 0.955043664954921 | 8.50839183469313E-14 | *TMEM59* | 0.87485032944984 | 8.24676152303022E-11 |

| | | | | | |
|---|---|---|---|---|---|
| *SUMO3* | 0.876323005399323 | 8.78593453617122E-14 | *NEDD8* | 0.829138103623307 | 8.82835808811237E-11 |
| *COPE* | 0.956823495722851 | 8.9459062803978E-14 | *MT-ND2* | 0.878366935640203 | 9.02996996179639E-11 |
| *RPL24* | 0.85528244496289 | 9.99021962521997E-14 | *RPS26* | 0.927392581423257 | 1.02872590839971E-10 |
| *CDKN2D* | 1.09252590075448 | 1.04436882697956E-13 | *EIF4G2* | 0.877230622088279 | 1.06020475362607E-10 |
| *EIF1* | 0.789870962935528 | 1.06967966182009E-13 | *PRKCB* | 0.975238010620195 | 1.11695223670908E-10 |
| *FERMT3* | 0.91628224898927 | 1.13670195661467E-13 | *M6PR* | 0.882453482967653 | 1.12546406948036E-10 |
| *PTMA* | 0.69085738187614 | 1.17427480039434E-13 | *ZFAT* | 0.68099195591287 | 1.157805612681E-10 |
| *EPHB1* | 0.790249018064499 | 1.30819627435462E-13 | *MGAT1* | 0.730227338340822 | 1.35597480778073E-10 |
| *RPL10* | 0.937094025296919 | 1.394833886035E-13 | *SEC31A* | 0.749431549237974 | 1.35967637127384E-10 |
| *XBP1* | 0.872896325441829 | 1.4669320654922E-13 | *PSMB3* | 0.825734305892937 | 1.42332710440772E-10 |
| *MOB1A* | 1.0296785579213 | 1.58832532501295E-13 | *RPL13* | 1.00288584304339 | 1.47032994076727E-10 |
| *RPL22L1* | 1.00570235936418 | 1.61528876457592E-13 | *EIF3L* | 0.841468816067144 | 1.62155938636947E-10 |
| *MGAT1* | 0.844015830611746 | 1.7835837866046E-13 | *CDK2AP2* | 0.802883280628198 | 1.62626319255549E-10 |
| *CIB2* | 0.756826016527049 | 1.85680038543344E-13 | *FABP5* | 0.626303296102056 | 1.64060713398283E-10 |
| *RPL32* | 0.814380558967458 | 1.8797504004082E-13 | *COX4I1* | 0.958073089854439 | 1.85002794196183E-10 |
| *ATP6V1G1* | 1.01759856888058 | 1.9610189470961E-13 | *RPS6* | 0.950977492680309 | 1.85209234624102E-10 |
| *HNRNPU* | 1.10674647363674 | 2.09886993873393E-13 | *IFNAR2* | 0.74400175343867 | 1.91210353033629E-10 |
| *RPS27A* | 0.83092868724823 | 2.21147331003602E-13 | *HM13* | 0.793640290976529 | 2.05839018231709E-10 |
| *BRK1* | 0.963253115201452 | 2.22840816572096E-13 | *EIF2AK4* | 0.697432013700322 | 2.06784521038999E-10 |
| *CCNI* | 0.911182182300611 | 2.39844677439932E-13 | *NCL* | 0.797529202982671 | 2.1581797990262E-10 |
| *TMEM59* | 0.901059374937987 | 2.48406561987682E-13 | *ATP5F1B* | 0.835128071551049 | 2.247946858997E-10 |
| *MYBL2* | 0.988047197238235 | 2.63060214686428E-13 | *DBI* | 0.820211952251826 | 2.35964077142409E-10 |
| *OGT* | 1.06848471665051 | 2.74050633860929E-13 | *HNRNPA2B1* | 0.804627749934897 | 2.44732655664047E-10 |
| *GTF2I* | 0.90618503499248 | 2.77509597725002E-13 | *TMED10* | 0.849734541519389 | 2.46789501933344E-10 |
| *CHD9* | 0.998236597562507 | 2.80995769161632E-13 | *NADK* | 0.725386075271323 | 2.48907545935742E-10 |
| *DSTN* | 1.27212719178431 | 3.08425698058895E-13 | *MT-CO2* | 0.830505621071727 | 2.53901396251161E-10 |
| *SLC25A3* | 1.05563915464886 | 3.08746934963571E-13 | *ARPC2* | 0.928299488667546 | 2.63650453943676E-10 |
| *PFN1* | 0.80321705876202 | 3.51412129836673E-13 | *SPCS2* | 0.829494478612061 | 2.82103346344846E-10 |
| *HNRNPM* | 0.988647195184453 | 3.60416744962673E-13 | *MLF2* | 0.761374200762452 | 2.93943857672565E-10 |
| *H2AFY* | 1.00029755492338 | 3.63676682038705E-13 | *CHD9* | 0.862646249937289 | 2.97184470224705E-10 |
| *RNF149* | 0.945105768491192 | 4.00593248008343E-13 | *RAC1* | 0.908160909189658 | 3.16286859969097E-10 |
| *HNRNPA3* | 0.918503463759732 | 4.00741183651862E-13 | *COX7A2* | 0.818963811119892 | 3.31498632247731E-10 |
| *SERP1* | 0.914036371842371 | 4.46506739710005E-13 | *SLC7A5* | 0.74433298866935 | 3.90945529278755E-10 |
| *BLNK* | 0.768053271950303 | 5.25125573732193E-13 | *FERMT3* | 0.87728481087041 | 4.50541622557393E-10 |
| *RBIS* | 0.931443938674625 | 5.28304291863604E-13 | *COMT* | 0.730945600613224 | 5.68229426472446E-10 |
| *ARHGAP27* | 0.827556851116903 | 5.56472895857992E-13 | *SERTAD2* | 0.692405196625489 | 5.77395458411782E-10 |
| *TMEM258* | 1.01696147677887 | 6.73810725944041E-13 | *RPS27A* | 0.894519446453 | 5.94899203328291E-10 |
| *DBNL* | 0.850578462792269 | 7.7217572099957E-13 | *GNB1* | 0.728858267178128 | 6.12081567853305E-10 |

| | | | | | |
|---|---|---|---|---|---|
| *UBB* | 0.856730430026524 | 7.8632404521781E-13 | *DOCK2* | 0.634881386082849 | 6.63373182624112E-10 |
| *PIM3* | 1.02613488481304 | 8.06449913116709E-13 | *SFT2D1* | 0.700933031546784 | 6.75659192307932E-10 |
| *RPS29* | 0.687422914942412 | 8.59811888202176E-13 | *NDUFB1* | 0.881036945731517 | 6.94461674260554E-10 |
| *FKBP1A* | 0.87905006638757 | 8.86850630068383E-13 | *PFN1* | 0.678490450046862 | 7.65412521373567E-10 |
| *HDGF* | 0.821014005250131 | 9.78460140208732E-13 | *RPL39* | 1.00199831368387 | 7.82763395764999E-10 |
| *LAP3* | 0.923334805227583 | 1.00787537818547E-12 | *COX5A* | 0.862383123663805 | 8.13020531606593E-10 |
| *SERTAD2* | 0.839809111940921 | 1.02055152148051E-12 | *LSP1* | 0.918031153213719 | 8.94199973793458E-10 |
| *SON* | 0.938097424906564 | 1.0580400593463E-12 | *XBP1* | 0.705727549500855 | 1.03365523972121E-09 |
| *MT-CYB* | 0.872526405892807 | 1.08901065546318E-12 | *RPL31* | 0.595437455066159 | 1.04449986535395E-09 |
| *MDM4* | 0.949118830997816 | 1.32298640241177E-12 | *SMIM3* | 0.74400175343867 | 1.0514219475657E-09 |
| *B2M* | 0.630700616298009 | 1.43654215962429E-12 | *EID1* | 0.827783834297185 | 1.15283313842144E-09 |
| *DDX17* | 1.04029673174682 | 1.47127582414752E-12 | *BCAP31* | 0.704062306462276 | 1.18202601354544E-09 |
| *SEPTIN2* | 0.917957899511356 | 1.55105142659091E-12 | *MED13L* | 0.784643737936016 | 1.20838463753576E-09 |
| *C11orf58* | 1.03217457072537 | 1.55180869257708E-12 | *PYCARD* | 0.809290523021336 | 1.24226614531409E-09 |
| *DOCK2* | 0.808941172949556 | 1.68170581749082E-12 | *S100A6* | 0.960022178307946 | 1.36159274219683E-09 |
| *MIS18BP1* | 0.981251451938625 | 1.91356121873357E-12 | *CDYL* | 0.77242583096682 | 1.36269573061602E-09 |
| *NUCB2* | 1.14670295875176 | 1.97995525449442E-12 | *ABHD15* | 0.716615865643591 | 1.4433334710071E-09 |
| *PACSIN1* | 0.748347862602744 | 2.04847209222234E-12 | *PPM1J* | 0.587755715612837 | 1.49366850956803E-09 |
| *GSN* | 0.827774010000565 | 2.07608764466038E-12 | *SCAMP4* | 0.601641241867787 | 1.49366850956807E-09 |
| *SMIM3* | 0.838087518609072 | 2.15845874668398E-12 | *SLC35E1* | 0.68099195591287 | 1.53822905956526E-09 |
| *PNISR* | 1.10792175654821 | 2.33171295721015E-12 | *RPL34* | 0.927677290551262 | 1.75451547189799E-09 |
| *MAP3K8* | 0.941372798747175 | 2.65824653448145E-12 | *MDFIC* | 0.804502207583305 | 2.05365188936033E-09 |
| *UQCRQ* | 0.860433677515681 | 2.7452086187582E-12 | *REEP5* | 0.747321865238969 | 2.40558496206693E-09 |
| *LINC00996* | 0.792999629080534 | 2.86644117837113E-12 | *DHRS7* | 0.758374487225234 | 2.56116666484792E-09 |
| *MARCH1* | 0.739819591221359 | 2.86644117837113E-12 | *MGST2* | 0.562419932070068 | 2.56579130040841E-09 |
| *SELENOF* | 0.86765473036892 | 2.88250426436931E-12 | *RPL17* | 1.01819852569068 | 2.70337644332562E-09 |
| *USP15* | 0.995131228754652 | 2.93137405242173E-12 | *SUMO3* | 0.830954696389442 | 2.78034468966495E-09 |
| *TM9SF3* | 0.968227653945521 | 3.07827015607929E-12 | *CCNI* | 0.78693704953165 | 2.85027633156503E-09 |
| *SKAP2* | 0.877495903167308 | 3.18844699182505E-12 | *SNRNP25* | 0.734585012117788 | 2.8590417825524E-09 |
| *HNRNPA1* | 0.889140684957222 | 3.62437876995468E-12 | *C17orf49* | 0.693451172639507 | 2.9393985791962E-09 |
| *REEP5* | 0.810220725176128 | 3.7753787948659E-12 | *CYTH4* | 0.76834937167879 | 3.23903835645163E-09 |
| *SELENOH* | 0.920206139082577 | 3.78350356518655E-12 | *ITGAE* | 0.744966444240839 | 3.26128680435457E-09 |
| *SEC63* | 0.819835814052849 | 3.98910550412181E-12 | *IRF2BPL* | 0.696367181799786 | 3.35400459660671E-09 |
| *GNG7* | 0.678666953601539 | 4.00424397019535E-12 | *MEF2D* | 0.743590670648913 | 3.39479226955151E-09 |
| *A1BG* | 0.877374223494407 | 4.56878822759426E-12 | *ATP5MC3* | 0.840494552742699 | 3.40681774138607E-09 |
| *PGK1* | 0.929191787110176 | 4.68925228009177E-12 | *KTN1* | 0.908591457327413 | 3.44951878148212E-09 |
| *VPS13C* | 0.925776647216301 | 4.92197182594329E-12 | *MXD4* | 0.798540873287159 | 3.67187310332374E-09 |
| *ATP5MD* | 0.883165757971282 | 5.19052471327536E-12 | *SLC20A1* | 0.761440790273017 | 3.76885718412036E-09 |

| | | | | | |
|---|---|---|---|---|---|
| *ALCAM* | 0.642522389866444 | 5.58432046396034E-12 | *H2AFY* | 0.815271578788518 | 4.72877192836954E-09 |
| *EIF2AK4* | 0.773800570785285 | 5.87295302368344E-12 | *CLIC3* | 0.732307441924668 | 4.80303546746583E-09 |
| *HSP90AB1* | 1.15248234636142 | 6.24784493761428E-12 | *RBBP4* | 0.752108082328903 | 4.8668991088208E-09 |
| *MAP2K3* | 0.891668558813916 | 6.8262147437178E-12 | *COPE* | 0.826543145041101 | 5.05349416003761E-09 |
| *TAPBP* | 0.955155220728485 | 7.04446104102743E-12 | *JTB* | 0.876210977519092 | 5.27860399391602E-09 |
| *MXD4* | 0.932583463271516 | 7.06703181599243E-12 | *SAP18* | 0.738774450435232 | 5.44333957547452E-09 |
| *M6PR* | 0.873169977764518 | 7.65190858747831E-12 | *SLC3A2* | 0.675830250797091 | 5.60454134829314E-09 |
| *HNRNPC* | 0.986186157598206 | 8.16849712020813E-12 | *BST2* | 0.891396026710252 | 5.68413374060127E-09 |
| *EIF3L* | 0.828981207684175 | 9.99304915842796E-12 | *NME4* | 0.617798129425676 | 6.29264267623939E-09 |
| *PHEX* | 0.699340521866289 | 1.08073060112512E-11 | *TIMM13* | 0.754365142243089 | 6.30470478238414E-09 |
| *ACTG1* | 1.17357077451256 | 1.21490409316105E-11 | *KIF5B* | 0.842177398694465 | 6.31812068082953E-09 |
| *CHMP4B* | 0.773957181189357 | 1.45158698509034E-11 | *SEPTIN9* | 0.744966444240839 | 6.48173222686354E-09 |
| *RASD1* | 1.23255944749345 | 1.49978765581516E-11 | *SLC25A3* | 0.979708049768543 | 6.52615469755259E-09 |
| *SCAMP4* | 0.690516671479283 | 1.49978765581516E-11 | *TM9SF3* | 0.757453324030659 | 7.05314599784927E-09 |
| *EEF1B2* | 0.942756328774276 | 1.77439924934243E-11 | *MYDGF* | 0.790795964917611 | 7.08418438396394E-09 |
| *GUK1* | 1.03087543057375 | 1.98169846803679E-11 | *SELENOF* | 0.756567730519603 | 7.37090667737407E-09 |
| *COX5A* | 0.90289581423717 | 2.06428707087875E-11 | *RBM39* | 0.463267375619264 | 7.50808020877657E-09 |
| *SRGN* | 1.26012295749231 | 2.09262482906717E-11 | *QKI* | 0.73565441072515 | 7.62397227712331E-09 |
| *ATP5MF* | 0.893317039329464 | 2.20835891507376E-11 | *PPM1K* | 0.819850759274815 | 7.7831003946176E-09 |
| *CREB3L2* | 0.830540709555724 | 2.21821480843875E-11 | *NDUFA11* | 0.753317008019115 | 7.86340686615245E-09 |
| *ZMYM2* | 0.871363673483371 | 2.40177773595888E-11 | *UBB* | 0.745226741046397 | 7.92581564525936E-09 |
| *UCP2* | 1.03206490594322 | 2.63255637457848E-11 | *RPS14* | 0.945269308895831 | 7.96724542778026E-09 |
| *SHTN1* | 0.812108452028239 | 2.8745420912293E-11 | *PHB2* | 0.596103058326357 | 8.0078359367566E-09 |
| *SLC9A7* | 0.713928058089335 | 2.87454209122938E-11 | *ARHGAP27* | 0.625850401720409 | 8.32160783209766E-09 |
| *DAZAP2* | 0.901290305907201 | 3.04068171426979E-11 | *ACTG1* | 1.03106481851361 | 8.71434302554025E-09 |
| *TTC3* | 1.01546132582651 | 3.09334605087664E-11 | *KRTCAP2* | 0.817296730079179 | 9.04926074545025E-09 |
| *ZNF207* | 0.900399823692507 | 3.21175165286411E-11 | *TPP1* | 0.664998139537424 | 9.68907534665029E-09 |
| *RPS27* | 0.740383155044217 | 3.22211920150217E-11 | *WAKMAR2* | 0.54240203849311 | 9.73923451803984E-09 |
| *CCNL1* | 0.963403057004064 | 3.28604986820159E-11 | *GABARAPL2* | 0.761858418019713 | 1.01803404175774E-08 |
| *MLF2* | 0.787013334014803 | 3.58609771685213E-11 | *KDELR1* | 0.664998139537424 | 1.04328326622034E-08 |
| *CYTH4* | 0.877683692954867 | 3.67202596646655E-11 | *PAIP1* | 0.636517326772265 | 1.07073650686697E-08 |
| *MAP4K4* | 0.852257291745227 | 3.94960257811142E-11 | *MT-CO3* | 0.880607898198616 | 1.08392927188773E-08 |
| *LAPTM5* | 0.930135727630243 | 3.96226528516781E-11 | *SKAP2* | 0.681388246271326 | 1.08609753984346E-08 |
| *TMEM170B* | 0.654671472532561 | 3.97017461911017E-11 | *DDX17* | 0.774825008005828 | 1.13190242113277E-08 |
| *SH3TC1* | 0.642522389866444 | 3.9701746191104E-11 | *DDT* | 0.53834476513195 | 1.23986075643522E-08 |
| *PAIP1* | 0.706343264147822 | 4.03882068529834E-11 | *CYP46A1* | 0.653220029124717 | 1.26772889375545E-08 |
| *BST2* | 0.939311213071232 | 4.11147518189705E-11 | *GDI2* | 0.697262474356935 | 1.35699823142171E-08 |
| *RPS18* | 0.680203222165424 | 4.26106013209252E-11 | *NDUFS6* | 0.693990878995789 | 1.37035330426011E-08 |

| | | | | | |
|---|---|---|---|---|---|
| *SRP72* | 0.859759310859172 | 4.34896900201487E-11 | *CBX6* | 0.738677571623713 | 1.46187442030794E-08 |
| *COX6A1* | 0.967756371990366 | 4.35201873323892E-11 | *ARF1* | 0.836567052584823 | 1.52186373632638E-08 |
| *ZDHHC17* | 0.771215772131979 | 4.40199070460021E-11 | *UBL5* | 0.75990845338877 | 1.61620208086748E-08 |
| *NDUFS6* | 0.756787416507224 | 4.99237638969328E-11 | *MARCH1* | 0.573735245297902 | 1.64848112899581E-08 |
| *TRIM8* | 0.82233832125602 | 5.04806618582072E-11 | *PHEX* | 0.607158246835352 | 1.64848112899586E-08 |
| *SYNGR2* | 0.781843407969453 | 5.74627953422184E-11 | *NUDT1* | 0.648191569474811 | 1.69587905681265E-08 |
| *GDI2* | 0.762033278397321 | 6.71654290188602E-11 | *ODC1* | 0.747590811511993 | 1.71165691385085E-08 |
| *ERCC1* | 0.812776430079885 | 7.00971879156356E-11 | *NDUFB8* | 0.796971766976937 | 1.7863087826289E-08 |
| *EIF3F* | 0.839646346465634 | 7.11732905092026E-11 | *PNN* | 0.724363733165788 | 1.9040260585862E-08 |
| *ZNRF2* | 0.693244216736888 | 7.43613666705636E-11 | *APEX1* | 0.74200078912222 | 1.96118415782567E-08 |
| *SLC39A6* | 0.651643779834993 | 7.53823371933324E-11 | *LAIR1* | 0.612404870655457 | 1.97280993055391E-08 |
| *XRCC5* | 0.827556851116902 | 9.97114677795104E-11 | *B2M* | 0.596407135688825 | 2.0127470402663E-08 |
| *JTB* | 0.920655366874122 | 1.0340913091425E-10 | *HNRNPD* | 0.621943063261616 | 2.10182842301365E-08 |
| *GNA12* | 0.608575057943107 | 1.03634035509758E-10 | *NCKAP1L* | 0.690620657110646 | 2.13641926480867E-08 |
| *PPM1J* | 0.633342963092049 | 1.03634035509767E-10 | *PECAM1* | 0.530837286860188 | 2.14141776382913E-08 |
| *S100A6* | 1.43585024500115 | 1.0366221338908E-10 | *TTC3* | 0.890615695633319 | 2.23011176809397E-08 |
| *ABHD15* | 0.770973322750535 | 1.05057131691745E-10 | *MTPN* | 0.741953924789811 | 2.26205242569729E-08 |
| *SCAMP5* | 0.723484396969216 | 1.07731100841362E-10 | *TMEM14C* | 0.658387336580699 | 2.2629208418467E-08 |
| *LAIR1* | 0.703466503392259 | 1.11701679717805E-10 | *NIN* | 0.728206594333702 | 2.37082177049186E-08 |
| *RBX1* | 0.797341176265763 | 1.1258336235372E-10 | *MIS18BP1* | 0.722040047391332 | 2.66990117779772E-08 |
| *ATP5ME* | 0.870616857319098 | 1.15430899649953E-10 | *CDC40* | 0.663847664962191 | 2.69413311838615E-08 |
| *TMSB10* | 0.730097194000608 | 1.15722633129113E-10 | *SET* | 0.774078192482138 | 2.73981826253366E-08 |
| *ANKRD12* | 1.16385767979083 | 1.22893771905045E-10 | *FCER1A* | 0.671767328258429 | 2.77896076769303E-08 |
| *FABP5* | 0.660707862263229 | 1.42258899127201E-10 | *RPL18* | 0.888942424321353 | 2.85741070144859E-08 |
| *RUBCN* | 0.781805043012042 | 1.46153150176641E-10 | *DAD1* | 0.731555842511529 | 2.88194284524657E-08 |
| *TLN1* | 0.908920002713719 | 1.49255732830129E-10 | *YWHAZ* | 0.817073404593606 | 2.89491425502391E-08 |
| *MTRNR2L1* | 1.4796335476966 | 1.51660788410052E-10 | *SLC39A6* | 0.562419932070068 | 3.60272197082883E-08 |
| *MAP3K2* | 0.924094807315339 | 1.53878624662354E-10 | *TLR7* | 0.545278799248674 | 3.60272197082904E-08 |
| *MOB1B* | 0.754763790734946 | 1.58556068530889E-10 | *SYNGR2* | 0.671031545832922 | 3.60912352513756E-08 |
| *TPM3* | 0.887153077691423 | 1.58916864390532E-10 | *LAMTOR1* | 0.734867128282208 | 3.69919450694736E-08 |
| *LAMTOR1* | 0.851343615311339 | 1.61950169926188E-10 | *UQCRQ* | 0.74038823861057 | 3.72873900390801E-08 |
| *HMGN1* | 1.00613877925434 | 1.70824562694857E-10 | *EEF1D* | 0.773483864687716 | 3.72902744558587E-08 |
| *RBBP4* | 0.86770399937517 | 1.7216108292448E-10 | *MDH2* | 0.734517087023452 | 4.06468206043276E-08 |
| *N4BP2* | 0.861799107156541 | 1.77285270977354E-10 | *TMEM14B* | 0.780912296854648 | 4.21789305507439E-08 |
| *PPM1K* | 0.930010008050111 | 1.78733886282419E-10 | *RBX1* | 0.712282487997143 | 4.27660792236141E-08 |
| *NHP2* | 0.858265400546702 | 1.79592390225857E-10 | *MLEC* | 0.661307069265069 | 4.29067042319736E-08 |
| *ETV6* | 0.68177143229294 | 1.81261211060184E-10 | *FKBP1A* | 0.77242583096682 | 4.52668665804244E-08 |
| *PSME1* | 0.877063893869009 | 1.8136248760368E-10 | *RABGAP1L* | 0.694988722836814 | 4.65941416227192E-08 |

| | | | | | |
|---|---|---|---|---|---|
| *MYDGF* | 0.876541943868654 | 1.81925361264775E-10 | *PCBP2* | 0.737935752739316 | 5.080002153133E-08 |
| *LSP1* | 0.975336351734126 | 1.8608335451337E-10 | *MOB1A* | 0.832896627799782 | 5.9557318511764E-08 |
| *KIF5B* | 0.974310922451692 | 1.87566168890249E-10 | *RPS15A* | 0.813622804907782 | 6.19178677973607E-08 |
| *DNASE1L3* | 0.702269852904961 | 1.94987755501298E-10 | *TAPBP* | 0.810697109102397 | 6.30288076790908E-08 |
| *CDC40* | 0.822466538021203 | 2.03768985861527E-10 | *ATP5F1C* | 0.631231281729958 | 6.31660467985916E-08 |
| *EEF1D* | 0.794067106037034 | 2.04526809784282E-10 | *RAC2* | 0.326836698781048 | 6.34403278044342E-08 |
| *SLC35E1* | 0.746026965620304 | 2.0579251402911E-10 | *TMEM219* | 0.650179731082149 | 6.42560095762865E-08 |
| *RPS17* | 0.785148552693125 | 2.17058274890403E-10 | *UVRAG* | 0.544034338183519 | 6.44048970468751E-08 |
| *GLIPR1* | 1.00672955416791 | 2.46064412037479E-10 | *ALCAM* | 0.527931555684777 | 7.80398310323863E-08 |
| *FCER1A* | 0.773634304213603 | 2.66865632924726E-10 | *TLR9* | 0.548149835103429 | 7.80398310323863E-08 |
| *IGF2R* | 0.812208886783467 | 2.70697663586686E-10 | *NDUFC2* | 0.806026127700588 | 7.91893899719683E-08 |
| *HYOU1* | 0.769140164891581 | 2.73115709373083E-10 | *MT-ND5* | 0.668399451982414 | 8.06653795442875E-08 |
| *SUMO2* | 0.968671628419216 | 2.75863063898864E-10 | *RPS17* | 0.565277932189218 | 8.22511910416886E-08 |
| *STRBP* | 0.720641647200391 | 2.81039642096931E-10 | *BTF3* | 0.866292183242098 | 8.28873013665978E-08 |
| *DPP7* | 0.665237640258418 | 2.81680121043013E-10 | *TRIM8* | 0.654172114581887 | 8.69368152812719E-08 |
| *ST3GAL2* | 0.691901751759336 | 2.95986308692278E-10 | *GLIPR1* | 0.820211952251826 | 8.98476212237648E-08 |
| *KTN1* | 0.986950904523555 | 2.96699495390374E-10 | *RPL28* | 0.743448993295366 | 8.99534528508493E-08 |
| *MEF2D* | 0.852442811586142 | 2.98202406335134E-10 | *TLN1* | 0.795036506838816 | 9.36879096575543E-08 |
| *CSDE1* | 0.882287322519607 | 3.05112917659572E-10 | *TAX1BP3* | 0.604276989772057 | 9.61152678689337E-08 |
| *CDK2AP2* | 0.837296784119072 | 3.13980688760136E-10 | *PTPRS* | 0.570914726235523 | 1.00779083981496E-07 |
| *PYCARD* | 0.851464116045666 | 3.29953736032254E-10 | *ARL6IP1* | 0.692964597578946 | 1.08797827728916E-07 |
| *RPS15A* | 0.734834713048843 | 3.32759483274707E-10 | *PPM1G* | 0.728724360696545 | 1.1094094296998E-07 |
| *GPM6B* | 0.713928058089335 | 3.6470509356472E-10 | *GRB2* | 0.789297839091516 | 1.12363505691575E-07 |
| *PET100* | 0.79008214569824 | 3.65473867429605E-10 | *MAP2K3* | 0.699532647981762 | 1.13705904498255E-07 |
| *GIT2* | 0.729153147055908 | 4.02950013775453E-10 | *GTF2I* | 0.711580275746293 | 1.24952456388688E-07 |
| *IRF2BPL* | 0.754390438554931 | 4.03358720107556E-10 | *RPS29* | 0.603322073338346 | 1.26619316645465E-07 |
| *DAD1* | 0.783695221790444 | 4.1069928398201E-10 | *ASPH* | 0.598832851081929 | 1.26640589178395E-07 |
| *PTP4A2* | 0.83602505215218 | 4.36567039017385E-10 | *TMED3* | 0.495576182372392 | 1.30019912326602E-07 |
| *SERBP1* | 0.9052473325619 | 4.63065048949428E-10 | *SNRPN* | 0.739405536358906 | 1.37934035790298E-07 |
| *SEPTIN9* | 0.825487481829439 | 4.68917040263968E-10 | *HNRNPM* | 0.705401700817168 | 1.41597099009952E-07 |
| *SLC38A2* | 1.01117979913007 | 4.72761929438794E-10 | *CHMP4B* | 0.627129953946981 | 1.42898232545803E-07 |
| *ATP13A2* | 0.740424259390928 | 5.359556066132E-10 | *XRCC5* | 0.691260291366697 | 1.46077574765367E-07 |
| *NCL* | 1.01144796429098 | 5.61798189966798E-10 | *EIF3I* | 0.64049717768481 | 1.59064862202618E-07 |
| *UVRAG* | 0.678888923759818 | 5.62198277000698E-10 | *DPP7* | 0.632172313033878 | 1.70294054916408E-07 |
| *TRIM44* | 0.761553670419773 | 6.1092105737876E-10 | *C1QBP* | 0.733191477760123 | 1.73627161812946E-07 |
| *ATP5MC3* | 0.858740628773 | 6.21869061261095E-10 | *ACTR2* | 0.750338685716696 | 1.82989201695867E-07 |
| *RPL13A* | 0.756993203990435 | 6.53522713211678E-10 | *RPL12* | 0.930655692078779 | 1.86176886951048E-07 |
| *ARPC5* | 0.855544715685618 | 6.73231109250485E-10 | *HNRNPU* | 0.686215477990645 | 1.92055600689533E-07 |

| | | | | | |
|---|---|---|---|---|---|
| *ZFAT* | 0.808941172949556 | 7.18315616160256E-10 | *UBE2D2* | 0.762613395355904 | 1.94000804059377E-07 |
| *LYPLA1* | 0.686084425164022 | 7.74053744799507E-10 | *NDUFA1* | 0.712118896264361 | 0.000000201034814410 9 |
| *TMEM109* | 0.766615763476034 | 7.97019359769759E-10 | *RPL32* | 0.844313517877649 | 2.1115552911556E-07 |
| *RBM25* | 0.927473836957836 | 8.13653479168078E-10 | *EIF1* | 0.652985032687796 | 2.12038046833594E-07 |
| *SMARCB1* | 0.690260101135215 | 8.45818632098842E-10 | *CDCA7L* | 0.480625840906421 | 2.15803709990842E-07 |
| *HDAC9* | 0.608575057943107 | 9.22860139512642E-10 | *NOTCH4* | 0.573735245297902 | 2.15803709990849E-07 |
| *WAC* | 0.743569919824782 | 9.45544819474664E-10 | *TXNL4A* | 0.67410557032763 | 2.2454500517255E-07 |
| *LIME1* | 0.995131228754652 | 9.60211783855294E-10 | *SEPTIN6* | 0.71921360064872 | 2.28384504799237E-07 |
| *SH3KBP1* | 0.876268265590134 | 1.04996953073688E-09 | *PRDX6* | 0.748750068336833 | 2.32194170910361E-07 |
| *BID* | 0.812776430079885 | 1.06051243483014E-09 | *TP53I13* | 0.64492553691103 | 2.61198248564757E-07 |
| *RPL23A* | 0.735118323578652 | 1.06771919930971E-09 | *PSME1* | 0.810945206052923 | 2.65181281879546E-07 |
| *CIITA* | 0.602316067471424 | 1.25401699652861E-09 | *ZDHHC17* | 0.593448604880379 | 2.70132366984606E-07 |
| *ETF1* | 0.691730988305164 | 1.25672598325429E-09 | *PRKCD* | 0.513314459612595 | 2.77635751872152E-07 |
| *N4BP2L1* | 0.754276197351174 | 1.27510807129287E-09 | *DNAJC3* | 0.64292391507935 | 2.8613252225521E-07 |
| *TIMM13* | 0.74481083552984 | 1.30323060576407E-09 | *RPN1* | 0.534702513662213 | 2.98845434894427E-07 |
| *PHB2* | 0.643132279979475 | 1.32368706151729E-09 | *SF3B5* | 0.652178982732445 | 3.02712898231638E-07 |
| *NDUFA11* | 0.769671055436997 | 1.34019767329417E-09 | *SMARCB1* | 0.603922562785656 | 3.12428292526904E-07 |
| *ITPR2* | 0.731365304464888 | 1.41908549161376E-09 | *N4BP2L1* | 0.668373344816519 | 3.21395167559213E-07 |
| *AC023590.1* | 0.671437649199621 | 1.47366272671555E-09 | *RPSA* | 0.717479780416853 | 3.24731291209805E-07 |
| *C15orf39* | 0.680243546574962 | 1.48556191744884E-09 | *ST13* | 0.749908394405392 | 3.39656135872094E-07 |
| *WAKMAR2* | 0.577004978942237 | 1.70163279882798E-09 | *ST3GAL2* | 0.601557488418465 | 3.48492753460273E-07 |
| *GRSF1* | 0.724052275363043 | 1.90540503511493E-09 | *NRP1* | 0.492598482572497 | 3.56854411055538E-07 |
| *OST4* | 0.853740782920557 | 2.00789251738542E-09 | *SLC9A7* | 0.519179081675814 | 3.56854411055568E-07 |
| *RPS25* | 0.668812458630512 | 2.07427387879568E-09 | *PIM3* | 0.843044113583892 | 3.57044087876311E-07 |
| *BTF3* | 0.787367761001204 | 2.10339308174025E-09 | *HNRNPC* | 0.745667106319989 | 3.69910499955262E-07 |
| *DHRS7* | 0.71555133410469 | 2.20096811316667E-09 | *EIF3H* | 0.767808066213955 | 3.87276974991519E-07 |
| *ZC3H15* | 0.722266761470668 | 2.26510697285728E-09 | *CLTC* | 0.585610684508714 | 4.37992005451973E-07 |
| *MGST2* | 0.544728575918481 | 2.30583976556388E-09 | *SCRN1* | 0.474602053278651 | 4.5825722080461E-07 |
| *DNAJC3* | 0.808941172949556 | 2.51830690682418E-09 | *SRSF9* | 0.685585067213348 | 4.65200987619802E-07 |
| *EID1* | 0.886383075649853 | 2.57800345453731E-09 | *ST6GALNAC 4* | 0.543230785043599 | 4.65324393652167E-07 |
| *RNF11* | 0.779193829555503 | 2.59157582311521E-09 | *ATP5ME* | 0.801542346381607 | 4.69178193981478E-07 |
| *CPNE3* | 0.676832111925826 | 2.80121547097517E-09 | *UQCR10* | 0.741590947127804 | 4.94564516043585E-07 |
| *SPCS2* | 0.898629060544123 | 2.82786002889235E-09 | *RPL23A* | 0.626088207289194 | 5.11312103078103E-07 |
| *EGLN3* | 0.624104756236711 | 3.12032001415418E-09 | *SNU13* | 0.700439718141092 | 5.30235427475072E-07 |
| *RPL8* | 0.73901621942151 | 3.12419351459178E-09 | *MAP3K8* | 0.708903660802483 | 5.34550352578466E-07 |
| *DBI* | 0.860987920719151 | 3.13092658637276E-09 | *SH3TC1* | 0.474602053278651 | 5.87940633774655E-07 |
| *SNRNP25* | 0.701789325869897 | 3.1483777060084E-09 | *MYCBP* | 0.462478494196161 | 5.87940633774673E-07 |

| | | | | | |
|---|---|---|---|---|---|
| *OSBPL8* | 0.837328005540259 | 3.18268292348252E-09 | *NEK8* | 0.536631250635163 | 5.87940633774673E-07 |
| *KLF10* | 0.849077923668007 | 3.21051178277905E-09 | *P2RY14* | 0.565729409282838 | 6.00049711674695E-07 |
| *MLEC* | 0.766788808664723 | 3.26873615875906E-09 | *MDM4* | 0.721475741491159 | 6.11485557171737E-07 |
| *SFT2D1* | 0.68041567507733 | 3.53570267919602E-09 | *ANAPC11* | 0.610087287762058 | 6.11973801993489E-07 |
| *KCTD5* | 0.716870550583672 | 3.70346559365281E-09 | *MX1* | 0.683565899576696 | 6.61845550917941E-07 |
| *AKIRIN1* | 0.548580901414087 | 3.78746846315912E-09 | *ZNRF2* | 0.524367304727442 | 6.86003270592386E-07 |
| *SEC31A* | 0.727904600858649 | 3.96367439656418E-09 | *STRBP* | 0.601557488418465 | 7.0106028638132E-07 |
| *TNFSF13B* | 0.671437649199621 | 4.00037241975514E-09 | *SEC11C* | 0.799695772534064 | 7.11933070224122E-07 |
| *PIK3AP1* | 0.671105861688036 | 4.18174454639137E-09 | *SMC6* | 0.623171301524527 | 7.59009577585485E-07 |
| *TLR7* | 0.627190736110454 | 4.21678728166839E-09 | *VPS13C* | 0.757094934998661 | 7.76923052228416E-07 |
| *LAMP5* | 0.63946906588889 | 4.21678728166864E-09 | *HNRNPK* | 0.76220634881626 | 9.0525979205512E-07 |
| *PLXNB2* | 0.621012161181329 | 4.21678728166864E-09 | *EIF3D* | 0.579229239761961 | 9.39045841356455E-07 |
| *CDYL* | 0.831145909190225 | 4.22119737691688E-09 | *COX7A2L* | 0.689066077953785 | 0.000000941125817481 7 |
| *UQCRB* | 0.830145017002259 | 4.35557853310857E-09 | *SON* | 0.74055178475353 | 9.42654272412373E-07 |
| *SLC3A2* | 0.644264249469395 | 4.36831497427023E-09 | *FMNL3* | 0.480625840906421 | 9.65189865627882E-07 |
| *TBL1XR1* | 0.827128306619206 | 4.40771014582816E-09 | *ZC3H15* | 0.580996165936148 | 9.71665119453195E-07 |
| *ATP5F1C* | 0.735355218621979 | 4.4317185317736E-09 | *RPS18* | 0.651641707243799 | 0.000001000554656471 7 |
| *PNN* | 0.895129554858177 | 4.46779472162867E-09 | *RPLP2* | 0.584193313055089 | 1.00372916168984E-06 |
| *HNRNPR* | 0.790148286445111 | 4.51000910134624E-09 | *NOP56* | 0.716891281105867 | 1.02337999248905E-06 |
| *RTN4* | 0.767878516327669 | 4.81422798632964E-09 | *PLAAT3* | 0.666021763137121 | 1.07234636163633E-06 |
| *RPL38* | 0.567777809048505 | 4.92958856628317E-09 | *BUD23* | 0.566631939835193 | 0.000001093608024388 7 |
| *MDFIC* | 0.78648632212186 | 4.97151624292403E-09 | *TRIM44* | 0.654996747379925 | 1.15439503318316E-06 |
| *MTPN* | 0.75988035449867 | 5.19442795296627E-09 | *HDGF* | 0.611928025488039 | 1.21656679973419E-06 |
| *KDELR1* | 0.709588362069637 | 5.26876711191625E-09 | *SERBP1* | 0.736454944385322 | 1.23199197529544E-06 |
| *RPL7* | 0.705957454913858 | 5.58749740538923E-09 | *PDXP* | 0.498547748903683 | 1.23502119263046E-06 |
| *CDCA7L* | 0.554487542380809 | 5.69092064470896E-09 | *PLXNB2* | 0.474602053278651 | 1.23502119263046E-06 |
| *NDUFB8* | 0.741313945884414 | 5.73091441429516E-09 | *TMEM170B* | 0.441013985007564 | 1.23502119263046E-06 |
| *GABARAPL2* | 0.736408963058707 | 5.95885213065775E-09 | *PET100* | 0.605182352237219 | 1.25891995489473E-06 |
| *GGNBP2* | 0.842990480136473 | 6.06023769202641E-09 | *NRDC* | 0.64049717768481 | 1.27832203664249E-06 |
| *PRDX6* | 0.728951074775882 | 6.48158521016645E-09 | *YPEL5* | 0.7865145160039 | 1.27920967389088E-06 |
| *TGFBR2* | 0.762033278397321 | 6.66569407684247E-09 | *UQCRB* | 0.726732500978839 | 1.29795767877179E-06 |
| *ASPH* | 0.674378924511743 | 6.86532347360125E-09 | *SEPTIN2* | 0.681388246271326 | 1.34514317476403E-06 |
| *ANXA11* | 0.677109621472147 | 7.00033680936341E-09 | *MAP4K4* | 0.753482806449555 | 1.35408307925427E-06 |
| *CAPRIN1* | 0.660113846297763 | 7.35371317011126E-09 | *ZMYM2* | 0.65739923759523 | 0.000001386511326184 7 |
| *SCAMP2* | 0.641432658686233 | 7.35918793455149E-09 | *PDIA4* | 0.667361741496178 | 1.40810578779791E-06 |
| *BCAP31* | 0.630436395614108 | 7.39830866298731E-09 | *HMGN3* | 0.68791136985266 | 1.42532510251864E-06 |

| | | | | | |
|---|---|---|---|---|---|
| *RPL30* | 0.639651838467706 | 7.61468136102797E-09 | *SCAMP2* | 0.590941353210578 | 1.44463204912956E-06 |
| *NOTCH2* | 0.586548751613108 | 7.67021097245865E-09 | *PSMA2* | 0.681494549557001 | 1.54490849099166E-06 |
| *ADPGK* | 0.669501540968418 | 7.68346507430777E-09 | *UBE2E2* | 0.453318494910685 | 1.57890157631536E-06 |
| *ELMO1* | 0.605164429780455 | 7.7169359456955E-09 | *ASIP* | 0.474602053278651 | 0.000001578901576315 4 |
| *GNA15* | 0.773957181189357 | 7.76970481538552E-09 | *EGLN3* | 0.498547748903683 | 0.000001578901576315 4 |
| *NUFIP2* | 0.844485289807223 | 8.01658113247613E-09 | *ERN1* | 0.670779744620853 | 1.58513796219951E-06 |
| *HMGN3* | 0.708110731494021 | 8.3747547067545E-09 | *COX6C* | 0.651623007073942 | 1.58988541227279E-06 |
| *ANXA5* | 0.771096361856907 | 8.45296957384331E-09 | *PEBP1* | 0.780206994938431 | 1.63516711926097E-06 |
| *CLTC* | 0.723237423581488 | 8.46098124191143E-09 | *N4BP2* | 0.626691378159779 | 1.68089008781097E-06 |
| *LMO4* | 0.710284120035352 | 8.54796273357946E-09 | *RAP1GDS1* | 0.543230785043599 | 1.71137769042593E-06 |
| *NR3C1* | 0.900823273957035 | 9.17333190506799E-09 | *SFPQ* | 0.736033319470832 | 1.72961321981653E-06 |
| *PPM1G* | 0.781628758472558 | 9.41041610771184E-09 | *ETF1* | 0.528230945433898 | 1.76047958036092E-06 |
| *EDEM1* | 0.643841976660617 | 9.41474626475364E-09 | *MAP3K2* | 0.644654803158466 | 1.77642465727342E-06 |
| *OS9* | 0.735355218621979 | 9.46277165216905E-09 | *ACADVL* | 0.641745723882069 | 1.85627638199207E-06 |
| *HNRNPUL1* | 0.738551845058158 | 9.64478607424E-09 | *RNF11* | 0.616861618493154 | 1.97030983736916E-06 |
| *PLXNA4* | 0.680243546574962 | 9.74676409303379E-09 | *MCOLN2* | 0.447179357813279 | 2.01677708689944E-06 |
| *KDELR2* | 0.740023232274826 | 9.98692230687607E-09 | *TMX1* | 0.564852124341917 | 2.02004474209624E-06 |
| *RPS20* | 0.830419152910677 | 1.02914508180423E-08 | *TBC1D4* | 0.593408949157093 | 2.14492117845377E-06 |
| *LHFPL2* | 0.570607207744086 | 1.03243240188791E-08 | *COX16* | 0.513640898134384 | 2.22244991350622E-06 |
| *MFSD2A* | 0.564180938584653 | 1.03243240188791E-08 | *NDUFB11* | 0.60001810832974 | 2.23665794155843E-06 |
| *NOTCH4* | 0.63946906588889 | 1.03243240188791E-08 | *SNRPD2* | 0.709845224375545 | 2.26284638423466E-06 |
| *TMED10* | 0.85474486256268 | 1.08610502081488E-08 | *ARPC5* | 0.672453944456471 | 2.55275859765859E-06 |
| *RPL27A* | 0.643558230688471 | 1.09031286805929E-08 | *SEMA7A* | 0.468553008605846 | 2.57386701932912E-06 |
| *RPS15* | 0.540789090230425 | 1.1588886383494E-08 | *MFSD2A* | 0.444099964881307 | 2.57386701932919E-06 |
| *ODC1* | 0.771444329950569 | 1.16800625280679E-08 | *MYCL* | 0.428603628558138 | 2.57386701932919E-06 |
| *RCC2* | 0.459915313082091 | 1.17511254079165E-08 | *TMEM8B* | 0.465518948516212 | 2.57386701932919E-06 |
| *SRSF5* | 0.764954462772247 | 1.19044313134514E-08 | *RPL8* | 0.715482617971092 | 2.63206769803415E-06 |
| *ARL6IP1* | 0.774734181364804 | 1.30788822541119E-08 | *AHCY* | 0.543230785043599 | 2.69034833510767E-06 |
| *RASSF5* | 0.740719681740868 | 1.33493776416911E-08 | *GIT2* | 0.562936194391157 | 2.70178150245964E-06 |
| *PGD* | 0.724331894520985 | 1.35441669941568E-08 | *CPNE3* | 0.548385556748889 | 2.70397825013392E-06 |
| *HEXB* | 0.557725916447354 | 1.38787707972417E-08 | *NENF* | 0.567482380386589 | 0.000002733242557575 7 |
| *NAMPT* | 1.33990878402998 | 1.38787707972421E-08 | *CIRBP* | 0.731618028982838 | 2.88218351114119E-06 |
| *FMNL3* | 0.544728575918481 | 1.38787707972425E-08 | *ELMO1* | 0.502993653934875 | 2.88352999991995E-06 |
| *EIF3H* | 0.814817739286225 | 1.40835908023172E-08 | *KCTD5* | 0.598752636740306 | 2.98140602207938E-06 |
| *SF3B5* | 0.713963457739459 | 1.49887124629045E-08 | *CSNK2B* | 0.563922652577207 | 3.05854344139772E-06 |
| *DDX3X* | 0.80667456597761 | 1.51113167679077E-08 | *GNA15* | 0.654996747379925 | 3.08753071332076E-06 |

| | | | | | |
|---|---|---|---|---|---|
| *SNU13* | 0.776937855615136 | 1.51371533643275E-08 | *RUBCN* | 0.66000611019367 | 3.08948351681857E-06 |
| *HADHA* | 0.703588172803327 | 1.51949543683488E-08 | *ATP5PB* | 0.583698037435576 | 0.000003090005995774<br>4 |
| *H2AFV* | 0.742252251167367 | 1.68748085098355E-08 | *LMO4* | 0.699938869310897 | 3.21028523176841E-06 |
| *NEDD8* | 0.743569919824782 | 1.71102916021926E-08 | *FIS1* | 0.580562207430579 | 3.49564905666881E-06 |
| *ULK1* | 0.712328524414549 | 1.76491328863817E-08 | *APH1A* | 0.564852124341917 | 3.54774712405071E-06 |
| *COMT* | 0.739665465368866 | 1.8361314406471E-08 | *NR3C1* | 0.795411866549764 | 3.59788321614091E-06 |
| *PEA15* | 0.501659854026595 | 1.86329191640316E-08 | *LMAN1* | 0.655746004753758 | 3.63364251253675E-06 |
| *SEMA7A* | 0.630270129042426 | 1.86329191640316E-08 | *DDX5* | 0.680277079670164 | 3.69606053396678E-06 |
| *TMED3* | 0.531613076128222 | 1.86329191640316E-08 | *PSMA6* | 0.693451172639507 | 4.03417548214187E-06 |
| *MCOLN2* | 0.54146086208457 | 1.86329191640327E-08 | *HHEX* | 0.437921389952182 | 0.000004181491195069<br>6 |
| *NEK8* | 0.645569265425308 | 1.86329191640327E-08 | *PEA15* | 0.453318494910685 | 0.000004181491195069<br>6 |
| *BAZ2B* | 0.644692793967092 | 1.86491416042476E-08 | *PLVAP* | 0.489614624133675 | 4.18149119506972E-06 |
| *PDIA3* | 0.854961337173468 | 2.01113507963496E-08 | *NECTIN1* | 0.441013985007564 | 4.18149119506984E-06 |
| *MDH2* | 0.715177336684349 | 2.03404989887092E-08 | *PNOC* | 0.545278799248674 | 4.18149119506984E-06 |
| *FTL* | 1.38984960395743 | 2.03859723076469E-08 | *PFDN5* | 0.692689037451876 | 4.28711688633306E-06 |
| *PRKAR1A* | 0.741654112982068 | 2.05644974430302E-08 | *SRP72* | 0.672051911559655 | 4.33740331801814E-06 |
| *SET* | 0.835178439332675 | 2.09332017755649E-08 | *RABAC1* | 0.630372960411103 | 4.46558126571273E-06 |
| *CHML* | 0.560957037710367 | 2.49836971375132E-08 | *DNAJC7* | 0.703833298161325 | 4.48930196577744E-06 |
| *TMEM14C* | 0.686084425164022 | 2.50179694749043E-08 | *LAPTM5* | 0.651112005777524 | 4.53336537714755E-06 |
| *KLC1* | 0.608243822712427 | 2.52165293720183E-08 | *UBXN4* | 0.648055137961078 | 4.86535490370017E-06 |
| *RAP1GDS1* | 0.608243822712427 | 2.52943480080572E-08 | *CMTM3* | 0.644239579370627 | 5.19224724817632E-06 |
| *RELT* | 0.617442759558891 | 2.55928222986735E-08 | *CDH1* | 0.428603628558138 | 5.32297327559026E-06 |
| *SLC2A1* | 0.607610367140937 | 2.6776547721351E-08 | *AC097375.1* | 0.41608558871171 | 5.32297327559056E-06 |
| *ATP5PB* | 0.716073743518676 | 2.85376567246536E-08 | *LYPLA1* | 0.593448604880379 | 5.51075700211898E-06 |
| *ITGAE* | 0.681896628200589 | 2.95368431153784E-08 | *HADHA* | 0.653370258999207 | 5.65121872158144E-06 |
| *TOMM20* | 0.758108535146365 | 3.04416537577505E-08 | *DNAJC4* | 0.596103058326357 | 5.73587347312786E-06 |
| *EWSR1* | 0.807390721386156 | 3.18988102224727E-08 | *PDIA3* | 0.67856521851833 | 5.99813233155384E-06 |
| *MYCBP* | 0.501659854026595 | 3.34567500256724E-08 | *SH3BGRL* | 0.674311890677054 | 6.34091785636059E-06 |
| *HIVEP1* | 0.686084425164022 | 3.50353843884353E-08 | *FLT3* | 0.431716240425474 | 6.77041360478488E-06 |
| *ABRACL* | 0.719487583431236 | 3.60782662005205E-08 | *APRT* | 0.748972895588939 | 6.97341457890431E-06 |
| *C17orf49* | 0.699759362061099 | 3.6785260174288E-08 | *APBB1IP* | 0.613061885486193 | 7.21678942354733E-06 |
| *RPL31* | 0.765151264060405 | 3.78219325498663E-08 | *PLXNA4* | 0.537550875971244 | 7.42475482620985E-06 |
| *NPC1* | 0.676000644434992 | 3.85652976470997E-08 | *RPS25* | 0.655496728338278 | 0.000007515828157521<br>9 |
| *BAZ2A* | 0.726360382755716 | 3.91736982176791E-08 | *KRT10* | 0.577106067868991 | 0.000007657171167253 |
| *UBXN4* | 0.74266795353039 | 4.04464906511602E-08 | *DDB1* | 0.666938419623963 | 7.88611702657792E-06 |
| *NRDC* | 0.691417869467063 | 4.15090973609056E-08 | *PGK1* | 0.736033319470831 | 7.94617231634032E-06 |

| | | | | | |
|---|---|---|---|---|---|
| *SMC6* | 0.691901751759336 | 4.28016336804379E-08 | *COX7C* | 0.613657681433425 | 7.97196249919282E-06 |
| *DNAJC7* | 0.790562643634701 | 4.39059703600655E-08 | *RPS10* | 0.32723231793223 | 8.24001249960321E-06 |
| *CHAMP1* | 0.54146086208457 | 4.47473478290991E-08 | *LAMTOR5* | 0.628946315013374 | 8.45909396563244E-06 |
| *NRP1* | 0.505018866116509 | 4.47473478291003E-08 | *ATP5MC2* | 0.622771008788437 | 8.48498599971998E-06 |
| *EIF4B* | 0.854481956859741 | 4.63896285579572E-08 | *TOMM20* | 0.654481407120511 | 8.53953518845562E-06 |
| *PBRM1* | 0.650927979750406 | 4.99677222073235E-08 | *ATF5* | 0.486624581527864 | 0.0000086043285516544 |
| *SND1* | 0.666145578110691 | 5.29054146552197E-08 | *AC023590.1* | 0.537550875971244 | 9.26954342037321E-06 |
| *RELL1* | 0.645083962582862 | 5.80997472571907E-08 | *MAPKAPK3* | 0.491288153282596 | 0.000009787011103206 |
| *PTGES3* | 0.777070657662534 | 5.85617313997704E-08 | *NAA38* | 0.554880395599843 | 0.0000107558824800755 |
| *TXNL4A* | 0.688753611940006 | 5.93466438976595E-08 | *ATP5F1D* | 0.601908680007562 | 0.0000109143040501784 |
| *XIST* | 0.872156849182932 | 5.94660661772433E-08 | *ADA* | 0.42548428671396 | 0.0000109260371478052 |
| *HHEX* | 0.570607207744086 | 5.97740325302168E-08 | *GALNT2* | 0.531008030104472 | 0.0000111420184307961 |
| *ARHGAP24* | 0.528315486027647 | 5.97740325302185E-08 | *CIAO2A* | 0.566631939835193 | 0.0000113162906922747 |
| *C4orf48* | 0.574191545093125 | 5.98233999527201E-08 | *RPL14* | 0.611117231619469 | 0.0000115180082162964 |
| *CBX6* | 0.779193829555503 | 6.0237333078616E-08 | *BID* | 0.569727642902059 | 0.000011750435584196 |
| *NDUFC2* | 0.811422162453311 | 6.09393366981414E-08 | *RPL30* | 0.664551054093173 | 0.0000118693066064089 |
| *SNHG9* | 0.761046482845244 | 6.16215287096677E-08 | *POLE4* | 0.491633791770327 | 0.0000123144738605302 |
| *RREB1* | 0.605164429780455 | 6.23023769213235E-08 | *TPM3* | 0.71307917981971 | 0.0000126930931846207 |
| *CAP1* | 0.785328615727605 | 6.47487510665146E-08 | *AP2S1* | 0.577244031075041 | 0.0000134643507358987 |
| *BUD23* | 0.660397781500373 | 6.60129899670606E-08 | *SEPTIN11* | 0.532828883475638 | 0.0000136540064569582 |
| *RABAC1* | 0.698362754942459 | 6.6630645626651E-08 | *ABRACL* | 0.603395550892713 | 0.0000141271396015412 |
| *SF1* | 0.847036475336555 | 6.85328517551938E-08 | *ING3* | 0.466651806871154 | 0.0000148847026467478 |
| *RILPL2* | 0.779193829555504 | 7.2280782735311E-08 | *SNHG9* | 0.702302462158282 | 0.0000149740121834233 |
| *RERE* | 0.647090293548158 | 7.62679293528176E-08 | *SND1* | 0.553034336434471 | 0.0000154818365487403 |
| *PFDN5* | 0.768546585355995 | 7.6438057130704E-08 | *UBE2E3* | 0.678843489334293 | 0.0000158595304616675 |
| *TP53I13* | 0.647949296277251 | 7.92912159618666E-08 | *PGLS* | 0.646392956388188 | 0.000015943761882343 |

| | | | | | |
|---|---|---|---|---|---|
| | | | | | 4 |
| *SHD* | 0.583374503817964 | 7.97488929793332E-08 | *OTULINL* | 0.684727957666821 | 0.000016939771350100 9 |
| *ARID1B* | 0.843661363433664 | 8.1205895829626E-08 | *RPS20* | 0.559800009553348 | 0.000017005989076402 |
| *P2RY14* | 0.641690305805569 | 8.12557705506257E-08 | *RFLNB* | 0.57472940753379 | 0.000017210725425514 2 |
| *ZBTB7A* | 0.770682329699986 | 8.31467414040844E-08 | *NMT1* | 0.567344065344933 | 0.000017543403433672 1 |
| *ATP5PF* | 0.73117231469256 | 8.41366815409676E-08 | *PROC* | 0.444099964881307 | 0.000017575099103020 4 |
| *RAC2* | 0.442813274151658 | 8.61390422802572E-08 | *ZDHHC24* | 0.381090167355506 | 0.000017575099103020 4 |
| *TMEM219* | 0.661836878917345 | 8.93494412453771E-08 | *TOR3A* | 0.374635145218207 | 0.000017575099103021 4 |
| *EIF3I* | 0.664551263614381 | 9.05097806714533E-08 | *SH3KBP1* | 0.654340827822871 | 0.000017657219610225 8 |
| *HSP90AA1* | 0.855266314458377 | 9.15030907995177E-08 | *TRIM38* | 0.536546213645902 | 0.000017746713691716 |
| *JMJD1C* | 0.814817739286225 | 9.25175566739101E-08 | *ANXA11* | 0.653654080888515 | 0.000017843685024655 7 |
| *SH3BGRL* | 0.796728488848015 | 9.30763982244922E-08 | *FAM118A* | 0.540393625740069 | 0.000018242266610049 7 |
| *ATP5F1D* | 0.741872203349769 | 9.87868410341555E-08 | *ZNF207* | 0.68454102299394 | 0.000018708266622384 8 |
| *AZIN1* | 0.560553543080163 | 1.01229028873691E-07 | *GRSF1* | 0.564852124341917 | 0.000019049599947114 1 |
| *AP2S1* | 0.659612213802831 | 1.01649187640546E-07 | *HNRNPAB* | 0.619949800280724 | 0.000019097831007513 3 |
| *AKAP13* | 0.874214021983925 | 1.01863837507084E-07 | *SMIM14* | 0.496204488272236 | 0.000019278453087411 6 |
| *COX16* | 0.521492868024751 | 1.05350346492637E-07 | *VPS36* | 0.575344498159559 | 0.000019708855511306 |
| *NECTIN1* | 0.570607207744086 | 1.06269517507742E-07 | *SLC2A1* | 0.533779779354525 | 0.000020447105545641 3 |
| *STAT2* | 0.525010341303976 | 1.06269517507742E-07 | *DGKZ* | 0.614481587641211 | 0.000020637118251593 2 |
| *NDUFB10* | 0.722432179694881 | 1.08694759281434E-07 | *ATP6V0E1* | 0.584904851721493 | 0.000020771328514502 2 |
| *PDIA4* | 0.735156345069182 | 1.12065156359761E-07 | *AC007381.1* | 0.437921389952182 | 0.000022263486121552 2 |
| *NDUFA1* | 0.825708624058248 | 1.12233583220747E-07 | *CIITA* | 0.42548428671396 | 0.000022263486121553 5 |
| *SNRPE* | 0.693657644868823 | 0.000000113268463816 1 | *CHAMP1* | 0.441013985007564 | 0.000022263486121554 1 |
| *NDUFB11* | 0.660773336634835 | 1.13331409722045E-07 | *ZRANB2* | 0.632574614991503 | 0.000022351657486209 8 |

| | | | | | |
|---|---|---|---|---|---|
| *COX7A2* | 0.757722299252679 | 1.16785603726091E-07 | *NPC1* | 0.513007442562894 | 0.0000230555996850073 |
| *BNIP2* | 0.695915434523716 | 1.26286105374338E-07 | *PDPK1* | 0.502993653934875 | 0.0000231289776344693 |
| *YWHAE* | 0.676293921971213 | 1.27129990178368E-07 | *TSPAN3* | 0.509946414576642 | 0.0000231319042125089 |
| *FUS* | 0.855175476902366 | 1.27452744543452E-07 | *WASF2* | 0.579424317179726 | 0.0000232608096646224 |
| *NOP56* | 0.731476327978037 | 1.3215223888773E-07 | *TALDO1* | 0.558628352907694 | 0.0000234617473563642 |
| *ING3* | 0.604888342316041 | 1.37513315636924E-07 | *HYOU1* | 0.56060437710497 | 0.0000239367706665711 |
| *SCRN1* | 0.505018866116509 | 1.41439177352449E-07 | *TOMM6* | 0.65767833744731 | 0.0000243795293606295 |
| *PLAUR* | 0.906573136031731 | 1.41439177352453E-07 | *GLRX* | 0.489573138172553 | 0.0000258849195855234 |
| *UBE2E2* | 0.477925205234234 | 1.41439177352453E-07 | *SEC63* | 0.543635638432221 | 0.0000265198469094551 |
| *SIRPB1* | 0.544728575918481 | 1.41439177352457E-07 | *GNA12* | 0.42548428671396 | 0.0000281802205893126 |
| *PDIA6* | 0.733390139942379 | 1.42935325139823E-07 | *RASSF5* | 0.598473962789103 | 0.0000284915500155275 |
| *RABGAP1L* | 0.75709253712705 | 1.48130748782517E-07 | *UQCR11* | 0.684934031534029 | 0.0000285399232661265 |
| *COX6C* | 0.736444182096166 | 1.48822356638022E-07 | *NDUFB10* | 0.639957347915784 | 0.0000296341588070143 |
| *GPATCH11* | 0.608243822712427 | 1.51709769758831E-07 | *WAC* | 0.500645899078452 | 0.0000302933030649063 |
| *GRINA* | 0.586548751613108 | 1.54922112294727E-07 | *EIF3F* | 0.732424529539137 | 0.0000304177887735503 |
| *KHSRP* | 0.558166950243285 | 1.60975766465215E-07 | *GPX4* | 0.670212272354868 | 0.0000306622094178668 |
| *ZNF791* | 0.878729503106418 | 1.63432224253596E-07 | *LSM4* | 0.577691021212476 | 0.0000313444963487361 |
| *IL10RA* | 0.781474770806742 | 1.71329905384528E-07 | *PSMB10* | 0.405909459378057 | 0.0000325613667718547 |
| *DNAJA1* | 0.780943611505463 | 1.75845640497099E-07 | *CREB3L2* | 0.575041442798527 | 0.0000330770911671084 |
| *KRTCAP2* | 0.75380851113559 | 1.80253781795515E-07 | *RREB1* | 0.467588317803676 | 0.0000337418030234782 |
| *NUDT1* | 0.674724562999474 | 1.84445220072551E-07 | *SUSD1* | 0.635922376439742 | 0.0000342801915296554 |
| *KIAA0100* | 0.635071259244699 | 1.86224327384846E-07 | *IGF2R* | 0.569136010726087 | 0.0000350035845069815 |
| *HNRNPL* | 0.675864582846143 | 1.97874613514191E-07 | *HEXB* | 0.387516436514939 | 0.000035641368945919 |

| | | | | | |
|---|---|---|---|---|---|
| | | | | | 1 |
| *DGKZ* | 0.727411287452957 | 2.00505030355567E-07 | *OS9* | 0.562699900059143 | 0.0000357273394764997 |
| *RRP7A* | 0.592780705324622 | 2.01933489144082E-07 | *MPG* | 0.510153445507741 | 0.0000364380581077533 |
| *GALNT2* | 0.654626993642424 | 2.04666362083857E-07 | *PRKAR1A* | 0.630280173644444 | 0.000036658702188298 |
| *PBX2* | 0.574003498293994 | 2.08078178010557E-07 | *GRASP* | 0.580996165936148 | 0.0000366996609380014 |
| *NME4* | 0.59898585485133 | 2.09593794576766E-07 | *ZNF791* | 0.666938419623963 | 0.0000370911576471607 |
| *COX7A2L* | 0.712079633696967 | 2.14811689360208E-07 | *ATP5PF* | 0.65705626955822 | 0.0000377373797192519 |
| *EIF3M* | 0.679984225627604 | 2.15425511295241E-07 | *RPL13A* | 0.475028645398896 | 0.0000378882904669178 |
| *FXR1* | 0.701438672041059 | 2.17328001338234E-07 | *ANAPC5* | 0.540704819667265 | 0.0000406185321296686 |
| *SUPT5H* | 0.714653577360793 | 2.18534366851215E-07 | *RRP7A* | 0.464598275197865 | 0.0000423029337942358 |
| *RFLNB* | 0.663274542255936 | 0.0000002238603351157 | *TCF3* | 0.327906947376784 | 0.0000432719576402527 |
| *DYNC1H1* | 0.746625967099073 | 2.29434948951559E-07 | *RPL36AL* | 0.613761724822863 | 0.0000436678727843643 |
| *LENG8* | 0.510377071127704 | 2.32291211927362E-07 | *TMEM179B* | 0.525448504762571 | 0.0000443736437540259 |
| *RPL36AL* | 0.688451439707938 | 2.44392919096003E-07 | *NIPSNAP2* | 0.387516436514939 | 0.0000450428698336025 |
| *ILF2* | 0.657352706620706 | 2.45811758263774E-07 | *CNDP2* | 0.487578601548187 | 0.0000455888429276041 |
| *PTPRS* | 0.511713518690518 | 2.49656278006414E-07 | *UBE2K* | 0.532086565455202 | 0.0000490284373380973 |
| *APBB1IP* | 0.722025306233136 | 2.52881841416906E-07 | *BAZ2B* | 0.52612352877343 | 0.0000498629360564241 |
| *NMT1* | 0.608081913162749 | 2.54439939349321E-07 | *RTF1* | 0.651385493827546 | 0.000050905422671453 |
| *SUSD1* | 0.704626485173427 | 2.61748756716445E-07 | *PRELID1* | 0.613381049758192 | 0.0000509490189915528 |
| *PSMA2* | 0.745822371556161 | 2.7242720541926E-07 | *NUDT22* | 0.534702513662213 | 0.0000509498192380423 |
| *PMEPA1* | 0.623542959095309 | 2.73608411164931E-07 | *C15orf39* | 0.464598275197865 | 0.0000539538788290104 |
| *CLINT1* | 0.795495641884604 | 2.82173633488822E-07 | *TAF10* | 0.658511231556162 | 0.0000547625650403985 |
| *VPS29* | 0.609912744678618 | 2.86630197042812E-07 | *RPL38* | 0.473915619316868 | 0.0000567694646876227 |
| *COPA* | 0.705713231912955 | 3.05123097652831E-07 | *ETV6* | 0.504428457674111 | 0.000056911089661724 |

| | | | | | |
|---|---|---|---|---|---|
| | | | | | 1 |
| *KDM5A* | 0.783752111517808 | 3.17055370545996E-07 | *H2AFV* | 0.638039015493771 | 0.0000588960955246792 |
| *TMX1* | 0.624325402416064 | 3.24303045515511E-07 | *TBL1XR1* | 0.59853798724341 | 0.000065698727885882 |
| *DDB1* | 0.762705706766934 | 3.25008748950234E-07 | *FYTTD1* | 0.638836043314889 | 0.0000666493646195045 |
| *GLRX5* | 0.642691829650642 | 3.29701576095631E-07 | *TRIR* | 0.601613039718635 | 0.0000671356265699096 |
| *DDT* | 0.527854757354085 | 3.30177215097186E-07 | *MT-ND3* | 0.600465317778929 | 0.0000701867534963393 |
| *FAM160A1* | 0.505018866116509 | 3.31103701483146E-07 | *ANAPC16* | 0.655585482356832 | 0.0000707723196824178 |
| *VPS36* | 0.655535361804415 | 3.35976211490238E-07 | *SLC7A11* | 0.387516436514939 | 0.0000717738217318265 |
| *UQCR10* | 0.745246497632166 | 3.47774562249581E-07 | *HLA-DOA* | 0.428603628558138 | 0.0000717738217318307 |
| *CSNK2B* | 0.598621583913683 | 3.51250085969016E-07 | *NUDT5* | 0.368151111648008 | 0.0000717738217318307 |
| *TRIM38* | 0.714098801333619 | 3.52709878386265E-07 | *HDAC9* | 0.447179357813279 | 0.0000717738217318327 |
| *TET2* | 0.638595499382703 | 3.5490693017806E-07 | *PDIA6* | 0.591587577203863 | 0.0000748903753769875 |
| *GLRX* | 0.626077115799745 | 3.78289612438311E-07 | *SUPT5H* | 0.598355510057735 | 0.0000752649533395301 |
| *SAP18* | 0.796178009186382 | 3.79249968092613E-07 | *IER3IP1* | 0.475457704246496 | 0.0000754852143113039 |
| *LAMTOR5* | 0.663123276720711 | 3.86712535531259E-07 | *NAPA* | 0.629823676170358 | 0.0000769098396830463 |
| *APRT* | 0.775721637648699 | 3.88151085449535E-07 | *KDELR2* | 0.575344498159559 | 0.0000793106773256894 |
| *IER3IP1* | 0.561727579527069 | 3.95765332756589E-07 | *NDUFA7* | 0.579396725593956 | 0.0000813466462498805 |
| *PRELID1* | 0.759505494633391 | 4.05197733176492E-07 | *CSNK1E* | 0.515484598118072 | 0.0000815956826291716 |
| *YWHAB* | 0.751419163453899 | 4.16746904356171E-07 | *AP3S1* | 0.581204181947018 | 0.0000820908241633464 |
| *AP3S1* | 0.692657079087508 | 4.21816432389082E-07 | *CYSTM1* | 0.467588317803676 | 0.000085088720173026 |
| *ACTN4* | 0.700439718141092 | 4.2728816954182E-07 | *PSMG2* | 0.504428457674111 | 0.0000855211806932696 |
| *USP24* | 0.592597796285913 | 4.32434007308405E-07 | *ATP5MC1* | 0.551556619373095 | 0.0000903000594396525 |
| *FLT3* | 0.488145047552113 | 4.3861363172015E-07 | *HHIP-AS1* | 0.381090167355506 | 0.0000904982490883054 |
| *ARHGAP17* | 0.607922402405675 | 4.50336929120694E-07 | *LHFPL2* | 0.387516436514939 | 0.000090498249088305 |

4

| | | | | | |
|---|---|---|---|---|---|
| *HNRNPAB* | 0.720300140501935 | 4.65261512512639E-07 | *SHD* | 0.525019960264874 | 0.0000904982490883054 |
| *NUDT22* | 0.580562456547376 | 4.78993926203109E-07 | *FAM160A1* | 0.35181237483326 | 0.0000904982490883079 |
| *IQSEC1* | 0.62486568316421 | 4.91405671809338E-07 | *PARVB* | 0.493203150742066 | 0.000099843405835112 |
| *ST13* | 0.774678348433011 | 4.93411536601606E-07 | *DYNLL1* | 0.563625434809182 | 0.000103344968865652 |
| *TMEM30A* | 0.616543491910945 | 4.99063732269485E-07 | *TUBB* | 0.603022472266147 | 0.000103952099476188 |
| *OTULINL* | 0.764294953176165 | 5.21653212574065E-07 | *CIAO2B* | 0.568285114186594 | 0.00010619390699467 |
| *PAK2* | 0.703868256399837 | 5.54108405561668E-07 | *MFSD12* | 0.449554394973415 | 0.0001105857034625 |
| *ADA2* | 0.668854054502302 | 5.59941255703977E-07 | *BORCS7* | 0.561720064300693 | 0.000111736496887706 |
| *RTF1* | 0.717008998358922 | 5.6107208339257E-07 | *APPL1* | 0.598473962789103 | 0.000112320363953927 |
| *ELF2* | 0.664815808410371 | 5.69251617346493E-07 | *ARPC5L* | 0.617302224032189 | 0.000113038300670461 |
| *FNBP1* | 0.700384988714827 | 5.76648063660377E-07 | *MYD88* | 0.361637804689334 | 0.000114021696535233 |
| *KCTD12* | 0.554487542380809 | 5.80365416687531E-07 | *PGD* | 0.498680661063793 | 0.000117046873610297 |
| *TLR9* | 0.505018866116509 | 5.80365416687531E-07 | *DNAJA1* | 0.590840942920943 | 0.000117658744962185 |
| *NREP* | 0.457265734668141 | 5.80365416687546E-07 | *AKIRIN1* | 0.329316517631455 | 0.000117808429371048 |
| *PLVAP* | 0.557725916447354 | 5.80365416687546E-07 | *SP110* | 0.673346057258817 | 0.00012273173434422 |
| *ANAPC11* | 0.605164429780455 | 5.84440199431302E-07 | *UBE2V1* | 0.600719686396727 | 0.000124900913383513 |
| *ATP5MC2* | 0.71169506884306 | 6.14604852132569E-07 | *ANKRD12* | 0.752607544006347 | 0.000130422923840579 |
| *PSMA6* | 0.716978433109806 | 6.32224896056371E-07 | *KIAA0100* | 0.454784560892255 | 0.000141064972079479 |
| *HDLBP* | 0.595886616192444 | 6.40510734126525E-07 | *GTF3A* | 0.657214572629744 | 0.000146168844923836 |
| *CIAO2A* | 0.62246832586805 | 6.53962754632297E-07 | *RPL7* | 0.472990777045141 | 0.0001504338566831 |
| *CORO1A* | 0.828510147618683 | 6.77647140639253E-07 | *IL16* | 0.473420538551528 | 0.000152736438198201 |
| *LSM4* | 0.589532610051929 | 7.51450365400222E-07 | *KIAA2013* | 0.475457704246496 | 0.000153280297414153 |
| *UBL5* | 0.784337132831463 | 7.56069873252781E-07 | *ILF2* | 0.593560863763903 | 0.000159137843898503 |
| *HLA-DOA* | 0.501659854026595 | 7.67055794200051E-07 | *EIF4B* | 0.63109847968256 | 0.000162337111393878 |
| *LMAN1* | 0.741225979356483 | 7.9185931654011E-07 | *LUC7L3* | 0.615161394142041 | 0.000166502287393625 |
| *PLAAT3* | 0.65006520236304 | 8.01692149475939E-07 | *LSM3* | 0.588773516527868 | 0.000166763344644963 |
| *AC103591.3* | 0.74318887610001 | 8.19568022373794E-07 | *PLIN3* | 0.378636808297922 | 0.000170927251121985 |
| *ICAM1* | 0.812302141367442 | 8.56490300434486E-07 | *KLC1* | 0.443492642186213 | 0.000173471382823902 |
| *COX7C* | 0.642805360639556 | 8.59615561048748E-07 | *NOTCH2* | 0.368151111648008 | 0.000180598100115074 |
| *ATP6V1F* | 0.651889505280426 | 8.77284611099184E-07 | *GRK3* | 0.355094958822562 | 0.00018059810011509 |
| *RPS3* | 0.56906863127854 | 8.86023730203906E-07 | *SLC35F3* | 0.358370090855423 | 0.000180598100115101 |
| *MAN1A1* | 0.665022809636193 | 8.93423012901859E-07 | *TCL1A* | 1.39747168531823 | 0.000184774647939658 |
| *FAM118A* | 0.561348197487965 | 8.94870194493841E-07 | *GLRX5* | 0.526840395889243 | 0.00018964621302115 |
| *PNRC1* | 0.851762455506148 | 9.0380206344199E-07 | *MIIP* | 0.394200012522864 | 0.00019421193168632 |
| *NAA38* | 0.629446710050821 | 9.22883108970908E-07 | *SNRPF* | 0.608776888361467 | 0.000199334324371326 |
| *TMEM14B* | 0.67449645088881 | 9.70025046439311E-07 | *ZDHHC4* | 0.474424501738104 | 0.000200005298380711 |

| | | | | | |
|---|---|---|---|---|---|
| *SEPTIN6* | 0.772961875843989 | 9.83621469612997E-07 | *FEZ2* | 0.529237753185402 | 0.000200072417763478 |
| *HSD17B4* | 0.460729589971713 | 1.01265866709397E-06 | *NFATC2IP* | 0.593696560619261 | 0.000202172965632331 |
| *MEF2A* | 0.62486568316421 | 1.01372061858285E-06 | *RELL1* | 0.490393617155242 | 0.000206618912137723 |
| *PLEKHO1* | 0.655458461133259 | 1.05949770749135E-06 | *ADI1* | 0.567482380386589 | 0.000207337828918397 |
| *AC007952.4* | 0.808941172949556 | 1.06147384865783E-06 | *RCC2* | 0.312771238259335 | 0.000207707176781691 |
| *FYTTD1* | 0.867730504157161 | 1.07250592337763E-06 | *ITCH* | 0.501675522238896 | 0.000215071120726943 |
| *WDR26* | 0.489687212360519 | 1.07690659104391E-06 | *GPAA1* | 0.437405312307298 | 0.000219795669085748 |
| *USP7* | 0.688681536650061 | 0.0000010897885668444 | *RUBCNL* | 0.355094958822562 | 0.000227037292614813 |
| *SRSF6* | 0.697282631768886 | 1.14429269811354E-06 | *HSD17B4* | 0.381090167355506 | 0.000227037292614826 |
| *ZFP91* | 0.670817668510711 | 1.16473365544294E-06 | *GTF2A2* | 0.483628329067944 | 0.000232623697277085 |
| *SERINC1* | 0.696444936769744 | 1.22819231748516E-06 | *SRSF5* | 0.540961504133896 | 0.000237047845425873 |
| *POLR2J3* | 0.770033830270027 | 1.29824561886692E-06 | *HNRNPA3* | 0.649077523191405 | 0.000250234658956954 |
| *LASP1* | 0.618131396822987 | 1.32202354612475E-06 | *PPP1R2* | 0.566719251545262 | 0.000257036979158849 |
| *ELL* | 0.446823987946494 | 1.33540973549469E-06 | *RTF2* | 0.510986444688406 | 0.000260791965055802 |
| *PPP1R16B* | 0.457265734668141 | 1.33540973549469E-06 | *ROMO1* | 0.563535195869926 | 0.000270140263927165 |
| *PSMG2* | 0.533283601512919 | 1.35394429044883E-06 | *AC103591.3* | 0.591014181653557 | 0.000271724439650231 |
| *ATAD2B* | 0.565175100820541 | 0.0000013715241083422 | *LSM7* | 0.608486782782189 | 0.000278127250670909 |
| *CMTM3* | 0.663931282925472 | 1.38012962592059E-06 | *NBDY* | 0.620269369457109 | 0.000280057182890564 |
| *CIAO2B* | 0.664551263614381 | 1.39132292094401E-06 | *TMEM256* | 0.504386889234741 | 0.000281885013806519 |
| *ERN1* | 0.788182612782758 | 1.42796369513311E-06 | *MRPS10* | 0.36489813385292 | 0.000285210671002315 |
| *POLB* | 0.677572925308505 | 1.46821988826418E-06 | *NSMCE4A* | 0.331958460024627 | 0.000285210671002323 |
| *MT-ND4L* | 0.659403882388622 | 1.50372966703198E-06 | *STAT2* | 0.371396771151662 | 0.000285210671002323 |
| *CSF2RA* | 0.605164429780455 | 1.51598693213781E-06 | *COL26A1* | 0.412939033394846 | 0.000285210671002332 |
| *ACBD5* | 0.561348197487965 | 1.53694757935508E-06 | *PLEKHD1* | 0.387516436514939 | 0.000285210671002332 |
| *SCAND1* | 0.620101311324134 | 0.0000015760411910876 | *PPP1R14B-AS1* | 0.36489813385292 | 0.000285210671002332 |
| *ACADVL* | 0.580648150173423 | 1.68451705673261E-06 | *HIP1* | 0.361637804689334 | 0.000285210671002349 |
| *APH1A* | 0.562800148840499 | 1.69256792054323E-06 | *UFC1* | 0.56825507337324 | 0.000291979423432817 |
| *SASH3* | 0.536471879143966 | 1.72524396244373E-06 | *PTPN2* | 0.603395550892713 | 0.000292704475852533 |
| *VASH2* | 0.57380963978243 | 1.75908124102835E-06 | *OSBPL8* | 0.569762666147576 | 0.000294476906110075 |
| *NUDT5* | 0.450312974026374 | 1.75908124102841E-06 | *TBCA* | 0.63252646657623 | 0.000301621982238222 |
| *LSM3* | 0.589331192284435 | 1.76571593641798E-06 | *GGNBP2* | 0.667223049713185 | 0.000311933622563042 |
| *COG3* | 0.625542883228971 | 1.94504552450176E-06 | *HNRNPR* | 0.564980540830576 | 0.000312076643607077 |
| *SNX18* | 0.589621585662731 | 2.12260814734012E-06 | *NEMF* | 0.497640419440668 | 0.000315683823178977 |
| *ROMO1* | 0.646619670467524 | 2.25820246394509E-06 | *C4orf48* | 0.451854804943074 | 0.000320879629836616 |
| *APEX1* | 0.69902348087152 | 0.0000022775661814013 | *POLB* | 0.474424501738104 | 0.00033308028259385 |
| *RNF144B* | 0.531613076128222 | 2.31463064817807E-06 | *SRGN* | 0.632856405601338 | 0.000335092248807666 |

| | | | | | |
|---|---|---|---|---|---|
| *ATF5* | 0.477925205234234 | 2.31463064817819E-06 | *UBE2L3* | 0.547493114101735 | 0.000345613504289228 |
| *SEC61A1* | 0.642691829650642 | 2.40159391571622E-06 | *TMSB10* | 0.551488585727019 | 0.000354553637199974 |
| *SEPTIN11* | 0.604888342316041 | 2.43073788762856E-06 | *CDH23* | 0.397102485344137 | 0.000358031881334609 |
| *CTDNEP1* | 0.622308514730707 | 2.48473474754368E-06 | *SMARCE1* | 0.558135208127336 | 0.000364562404179613 |
| *SNRPN* | 0.687301375924404 | 2.50911403415297E-06 | *RBM25* | 0.645446393656543 | 0.000371227011864053 |
| *UQCR11* | 0.69227625887829 | 2.51830325266071E-06 | *PARP10* | 0.50024304291884 | 0.000371336371269277 |
| *ZRANB2* | 0.757301223706662 | 2.53247734191667E-06 | *ENO1* | 0.594183304162991 | 0.000372754961528845 |
| *PEBP1* | 0.853394771396921 | 2.57703841307411E-06 | *HSP90AA1* | 0.662077095729761 | 0.000373923355129784 |
| *SEC11C* | 0.774678348433011 | 2.58085805987281E-06 | *PTGDS* | 2.8141600421878 | 0.000386154347695406 |
| *ITCH* | 0.636493182789512 | 2.58588446153583E-06 | *AKR7A2* | 0.420770528627201 | 0.000387271171962681 |
| *SMARCE1* | 0.633940806200634 | 0.0000025920814285882 | *ZNF652* | 0.538135460055132 | 0.000388861927166356 |
| *TBC1D1* | 0.545371344938197 | 2.69572369293261E-06 | *TOMM7* | 0.436957895637287 | 0.000403066605744475 |
| *CSNK1E* | 0.553312422139176 | 2.78123441388947E-06 | *CAPN15* | 0.470572176242498 | 0.000405125828992194 |
| *APPL1* | 0.6810134356166 | 2.79214199107388E-06 | *SCAND1* | 0.531008030104472 | 0.000412923909748238 |
| *ELOB* | 0.582796616752 | 2.79624588466219E-06 | *GPATCH11* | 0.452575746948652 | 0.000413239793226036 |
| *TRABD* | 0.671211148682492 | 2.80622189406724E-06 | *TCEA1* | 0.599894637488731 | 0.000419519624017159 |
| *PTPN2* | 0.696444936769744 | 0.0000028353246566375 | *PTGES3* | 0.589724145763652 | 0.000447032756784653 |
| *IL16* | 0.571536180758083 | 0.0000028496131513632 | *TP53I11* | 0.371396771151662 | 0.000449125581722111 |
| *KRT10* | 0.564817369399409 | 2.91697647047079E-06 | *PTP4A2* | 0.556932461045679 | 0.000453917374334482 |
| *PTEN* | 0.583798140597073 | 3.07529503585195E-06 | *SRM* | 0.480183771026789 | 0.000460454668349597 |
| *UBE2K* | 0.597953514885357 | 3.07780536672307E-06 | *HINT2* | 0.497487177628332 | 0.0004642716943635 |
| *UBE2D2* | 0.724450864961072 | 0.0000031265454651153 | *KLF10* | 0.624984178313485 | 0.00047939349790322 |
| *BOD1L1* | 0.683845952968023 | 3.18883387900013E-06 | *RPS15* | 0.572321005303614 | 0.000484927425850458 |
| *OTUB1* | 0.401988403802399 | 0.0000032893825253712 | *MRPL36* | 0.475457704246496 | 0.000494701663075534 |
| *GPX4* | 0.800759179390437 | 0.0000033426878681151 | *PTRHD1* | 0.591084983559732 | 0.000511022280837989 |
| *KCTD20* | 0.528650367904906 | 3.44888797105905E-06 | *ACAA1* | 0.505819698013638 | 0.000514441613594486 |
| *PDPK1* | 0.580474237203423 | 3.50689648401482E-06 | *PARVG* | 0.578942507168174 | 0.00051449060332267 |
| *YTHDF2* | 0.663123276720711 | 3.94945769567602E-06 | *ULK1* | 0.521001603085638 | 0.00052432010444362 |
| *ENO1* | 0.734379982204803 | 3.97711722435726E-06 | *SF1* | 0.697952312281998 | 0.000531495743363089 |
| *SLC15A3* | 0.474502439593445 | 3.99451976973838E-06 | *STUB1* | 0.425568681989709 | 0.000531740570604737 |
| *SLC7A11* | 0.515049231253774 | 3.99451976973838E-06 | *RPL27A* | 0.332099145876495 | 0.00053648686727247 |
| *ARMCX6* | 0.429251358498545 | 3.99451976973849E-06 | *HDLBP* | 0.437405312307298 | 0.000537381122350168 |
| *RBM47* | 0.446823987946494 | 3.99451976973849E-06 | *ARPC4* | 0.566496596241033 | 0.000539866090309556 |
| *ZDHHC24* | 0.432783049468798 | 3.99451976973849E-06 | *YWHAB* | 0.593043258577298 | 0.000549034396420839 |

| | | | | | |
|---|---|---|---|---|---|
| *GRASP* | 0.696731669363531 | 4.00034860158019E-06 | *ARMCX6* | 0.338606836033993 | 0.000562997989108926 |
| *ARFGAP3* | 0.624325402416064 | 4.20096116852254E-06 | *SLC12A2* | 0.335286477742381 | 0.000562997989108943 |
| *PCF11* | 0.771962260324428 | 4.48803803581094E-06 | *ANXA5* | 0.487115909430871 | 0.000593536256381374 |
| *KLF13* | 0.681896628200588 | 4.49835576974384E-06 | *TGFBR2* | 0.528756749999957 | 0.000596576624676519 |
| *SRSF7* | 0.936294790640232 | 4.59224976033313E-06 | *AZIN1* | 0.353956018966386 | 0.000604350828319128 |
| *RRAGC* | 0.564853680513789 | 4.74410226979894E-06 | *CAPRIN1* | 0.440709375680532 | 0.000635154656874762 |
| *TBC1D4* | 0.567973022357968 | 4.75017508246894E-06 | *ARMH1* | 0.508640217076017 | 0.000654315359210476 |
| *SP110* | 0.71779328489136 | 4.84645970374387E-06 | *TNFSF13B* | 0.440452187866162 | 0.000666082551084325 |
| *ABI2* | 0.545371344938197 | 4.94508381946547E-06 | *SYS1* | 0.513943953495416 | 0.000687335218036622 |
| *HNRNPH1* | 0.734206374782194 | 5.00135800610526E-06 | *CSF2RA* | 0.42515305148328 | 0.000692496330657972 |
| *PARVB* | 0.553312422139176 | 5.01483282151287E-06 | *LINC01184* | 0.315202231699049 | 0.000705247646366013 |
| *TYMP* | 0.710606744298091 | 5.09581597051755E-06 | *ARHGAP24* | 0.381090167355506 | 0.000705247646366034 |
| *LSM7* | 0.687189515253206 | 5.16282696745783E-06 | *IFIT2* | 0.403457980383961 | 0.000705247646366034 |
| *ERGIC3* | 0.600790674188887 | 5.22055791429703E-06 | *LEPROT* | 0.382699420217163 | 0.000721697636333686 |
| *PDXP* | 0.453793542761336 | 0.0000052391462332891 | *FBXW5* | 0.443572567675327 | 0.000730433368942952 |
| *RUBCNL* | 0.443326543710267 | 0.0000052391462332891 | *C18orf32* | 0.574001765897903 | 0.000731008396517419 |
| *TEX2* | 0.460729589971713 | 5.23914623328926E-06 | *DAZAP2* | 0.568396129156514 | 0.000744596106259323 |
| *CYC1* | 0.45045617667088 | 5.33804757344863E-06 | *MTRNR2L1* | 0.800229543906332 | 0.000764132403207408 |
| *GHITM* | 0.628217930914181 | 5.34875776932515E-06 | *SRSF7* | 0.688813432141131 | 0.000775232839593634 |
| *LRRFIP1* | 0.758520261290754 | 5.35968992439801E-06 | *CALR* | 0.592963350795789 | 0.00078755317576462 |
| *C18orf32* | 0.611399615563769 | 5.44258390413375E-06 | *PMEPA1* | 0.464598275197865 | 0.000804393447537403 |
| *SRSF9* | 0.72003175708736 | 5.50577015243303E-06 | *ITPR2* | 0.537427150998749 | 0.000810527311248504 |
| *NUCKS1* | 0.582293010943373 | 0.0000056436855464271 | *ZNF511* | 0.434351988329744 | 0.000830091669320035 |
| *C6orf62* | 0.644617807868638 | 5.70867186925298E-06 | *NDUFB4* | 0.542331801885376 | 0.000858565669569013 |
| *TFRC* | 0.60059652609874 | 5.88116177599693E-06 | *SIRPB1* | 0.374635145218207 | 0.000882825738885619 |
| *ANP32A* | 0.700091283157901 | 6.33540701826843E-06 | *TEX2* | 0.338606836033993 | 0.000882825738885696 |
| *BCL3* | 0.604360027729568 | 6.40362056594348E-06 | *ADPGK* | 0.516668590831952 | 0.000892412223719468 |
| *TTC7A* | 0.493022924923775 | 6.60746453189575E-06 | *EIF3M* | 0.516323984347359 | 0.00089520848991166 |
| *SINHCAF* | 0.632904387304847 | 6.64368274068482E-06 | *HIVEP1* | 0.485920140575933 | 0.000904658035738096 |
| *TSPAN3* | 0.592510309851825 | 6.72304721987999E-06 | *LDLRAD4* | 0.572035298364526 | 0.000913197673601282 |
| *BORCS7* | 0.580562456547376 | 0.0000067598257051671 | *TMEM243* | 0.517414169681046 | 0.00091557628410592 |
| *CDKN1A* | 0.686084425164022 | 7.42084048823739E-06 | *CHPT1* | 0.442330872028251 | 0.000950105336400599 |
| *CSNK1A1* | 0.628052303792607 | 7.68104426317144E-06 | *ELOB* | 0.416514133209407 | 0.000969992702974914 |
| *NASP* | 0.662497604846406 | 8.17776845538287E-06 | *CORO1A* | 0.71324813147065 | 0.000970975087410355 |
| *LDLRAD4* | 0.669010911805081 | 8.48247771062142E-06 | *RALBP1* | 0.356249022591717 | 0.00100840803143295 |
| *ARPC5L* | 0.663123276720711 | 8.57387734849689E-06 | *SNX18* | 0.430880675849333 | 0.00101282506809334 |

| | | | | | |
|---|---|---|---|---|---|
| *SMIM14* | 0.577813941635499 | 8.69420298831931E-06 | *CYC1* | 0.386418077085496 | 0.00103235018121008 |
| *UBE2I* | 0.678852855932946 | 8.72378665186369E-06 | *UNC119* | 0.481932038407045 | 0.00105399771451888 |
| *OIP5-AS1* | 0.626626190988443 | 8.83128547816448E-06 | *RWDD1* | 0.553700615964573 | 0.00106298373044965 |
| *ITPR1* | 0.592597796285913 | 8.87914439763675E-06 | *POLR2L* | 0.526840395889243 | 0.00106470312758481 |
| *AC007381.1* | 0.418604114299036 | 8.98428093395687E-06 | *HLA-B* | 0.488562498942364 | 0.00107556386916403 |
| *NIPSNAP2* | 0.450312974026374 | 8.98428093395687E-06 | *SLC38A2* | 0.637130326586412 | 0.00108163170645772 |
| *ADA* | 0.418604114299036 | 8.98428093395711E-06 | *BAZ2A* | 0.563625434809182 | 0.00108626138580269 |
| *CDH1* | 0.432783049468798 | 8.98428093395711E-06 | *SINHCAF* | 0.516668590831952 | 0.00108709865228165 |
| *TBC1D9* | 0.429251358498545 | 8.98428093395711E-06 | *TBC1D1* | 0.412795844963297 | 0.00109498494073533 |
| *PABPN1* | 0.602728678668542 | 9.46267089827124E-06 | *KCTD12* | 0.338606836033993 | 0.0011043574193298 |
| *NDUFB7* | 0.493284957469123 | 9.48230126585476E-06 | *SH3BGRL3* | 0.656475676305838 | 0.001126562688735 |
| *TMEM256* | 0.562453075995645 | 9.50072755585521E-06 | *COG3* | 0.501675522238896 | 0.00114857046553204 |
| *TMEM50A* | 0.595886616192444 | 0.0000103973820750128 | *HIGD2A* | 0.560837728507159 | 0.00114891803757479 |
| *RPSA* | 0.652392130352795 | 0.0000104943082584665 | *CD99* | 0.4578416175425 | 0.0011517850913691 |
| *TMEM179B* | 0.540353928734502 | 0.0000106718657195469 | *NDUFB7* | 0.421865822342508 | 0.00116013217166666 |
| *LAMP2* | 0.566084649053392 | 0.0000108653699467557 | *CLINT1* | 0.56533070953154 | 0.00116793289914252 |
| *SNRNP27* | 0.597953514885357 | 0.0000109259379887894 | *RER1* | 0.428646312474794 | 0.00117109930603616 |
| *MAP2K6* | 0.509586769798224 | 0.000011306138069112 | *EWSR1* | 0.547847889719135 | 0.00118844198872558 |
| *FAM49B* | 0.678325001705914 | 0.0000114124153353784 | *TRMT112* | 0.507566383724699 | 0.00120574366559296 |
| *MPZL1* | 0.502984034973977 | 0.0000115340085751214 | *FXR1* | 0.539341408465734 | 0.00122107005278777 |
| *VAPA* | 0.703082501515919 | 0.0000116158711893122 | *TMEM30A* | 0.459924932042989 | 0.00122610495069374 |
| *HNRNPDL* | 0.690086355721518 | 0.0000118055937846295 | *RBCK1* | 0.491633791770327 | 0.00124269243627638 |
| *ST6GALNAC4* | 0.472891969817541 | 0.0000118261803671133 | *CRYBG3* | 0.397292607578246 | 0.00128470366847669 |
| *RUFY3* | 0.577427361644559 | 0.0000121079002095892 | *RNF149* | 0.587057918723349 | 0.00131447298586771 |
| *DNAJC4* | 0.638595499382703 | 0.0000121758354567815 | *PPP6R1* | 0.432604326043477 | 0.00135877366601452 |
| *C1QBP* | 0.650338763879079 | 0.0000123687003997618 | *CHML* | 0.35181237483326 | 0.00138053820733957 |
| *WDR1* | 0.657807434782929 | 0.0000125401456788737 | *DACH1* | 0.358370090855423 | 0.00138053820733961 |
| *RPS6KA3* | 0.712766467816528 | 0.0000126130442232299 | *ELL* | 0.331958460024627 | 0.00138053820733961 |

| | | | | | |
|---|---|---|---|---|---|
| *NFKBID* | 0.639935964360132 | 0.0000127387079413441 | *QDPR* | 0.325279304387 | 0.00138053820733961 |
| *MEX3C* | 0.632261725657875 | 0.0000127621892891535 | *CDK2AP1* | 0.3452247147985 | 0.00138053820733965 |
| *PHC3* | 0.662497604846406 | 0.0000131633606067418 | *ARHGAP9* | 0.36549013018494 | 0.00141660713489872 |
| *NOL7* | 0.599061362926657 | 0.0000133463611355912 | *PTBP1* | 0.530399811275527 | 0.00142788655426679 |
| *MTRNR2L8* | 0.790728859921981 | 0.0000134254881588177 | *ADD1* | 0.433672593430969 | 0.00143185191331453 |
| *PPP1R9B* | 0.532461236050811 | 0.0000142191338047969 | *NDUFAF3* | 0.483628329067944 | 0.00145243629446997 |
| *GPAA1* | 0.519434555754571 | 0.0000144642338358672 | *FAM49B* | 0.612591181114925 | 0.00149437103350766 |
| *TKT* | 0.852623611766953 | 0.0000145097607748221 | *ARHGAP17* | 0.448877105143179 | 0.00152735770578872 |
| *ATP6V0E1* | 0.554320418715301 | 0.0000147134511926236 | *MTHFD2L* | 0.382084329591394 | 0.00154876913659006 |
| *FGD2* | 0.415037499278844 | 0.0000153429707035041 | *EIF3K* | 0.566673581463716 | 0.00157703946972607 |
| *WASF2* | 0.660430880325516 | 0.0000153882508528411 | *RILPL2* | 0.50024304291884 | 0.00158767391952112 |
| *SLC38A10* | 0.486343769286148 | 0.0000154188071263722 | *MEF2A* | 0.484758082485809 | 0.00164893276938196 |
| *PPP3R1* | 0.514004539018036 | 0.0000160394055094809 | *NOP10* | 0.447345624384961 | 0.00167053516421586 |
| *ZSCAN16-AS1* | 0.565487136085278 | 0.0000161256488003678 | *DPPA4* | 0.3452247147985 | 0.00172462272393884 |
| *HLA-B* | 0.469909943920364 | 0.0000167704839783024 | *LCNL1* | 0.406625259462644 | 0.00172462272393895 |
| *AP2M1* | 0.509104666187759 | 0.0000168909456335422 | *SNRPE* | 0.505522480245614 | 0.0017346751864785 |
| *DDOST* | 0.592342715388082 | 0.0000170012283903129 | *NUCKS1* | 0.480005993403485 | 0.00176065754151964 |
| *SYS1* | 0.580648150173423 | 0.0000170206176193639 | *YTHDF2* | 0.495380469825 | 0.00178506244774086 |
| *STUB1* | 0.408944252023498 | 0.0000171416605708781 | *LENG8* | 0.335951161025684 | 0.00179446622432197 |
| *EIF3A* | 0.749810022774409 | 0.0000172882807440205 | *H1FX* | 0.747267131660007 | 0.00179751880900593 |
| *NAGK* | 0.543480029721222 | 0.000017950483341047 | *AIF1* | 0.572035298364526 | 0.00181091355350121 |
| *RAB5C* | 0.56579019144631 | 0.0000180722083785335 | *NUFIP2* | 0.553169262321021 | 0.00188345450160014 |
| *RTF2* | 0.540006165676077 | 0.0000180779717754535 | *USP15* | 0.669352040356995 | 0.00191515948527112 |

| | | | | | |
|---|---|---|---|---|---|
| *RPL36* | 0.385231322980974 | 0.0000192831473404238 | *SRRM2* | 0.531268107930052 | 0.00199262963894669 |
| *ARGLU1* | 0.682705746709052 | 0.0000196062486447804 | *DDOST* | 0.474261935391559 | 0.00209404541447789 |
| *PSMA4* | 0.540653985642458 | 0.0000197668755538706 | *PLEKHO1* | 0.472367689904112 | 0.00209594336635219 |
| *CIRBP* | 0.730566861074852 | 0.0000199647049877045 | *CD300A* | 0.334144889051902 | 0.00211555930596552 |
| *GRK3* | 0.436306116032495 | 0.0000200197863118509 | *PABPN1* | 0.510566873639676 | 0.00213195259539303 |
| *CUX2* | 0.429251358498545 | 0.0000200197863118515 | *SUPT4H1* | 0.426821973449471 | 0.00213264741484118 |
| *MYB* | 0.488145047552113 | 0.0000200197863118515 | *PFDN2* | 0.499141328439269 | 0.0022166470913807 |
| *HIGD2A* | 0.694801642265131 | 0.0000205616299797665 | *PBRM1* | 0.502993653934875 | 0.00231864256499824 |
| *TALDO1* | 0.712960131155417 | 0.0000206768861856628 | *NDUFA13* | 0.465974464504069 | 0.00239558602634091 |
| *RNPS1* | 0.57018530540422 | 0.0000212650303540805 | *COX14* | 0.500391858746595 | 0.00252900606222005 |
| *VTI1B* | 0.507520505447837 | 0.0000215634736635362 | *C6orf62* | 0.461527036639491 | 0.00254137046978374 |
| *NONO* | 0.59735550656969 | 0.0000219253286237042 | *CLTA* | 0.47330196899807 | 0.00257842399507163 |
| *STARD7* | 0.481195751466879 | 0.0000220470346011375 | *NDUFC1* | 0.499141328439269 | 0.00260259185518494 |
| *AIF1* | 1.04708208472998 | 0.0000226203038521259 | *ABI2* | 0.390912727064847 | 0.00260974650828207 |
| *ACAP2* | 0.646833609081528 | 0.0000229991120284368 | *CHAF1A* | 0.409689934095475 | 0.00261236009693501 |
| *PTBP1* | 0.583722707534345 | 0.000023106512877952 | *SETBP1* | 0.466651806871155 | 0.00261535138890499 |
| *PRKRA* | 0.459915313082091 | 0.000023567449951178 | *USP24* | 0.459924932042989 | 0.00263875711908013 |
| *MIIP* | 0.413544357238823 | 0.0000239433227264399 | *KHSRP* | 0.387759294025973 | 0.00265108124966822 |
| *ANP32B* | 0.703718099679637 | 0.0000241624976950685 | *KCNK17* | 0.328622747461371 | 0.00268606979561305 |
| *NDUFA3* | 0.609268828113191 | 0.0000243126369106259 | *LTK* | 0.315202231699049 | 0.00268606979561305 |
| *MAPKAPK3* | 0.499671300933141 | 0.000024925477129165 | *MICAL1* | 0.30844486552374 | 0.00268606979561321 |
| *KIAA2013* | 0.54282737418379 | 0.0000258879884983082 | *TNKS2* | 0.426821973449471 | 0.00271218764590807 |
| *SRSF3* | 0.669821300839667 | 0.0000260605841569213 | *COX7B* | 0.556453981673357 | 0.00279777845622619 |
| *PLEKHD1* | 0.432783049468798 | 0.000026095942680698 | *MARCH2* | 0.444099964881307 | 0.00283447640728298 |

| | | | | | |
|---|---|---|---|---|---|
| | | 5 | | | |
| *AL138756.1* | 0.450312974026374 | 0.0000260959426806992 | *PSENEN* | 0.494956334869457 | 0.00285909942929325 |
| *KIAA0930* | 0.466118741222114 | 0.0000262796304269163 | *RPS3* | 0.633107219605548 | 0.00289710789356577 |
| *UBE2L3* | 0.530858871051769 | 0.0000263422166364447 | *DYNLRB1* | 0.475808824608645 | 0.00291301837833182 |
| *UNC119* | 0.534855517431615 | 0.0000270127140763148 | *ZFP91* | 0.567344065344933 | 0.00293854075254836 |
| *C1GALT1* | 0.541034092360739 | 0.0000272299063160207 | *NOL7* | 0.476955793033352 | 0.00298953874391966 |
| *RIPOR1* | 0.514004539018036 | 0.0000272568140082044 | *TRABD* | 0.532287960728913 | 0.00302410293386257 |
| *PSMA3* | 0.565253973249879 | 0.0000274997707957312 | *ATP2A3* | 0.42586922780422 | 0.00307973864199153 |
| *LUC7L3* | 0.720460004765037 | 0.0000295009118657235 | *MEX3C* | 0.43787778489737 | 0.00308111667981758 |
| *PPP6R1* | 0.52949406222332 | 0.0000295422366472931 | *HNRNPDL* | 0.567007827657045 | 0.00318983459413896 |
| *CALR* | 0.74266795353039 | 0.0000314627723086959 | *PSMA4* | 0.417505754869593 | 0.00318993015705053 |
| *TOMM6* | 0.642947921786024 | 0.0000320996018901471 | *SPNS3* | 0.448877105143179 | 0.00321992906946775 |
| *POLR1D* | 0.586548751613108 | 0.0000330145590537597 | *DKC1* | 0.343795059085621 | 0.00323811839464652 |
| *COX7B* | 0.66389766189936 | 0.000033868597214026 | *COPS9* | 0.573941408402198 | 0.00324898685518349 |
| *METAP2* | 0.601024074128987 | 0.0000339120953742378 | *ARID1B* | 0.586374801364476 | 0.00332966289495926 |
| *PROC* | 0.411462045055016 | 0.0000339824194191344 | *CUX2* | 0.301655699861101 | 0.00334888624321987 |
| *CDK2AP1* | 0.400682206301774 | 0.0000339824194191364 | *CHCHD10* | 0.512187008017497 | 0.0034770286109711 |
| *TRIM22* | 0.647610277349386 | 0.0000342887654031621 | *CCS* | 0.421797571086895 | 0.0036109805860759 |
| *OXR1* | 0.462720164760249 | 0.0000343633497583629 | *PHPT1* | 0.492267247341817 | 0.00365903616350209 |
| *DYNLL1* | 0.575211207678827 | 0.0000345979183345305 | *MOB1B* | 0.421797571086895 | 0.00368915589244407 |
| *CPT1A* | 0.574374037667005 | 0.0000354094130690017 | *TMEM165* | 0.418757116766843 | 0.00371507768659776 |
| *SETD2* | 0.517108253115259 | 0.0000354163909413304 | *ABHD14B* | 0.356562355159499 | 0.0037605034496505 |
| *MCL1* | 1.0808495294166 | 0.0000355382061736642 | *EIF1AX* | 0.563681580633979 | 0.00377506554470821 |

| | | | | | |
|---|---|---|---|---|---|
| *PTPN18* | 0.515143083390216 | 0.0000368130336516929 | *SASH3* | 0.412656917230426 | 0.00390107537243516 |
| *SAT1* | 1.44952223069142 | 0.0000368603666067168 | *OIP5-AS1* | 0.471495501100183 | 0.00393918501018774 |
| *SH3BGRL3* | 0.792236096360469 | 0.000036988543789396 | *TYMP* | 0.446959619020547 | 0.00396401981199784 |
| *TBCA* | 0.672909036416289 | 0.0000370449157337461 | *SNHG6* | 0.488625363646638 | 0.00404397600350991 |
| *SRRM2* | 0.663364348663939 | 0.0000373538081466505 | *MZT2B* | 0.597923493950726 | 0.00411169886526898 |
| *ATPAF2* | 0.342461259177341 | 0.0000375883646749535 | *PCF11* | 0.511942610662358 | 0.00413144977830559 |
| *PRPF38B* | 0.739004739439805 | 0.0000386499652894202 | *EMC7* | 0.493988840673666 | 0.00414657963873985 |
| *PARP10* | 0.52949406222332 | 0.0000388752886600848 | *ATP6V1F* | 0.487312031702606 | 0.00417248455669051 |
| *TSPYL2* | 0.88309469566782 | 0.0000396069595680122 | *PTOV1* | 0.287980762964025 | 0.00417255019865402 |
| *NOP10* | 0.547020387426471 | 0.0000400441577532163 | *CCDC189* | 0.321928094887362 | 0.00417255019865414 |
| *TAX1BP3* | 0.532461236050811 | 0.0000404689590356405 | *SLC15A3* | 0.294834434005087 | 0.00417255019865414 |
| *CDC14A* | 0.69346395552962 | 0.0000406627413733453 | *ZNF22* | 0.420770528627201 | 0.00419489842317961 |
| *TMEM248* | 0.574723316216318 | 0.0000420068967088024 | *SNX2* | 0.36549013018494 | 0.00420652792082051 |
| *CAPNS1* | 0.649558549138908 | 0.0000422002303392592 | *ADAM19* | 0.466651806871155 | 0.00420861231708791 |
| *CLTA* | 0.606448309050812 | 0.0000424148350800191 | *IQSEC1* | 0.412391105153968 | 0.00432596044657429 |
| *CD53* | 0.400682206301774 | 0.0000429130018054102 | *UBE2I* | 0.510189155339079 | 0.00435003119588159 |
| *KHDRBS1* | 0.634685272657378 | 0.0000430057520877755 | *PHYKPL* | 0.432604326043477 | 0.00435829143745073 |
| *ATP2A3* | 0.490236078656235 | 0.0000432839963678385 | *NAA10* | 0.442330872028251 | 0.00436547566704068 |
| *ZNF106* | 0.597953514885357 | 0.0000435638149263895 | *ANP32A* | 0.498294317399054 | 0.00437058792596429 |
| *SLC12A2* | 0.389821213670007 | 0.0000442086762982449 | *SNRNP27* | 0.492267247341817 | 0.0045290163714487 |
| *DPPA4* | 0.432783049468798 | 0.0000442086762982475 | *SMIM19* | 0.462836527462892 | 0.00456853991163363 |
| *ADD1* | 0.521492868024751 | 0.0000442163580303816 | *SNF8* | 0.43104381205586 | 0.00463223515231219 |
| *PSMB6* | 0.575660435470371 | 0.000045055278389977 | *FBXW11* | 0.433452720681798 | 0.00464454654725513 |

| | | | | | |
|---|---|---|---|---|---|
| *ISCU* | 0.629298399072445 | 0.0000452531126602985 | *KDM1A* | 0.397198989912278 | 0.00494282072864144 |
| *NEMF* | 0.600790674188887 | 0.0000461157182224904 | *PSMA7* | 0.570107849793412 | 0.00496289270883302 |
| *GNAQ* | 0.586548751613108 | 0.0000469554975155503 | *ABCE1* | 0.460376409136609 | 0.00502813681395764 |
| *IKZF1* | 0.770057034983381 | 0.0000477147488215437 | *KPNA2* | 0.301503337369182 | 0.00515172492256312 |
| *PABPC4* | 0.583601467562771 | 0.0000491133420565897 | *PSME2* | 0.654362456298752 | 0.00515611012613512 |
| *NDUFAF3* | 0.535607185949932 | 0.000050841294390786 | *ARMC5* | 0.311827504918392 | 0.00519545548977935 |
| *TNKS2* | 0.526427759175537 | 0.0000512101913000679 | *FGD2* | 0.305054276322966 | 0.00519545548977966 |
| *PTPN1* | 0.586548751613108 | 0.0000517849803262752 | *FUT7* | 0.335286477742381 | 0.00519545548977966 |
| *HOOK3* | 0.457265734668141 | 0.0000521632255028322 | *RNF144B* | 0.315202231699049 | 0.00519545548977966 |
| *TCF3* | 0.434545658168058 | 0.0000529129305216545 | *VASH2* | 0.335286477742381 | 0.00519545548977966 |
| *RALBP1* | 0.474936760995197 | 0.0000529191628906183 | *TBC1D9* | 0.30844486552374 | 0.00519545548977981 |
| *PPP1CA* | 0.601735155427784 | 0.000054122689645114 | *CNP* | 0.390912727064847 | 0.0052275571590591 |
| *ICOSLG* | 0.538930731380368 | 0.0000542923865063786 | *ELF2* | 0.517000511928754 | 0.00525230126341224 |
| *CD300A* | 0.447814259323569 | 0.0000552077423966587 | *VMA21* | 0.486410769739271 | 0.00538046996326181 |
| *ARPC4* | 0.642141077576425 | 0.0000555956263250351 | *TBCB* | 0.486873988571598 | 0.0054855082299881 |
| *POLR2L* | 0.54737815433243 | 0.0000558447535677088 | *SEC62* | 0.624957920998542 | 0.00555661190587012 |
| *TOR3A* | 0.393450637070698 | 0.0000574561824295603 | *MCFD2* | 0.346997491123965 | 0.00561282092611383 |
| *IFIT2* | 0.450312974026374 | 0.0000574561824295619 | *CYSLTR1* | 0.41571024120798 | 0.00563315787987984 |
| *SLC25A28* | 0.389821213670007 | 0.0000574561824295636 | *RNF5* | 0.465742258638303 | 0.00566679630788463 |
| *TNIP1* | 0.525962598752069 | 0.0000579947766227017 | *PRKRA* | 0.391283266437858 | 0.00570737262150742 |
| *PSMB10* | 0.414782403346628 | 0.0000585761377332205 | *TMED9* | 0.52210247688258 | 0.00571059560964308 |
| *SP3* | 0.589155247427589 | 0.0000586543364804687 | *RAB5C* | 0.436566292195899 | 0.00580856786511131 |
| *NDUFA13* | 0.523354925125914 | 0.0000599840758781144 | *OCIAD1* | 0.488015090458889 | 0.00612618764904355 |

| | | | | | |
|---|---|---|---|---|---|
| *GADD45B* | 0.899863716195762 | 0.0000611129883525532 | *TMEM248* | 0.478047296804644 | 0.00612698078715357 |
| *SETX* | 0.563783860481013 | 0.0000633948799213936 | *MT-ATP6* | 0.447775166264679 | 0.00617891527752446 |
| *MPG* | 0.49118176863309 | 0.0000644721401697619 | *CRTAP* | 0.39747168531823 | 0.0062140613450559 |
| *ARL5A* | 0.505680905917445 | 0.0000662032538352006 | *UBL7* | 0.378257426258818 | 0.00622687504165318 |
| *FIS1* | 0.541694515828847 | 0.0000666172326912677 | *POLE3* | 0.459601058421494 | 0.00625275282921415 |
| *MAP7D1* | 0.507891807738109 | 0.0000669320657442477 | *GLCE* | 0.375076179014138 | 0.00638174342280257 |
| *CCS* | 0.464648698186829 | 0.0000672585691490392 | *PPP1R9B* | 0.371887901383091 | 0.00642381504776697 |
| *ZFP36* | 1.07931755412452 | 0.0000685105239167656 | *LRRK1* | 0.301655699861101 | 0.00646501283276454 |
| *RER1* | 0.415917757966401 | 0.0000687163604943476 | *NREP* | 0.291411668364298 | 0.00646501283276474 |
| *PSMA7* | 0.689251704242705 | 0.0000692952219819659 | *PDK2* | 0.291411668364298 | 0.00646501283276474 |
| *YME1L1* | 0.605594507303654 | 0.0000706493498538459 | *RBM47* | 0.294834434005087 | 0.00646501283276474 |
| *MT-ATP8* | 0.840357692233899 | 0.0000708716833553191 | *MILR1* | 0.315202231699049 | 0.00646501283276492 |
| *TRAPPC1* | 0.521302726997669 | 0.0000718712502367184 | *TRAPPC12* | 0.294834434005087 | 0.00646501283276492 |
| *GLG1* | 0.592108276955008 | 0.0000729265635468819 | *PIK3AP1* | 0.457583194244928 | 0.00647874904317279 |
| *EIF3D* | 0.515320403610311 | 0.0000731832217926759 | *CCDC18-AS1* | 0.533050887791055 | 0.0067563258277968 |
| *TMEM165* | 0.501007198489163 | 0.0000738234093825548 | *RTN4* | 0.514966295600951 | 0.0067885934405413 |
| *ANAPC16* | 0.646283417106047 | 0.0000744032401475031 | *SLC66A2* | 0.409419538622678 | 0.00685101429185204 |
| *LTK* | 0.382534859591515 | 0.000074601260801093 | *DST* | 0.40345798038396 | 0.00708209402336372 |
| *LRRK1* | 0.397070952749395 | 0.0000746012608010952 | *ZBTB18* | 0.428225885532903 | 0.00712663495690143 |
| *MYD88* | 0.418604114299036 | 0.0000746012608010973 | *BOD1L1* | 0.511912971623397 | 0.00728434921690561 |
| *PSME2* | 0.699759362061099 | 0.0000746825861287468 | *OXR1* | 0.394059282381711 | 0.00751319893917191 |
| *ADI1* | 0.586548751613108 | 0.0000747335014039694 | *ANP32B* | 0.582027873114633 | 0.00759818986863595 |
| *CUEDC1* | 0.489687212360519 | 0.0000767599631431332 | *NPM1* | 0.575372906336088 | 0.00760965189817794 |

| | | | | | |
|---|---|---|---|---|---|
| *ZNF511* | 0.445606143896402 | 0.0000779121261033862 | *AC005224.2* | 0.553034336434471 | 0.00778201620845144 |
| *NDUFB4* | 0.67667039475615 | 0.0000792400119585112 | *CLIC1* | 0.505739030673692 | 0.00788637109269545 |
| *NFATC2IP* | 0.637837980309962 | 0.0000795015711369972 | *MRPS21* | 0.506835720229269 | 0.00788934263628128 |
| *MX1* | 0.663274542255936 | 0.0000815005019572939 | *MAGED1* | 0.368692562223284 | 0.00790680983282098 |
| *FIP1L1* | 0.583722707534345 | 0.0000818466323556606 | *SLC25A28* | 0.311827504918392 | 0.00803973421650852 |
| *ARHGAP9* | 0.437219792466383 | 0.0000829165093289484 | *EIF4EBP3* | 0.291411668364298 | 0.008039734216509 |
| *MTMR10* | 0.342461259177341 | 0.0000871146967627198 | *FBH1* | 0.274174963438994 | 0.00803973421650948 |
| *NENF* | 0.520275532193942 | 0.0000877010635978363 | *SLC2A8* | 0.298249098417852 | 0.00803973421650948 |
| *HINT2* | 0.504779195870774 | 0.0000878906371185381 | *EDEM1* | 0.409597117481368 | 0.00831915311221314 |
| *PGLS* | 0.605914076480039 | 0.0000885740357420036 | *IL10RA* | 0.53311364231121 | 0.00848991975644219 |
| *RBM4* | 0.576075420383599 | 0.0000904250410043516 | *RERE* | 0.38866097845588 | 0.00864594009065402 |
| *FBXW11* | 0.518279475596079 | 0.0000939227808849158 | *ARF5* | 0.400378587451989 | 0.00868310983272325 |
| *AHI1* | 0.511909950720352 | 0.0000946828042768524 | *NUMA1* | 0.327044864864793 | 0.0087773674154708 |
| *H1FX* | 0.804140186685735 | 0.0000948028880761213 | *CAP1* | 0.541812902825403 | 0.00889193192056745 |
| *EIF5* | 0.713439384071789 | 0.0000948235974904735 | *RAB18* | 0.409507025056766 | 0.00891692908734045 |
| *PPP1R2* | 0.606357232693288 | 0.0000952761333260599 | *NAP1L1* | 0.532624001501957 | 0.00901163209587517 |
| *CLIC3* | 0.646503155898778 | 0.0000963873201808449 | *SLC1A5* | 0.400378587451989 | 0.00923288229432892 |
| *ARMC5* | 0.393450637070698 | 0.0000967697894683014 | *PPP3R1* | 0.381762909284642 | 0.00923365762557867 |
| *MICAL1* | 0.400682206301774 | 0.000096769789468307 | *FAAP20* | 0.403457980383961 | 0.00926502546835116 |
| *PSEN1* | 0.389821213670007 | 0.000096769789468307 | *CYB5R3* | 0.387994862996156 | 0.00928541520721927 |
| *MIA3* | 0.614893377307069 | 0.000098490990426829 | *HNRNPUL1* | 0.485792913181267 | 0.00928775221424081 |
| *SRSF4* | 0.601735155427784 | 0.00010023007210732 | *PIGT* | 0.388445409528936 | 0.00933989745385369 |
| *SPNS3* | 0.497739484655252 | 0.000101133647063928 | *XRCC6* | 0.485071745937613 | 0.00937272409619371 |
| *CNOT7* | 0.528132065387786 | 0.000101275849110269 | *MYL6B* | 0.433452720681798 | 0.00942000868014893 |
| *COPS9* | 0.586548751613108 | 0.000102427748083651 | *TMCO1* | 0.523543490860439 | 0.0095016521144381 |

| | | | | | |
|---|---|---|---|---|---|
| *TMEM19* | 0.497739484655252 | 0.000103235109622811 | *CUEDC1* | 0.390912727064847 | 0.00950256151474351 |
| *BTK* | 0.431767236431337 | 0.000103277865019221 | *MAP2K6* | 0.378257426258818 | 0.00951205414797907 |
| *TRA2B* | 0.609268828113191 | 0.000103941926391967 | *CDC14A* | 0.498052912664605 | 0.00953633677377728 |
| *CNDP2* | 0.544113485292711 | 0.000105722370498417 | *ACBD5* | 0.384598953132645 | 0.00970771060787348 |
| *PTRHD1* | 0.566370869675477 | 0.00010724707263119 | *LASP1* | 0.443535419759296 | 0.00983152923511679 |
| *VMA21* | 0.547756268616049 | 0.00010895093201253 | *ARL6IP4* | 0.559577182301243 | 0.00986154701695416 |
| *FBRS* | 0.568514828782846 | 0.000108969087999833 | *RNASEH2B* | 0.429349513515984 | 0.00995638649514041 |
| *ZNF22* | 0.53633730565934 | 0.000112213847432977 | *CCDC112* | 0.291411668364298 | 0.00999179183522215 |
| *BRD4* | 0.636560519780931 | 0.000112388174238192 | *YBEY* | 0.287980762964025 | 0.00999179183522245 |
| *ACAA1* | 0.515098228351399 | 0.000115221573262821 | *THBD* | 0.315202231699049 | 0.00999179183522274 |
| *METTL7A* | 0.489687212360519 | 0.000122510427807393 | *ZNF787* | 0.301655699861101 | 0.00999179183522274 |
| *SNX2* | 0.466118741222114 | 0.000124310431265592 | *B3GALNT2* | 0.298249098417852 | 0.00999179183522303 |
| *PARVG* | 0.569740463926554 | 0.000125359666254844 | *NUDT17* | 0.294834434005087 | 0.00999179183522333 |
| *MRPS10* | 0.382534859591515 | 0.000125406981639418 | *SLC12A9* | 0.274174963438994 | 0.00999179183522363 |
| *SCAF8* | 0.378877835983425 | 0.000125406981639418 | *IKZF1* | 0.606991374469093 | 0.00999884610516042 |
| *MYCL* | 0.400682206301774 | 0.000125406981639426 | *CMPK1* | 0.464270222205515 | 0.0100068101752305 |
| *NAP1L1* | 0.682896147532975 | 0.000125619608235207 | *IFNAR1* | 0.403457980383961 | 0.0101423547611096 |
| *AEBP1* | 0.515098228351399 | 0.000125820880064458 | *YWHAE* | 0.444680643110474 | 0.0102195539806966 |
| *ZBTB18* | 0.51615942372171 | 0.00012590109714052 | *TBCC* | 0.421016351768265 | 0.0104024887192996 |
| *SRPRA* | 0.592034290696837 | 0.000129430695757758 | *LSM8* | 0.327569672088073 | 0.0104477870287836 |
| *CYSLTR1* | 0.510766164951492 | 0.000131991813151315 | *RIPOR1* | 0.406530814433584 | 0.010457163848612 |
| *CRYBG3* | 0.44102509366093 | 0.000133510671748306 | *SELPLG* | 0.542932399305019 | 0.0105660432064794 |
| *UFM1* | 0.547868283449048 | 0.00014178689210509 | *TMEM230* | 0.495380469825 | 0.0105811411378165 |
| *FAM214A* | 0.622792272212937 | 0.000143286429783421 | *ZNF106* | 0.508327098237916 | 0.0107230421807049 |
| *ANAPC5* | 0.558036063870695 | 0.000144301879603497 | *MIA3* | 0.486873988571598 | 0.010912393985259 |
| *SYNCRIP* | 0.608993898137916 | 0.000144666293411929 | *RAD23A* | 0.475056777431752 | 0.0110873014681454 |
| *CDV3* | 0.674943867558822 | 0.000144849665868335 | *SNRPD1* | 0.361637804689334 | 0.0111651328319847 |
| *GLO1* | 0.566649194175403 | 0.000146081291312599 | *ERGIC3* | 0.459601058421494 | 0.0112466869210819 |
| *INPP4A* | 0.532553861972378 | 0.000148061890837068 | *FUS* | 0.599934919917383 | 0.0116687548807349 |
| *BCLAF1* | 0.723099267484578 | 0.000156746615804084 | *TMEM19* | 0.427859323874096 | 0.0116694245600581 |
| *IQGAP2* | 0.632645166202562 | 0.000159641983855418 | *LYSMD2* | 0.369727123071411 | 0.0119475933901245 |
| *ASIP* | 0.400682206301774 | 0.000162366327129109 | *FIP1L1* | 0.431418061724848 | 0.0119713669198138 |
| *FEM1B* | 0.407877707705978 | 0.000162366327129109 | *AC011893.1* | 0.298249098417852 | 0.0124101580534375 |
| *MILR1* | 0.378877835983425 | 0.000162366327129109 | *BCOR* | 0.284541678997254 | 0.0124101580534375 |
| *CHPF2* | 0.345540652109313 | 0.000162366327129114 | *NUP42* | 0.274174963438994 | 0.0124101580534375 |
| *PLEKHM2* | 0.36415633027666 | 0.000162366327129114 | *REPS1* | 0.291411668364298 | 0.0124101580534375 |
| *EIF1AX* | 0.622731087919754 | 0.000162911157709101 | *TFPT* | 0.281094377378785 | 0.0124101580534375 |
| *MFSD12* | 0.459313563317 | 0.000165494902313261 | *IMPACT* | 0.294834434005087 | 0.0124101580534379 |

| | | | | | |
|---|---|---|---|---|---|
| *DNAJB11* | 0.548309947325416 | 0.000166625083477699 | *PPP1R16B* | 0.294834434005087 | 0.0124101580534379 |
| *TRIR* | 0.617754619450918 | 0.000167721117074841 | *RAB34* | 0.277638818742566 | 0.0124101580534379 |
| *RNF114* | 0.527292166056333 | 0.000168190507012846 | *DEF8* | 0.394200012522865 | 0.0129690919674767 |
| *ATP5MC1* | 0.518107695511231 | 0.000169433670883017 | *LSM10* | 0.433022437926099 | 0.0129888798942035 |
| *ABHD14B* | 0.324129436405083 | 0.000170278501057395 | *CTDNEP1* | 0.484681775657934 | 0.0129922475287456 |
| *TCF25* | 0.544728575918481 | 0.000171529067331219 | *FAM214A* | 0.464270222205515 | 0.0130277674034507 |
| *UHRF1BP1L* | 0.445606143896402 | 0.000174404776247134 | *ZCRB1* | 0.400378587451989 | 0.0130797388151789 |
| *STK24* | 0.471327853824456 | 0.000175733293056496 | *ZBTB7A* | 0.458175150093029 | 0.0130957502974966 |
| *EMB* | 0.543815766624575 | 0.000182646964330345 | *PSMB1* | 0.467482892319257 | 0.0135916562699792 |
| *ADAM19* | 0.474655871542211 | 0.000182649649768113 | *CCDC85B* | 0.458599534576422 | 0.0138432465137158 |
| *NFX1* | 0.396099222779414 | 0.000187812539440172 | *STARD7* | 0.412140223483761 | 0.0138474509402586 |
| *PPP1R15A* | 0.805666040916695 | 0.000189922575743234 | *SMDT1* | 0.470572176242497 | 0.0139118517294654 |
| *MCFD2* | 0.44102509366093 | 0.000191464863674409 | *TRIM22* | 0.52310122738345 | 0.0140062096187402 |
| *PI4KA* | 0.510997718605931 | 0.000196365887983673 | *SNRPB* | 0.524099286946526 | 0.0140636633482364 |
| *PRDX1* | 0.586548751613108 | 0.000197549968794222 | *SREK1IP1* | 0.421269256500421 | 0.0141861622499799 |
| *KARS* | 0.456602579041255 | 0.000198237392260345 | *RTRAF* | 0.469800475212657 | 0.0142378458220373 |
| *UFC1* | 0.598621583913683 | 0.000204061007122658 | *ACYP2* | 0.378257426258818 | 0.0144589828828644 |
| *UBTF* | 0.513019716191143 | 0.000204199043269373 | *ZSCAN16-AS1* | 0.456245394743396 | 0.0146761408382443 |
| *SETBP1* | 0.536471879143966 | 0.000204280744326234 | *ISCU* | 0.510566873639676 | 0.0146781202294635 |
| *CMPK1* | 0.560912824626258 | 0.00020449587439363 | *VIPR2* | 0.40345798038396 | 0.0149446820596014 |
| *COL26A1* | 0.360452361513241 | 0.000210022863585628 | *DCPS* | 0.36228057370905 | 0.0150168306617197 |
| *LRP8* | 0.411462045055016 | 0.000210022863585628 | *OTULIN* | 0.452193646927643 | 0.0151579320527178 |
| *LILRB2* | 0.418604114299036 | 0.00021002286358564 | *ITPR1* | 0.42451959591179 | 0.0153997036501799 |
| *PNOC* | 0.446823987946494 | 0.00021002286358564 | *AL138756.1* | 0.291411668364298 | 0.0154044564886327 |
| *CYSTM1* | 0.442158842277933 | 0.000216938884910409 | *ARMC10* | 0.267222202797227 | 0.0154044564886332 |
| *EMC10* | 0.513150673849371 | 0.000217903133821373 | *SLC12A3* | 0.298249098417852 | 0.0154044564886332 |
| *EPN1* | 0.445606143896402 | 0.000218996234312251 | *NUB1* | 0.410888185859449 | 0.0157671742972393 |
| *SIRT2* | 0.38906167203948 | 0.000220144645703832 | *BNIP2* | 0.505457693886356 | 0.0158877409386173 |
| *TCEA1* | 0.674904626033954 | 0.000221133159683601 | *CLTB* | 0.421016351768265 | 0.0159706268815138 |
| *EIF4G3* | 0.490848335192522 | 0.000226998017541447 | *UPF3A* | 0.386418077085496 | 0.0162923177827824 |
| *LILRB1* | 0.417008212542395 | 0.000228329729797645 | *GHITM* | 0.490609929479043 | 0.0163523618676896 |
| *OCIAD1* | 0.557224672133164 | 0.000230109301013189 | *COPA* | 0.48309409649717 | 0.016650502569162 |
| *ZNF652* | 0.603389740990421 | 0.000231123985044874 | *PRDX5* | 0.527125316407773 | 0.0166852778905041 |
| *PSENEN* | 0.51013563609394 | 0.000231240497930726 | *ESD* | 0.441774911935063 | 0.0167244250621648 |
| *SREK1IP1* | 0.541034092360739 | 0.000231784803706845 | *ZNF800* | 0.403457980383961 | 0.0169801097442991 |
| *ERO1B* | 0.548848042427884 | 0.000235148421281842 | *CDV3* | 0.553874822937059 | 0.0173634911920993 |
| *DDX21* | 0.787921593868035 | 0.000235491409550906 | *GNB2* | 0.375712989901056 | 0.0174312713277597 |
| *SNRPB* | 0.564689317649368 | 0.000237023722007188 | *PHF14* | 0.375918956785679 | 0.0177309591597437 |

| | | | | | |
|---|---|---|---|---|---|
| *UBE2G2* | 0.528948484990179 | 0.000238002463950735 | *PRMT9* | 0.439906951638626 | 0.018169785733531 |
| *TMCO1* | 0.614964433777194 | 0.00024962695357278 | *RAB14* | 0.418644384198637 | 0.0183115840726488 |
| *RBM23* | 0.476459299901452 | 0.000249913154719983 | *ATAD2B* | 0.382084329591394 | 0.0186921299606643 |
| *SIN3A* | 0.262441907193779 | 0.000251869849803444 | *CCT8* | 0.474112533947746 | 0.0188439539042541 |
| *NRIP1* | 0.514096957264817 | 0.000255127539031147 | *SLC44A2* | 0.436945635693281 | 0.0190618579256063 |
| *CELF1* | 0.651889505280426 | 0.000257425302158029 | *GCA* | 0.267222202797227 | 0.0191096830196191 |
| *FAM53C* | 0.559803896380579 | 0.000260318587967584 | *DAPK1* | 0.267222202797227 | 0.0191096830196203 |
| *CCDC18-AS1* | 0.572925467749133 | 0.000260359383005888 | *GIT1* | 0.277638818742566 | 0.0191096830196203 |
| *SELPLG* | 0.610751099938733 | 0.00026323975564161 | *PLEKHM2* | 0.274174963438994 | 0.0191096830196208 |
| *RAB18* | 0.502302623046311 | 0.00026642882130109 | *TUBB6* | 0.267222202797227 | 0.0191096830196208 |
| *TMED9* | 0.589122695276934 | 0.000268403179900452 | *WDR83OS* | 0.491994654985618 | 0.0192624382083672 |
| *BRI3BP* | 0.469509330422888 | 0.000269065185304311 | *ADA2* | 0.520020665495528 | 0.0194192075981269 |
| *ZNF787* | 0.371535860642257 | 0.00027141731576216 | *PSMB6* | 0.459311215117478 | 0.0197312298463462 |
| *ABCA7* | 0.353015772671459 | 0.000271417315762168 | *USP7* | 0.447345624384961 | 0.0200446113197466 |
| *HIP1* | 0.404284442981969 | 0.000271417315762177 | *SNRPC* | 0.43787778489737 | 0.0209226412470305 |
| *FAM204A* | 0.501052531331501 | 0.000272062661304644 | *SETD2* | 0.387094534175073 | 0.0211283697190129 |
| *WDFY2* | 0.457265734668141 | 0.000273394307100439 | *C4orf3* | 0.484987865880559 | 0.0212746171866207 |
| *LINC-PINT* | 0.68149715566837 | 0.000276541164554851 | *TMEM50A* | 0.447284221348587 | 0.0212966779157475 |
| *FBXW5* | 0.470845530426611 | 0.000277940297799102 | *FBRS* | 0.43886331651516 | 0.0213175415373412 |
| *ENSA* | 0.509586769798223 | 0.000279269175834056 | *TNNI2* | 0.400331879397241 | 0.0214310919705677 |
| *SF3B2* | 0.563465138500066 | 0.000291258824623704 | *KLF16* | 0.418757116766843 | 0.0214322983108214 |
| *UBL7* | 0.414279809702496 | 0.000296154139910813 | *EDF1* | 0.514489292772704 | 0.0215852157280943 |
| *MAF1* | 0.435148161848098 | 0.000298373830859682 | *RHOF* | 0.41470712151692 | 0.0217303783360783 |
| *SEC13* | 0.457938476597277 | 0.000302137471538636 | *SMIM7* | 0.438363595332162 | 0.0228425611029532 |
| *TAX1BP1* | 0.646191698040509 | 0.000314443964956491 | *SLC30A5* | 0.342871827522922 | 0.0231726762641233 |
| *TMEM107* | 0.499416159887133 | 0.000314609557696814 | *LTV1* | 0.465946667179555 | 0.023245483277191 |
| *ME2* | 0.38075829761379 | 0.000321559358306928 | *TRAPPC1* | 0.425372756812401 | 0.0235852749339681 |
| *TTC17* | 0.454758879057567 | 0.000324591536650362 | *GPX7* | 0.263733216717347 | 0.0236919905687881 |
| *AKAP8* | 0.524621002469906 | 0.000327306751728376 | *ZNF852* | 0.274174963438994 | 0.0236919905687881 |
| *PLIN3* | 0.374882736869718 | 0.000327495890622768 | *PALD1* | 0.287980762964025 | 0.0236919905687887 |
| *DKC1* | 0.392584738603177 | 0.000331818829331328 | *FEM1B* | 0.270702771532189 | 0.0236919905687895 |
| *DDAH2* | 0.501659854026595 | 0.00033195280821135 | *PRDX3* | 0.333068652492563 | 0.023833629379634 |
| *LEPROT* | 0.473278052656772 | 0.000343826208012421 | *RNPS1* | 0.424991141933602 | 0.0240546416431079 |
| *AHCY* | 0.459313563317 | 0.00034644121889945 | *TMBIM4* | 0.494223444307143 | 0.0245260380351509 |
| *CDH23* | 0.378877835983425 | 0.000350439367925155 | *AHI1* | 0.403457980383961 | 0.0247722169202388 |
| *AP1S3* | 0.382534859591515 | 0.000350439367925165 | *SGSM3* | 0.384882251128821 | 0.0250799549661575 |
| *CLMN* | 0.393450637070698 | 0.000350439367925165 | *BLOC1S1* | 0.507794640198696 | 0.0252729298221708 |
| *TP53I11* | 0.36415633027666 | 0.000350439367925165 | *MT-CYB* | 0.337218056872685 | 0.0258116430748493 |

| | | | | | |
|---|---|---|---|---|---|
| *NSMCE4A* | 0.353015772671459 | 0.000350439367925176 | *BRI3BP* | 0.375076179014138 | 0.0263702288514128 |
| *PTOV1* | 0.360452361513241 | 0.000350439367925176 | *RRAGC* | 0.421797571086895 | 0.0269419756220365 |
| *KCNK17* | 0.400682206301774 | 0.000350439367925186 | *SERINC1* | 0.455427970682819 | 0.0273428532310669 |
| *TAF10* | 0.611466383817437 | 0.000355579241353438 | *LILRB1* | 0.321176756423602 | 0.027576065416829 |
| *CIR1* | 0.62246832586805 | 0.000357022959221806 | *KIAA0930* | 0.359063861025507 | 0.0278243245814051 |
| *GINM1* | 0.487652996032716 | 0.00035703126075902 | *GNAQ* | 0.420298969761274 | 0.0279687283111026 |
| *LINC00909* | 0.438703283641714 | 0.000360050511599983 | *MAN1A1* | 0.43886331651516 | 0.0284156681798279 |
| *NAPA* | 0.569740463926554 | 0.000366479697322563 | *TMEM208* | 0.366621212203589 | 0.0293292873646084 |
| *PXK* | 0.435239428338142 | 0.000368859048626 | *FAM49A* | 0.281094377378785 | 0.0293557741711688 |
| *TUBGCP3* | 0.442158842277933 | 0.000369935456386329 | *SMIM13* | 0.274174963438994 | 0.0293557741711688 |
| *SMARCC1* | 0.518189958949081 | 0.000370276217149847 | *ENPP2* | 0.270702771532189 | 0.0293557741711723 |
| *NUB1* | 0.564025734986968 | 0.000378550095838965 | *TM2D1* | 0.382396364856131 | 0.0296598561457455 |
| *CRTAP* | 0.484450563994372 | 0.000384946895071419 | *EIF6* | 0.432604326043477 | 0.0301715224801435 |
| *NUS1* | 0.527781343363234 | 0.000394645413345556 | *POLR2I* | 0.461560934947529 | 0.0302302319309233 |
| *DEF8* | 0.4679921412612 | 0.000395079628661957 | *VAV3* | 0.280601232598427 | 0.0309356226544439 |
| *SNRPD2* | 0.643379974209784 | 0.00039518166064352 | *CSDE1* | 0.548574688243771 | 0.0314020497993509 |
| *TLE3* | 0.417008212542395 | 0.000395500042038014 | *PSMA3* | 0.424443005513297 | 0.0314932370390278 |
| *NAGA* | 0.43656482580471 | 0.000402359644433317 | *VPS29* | 0.415187272966068 | 0.0325007380075665 |
| *GTF2A2* | 0.523354925125914 | 0.000404916260917116 | *RAB5IF* | 0.286920191688673 | 0.0336266629226996 |
| *PAG1* | 0.608081913162749 | 0.000407314014583432 | *SMU1* | 0.422823305250892 | 0.034542579934369 |
| *IFNAR1* | 0.518107695511231 | 0.000420795373798209 | *PBX2* | 0.346124805318009 | 0.0347666781907794 |
| *BLOC1S1* | 0.550022875587994 | 0.000422150640416134 | *PGP* | 0.339611498359335 | 0.0349628074788227 |
| *PSMB4* | 0.533368713753933 | 0.00042394849207509 | *SIRT2* | 0.294885458604675 | 0.0356048554321852 |
| *OTULIN* | 0.642831227210138 | 0.000424748094042124 | *LPGAT1* | 0.447140419201357 | 0.0356574434146388 |
| *ARHGDIA* | 0.617021103837378 | 0.000428440669720956 | *AC008764.7* | 0.263733216717347 | 0.0363523447916899 |
| *AP2B1* | 0.533368713753933 | 0.000429340926673334 | *OAS1* | 0.256729828979585 | 0.0363523447916899 |
| *ABCE1* | 0.542336046280783 | 0.00043815800894002 | *TM9SF1* | 0.26023577248112 | 0.0363523447916899 |
| *FEZ2* | 0.503559386942076 | 0.000444288257876014 | *LRP8* | 0.270702771532189 | 0.0363523447916911 |
| *LPGAT1* | 0.565525501042689 | 0.00044611619943524 | *PLAUR* | 0.263733216717347 | 0.0363523447916921 |
| *SNHG6* | 0.608659453218856 | 0.000449093616219622 | *AEBP1* | 0.462836527462892 | 0.036570344733523 |
| *TMEM230* | 0.519951002884084 | 0.000450114809689204 | *ARFGAP3* | 0.458341887457085 | 0.0366989834889275 |
| *GIT1* | 0.353015772671459 | 0.000452060509785987 | *POLR2J3* | 0.537875016404936 | 0.0368844627801688 |
| *FAM49A* | 0.356738858714177 | 0.000452060509786001 | *AP2M1* | 0.418460143723207 | 0.0368968833270543 |
| *AKR7A2* | 0.377639559640282 | 0.000455764490100657 | *NAGK* | 0.397383465974276 | 0.0371228849302905 |
| *CLPX* | 0.442158842277933 | 0.000461887997995462 | *ATOX1* | 0.395299456996797 | 0.0371984000334189 |
| *TCL1A* | 1.22333847029161 | 0.000490933054836247 | *CDC123* | 0.271668107828419 | 0.0374571810748499 |
| *RPL14* | 0.521875522275375 | 0.000500210064386343 | *UBE2G2* | 0.431418061724848 | 0.0386489718992273 |
| *PHPT1* | 0.501052531331501 | 0.000502029282257845 | *SDHC* | 0.372364211282392 | 0.0388779581841578 |

| | | | | | |
|---|---|---|---|---|---|
| *EIF1B* | 0.601371091318631 | 0.000504859591453881 | *ARHGAP4* | 0.458846550250493 | 0.0392536251962848 |
| *DNM2* | 0.535524748588641 | 0.00051164087108955 | *KHDRBS1* | 0.538617563665596 | 0.0398275767865703 |
| *DST* | 0.526885830314768 | 0.000537139013556738 | *SEC61A1* | 0.434606678954625 | 0.0407128497711047 |
| *YY1* | 0.612083843720245 | 0.000539064273705439 | *EIF2S2* | 0.515219956268135 | 0.0424478152307175 |
| *CMTM6* | 0.586548751613108 | 0.000543138986691278 | *ICAM1* | 0.415187272966068 | 0.0429080167236294 |
| *TNPO1* | 0.455436837265949 | 0.000554388962823108 | *TTC7A* | 0.342871827522922 | 0.043169975081765 |
| *CD99* | 0.506427885703936 | 0.000561905160487597 | *BZW1* | 0.306398037082254 | 0.0441127390726168 |
| *TMED5* | 0.560912824626258 | 0.000567679721158084 | *MCC* | 0.263733216717347 | 0.0449905436496244 |
| *RAB14* | 0.526806667921218 | 0.000568587755043862 | *RMC1* | 0.270702771532189 | 0.0449905436496244 |
| *POLE3* | 0.551783333452431 | 0.000571848036761781 | *CCR2* | 0.287980762964025 | 0.0449905436496257 |
| *CTNNA1* | 0.345540652109313 | 0.000582629634728351 | *GAB1* | 0.256729828979585 | 0.0449905436496257 |
| *BACH1* | 0.382534859591515 | 0.000582629634728369 | *SLC2A6* | 0.281094377378785 | 0.0449905436496257 |
| *MBOAT7* | 0.367850813834401 | 0.000582629634728369 | *CLCN5* | 0.270702771532189 | 0.0449905436496297 |
| *REPS1* | 0.338026598452006 | 0.000582629634728369 | *TLE3* | 0.271480854269731 | 0.0453646930966167 |
| *SLC12A9* | 0.330473204020371 | 0.000582629634728402 | *EIF4EBP1* | 0.369217655965528 | 0.0484000695770726 |
| *MRPL14* | 0.330473204020371 | 0.00058262963472842 | *PAK2* | 0.466754510503548 | 0.0489529139129419 |
| *EIF2S3* | 0.582336500326193 | 0.000596495922603233 | | | |
| *NBDY* | 0.578169085997135 | 0.000598019537540686 | | | |
| *SEC62* | 0.653662947471645 | 0.000602172540941948 | | | |
| *PRRC2B* | 0.441118312091544 | 0.000616278176271502 | | | |
| *ZDHHC4* | 0.419651442868363 | 0.000618381886455257 | | | |
| *PRKDC* | 0.488368357674204 | 0.000619326981295154 | | | |
| *DNAJC1* | 0.512089759412663 | 0.000622237785849056 | | | |
| *PCBP1* | 0.638950449572927 | 0.000622585965404933 | | | |
| *CACUL1* | 0.371316121849526 | 0.000644130389973652 | | | |
| *WDR43* | 0.591757042615869 | 0.000657890236788019 | | | |
| *RASGEF1B* | 0.645083962582862 | 0.000662587128523687 | | | |
| *AKAP9* | 0.734694369388644 | 0.00067139901388116 | | | |
| *LAMP1* | 0.554066440497817 | 0.000694700536481567 | | | |
| *PLEKHB2* | 0.423911072797083 | 0.000697514167109428 | | | |
| *BTAF1* | 0.507891807738109 | 0.000699155113897287 | | | |
| *TUBB* | 0.625648571302048 | 0.000701782654863688 | | | |
| *POLR2G* | 0.443275047394262 | 0.000703621968278795 | | | |
| *ZNF655* | 0.463692003827574 | 0.000707948709506294 | | | |
| *UBE2E3* | 0.583683417918718 | 0.000715691749699834 | | | |
| *NUMA1* | 0.416623750170795 | 0.000743561175676165 | | | |
| *TMEM8B* | 0.375211518785656 | 0.000750248014067358 | | | |
| *BCOR* | 0.353015772671459 | 0.000750248014067381 | | | |

| | | |
|---|---|---|
| *HHIP-AS1* | 0.356738858714177 | 0.000750248014067402 |
| *SLC35F3* | 0.356738858714177 | 0.000750248014067402 |
| *CLCN5* | 0.371535860642257 | 0.000750248014067424 |
| *BRD2* | 0.58442245050472 | 0.000754819973927575 |
| *SLC66A2* | 0.448746502312693 | 0.0007588602425949 |
| *ATRAID* | 0.458483712824813 | 0.00077324585629244 |
| *DCPS* | 0.424797681616496 | 0.000784265925836984 |
| *PSMD9* | 0.526427759175537 | 0.000795456578380502 |
| *ZCRB1* | 0.477976229833822 | 0.000798477605461098 |
| *AC005224.2* | 0.697355537499042 | 0.000808300134833422 |
| *NCOA1* | 0.49801207701145 | 0.000822073918231897 |
| *POM121* | 0.389435738725018 | 0.000826142879972809 |
| *SUSD6* | 0.441025093660931 | 0.000828035386722362 |
| *MED30* | 0.304363245434402 | 0.000830257504760986 |
| *MLXIP* | 0.495818975437534 | 0.000839246491427728 |
| *GNB2* | 0.589294128351925 | 0.000853232664774146 |
| *RBM5* | 0.558803761130204 | 0.000854583219366118 |
| *AKR1A1* | 0.468650498147765 | 0.000868990612792027 |
| *SCFD1* | 0.49403661282357 | 0.000883995779954373 |
| *MTHFD2L* | 0.426440608531002 | 0.000894498339528951 |
| *SMG7* | 0.434203827804916 | 0.00089684575706651 |
| *FMNL1* | 0.463300976574635 | 0.000900093646778591 |
| *CTNNBL1* | 0.463692003827574 | 0.000925330435013421 |
| *LSM10* | 0.528650367904906 | 0.000946139342034725 |
| *MALAT1* | 0.505851139364285 | 0.000951574798057288 |
| *UQCRC2* | 0.507891807738109 | 0.00096405258580204 |
| *ATP6V0A1* | 0.341788517248205 | 0.00096524460213082 |
| *P2RX1* | 0.341788517248205 | 0.00096524460213082 |
| *PALD1* | 0.345540652109313 | 0.00096524460213082 |
| *ZDHHC14* | 0.322880054698141 | 0.00096524460213082 |
| *COX5B* | 0.555396231288169 | 0.000968028532381136 |
| *SRSF2* | 0.634356142495238 | 0.000972460615226542 |
| *PAPOLA* | 0.53165050546421 | 0.00099443338790754 |
| *CNP* | 0.407225052168546 | 0.00102959781684755 |
| *THOC2* | 0.566084649053392 | 0.00103713355896904 |
| *AC016831.5* | 0.504389646782167 | 0.00104945963836077 |
| *KDM2A* | 0.527981132182554 | 0.0010535354400477 |
| *AL355075.4* | 0.614382959809833 | 0.00105941095421859 |

| | | |
|---|---|---|
| *UBE2V1* | 0.604927280927962 | 0.00110536566031039 |
| *PRPF40A* | 0.602668416976385 | 0.00110906940739969 |
| *LTV1* | 0.521230413269132 | 0.00110994926916313 |
| *SMIM19* | 0.439923837894625 | 0.00111906129084655 |
| *CALHM6* | 0.49801207701145 | 0.0011213226283493 |
| *CCT8* | 0.572555479533608 | 0.00114010286503511 |
| *PGGT1B* | 0.481241007518482 | 0.00114651234709989 |
| *ATOX1* | 0.45904427819426 | 0.00117221483131592 |
| *RHEB* | 0.500392107863393 | 0.00119467920296092 |
| *EIF4G1* | 0.495862009488482 | 0.00124073658146315 |
| *DAPK1* | 0.334254844562798 | 0.00124077806858452 |
| *GAB1* | 0.34928305379513 | 0.00124077806858452 |
| *MAST3* | 0.319068440748122 | 0.00124077806858459 |
| *TRIT1* | 0.338026598452006 | 0.00124077806858459 |
| *COX14* | 0.521025178893526 | 0.00125134897768355 |
| *AUP1* | 0.498533753039875 | 0.00126518201458296 |
| *HCLS1* | 0.591506471409075 | 0.00128706342686565 |
| *FBRSL1* | 0.417794293878733 | 0.00129538045516907 |
| *PHF14* | 0.433197974706311 | 0.00130421727516699 |
| *AURKAIP1* | 0.4987423706443 | 0.0013253342871441 |
| *EVI2B* | 0.728951074775882 | 0.00134373993015732 |
| *ZNF800* | 0.513382339051136 | 0.00136893231511095 |
| *RAB8A* | 0.510377071127704 | 0.00138978398957382 |
| *TBC1D9B* | 0.441025093660931 | 0.00139596664045737 |
| *MRPS21* | 0.544728575918481 | 0.00142057618229521 |
| *DYNLRB1* | 0.52765506255954 | 0.00143637566777664 |
| *EMILIN2* | 0.399605166280949 | 0.00150079914846539 |
| *TMEM243* | 0.493251116411667 | 0.00152674184076951 |
| *FAAP20* | 0.453555415329767 | 0.00153251409997136 |
| *BIRC6* | 0.520835910931183 | 0.00154947404218733 |
| *TMBIM4* | 0.53754685113911 | 0.00157536710936617 |
| *BZW1* | 0.463057014802847 | 0.00157665009926537 |
| *FNBP4* | 0.581387046497329 | 0.00157693000575946 |
| *NIPSNAP3A* | 0.311414868204686 | 0.00159359878720178 |
| *IL13RA1* | 0.360452361513241 | 0.00159359878720182 |
| *MCC* | 0.356738858714177 | 0.00159359878720182 |
| *CHAF1A* | 0.431767236431337 | 0.00161636650242069 |
| *TOP1* | 0.568170222298253 | 0.0016488358899701 |

| | | |
|---|---|---|
| *ARL6IP4* | 0.605943002406414 | 0.00166533441957236 |
| *UHMK1* | 0.505680905917445 | 0.00166723406078806 |
| *PRDX3* | 0.407225052168546 | 0.00170028551557665 |
| *MICOS10* | 0.556405878390426 | 0.00173269412244216 |
| *SUZ12* | 0.495818975437534 | 0.00176607599793756 |
| *UBXN1* | 0.464483388949387 | 0.00179384659970133 |
| *SRM* | 0.495862009488482 | 0.00183692067221727 |
| *KDM3A* | 0.453282220749643 | 0.00186940823742539 |
| *FOXN2* | 0.478024294834939 | 0.00187317333069459 |
| *PRDX5* | 0.517896564257767 | 0.0018735156332473 |
| *RALY* | 0.476631059535072 | 0.00188547156522109 |
| *AKIRIN2* | 0.552265276012747 | 0.00189446992715363 |
| *POLE4* | 0.416244841425114 | 0.00194218861397361 |
| *MORF4L1* | 0.573948714833474 | 0.00196031401254213 |
| *SNRPF* | 0.584122014218568 | 0.0019688858896018 |
| *PUM1* | 0.540653985642458 | 0.00197682016892261 |
| *RSU1* | 0.457265734668141 | 0.00198269880061623 |
| *GM2A* | 0.311414868204686 | 0.00204501258470832 |
| *SNX4* | 0.345540652109313 | 0.00204501258470832 |
| *CCDC112* | 0.389821213670007 | 0.00204501258470844 |
| *SLC2A6* | 0.382534859591515 | 0.00204501258470844 |
| *PPP1R14B-AS1* | 0.319068440748122 | 0.0020450125847085 |
| *KDM1A* | 0.417794293878733 | 0.00204769569189692 |
| *LETM1* | 0.438703283641714 | 0.00205500147914412 |
| *SNRPD1* | 0.476856463026022 | 0.00209990347811451 |
| *AKAP17A* | 0.490533616046215 | 0.00218537092374019 |
| *UQCRFS1* | 0.502186481169106 | 0.00218638479397631 |
| *KLF16* | 0.461297488687192 | 0.00220628576477007 |
| *HNRNPA0* | 0.536305996833486 | 0.00221828621806554 |
| *SMIM7* | 0.470089775025725 | 0.0022556570174306 |
| *TOPORS* | 0.390628541637851 | 0.00227545729521366 |
| *ATP6AP1* | 0.438853697554261 | 0.00230943738718592 |
| *TDG* | 0.410806423575481 | 0.00231121524999829 |
| *OGFRL1* | 0.491911877475032 | 0.00231946635714981 |
| *GRK2* | 0.475303294197863 | 0.00232345681086675 |
| *RAB5IF* | 0.44777244874146 | 0.00237248858409052 |
| *PSMD3* | 0.36415633027666 | 0.0023869763698199 |
| *F11R* | 0.279158677268823 | 0.00243543730265822 |

| | | |
|---|---|---|
| *TENT2* | 0.278153812379438 | 0.00245705575521614 |
| *RPL7L1* | 0.478024294834939 | 0.00246793802647993 |
| *IREB2* | 0.464137211373428 | 0.00247206796430818 |
| *UBE2D3* | 0.581343592481847 | 0.00248319457632043 |
| *PHYKPL* | 0.444326678960643 | 0.00249034842147057 |
| *RHOF* | 0.517147232757829 | 0.00249196956332735 |
| *ARL8A* | 0.412830177012349 | 0.00250353676651529 |
| *SDHB* | 0.425556874940803 | 0.00253038377364338 |
| *SUPT4H1* | 0.476459299901452 | 0.00253659455657131 |
| *SLC44A2* | 0.493251116411667 | 0.00254373603165797 |
| *ZNF280D* | 0.422213431866433 | 0.00254476554878573 |
| *DDX39B* | 0.559531200974628 | 0.00259293415992916 |
| *XRCC6* | 0.537375909631406 | 0.00259521875807778 |
| *SCP2* | 0.491268248179999 | 0.00260767008181446 |
| *GALNT7* | 0.319068440748122 | 0.00262209880711711 |
| *ABCD4* | 0.307572801910292 | 0.00262209880711727 |
| *ZNF852* | 0.315246729795713 | 0.00262209880711742 |
| *NDUFC1* | 0.442158842277933 | 0.00265620615611981 |
| *SFSWAP* | 0.443144626423488 | 0.00269407775382517 |
| *EIF3K* | 0.533259048971782 | 0.00269903616911769 |
| *CNOT2* | 0.56579019144631 | 0.00272191961099165 |
| *ADGRG5* | 0.400135627382226 | 0.00276722361236371 |
| *NOP58* | 0.544728575918481 | 0.00281344040252096 |
| *AFF4* | 0.504389646782167 | 0.0028422381024435 |
| *DUT* | 0.563783860481013 | 0.00284950284386424 |
| *SNW1* | 0.525619752970776 | 0.00292884624631586 |
| *KPNA2* | 0.34972925964138 | 0.00294123388710284 |
| *BAG6* | 0.430781062164127 | 0.00296169691508005 |
| *EIF4EBP1* | 0.423911072797083 | 0.00297184052145068 |
| *CARD11* | 0.423862576400927 | 0.00299088921167232 |
| *GTF3A* | 0.588529115741348 | 0.00299140120798008 |
| *MRPL54* | 0.51013563609394 | 0.0030639684278356 |
| *INTS10* | 0.479633547696596 | 0.00309756319541714 |
| *NDUFAB1* | 0.485619842762327 | 0.003109533615762 |
| *EIF3E* | 0.533205112414243 | 0.00311727930058437 |
| *ARF3* | 0.431270526135197 | 0.00311752909077318 |
| *PIAS1* | 0.497518426008896 | 0.00313110986608915 |
| *PSMB7* | 0.437813372001969 | 0.00314544707191273 |

| | | |
|---|---|---|
| *WAPL* | 0.457891768542529 | 0.00321186752244641 |
| *PFDN2* | 0.519041932254831 | 0.00327368100534182 |
| *ILF3* | 0.536890865305796 | 0.00330766070564276 |
| *RWDD1* | 0.522304163942619 | 0.00331443417855523 |
| *SH3BP4* | 0.330473204020371 | 0.00335924874039088 |
| *SMIM13* | 0.299857836782287 | 0.00335924874039088 |
| *SYCP2L* | 0.315246729795713 | 0.00335924874039088 |
| *MYO15B* | 0.307572801910292 | 0.00335924874039118 |
| *HSPA9* | 0.560553543080163 | 0.00340550122184895 |
| *SLC30A5* | 0.410756743138799 | 0.00342361605583897 |
| *GLS* | 0.53545076233047 | 0.00345262964817008 |
| *UBALD2* | 0.586548751613107 | 0.00353059426517463 |
| *NME1* | 0.410756743138799 | 0.00356675357274745 |
| *TOMM7* | 0.420739858643028 | 0.00357195973076268 |
| *NDUFA7* | 0.497050600774005 | 0.00357910679635412 |
| *MTR* | 0.420463771178614 | 0.00362292286206869 |
| *NORAD* | 0.534775101889057 | 0.00369940979606844 |
| *HEBP2* | 0.544728575918481 | 0.00379026814632942 |
| *ANXA2* | 0.697282631768886 | 0.00385435705420019 |
| *MAPRE1* | 0.496606626300309 | 0.00392525599362564 |
| *IQGAP1* | 0.553426681438399 | 0.00397153286703039 |
| *SPAG7* | 0.50524864951126 | 0.00399953877362207 |
| *ADAR* | 0.550479496905586 | 0.00402431479764602 |
| *NCOA4* | 0.537221039249539 | 0.00415080849861572 |
| *SRRM1* | 0.455372847835184 | 0.00422946250569843 |
| *NDRG1* | 0.448472616210135 | 0.00425780557530688 |
| *DACH1* | 0.345540652109313 | 0.00430010730897298 |
| *P3H2* | 0.311414868204686 | 0.00430010730897323 |
| *KCNK10* | 0.311414868204686 | 0.00430010730897335 |
| *KYNU* | 0.307572801910292 | 0.00430010730897335 |
| *SLC2A8* | 0.284303022587445 | 0.00430010730897335 |
| *MTMR14* | 0.452029086301994 | 0.00438207225908117 |
| *SAFB* | 0.42889258750406 | 0.00453960137281138 |
| *SSRP1* | 0.335587178079889 | 0.00459202951887685 |
| *PA2G4* | 0.514461136249684 | 0.00460571729110085 |
| *NUDT3* | 0.281694170084687 | 0.00462387506307905 |
| *MYO9B* | 0.466254517895396 | 0.00462892827310146 |
| *STT3A* | 0.371424330740699 | 0.00468145587424976 |

| | | |
|---|---|---|
| *ETV3* | 0.487652996032716 | 0.00482865930893096 |
| *ARF5* | 0.410072165332018 | 0.00493206297044827 |
| *ZFAND6* | 0.515143083390216 | 0.00497663571050609 |
| *CCDC47* | 0.449625672466009 | 0.00498682977495279 |
| *GORASP2* | 0.497217860068249 | 0.00505883685516826 |
| *THUMPD3-AS1* | 0.406969413815633 | 0.00507425690129739 |
| *GTF3C6* | 0.491911877475032 | 0.00508232820087282 |
| *TRAPPC5* | 0.517147232757829 | 0.00517267381063802 |
| *CCT3* | 0.514181774281267 | 0.00526254196344962 |
| *SPG7* | 0.454041833793855 | 0.00533388615593033 |
| *TRMT112* | 0.466361190603558 | 0.00537097379847687 |
| *BCL2L11* | 0.421300237380269 | 0.00543001227517133 |
| *LBR* | 0.468650498147765 | 0.00543839688252857 |
| *MSRB3* | 0.292101393147852 | 0.00550002211982141 |
| *AC008764.7* | 0.319068440748122 | 0.00550002211982157 |
| *AC097375.1* | 0.311414868204686 | 0.00550002211982157 |
| *RAB34* | 0.29598482763508 | 0.00550002211982157 |
| *GCA* | 0.341788517248205 | 0.00550002211982174 |
| *OAS1* | 0.311414868204686 | 0.00550002211982174 |
| *SLC18B1* | 0.299857836782287 | 0.00550002211982174 |
| *NANS* | 0.410756743138799 | 0.00560109877928909 |
| *ASAH1* | 0.595481876383116 | 0.00563019413821176 |
| *ELP2* | 0.406352893930867 | 0.00563491481587732 |
| *DDX1* | 0.460482447351684 | 0.00563795503106989 |
| *GNG10* | 0.43656482580471 | 0.00577873797990855 |
| *NHLRC3* | 0.375044646419396 | 0.00588778804116942 |
| *ZNF451* | 0.357730061117227 | 0.00589925413937095 |
| *NAA10* | 0.446618490468633 | 0.00605068307717078 |
| *TXNRD1* | 0.353185212455657 | 0.00608995446448434 |
| *GNS* | 0.360615972571411 | 0.00614554991477162 |
| *ESD* | 0.435540963513425 | 0.00628849644682097 |
| *ZNHIT1* | 0.523354925125914 | 0.00644737965889861 |
| *SLC43A2* | 0.396099222779414 | 0.00657236998955 |
| *HSPA5* | 0.546299283793708 | 0.00658101206206356 |
| *C19orf53* | 0.511342407423934 | 0.00659303492193528 |
| *DHX9* | 0.514259890806228 | 0.00662591210276931 |
| *SLAMF7* | 0.463692003827574 | 0.00672428931636995 |
| *ATF4* | 0.618609960845406 | 0.00674048578552892 |

| | | |
|---|---|---|
| *NAA50* | 0.36415633027666 | 0.00678689531670447 |
| *BICD2* | 0.408761632397825 | 0.00689898166853448 |
| *KMT2E* | 0.577063542949669 | 0.00695691633713713 |
| *AC011893.1* | 0.322880054698141 | 0.00702913038068551 |
| *THBD* | 0.389821213670007 | 0.00702913038068572 |
| *DLD* | 0.292101393147852 | 0.00702913038068593 |
| *EAF2* | 0.322880054698141 | 0.00702913038068613 |
| *FARP2* | 0.29598482763508 | 0.00702913038068613 |
| *PDCD11* | 0.330473204020371 | 0.00702913038068613 |
| *RIOK2* | 0.299857836782287 | 0.00702913038068613 |
| *SPATS2* | 0.334254844562798 | 0.00702913038068613 |
| *SRC* | 0.276462269370262 | 0.00702913038068613 |
| *ATP11B* | 0.491911877475032 | 0.00705846059301235 |
| *PPP6C* | 0.396577807969037 | 0.00734365522015012 |
| *LARS* | 0.521230413269132 | 0.00739456301256641 |
| *MIR22HG* | 0.371424330740699 | 0.00750202818427893 |
| *MYL6B* | 0.426440608531002 | 0.00755668755307195 |
| *VPS13D* | 0.371424330740699 | 0.00756326026283055 |
| *ERBIN* | 0.488368357674204 | 0.00756519139132721 |
| *DCTN3* | 0.493251116411667 | 0.00757873665247108 |
| *HSD17B12* | 0.385217945804489 | 0.00760687507782211 |
| *TRIP12* | 0.474936760995197 | 0.00777528088861446 |
| *ANTXR2* | 0.474800393960652 | 0.00789940888083673 |
| *PAIP2* | 0.524290828526163 | 0.00802821469285074 |
| *UPF3A* | 0.390269170595791 | 0.00806562647949115 |
| *NCOA3* | 0.479633547696596 | 0.00820182209978053 |
| *CYB5R4* | 0.405976505971287 | 0.00829043900898716 |
| *CHRAC1* | 0.482629800156515 | 0.00831468850449366 |
| *ELOA* | 0.446618490468633 | 0.00841632001044012 |
| *POLR2I* | 0.508232507950081 | 0.00847244646904467 |
| *VDAC3* | 0.40212418047568 | 0.00852873601101801 |
| *CGAS* | 0.450810712530842 | 0.00860334761412659 |
| *WBP11* | 0.472592562316022 | 0.00878452062530178 |
| *CARHSP1* | 0.412090807340362 | 0.0088118140684856 |
| *MZT2B* | 0.584465437112619 | 0.00888488029003954 |
| *RABEP1* | 0.284303022587445 | 0.00897624732342717 |
| *TRAK1* | 0.315246729795713 | 0.00897624732342717 |
| *CCDC189* | 0.299857836782287 | 0.00897624732342744 |

| | | |
|---|---|---|
| *RABGEF1* | 0.292101393147852 | 0.00897624732342744 |
| *CCDC183* | 0.292101393147852 | 0.00897624732342771 |
| *SCPEP1* | 0.424797681616496 | 0.00921353773392162 |
| *PTGDS* | 2.36119695479612 | 0.00929794517023611 |
| *PICALM* | 0.473278052656772 | 0.00934515801491526 |
| *TMEM259* | 0.425296625090564 | 0.00937826257256547 |
| *SND1-IT1* | 0.378655899971775 | 0.00940966891387678 |
| *SNRPG* | 0.589000230607204 | 0.00950461150381631 |
| *TTYH2* | 0.371424330740699 | 0.00955602609300831 |
| *ARMH1* | 0.476365833862685 | 0.00956114384953831 |
| *POLR2A* | 0.426892085624622 | 0.00959667325960711 |
| *TTC37* | 0.468650498147765 | 0.00962861166607081 |
| *GNPTG* | 0.256136459352969 | 0.00962867673039453 |
| *U2SURP* | 0.546174403706992 | 0.00967762617253618 |
| *SETD7* | 0.413544357238823 | 0.00968531313425412 |
| *BTF3L4* | 0.419651442868363 | 0.00975704565934881 |
| *ANKRD10* | 0.418363404355892 | 0.00991287033074058 |
| *ATP5PO* | 0.530858871051769 | 0.00994441730250541 |
| *HVCN1* | 0.38906167203948 | 0.0101870946987033 |
| *DIDO1* | 0.40043171666821 | 0.0102395447413037 |
| *DNAJB14* | 0.475303294197863 | 0.0103426735612558 |
| *CHPT1* | 0.36415633027666 | 0.0104377399873026 |
| *LARP7* | 0.469238376334217 | 0.0107091237764463 |
| *RNF5* | 0.433197974706311 | 0.010776011082999 |
| *CTCF* | 0.490236078656235 | 0.0108646756598384 |
| *ATP6V1H* | 0.381989623363978 | 0.0108712249222838 |
| *DYRK1A* | 0.445456432378508 | 0.0110391018236294 |
| *GOLGA4* | 0.640914454729656 | 0.011276482040188 |
| *C4orf3* | 0.492480427252199 | 0.0113482245739492 |
| *IMPA2* | 0.288207477043361 | 0.011453761139752 |
| *LRRK2* | 0.411462045055016 | 0.011453761139752 |
| *QDPR* | 0.303720476414686 | 0.011453761139752 |
| *COL24A1* | 0.299857836782287 | 0.0114537611397524 |
| *LINC00877* | 0.284303022587445 | 0.0114537611397524 |
| *PON2* | 0.292101393147852 | 0.0114537611397527 |
| *CRIM1* | 0.303720476414686 | 0.0114537611397534 |
| *TET3* | 0.303720476414686 | 0.0114537611397537 |
| *DEDD2* | 0.487214726680998 | 0.0115957664789633 |

| | | |
|---|---|---|
| *WDR83OS* | 0.472592562316022 | 0.0116206026993711 |
| *CYFIP2* | 0.515320403610311 | 0.0117273851382611 |
| *EDF1* | 0.515582230258964 | 0.0117285034776381 |
| *SH3GLB1* | 0.506450212741782 | 0.0117702342519045 |
| *ZDHHC7* | 0.375044646419396 | 0.0118426640231082 |
| *SRSF10* | 0.610594225579759 | 0.011865144287059 |
| *LSM1* | 0.414782403346628 | 0.011901211471686 |
| *PARL* | 0.371424330740699 | 0.0119426460758281 |
| *RHOQ* | 0.38225813665197 | 0.0120387174421065 |
| *ARHGAP18* | 0.437618492217681 | 0.0121295685905147 |
| *REL* | 0.557500219102609 | 0.0123058824993336 |
| *MRPL36* | 0.406591596597056 | 0.0123218826181813 |
| *SELENOT* | 0.292897647106838 | 0.0123318651079294 |
| *TUT4* | 0.549711983432736 | 0.012334008908056 |
| *RNASEH2B* | 0.455995197341736 | 0.0123753410047127 |
| *NPLOC4* | 0.345824507504402 | 0.0124839503490567 |
| *EFCAB14* | 0.494965201452586 | 0.0125411774403351 |
| *STAG2* | 0.542074122472503 | 0.0126048928781897 |
| *EMC7* | 0.495818975437534 | 0.012752907479255 |
| *FAM120A* | 0.489433551693335 | 0.0127615531507174 |
| *CTSD* | 0.608725542457783 | 0.0128542677648964 |
| *RBBP6* | 0.542369303044559 | 0.0129598340771609 |
| *CYB5R3* | 0.38906167203948 | 0.0129603316621642 |
| *DARS* | 0.482712940628568 | 0.0130584661480667 |
| *CCDC12* | 0.547523594631045 | 0.0131116506541644 |
| *RNF4* | 0.344791005409729 | 0.0131728858153022 |
| *MAD2L2* | 0.392584738603177 | 0.0131910118114187 |
| *PSMB1* | 0.504229213638728 | 0.0135056056868828 |
| *RANBP1* | 0.485619842762327 | 0.0135591501282504 |
| *CHMP3* | 0.399940905416633 | 0.0136582982821679 |
| *MSL1* | 0.442158842277933 | 0.0137166024307389 |
| *MPRIP* | 0.427987942145896 | 0.0139243649547041 |
| *STAT1* | 0.52478972886515 | 0.0139658293611602 |
| *USP8* | 0.403191043460743 | 0.0139932192794952 |
| *CAPZB* | 0.480879936578342 | 0.0143726355476328 |
| *CCAR1* | 0.465497065108891 | 0.0145325416672761 |
| *CENPB* | 0.288207477043361 | 0.0146037911708643 |
| *B3GALNT2* | 0.307572801910292 | 0.0146037911708647 |

| | | |
|---|---|---|
| *LINC01724* | 0.272525854811091 | 0.0146037911708647 |
| *PNO1* | 0.280387972584062 | 0.0146037911708647 |
| *ADGRE2* | 0.322880054698141 | 0.0146037911708652 |
| *YBEY* | 0.292101393147852 | 0.0146037911708652 |
| *NCF4* | 0.385851401375979 | 0.0147173146838118 |
| *UBE3A* | 0.508232507950081 | 0.0149714898135614 |
| *RNF13* | 0.406930353966951 | 0.0150340425907145 |
| *EMC3* | 0.494965201452586 | 0.0154634506054562 |
| *AIMP1* | 0.495003809736281 | 0.0156580178721801 |
| *COLGALT1* | 0.345824507504402 | 0.015741073140454 |
| *ANAPC15* | 0.399090120386047 | 0.0160979507698238 |
| *RGS1* | 0.774678348433011 | 0.0164895664356719 |
| *AC118549.1* | 0.385529981069227 | 0.0166766267111449 |
| *JARID2* | 0.487013078062193 | 0.016702757356001 |
| *MAPK1IP1L* | 0.517072233015615 | 0.0168564670841589 |
| *TBCC* | 0.447572338464297 | 0.0174083206281359 |
| *RBCK1* | 0.449954203031889 | 0.0174362808652306 |
| *SDHC* | 0.367740667625699 | 0.0175710758993963 |
| *TNNI2* | 0.428286667696375 | 0.0178902720424306 |
| *TAP2* | 0.329390912115983 | 0.0181866614724141 |
| *TMEM160* | 0.499933109307127 | 0.0182070037866627 |
| *SENP6* | 0.526044012653554 | 0.0182604434581982 |
| *RAD23A* | 0.45045617667088 | 0.0184408081676378 |
| *ALYREF* | 0.422213431866433 | 0.0185369519201481 |
| *KLHDC3* | 0.272525854811091 | 0.0186059306834016 |
| *EIF4EBP3* | 0.280387972584062 | 0.0186059306834027 |
| *LCNL1* | 0.345540652109313 | 0.0186059306834027 |
| *OSGIN2* | 0.284303022587445 | 0.0186059306834027 |
| *BRPF1* | 0.29598482763508 | 0.0186059306834033 |
| *MCTP1* | 0.292101393147852 | 0.0186059306834044 |
| *SPRED2* | 0.276462269370262 | 0.0186059306834044 |
| *ZC3H14* | 0.264620656725745 | 0.0186059306834044 |
| *TNFSF13* | 0.375044646419396 | 0.0187068507847598 |
| *KCNAB2* | 0.326188480077639 | 0.0189705714553222 |
| *CAPZA1* | 0.546747743555133 | 0.0190328695652393 |
| *AC253572.2* | 0.499085910362768 | 0.0194803071740688 |
| *ZSWIM8* | 0.39960516628095 | 0.0195192275321131 |
| *RAB7A* | 0.442158842277933 | 0.0198041974104724 |

| | | |
|---|---|---|
| *SNHG14* | 0.53051471669878 | 0.0199127823500207 |
| *EIF2S2* | 0.535217759537394 | 0.0202026869739997 |
| *CAPN15* | 0.42889258750406 | 0.020404569101267 |
| *ERGIC1* | 0.436801632108426 | 0.0204422106531766 |
| *ITGB1BP1* | 0.402537422004516 | 0.0205872982442183 |
| *ENY2* | 0.486467738784905 | 0.0206165177360625 |
| *WSB1* | 0.517961666355695 | 0.0206900525225071 |
| *RPS19BP1* | 0.476785185387565 | 0.0210705361297701 |
| *GNL3* | 0.473931188332412 | 0.021073715230715 |
| *CTSH* | 0.438703283641714 | 0.0210904441494201 |
| *SNF8* | 0.427721459054126 | 0.0211780342495198 |
| *LAMTOR4* | 0.499229027771833 | 0.0212682685526182 |
| *SEPHS1* | 0.385529981069227 | 0.0213905953006181 |
| *DDX18* | 0.522304163942619 | 0.0214312018891951 |
| *BAX* | 0.443144626423488 | 0.0214538036038204 |
| *NAA25* | 0.367740667625699 | 0.0215622382402806 |
| *MCM5* | 0.385529981069227 | 0.0216073025947329 |
| *VCP* | 0.504571449446498 | 0.0216620082373642 |
| *REST* | 0.558284899394386 | 0.0217445863740314 |
| *MARCH2* | 0.408134027337262 | 0.0218436174142477 |
| *PSMC1* | 0.472012320037328 | 0.022019600453567 |
| *CLIC1* | 0.574723316216318 | 0.0221411275691351 |
| *TMEM242* | 0.259384008359598 | 0.0222737716095428 |
| *NSA2* | 0.551487608322587 | 0.0223453038314329 |
| *NCOR2* | 0.403150461892523 | 0.0226940122229407 |
| *PSMA1* | 0.506080217625342 | 0.0229082989668114 |
| *DOCK8* | 0.494162321449178 | 0.0229487703693667 |
| *ARHGAP4* | 0.481152973893797 | 0.0230481487018535 |
| *MYO9A* | 0.389435738725018 | 0.023067950893209 |
| *ARID1A* | 0.385575313911566 | 0.0235757776369647 |
| *SERPING1* | 0.284303022587445 | 0.0236869762557346 |
| *RALGPS2* | 0.276462269370262 | 0.0236869762557353 |
| *DAPK2* | 0.288207477043361 | 0.023686976255736 |
| *NUP42* | 0.272525854811091 | 0.023686976255736 |
| *FUT7* | 0.307572801910292 | 0.0236869762557367 |
| *LINC01226* | 0.311414868204686 | 0.0236869762557367 |
| *SNRPC* | 0.440018344791968 | 0.0240328178363336 |
| *MAPKAP1* | 0.40940741137156 | 0.0241614288247251 |

| | | |
|---|---|---|
| *STK11IP* | 0.360508553261516 | 0.0243345201728417 |
| *ATP6V1A* | 0.356851529653426 | 0.0245440250370449 |
| *COX17* | 0.476785185387565 | 0.024769816032828 |
| *AHCYL1* | 0.353185212455657 | 0.0249671710583351 |
| *PNPLA2* | 0.454758879057567 | 0.0249711378517202 |
| *LTA4H* | 0.537046790195533 | 0.0249765309203342 |
| *DDX27* | 0.497050600774005 | 0.0250179556286844 |
| *RNF7* | 0.415159233710763 | 0.0250952283827006 |
| *CTNNB1* | 0.494552967532335 | 0.0253073761533443 |
| *NIPBL* | 0.554678236326086 | 0.0255031223756278 |
| *ZDHHC21* | 0.378440556282908 | 0.0256231742749046 |
| *ADD3* | 0.520835910931183 | 0.0257622535165472 |
| *SURF4* | 0.40093617955142 | 0.0263177432597281 |
| *ALG2* | 0.42060959860584 | 0.0263901253506941 |
| *LMBRD1* | 0.423862576400927 | 0.0264032844518653 |
| *SPPL3* | 0.324726185448122 | 0.0264254848662798 |
| *SVIP* | 0.475303294197863 | 0.0266161231690228 |
| *PHKB* | 0.385529981069227 | 0.0266359102488884 |
| *MTMR1* | 0.374882736869718 | 0.0269840767242411 |
| *PCMTD1* | 0.487013078062193 | 0.0272963419468654 |
| *RBM6* | 0.532636469263814 | 0.0273058809540448 |
| *WASHC4* | 0.374882736869718 | 0.0275191539629563 |
| *KIAA0232* | 0.305561676365811 | 0.0278544826189731 |
| *SMU1* | 0.518107695511231 | 0.0279426097667649 |
| *TCOF1* | 0.473278052656772 | 0.0285275807639556 |
| *RAN* | 0.553279905205813 | 0.0286202228518482 |
| *CSRNP1* | 0.474936760995197 | 0.0289471903640967 |
| *PLEC* | 0.374725571986847 | 0.0293069611892386 |
| *KPNA4* | 0.458483712824813 | 0.029365025098511 |
| *TARBP1* | 0.412830177012349 | 0.0297531719689569 |
| *TOMM22* | 0.426892085624622 | 0.0301232922733044 |
| *PDK2* | 0.288207477043361 | 0.0301331464956141 |
| *ATP13A1* | 0.2606517545228 | 0.030133146495615 |
| *SLC23A2* | 0.280387972584062 | 0.030133146495615 |
| *TFPT* | 0.264620656725745 | 0.030133146495615 |
| *WDR41* | 0.2606517545228 | 0.030133146495615 |
| *DENND5B* | 0.272525854811091 | 0.0301331464956159 |
| *EBNA1BP2* | 0.268578670294429 | 0.0301331464956159 |

| | | |
|---|---|---|
| *GAB2* | 0.2606517545228 | 0.0301331464956159 |
| *PGAM5* | 0.272525854811091 | 0.0301331464956159 |
| *PRDX4* | 0.268578670294429 | 0.0301331464956159 |
| *ARMC10* | 0.280387972584062 | 0.0301331464956168 |
| *EEF2K* | 0.276462269370262 | 0.0301331464956168 |
| *TM9SF1* | 0.29598482763508 | 0.0301331464956177 |
| *GNPTAB* | 0.447987921080131 | 0.0302737699808836 |
| *PTTG1IP* | 0.392170706446256 | 0.030552320201064 |
| *IRAK1* | 0.446430333306202 | 0.0309384604671702 |
| *PDCD4* | 0.599644409224791 | 0.0312199468855636 |
| *SMC3* | 0.513271144396883 | 0.0312208223803303 |
| *PGP* | 0.349509554312259 | 0.0313100327653374 |
| *FLII* | 0.385529981069227 | 0.0313863243915238 |
| *TMEM141* | 0.408761632397825 | 0.0316793035001342 |
| *ACYP2* | 0.334712552384178 | 0.0322993986782405 |
| *MKRN1* | 0.378440556282908 | 0.0324309489499712 |
| *RAB4B* | 0.330989466341461 | 0.032529951940208 |
| *SENP2* | 0.371316121849526 | 0.0327977869442628 |
| *AL360012.1* | 0.339554357043582 | 0.0328956987552444 |
| *LGALS1* | 0.750354146675522 | 0.0330929939823062 |
| *DIS3* | 0.417349269442253 | 0.0333561588559075 |
| *ST3GAL4* | 0.33197592452751 | 0.0335647528587587 |
| *SIDT1* | 0.450810712530842 | 0.0338224216968561 |
| *NAPG* | 0.346367016833131 | 0.0338910316134441 |
| *ZBTB33* | 0.371316121849526 | 0.0340182607890859 |
| *RGS10* | 0.63827991477226 | 0.0342461643834579 |
| *TMEM50B* | 0.356960828872456 | 0.0348210011734553 |
| *PRRC2A* | 0.401454871699635 | 0.0350836645755489 |
| *ARF4* | 0.437992211104824 | 0.0352581001161603 |
| *PRPF8* | 0.476856463026022 | 0.0356821031558983 |
| *DNAJC8* | 0.561418036289426 | 0.035797934673293 |
| *KDM6B* | 0.610439371821129 | 0.0366704125464721 |
| *DCLRE1C* | 0.271525352366395 | 0.0371751895052674 |
| *DUSP3* | 0.256671903610254 | 0.0383054184862232 |
| *SLC12A3* | 0.280387972584062 | 0.0383054184862232 |
| *TUBB6* | 0.256671903610254 | 0.0383054184862232 |
| *CDC16* | 0.2606517545228 | 0.0383054184862244 |
| *TRPM2* | 0.256671903610254 | 0.0383054184862244 |

| | | |
|---|---|---|
| *HDAC5* | 0.25268104341422 | 0.0383054184862255 |
| *NDUFA2* | 0.456753322998436 | 0.0384334193383016 |
| *HDAC2* | 0.444326678960643 | 0.0384390974820909 |
| *DSE* | 0.367794907340008 | 0.0389536373454924 |
| *STAT6* | 0.342130023946661 | 0.0403625131147593 |
| *UBXN6* | 0.325305401305302 | 0.0407406091488549 |
| *CAPN1* | 0.346626405719083 | 0.040977928805649 |
| *RPA1* | 0.330989466341461 | 0.0412239560162008 |
| *SNAP29* | 0.392170706446256 | 0.0417286984678597 |
| *CNOT1* | 0.427581710418293 | 0.0419882075451801 |
| *ZFAS1* | 0.518733367385536 | 0.0432254757042761 |
| *PYURF* | 0.437219792466383 | 0.0435334589406768 |
| *PLEKHA2* | 0.271868740228614 | 0.0437632091519981 |
| *LARP1* | 0.398103662199998 | 0.0443476859768112 |
| *EIF5A* | 0.539754540134167 | 0.0445824586770475 |
| *PCM1* | 0.632842403887043 | 0.04497378907149 |
| *TMEM147* | 0.282810375201016 | 0.0463808494546423 |
| *VDAC2* | 0.47001096291782 | 0.0466763542701241 |
| *EIF3G* | 0.265975936337757 | 0.0478921651806159 |
| *STXBP2* | 0.564395609374557 | 0.0480533462425738 |
| *LINC01184* | 0.268578670294429 | 0.0486587671078259 |
| *DHX37* | 0.276462269370262 | 0.0486587671078286 |
| *ZNF467* | 0.2606517545228 | 0.0486587671078286 |
| *SLC43A3* | 0.272525854811091 | 0.0486587671078302 |
| *SNTA1* | 0.264620656725745 | 0.0486587671078302 |
| *IMPACT* | 0.276462269370262 | 0.0486587671078316 |
| *CCR2* | 0.272525854811091 | 0.0486587671078345 |
| *PITPNB* | 0.298567988649083 | 0.0488305363921709 |
| *SFMBT2* | 0.375044646419396 | 0.0491761765342327 |
| *STOML2* | 0.424697872211711 | 0.0492230979538134 |
| *NDUFV2* | 0.505433017128651 | 0.0493662088641161 |
| *LIMD1* | 0.38906167203948 | 0.0493870563412784 |
| *MRPS12* | 0.345824507504402 | 0.0495012300401732 |
| *VAV3* | 0.308122295362332 | 0.0495891865673816 |
| *IST1* | 0.405378993003173 | 0.0496800332317121 |
| *SRGAP2* | 0.342130023946661 | 0.0499357857483282 |
