## Supplementary Table 19 for "Heterogeneity of circulating epithelial cells in breast cancer at single-cell resolution: identifying tumor and hybrid cells"

Supplementary Table 19. Specific upregulated genes in diploid cells of CD45$^{+}$ CEC clusters

| Genes | Cluster 1 | | Genes | Cluster 2 | | Genes | Cluster 3 | |
|---|---|---|---|---|---|---|---|---|
| | LogFC | Adjusted P-value | | LogFC | Adjusted P-value | | LogFC | Adjusted P-value |
| *CCNL1* | 0.963403057004064 | 3.28604986820159E-11 | *RPS10* | 0.32723231793223 | 8.24001249960321E-06 | *GAPDH* | 2.19186755432299 | 1.92698334140158E-10 |
| *NAMPT* | 1.33990878402998 | 1.38787707972421E-08 | *CHCHD10* | 0.512187008017497 | 0.0034770286109711 | *YBX3* | 1.20391518208752 | 5.10869890823066E-09 |
| *DDX3X* | 0.80667456597761 | 1.51113167679077E-08 | *TBCB* | 0.486873988571598 | 0.0054855082299881 | *GIHCG* | 1.93118764160785 | 1.47469152994057E-08 |
| *RELT* | 0.617442759558891 | 2.55928222986735E-08 | *GLCE* | 0.375076179014138 | 0.00638174342280257 | *SERPINB1* | 2.51419394536368 | 5.15533788404645E-08 |
| *XIST* | 0.872156849182932 | 5.94660661772433E-08 | *TRAPPC12* | 0.294834434005087 | 0.00646501283276492 | *CDK6* | 1.91332505373316 | 6.07050932474477E-08 |
| *JMJD1C* | 0.814817739286225 | 9.25175566739101E-08 | *MAGED1* | 0.368692562223284 | 0.00790680983282098 | *PRSS57* | 1.47754470316051 | 1.78181497466538E-07 |
| *AKAP13* | 0.874214021983925 | 1.01863837507084E-07 | *FBH1* | 0.274174963438994 | 0.00803973421650948 | *UQCRH* | 1.47788629071766 | 2.89842425051721E-07 |
| *GRINA* | 0.586548751613108 | 1.54922112294727E-07 | *SLC1A5* | 0.400378587451989 | 0.00923288229432892 | *ANKRD28* | 1.16438681790088 | 3.32343409792001E-06 |
| *HNRNPL* | 0.675864582846143 | 1.97874613514191E-07 | *PIGT* | 0.388445409528936 | 0.00933989745385369 | *IMPDH2* | 1.12066544047156 | 0.0000111375314440676 |
| *DYNC1H1* | 0.746625967099073 | 2.29434948951559E-07 | *NUDT17* | 0.294834434005087 | 0.00991179183522333 | *GLUL* | 1.37779045601008 | 0.0000130840642868637 |
| *KDM5A* | 0.783752111517808 | 3.17055370545996E-07 | *LSM8* | 0.327569672088073 | 0.0104477870287836 | *LAPTM4B* | 0.869939459435627 | 0.0000153704554213148 |
| *TET2* | 0.638595499382703 | 3.5490693017806E-07 | *LYSMD2* | 0.369727123071411 | 0.0119475933901245 | *GAS5* | 1.51986747249927 | 0.000023148180965188 |
| *ACTN4* | 0.700439718141092 | 4.2728816954182E-07 | *CCDC85B* | 0.458599534576422 | 0.0138432465137158 | *GATA2* | 1.0741890089293 | 0.0000549542388645328 |
| *FNBP1* | 0.700384988714827 | 5.76648063660377E-07 | *SMDT1* | 0.470572176242497 | 0.0139118517294654 | *SLC40A1* | 1.12256664220626 | 0.000168059620060165 |
| *PNRC1* | 0.851762455506148 | 9.0380206344199E-07 | *RTRAF* | 0.469800475212657 | 0.0142378458220373 | *LMO2* | 0.814444346843923 | 0.00041508720622197 |
| *AC007952.4* | 0.808941172949556 | 1.06147384865783E-06 | *VIPR2* | 0.40345798038396 | 0.0149446820596014 | *SMIM24* | 0.726981505593584 | 0.00041508720622197 |
| *WDR26* | 0.489687212360519 | 1.07690659104391E-06 | *CLTB* | 0.421016351768265 | 0.0159706268815138 | *PBX1* | 0.633872101202103 | 0.000415087206221973 |
| *SRSF6* | 0.697282631768886 | 1.14429269811354E-06 | *PRMT9* | 0.439906951638626 | 0.018169785733531 | *CYTOR* | 1.08072888885397 | 0.00113113038366498 |
| *MT-ND4L* | 0.659403882388622 | 1.50372966703198E-06 | *GPX7* | 0.263733216717347 | 0.0236919905687881 | *HSBP1* | 0.568777072980218 | 0.00123130113732357 |
| *PTEN* | 0.583798140597073 | 3.07529503585195E-06 | *SGSM3* | 0.384882251128821 | 0.0250799549661575 | *LDHB* | 1.34041718199238 | 0.00160332704556095 |
| *OTUB1* | 0.401988403802399 | 0.0000032893825253712 | *TMEM208* | 0.366621212203589 | 0.0293292873646084 | *IL18* | 0.798737345584202 | 0.00182444488232825 |
| *KCTD20* | 0.528650367904906 | 3.44888797105905E-06 | *ENPP2* | 0.270702771532189 | 0.0293557741711723 | *HSPD1* | 1.087424836086 | 0.00198493352817093 |
| *KLF13* | 0.681896628200588 | 4.49835576974384E-06 | *TM2D1* | 0.382396364856131 | 0.0296598561457455 | *CNRIP1* | 0.601450623509725 | 0.00201607788311072 |
| *HNRNPH1* | 0.734206374782194 | 5.00135800610526E-06 | *EIF6* | 0.432604326043477 | 0.0301715224801435 | *UROD* | 0.499571009490512 | 0.00201607788311072 |
| *LRRFIP1* | 0.758520261290754 | 5.35968992439801E-06 | *CDC123* | 0.271668107828419 | 0.0374571810748499 | *FHL1* | 0.869939459435627 | 0.00201607788311073 |
| *TFRC* | 0.60059652609874 | 5.88116177599693E-06 | *RMC1* | 0.270702771532189 | 0.0449905436496244 | *DIPK1B* | 0.601450623509725 | 0.00201607788311074 |
| *BCL3* | 0.604360027729568 | 6.40362056594348E-06 | | | | *SNHG8* | 1.4045208216246 | 0.00283810187829936 |
| *CDKN1A* | 0.686084425164022 | 7.42084048823739E-06 | | | | *BZW2* | 0.643554654599442 | 0.00853681206393322 |
| *CSNK1A1* | 0.628052303792607 | 7.68104426317144E-06 | | | | *SPN* | 0.784114736782715 | 0.00916094838473162 |
| *NASP* | 0.662497604846406 | 8.17776845538287E-06 | | | | *HBD* | 1.35147237050138 | 0.00940556689698754 |
| *LAMP2* | 0.566084649053392 | 0.0000108653699467557 | | | | *CYTL1* | 1.22948184612277 | 0.00940556689698757 |
| *MPZL1* | 0.502984034973977 | 0.0000115340085751214 | | | | *EREG* | 0.756728848987636 | 0.00940556689698757 |
| *VAPA* | 0.703082501515919 | 0.0000116158711893122 | | | | *SNHG19* | 0.534336427651188 | 0.00940556689698757 |
| *RUFY3* | 0.577427361644559 | 0.0000121079002095892 | | | | *SYPL1* | 0.749349318622038 | 0.00969237271459255 |
| *WDR1* | 0.657807434782929 | 0.0000125401456788737 | | | | *PARP1* | 0.763024255519115 | 0.0152788831988103 |

| | | |
|---|---|---|
| *RPS6KA3* | 0.712766467816528 | 0.0000126130442232299 |
| *NFKBID* | 0.639935964360132 | 0.0000127387079413441 |
| *PHC3* | 0.662497604846406 | 0.0000131633606067418 |
| *MTRNR2L8* | 0.790728859921981 | 0.0000134254881588177 |
| *SLC38A10* | 0.486343769286148 | 0.0000154188071263722 |
| *EIF3A* | 0.749810022774409 | 0.0000172882807440205 |
| *RPL36* | 0.385231322980974 | 0.0000192831473404238 |
| *ARGLU1* | 0.682705746709052 | 0.0000196062486447804 |
| *VTI1B* | 0.507520505447837 | 0.0000215634736635362 |
| *NONO* | 0.59735550656969 | 0.0000219253286237042 |
| *ACAP2* | 0.646833609081528 | 0.0000229991120284368 |
| *NDUFA3* | 0.609268828113191 | 0.0000243126369106259 |
| *SRSF3* | 0.669821300839667 | 0.0000260605841569213 |
| *C1GALT1* | 0.541034092360739 | 0.0000272299063160207 |
| *POLR1D* | 0.586548751613108 | 0.0000330145590537597 |
| *METAP2* | 0.601024074128987 | 0.0000339120953742378 |
| *CPT1A* | 0.574374037667005 | 0.0000354094130690017 |
| *MCL1* | 1.0808495294166 | 0.0000355382061736642 |
| *PTPN18* | 0.515143083390216 | 0.0000368130336516929 |
| *SAT1* | 1.44952223069142 | 0.0000368603666067168 |
| *ATPAF2* | 0.342461259177341 | 0.0000375883646749535 |
| *PRPF38B* | 0.739004739439805 | 0.0000386499652894202 |
| *TSPYL2* | 0.88309469566782 | 0.0000396069595680122 |
| *CAPNS1* | 0.649558549138908 | 0.0000422002303392592 |
| *CD53* | 0.400682206301774 | 0.0000429130018054102 |
| *PABPC4* | 0.583601467562771 | 0.0000491133420565897 |
| *PTPN1* | 0.586548751613108 | 0.0000517849803262752 |
| *HOOK3* | 0.457265734668141 | 0.0000521632255028322 |
| *PPP1CA* | 0.601735155427784 | 0.000054122689645114 |
| *ICOSLG* | 0.538930731380368 | 0.0000542923865063786 |
| *TNIP1* | 0.525962598752069 | 0.0000579947766227017 |
| *SP3* | 0.589155247427589 | 0.0000586543364804687 |
| *GADD45B* | 0.899863716195762 | 0.0000611129883525532 |
| *SETX* | 0.563783860481013 | 0.0000633948799213936 |
| *ARL5A* | 0.505680905917445 | 0.0000662032538352006 |
| *MAP7D1* | 0.507891807738109 | 0.0000669320657442477 |
| *ZFP36* | 1.07931755412452 | 0.0000685105239167656 |
| *YME1L1* | 0.605594507303654 | 0.0000706493498538459 |

| | | |
|---|---|---|
| *LYL1* | 1.01238372445583 | 0.018327963990973 |
| *TPI1* | 1.11191939800675 | 0.0240508519291889 |
| *ITGA4* | 0.787532512509926 | 0.0250018549786163 |
| *NDUFS5* | 0.820432416683521 | 0.0269790314122695 |
| *ALDH1A1* | 0.814444346843923 | 0.0422880476340244 |
| *LINC02573* | 0.568283759574526 | 0.0422880476340244 |
| *MAP7D3* | 0.46394709975979 | 0.0422880476340244 |
| *RAB13* | 0.46394709975979 | 0.0422880476340244 |
| *NOA1* | 0.46394709975979 | 0.0422880476340246 |
| *LGALS9* | 0.512310121321189 | 0.0439212674317261 |
| *PRKACB* | 0.886852842371973 | 0.0496847601928101 |

| | | |
|---|---|---|
| *MT-ATP8* | 0.840357692233899 | 0.0000708716833553191 |
| *GLG1* | 0.592108276955008 | 0.0000729265635468819 |
| *MTMR10* | 0.342461259177341 | 0.0000871146967627198 |
| *RBM4* | 0.576075420383599 | 0.0000904250410043516 |
| *EIF5* | 0.713439384071789 | 0.0000948235974904735 |
| *PSEN1* | 0.389821213670007 | 0.000096769789468307 |
| *SRSF4* | 0.601735155427784 | 0.00010023007210732 |
| *CNOT7* | 0.528132065387786 | 0.000101275849110269 |
| *BTK* | 0.431767236431337 | 0.000103277865019221 |
| *TRA2B* | 0.609268828113191 | 0.000103941926391967 |
| *BRD4* | 0.636560519780931 | 0.000112388174238192 |
| *METTL7A* | 0.489687212360519 | 0.000122510427807393 |
| *SCAF8* | 0.378877835983425 | 0.000125406981639418 |
| *SRPRA* | 0.592034290696837 | 0.000129430695757758 |
| *UFM1* | 0.547868283449048 | 0.00014178689210509 |
| *SYNCRIP* | 0.608993898137916 | 0.000144666293411929 |
| *GLO1* | 0.566649194175403 | 0.000146081291312599 |
| *INPP4A* | 0.532553861972378 | 0.000148061890837068 |
| *BCLAF1* | 0.723099267484578 | 0.000156746615804084 |
| *IQGAP2* | 0.632645166202562 | 0.000159641983855418 |
| *CHPF2* | 0.345540652109313 | 0.000162366327129114 |
| *DNAJB11* | 0.548309947325416 | 0.000166625083477699 |
| *RNF114* | 0.527292166056333 | 0.000168190507012846 |
| *TCF25* | 0.544728575918481 | 0.000171529067331219 |
| *UHRF1BP1L* | 0.445606143896402 | 0.000174404776247134 |
| *STK24* | 0.471327853824456 | 0.000175733293056496 |
| *EMB* | 0.543815766624575 | 0.000182646964330345 |
| *NFX1* | 0.396099222779414 | 0.000187812539440172 |
| *PPP1R15A* | 0.805666040916695 | 0.000189922575743234 |
| *PI4KA* | 0.510997718605931 | 0.000196365887983673 |
| *KARS* | 0.456602579041255 | 0.000198237392260345 |
| *UBTF* | 0.513019716191143 | 0.000204199043269373 |
| *LILRB2* | 0.418604114299036 | 0.00021002286358564 |
| *EMC10* | 0.513150673849371 | 0.000217903133821373 |
| *EPN1* | 0.445606143896402 | 0.000218996234312251 |
| *EIF4G3* | 0.490848335192522 | 0.000226998017541447 |
| *ERO1B* | 0.548848042427884 | 0.000235148421281842 |
| *DDX21* | 0.787921593868035 | 0.000235491409550906 |

| | | |
|---|---|---|
| *RBM23* | 0.476459299901452 | 0.000249913154719983 |
| *SIN3A* | 0.262441907193779 | 0.000251869849803444 |
| *NRIP1* | 0.514096957264817 | 0.000255127539031147 |
| *CELF1* | 0.651889505280426 | 0.000257425302158029 |
| *FAM53C* | 0.559803896380579 | 0.000260318587967584 |
| *ABCA7* | 0.353015772671459 | 0.000271417315762168 |
| *FAM204A* | 0.501052531331501 | 0.000272062661304644 |
| *WDFY2* | 0.457265734668141 | 0.000273394307100439 |
| *LINC-PINT* | 0.68149715566837 | 0.000276541164554851 |
| *SF3B2* | 0.563465138500066 | 0.000291258824623704 |
| *MAF1* | 0.435148161848098 | 0.000298373830859682 |
| *SEC13* | 0.457938476597277 | 0.000302137471538636 |
| *TAX1BP1* | 0.646191698040509 | 0.000314443964956491 |
| *TMEM107* | 0.499416159887133 | 0.000314609557696814 |
| *ME2* | 0.38075829761379 | 0.000321559358306928 |
| *TTC17* | 0.454758879057567 | 0.000324591536650362 |
| *AKAP8* | 0.524621002469906 | 0.000327306751728376 |
| *DDAH2* | 0.501659854026595 | 0.00033195280821135 |
| *AP1S3* | 0.382534859591515 | 0.000350439367925165 |
| *CLMN* | 0.393450637070698 | 0.000350439367925165 |
| *CIR1* | 0.62246832586805 | 0.000357022959221806 |
| *GINM1* | 0.487652996032716 | 0.00035703126075902 |
| *LINC00909* | 0.438703283641714 | 0.000360050511599983 |
| *PXK* | 0.435239428338142 | 0.000368859048626 |
| *TUBGCP3* | 0.442158842277933 | 0.000369935456386329 |
| *SMARCC1* | 0.518189958949081 | 0.000370276217149847 |
| *NUS1* | 0.527781343363234 | 0.000394645413345556 |
| *NAGA* | 0.43656482580471 | 0.000402359644433317 |
| *PAG1* | 0.608081913162749 | 0.000407314014583432 |
| *PSMB4* | 0.533368713753933 | 0.00042394849207509 |
| *ARHGDIA* | 0.617021103837378 | 0.000428440669720956 |
| *AP2B1* | 0.533368713753933 | 0.000429340926673334 |
| *CLPX* | 0.442158842277933 | 0.000461887997995462 |
| *EIF1B* | 0.601371091318631 | 0.000504859591453881 |
| *DNM2* | 0.535524748588641 | 0.00051164087108955 |
| *YY1* | 0.612083843720245 | 0.000539064273705439 |
| *CMTM6* | 0.586548751613108 | 0.000543138986691278 |
| *TNPO1* | 0.455436837265949 | 0.000554388962823108 |

| | | |
|---|---|---|
| *TMED5* | 0.560912824626258 | 0.000567679721158084 |
| *CTNNA1* | 0.345540652109313 | 0.000582629634728351 |
| *BACH1* | 0.382534859591515 | 0.000582629634728369 |
| *MRPL14* | 0.330473204020371 | 0.00058262963472842 |
| *EIF2S3* | 0.582336500326193 | 0.000596495922603233 |
| *PRRC2B* | 0.441118312091544 | 0.000616278176271502 |
| *PRKDC* | 0.488368357674204 | 0.000619326981295154 |
| *DNAJC1* | 0.512089759412663 | 0.000622237785849056 |
| *PCBP1* | 0.638950449572927 | 0.000622585965404933 |
| *CACUL1* | 0.371316121849526 | 0.000644130389973652 |
| *WDR43* | 0.591757042615869 | 0.000657890236788019 |
| *RASGEF1B* | 0.645083962582862 | 0.000662587128523687 |
| *AKAP9* | 0.734694369388644 | 0.00067139901388116 |
| *LAMP1* | 0.554066440497817 | 0.000694700536481567 |
| *PLEKHB2* | 0.423911072797083 | 0.000697514167109428 |
| *BTAF1* | 0.507891807738109 | 0.000699155113897287 |
| *POLR2G* | 0.443275047394262 | 0.000703621968278795 |
| *ZNF655* | 0.463692003827574 | 0.000707948709506294 |
| *BRD2* | 0.58442245050472 | 0.000754819973927575 |
| *ATRAID* | 0.458483712824813 | 0.00077324585629244 |
| *PSMD9* | 0.526427759175537 | 0.000795456578380502 |
| *NCOA1* | 0.49801207701145 | 0.000822073918231897 |
| *POM121* | 0.389435738725018 | 0.000826142879972809 |
| *SUSD6* | 0.441025093660931 | 0.000828035386722362 |
| *MED30* | 0.304363245434402 | 0.000830257504760986 |
| *MLXIP* | 0.495818975437534 | 0.000839246491427728 |
| *RBM5* | 0.558803761130204 | 0.000854583219366118 |
| *AKR1A1* | 0.468650498147765 | 0.000868990612792027 |
| *SCFD1* | 0.49403661282357 | 0.000883995779954373 |
| *SMG7* | 0.434203827804916 | 0.00089684575706651 |
| *FMNL1* | 0.463300976574635 | 0.000900093646778591 |
| *MALAT1* | 0.505851139364285 | 0.000951574798057288 |
| *UQCRC2* | 0.507891807738109 | 0.00096405258580204 |
| *ATP6V0A1* | 0.341788517248205 | 0.00096524460213082 |
| *P2RX1* | 0.341788517248205 | 0.00096524460213082 |
| *ZDHHC14* | 0.322880054698141 | 0.00096524460213082 |
| *COX5B* | 0.555396231288169 | 0.000968028532381136 |
| *SRSF2* | 0.634356142495238 | 0.000972460615226542 |

| | | |
|---|---|---|
| *PAPOLA* | 0.53165050546421 | 0.00099443338790754 |
| *THOC2* | 0.566084649053392 | 0.00103713355896904 |
| *AC016831.5* | 0.504389646782167 | 0.00104945963836077 |
| *KDM2A* | 0.527981132182554 | 0.0010535354400477 |
| *AL355075.4* | 0.614382959809833 | 0.00105941095421859 |
| *PRPF40A* | 0.602668416976385 | 0.00110906940739969 |
| *CALHM6* | 0.49801207701145 | 0.0011213226283493 |
| *PGGT1B* | 0.481241007518482 | 0.00114651234709989 |
| *RHEB* | 0.500392107863393 | 0.00119467920296092 |
| *EIF4G1* | 0.495862009488482 | 0.00124073658146315 |
| *MAST3* | 0.319068440748122 | 0.00124077806858459 |
| *TRIT1* | 0.338026598452006 | 0.00124077806858459 |
| *AUP1* | 0.498533753039875 | 0.00126518201458296 |
| *HCLS1* | 0.591506471409075 | 0.00128706342686565 |
| *FBRSL1* | 0.417794293878733 | 0.00129538045516907 |
| *AURKAIP1* | 0.4987423706443 | 0.0013253342871441 |
| *EVI2B* | 0.728951074775882 | 0.00134373993015732 |
| *RAB8A* | 0.510377071127704 | 0.00138978398957382 |
| *TBC1D9B* | 0.441025093660931 | 0.00139596664045737 |
| *EMILIN2* | 0.399605166280949 | 0.00150079914846539 |
| *BIRC6* | 0.520835910931183 | 0.00154947404218733 |
| *FNBP4* | 0.581387046497329 | 0.00157693000575946 |
| *NIPSNAP3A* | 0.311414868204686 | 0.00159359878720178 |
| *IL13RA1* | 0.360452361513241 | 0.00159359878720182 |
| *TOP1* | 0.568170222298253 | 0.0016488358899701 |
| *UHMK1* | 0.505680905917445 | 0.00166723406078806 |
| *MICOS10* | 0.556405878390426 | 0.00173269412244216 |
| *SUZ12* | 0.495818975437534 | 0.00176607599793756 |
| *UBXN1* | 0.464483388949387 | 0.00179384659970133 |
| *KDM3A* | 0.453282220749643 | 0.00186940823742539 |
| *FOXN2* | 0.478024294834939 | 0.00187317333069459 |
| *RALY* | 0.476631059535072 | 0.00188547156522109 |
| *AKIRIN2* | 0.552265276012747 | 0.00189446992715363 |
| *MORF4L1* | 0.573948714833474 | 0.00196031401254213 |
| *PUM1* | 0.540653985642458 | 0.00197682016892261 |
| *RSU1* | 0.457265734668141 | 0.00198269880061623 |
| *GM2A* | 0.311414868204686 | 0.00204501258470832 |
| *SNX4* | 0.345540652109313 | 0.00204501258470832 |

| | | |
|---|---|---|
| *LETM1* | 0.438703283641714 | 0.00205500147914412 |
| *AKAP17A* | 0.490533616046215 | 0.00218537092374019 |
| *UQCRFS1* | 0.502186481169106 | 0.00218638479397631 |
| *HNRNPA0* | 0.536305996833486 | 0.00221828621806554 |
| *TOPORS* | 0.390628541637851 | 0.00227545729521366 |
| *ATP6AP1* | 0.438853697554261 | 0.00230943738718592 |
| *TDG* | 0.410806423575481 | 0.00231121524999829 |
| *OGFRL1* | 0.491911877475032 | 0.00231946635714981 |
| *GRK2* | 0.475303294197863 | 0.00232345681086675 |
| *PSMD3* | 0.36415633027666 | 0.0023869763698199 |
| *F11R* | 0.279158677268823 | 0.00243543730265822 |
| *TENT2* | 0.278153812379438 | 0.00245705575521614 |
| *RPL7L1* | 0.478024294834939 | 0.00246793802647993 |
| *IREB2* | 0.464137211373428 | 0.00247206796430818 |
| *UBE2D3* | 0.581343592481847 | 0.00248319457632043 |
| *ARL8A* | 0.412830177012349 | 0.00250353676651529 |
| *SDHB* | 0.425556874940803 | 0.00253038377364338 |
| *ZNF280D* | 0.422213431866433 | 0.00254476554878573 |
| *DDX39B* | 0.559531200974628 | 0.00259293415992916 |
| *SCP2* | 0.491268248179999 | 0.00260767008181446 |
| *GALNT7* | 0.319068440748122 | 0.00262209880711711 |
| *ABCD4* | 0.307572801910292 | 0.00262209880711727 |
| *SFSWAP* | 0.443144626423488 | 0.00269407775382517 |
| *CNOT2* | 0.56579019144631 | 0.00272191961099165 |
| *ADGRG5* | 0.400135627382226 | 0.00276722361236371 |
| *NOP58* | 0.544728575918481 | 0.00281344040252096 |
| *AFF4* | 0.504389646782167 | 0.0028422381024435 |
| *SNW1* | 0.525619752970776 | 0.00292884624631586 |
| *BAG6* | 0.430781062164127 | 0.00296169691508005 |
| *CARD11* | 0.423862576400927 | 0.00299088921167232 |
| *MRPL54* | 0.51013563609394 | 0.0030639684278356 |
| *INTS10* | 0.479633547696596 | 0.00309756319541714 |
| *NDUFAB1* | 0.485619842762327 | 0.003109533615762 |
| *ARF3* | 0.431270526135197 | 0.00311752909077318 |
| *PIAS1* | 0.497518426008896 | 0.00313110986608915 |
| *PSMB7* | 0.437813372001969 | 0.00314544707191273 |
| *WAPL* | 0.457891768542529 | 0.00321186752244641 |
| *ILF3* | 0.536890865305796 | 0.00330766070564276 |

| | | |
|---|---|---|
| *MRPS12* | 0.345824507504402 | 0.0495012300401732 |
| *IST1* | 0.405378993003173 | 0.0496800332317121 |
| *SRGAP2* | 0.342130023946661 | 0.0499357857483282 |
