## Supplementary Table 20 for "Heterogeneity of circulating epithelial cells in breast cancer at single-cell resolution: identifying tumor and hybrid cells"

Supplementary Table 20. Signaling pathways enriched in diploid CD45$^{+}$ CEC clusters

| Cluster 1 | | |
|---|---|---|
| Term | Adjusted P-value | Genes |
| Ribosome | 2.0226979372268556E-43 | *RPL4;RPL5;RPL30;RPL3;RPL32;RPL31;RPL34;RPLP1;MRPS12;RPLP0;MRPS10;MRPL36;RPL10A;RPL8;RPL9;RPL6;RPL7;RPS15;RPS4X;RPS14;RPL7A;RPS17;RPS16;RPL18A;RPS19;RPL36AL;RPS18;RPL36;RPL35;RPLP2;RPL38;RPL37;RPS11;RPL39;RPS13;RPS12;RPS9;RPL21;RPS7;RPS8;RPL23;RPS5;RPL22;RPS6;MRPS21;RPL13A;RPSA;RPS3A;RPL37A;RPL24;RPL27;RPL26;RPL29;RPL28;UBA52;RPL10;RPL12;RPL11;RPL36A;MRPL14;RPS15A;RPL14;RPS3;RPL13;RPL15;RPS2;RPL18;RPS27A;RPL17;RPL19;RPL41;RPL35A;RPL23A;RPS26;RPS25;RPS28;RPS27;RPS29;RPL27A;RPS20;RPL22L1;FAU;RPS21;RPS24;RPS23* |
| Coronavirus disease | 4.8179836541493774E-36 | *RPL4;RPL5;NRP1;RPL30;RPL3;RPL32;RPL31;RPL34;RPLP1;RPLP0;CGAS;ADAR;RPL8;RPL10A;RPL9;RPL6;RPL7;RPS15;RPS4X;RPS14;RPL7A;RPS17;RPS16;RPL18A;RPS19;RPL36AL;RPS18;RPL36;RPL35;RPLP2;RPL38;RPL37;RPS11;RPL39;RPS13;RPS12;IFNAR2;RPS9;RPL21;SYK;RPS7;PRKCB;RPS8;RPL23;RPS5;RPL22;RPS6;CYBB;RPL13A;RPSA;RPS3A;OAS1;RPL37A;RPL24;RPL27;TLR7;RPL26;RPL29;RPL28;UBA52;IFNAR1;RPL10;RPL12;RPL11;RPL36A;RPS15A;IRAK1;RPL14;RPS3;RPL13;RPL15;RPS2;RPL18;RPS27A;RPL17;RPL19;RPL41;STAT1;STAT2;MX1;RPL35A;RPL23A;RPS26;RPS25;RPS28;RPS27;RPS29;RPL27A;RPS20;RPL22L1;FAU;RPS21;RPS24;MYD88;RPS23* |

| | | |
|---|---|---|
| Parkinson disease | 4.639675571311284E-30 | *COX7B;NDUFA13;NDUFA11;COX4I1;LRRK2;PARK7;COX6A1;COX7C;UBE2L3;PSMD9;TUBA1B;TUBB6;KIF5B;PSMD3;UQCRFS1;COX8A;TUBB;NDUFC2;SDHC;NDUFC1;SDHB;COX6B1;ERN1;PSMA6;COX7A2L;PSMA3;PSMA4;PSMA1;PSMA2;NDUFS6;VDAC3;VDAC2;UQCRC2;SLC25A5;UBA52;ATF4;SLC25A6;NDUFB8;NDUFB7;NDUFB10;UQCRB;NDUFB11;NDUFB4;ATP5MC2;ITPR1;ATP5MC3;ITPR2;NDUFB1;UQCR11;COX7A2;TXN;UQCR10;COX5B;KLC1;COX5A;PSMA7;UBE2J1;GNAI2;ATP5MC1;PSMB6;PSMB7;ATP5F1A;PSMB4;ATP5F1B;UBB;PSMB3;PSMB1;UBC;CYC1;RPS27A;ATP5PF;XBP1;NDUFA7;HSPA5;ATP5PB;NDUFA4;NDUFA3;NDUFA2;NDUFA1;UBE2G2;ATP5F1C;COX6C;ATP5F1D;ATP5F1E;UQCRQ;PSMC1;NDUFAB1;ATP5PO;GNAS;BAX;CALM2* |
| Oxidative phosphorylation | 2.276112491576161E-26 | *NDUFA13;COX7B;NDUFA11;COX4I1;COX6A1;COX7C;UQCRFS1;COX8A;ATP6V1G1;ATP6V0B;ATP6V0E1;ATP6AP1;NDUFC2;SDHC;NDUFC1;SDHB;COX6B1;COX7A2L;NDUFS6;UQCRC2;ATP6V0C;ATP6V1A;NDUFB8;NDUFB7;NDUFB10;UQCRB;NDUFB11;NDUFB4;COX17;ATP5MC2;ATP5MC3;NDUFB1;COX7A2;UQCR11;UQCR10;COX5B;COX5A;ATP5MC1;ATP5F1A;ATP5F1B;ATP6V1H;CYC1;ATP5MG;ATP5MF;ATP5ME;ATP6V0A1;ATP6V1F;ATP5PF;NDUFA7;ATP5PB;NDUFA4;NDUFA3;NDUFA2;NDUFA1;ATP5F1C;COX6C;ATP5F1D;ATP5F1E;UQCRQ;NDUFAB1;ATP5PO* |

| | | |
|---|---|---|
| Huntington disease | 5.4156438769751104E-24 | *COX7B;NDUFA13;HIP1;NDUFA11;COX4I1;CLTC;CLTA;COX6A1;COX7C;PSMD9;TUBA1B;TUBB6;SIN3A;KIF5B;CREB3L2;PSMD3;UQCRFS1;AP2M1;COX8A;TUBB;NDUFC2;SDHC;NDUFC1;SDHB;COX6B1;ERN1;PSMA6;COX7A2L;PSMA3;PSMA4;PSMA1;PSMA2;NDUFS6;VDAC3;VDAC2;ULK1;UQCRC2;SLC25A5;SLC25A6;HDAC2;NDUFB8;NDUFB7;NDUFB10;UQCRB;NDUFB11;NDUFB4;DCTN3;ATP5MC2;ITPR1;ATP5MC3;NDUFB1;UQCR11;COX7A2;UQCR10;COX5B;KLC1;COX5A;PSMA7;ATP5MC1;PSMB6;PSMB7;ATP5F1A;PSMB4;ATP5F1B;POLR2A;PSMB3;ATG101;PSMB1;AP2S1;POLR2G;CYC1;POLR2I;POLR2L;POLR2J3;ATP5PF;NDUFA7;ATP5PB;NDUFA4;NDUFA3;NDUFA2;NDUFA1;AP2B1;ATP5F1C;COX6C;ATP5F1D;ATP5F1E;REST;UQCRQ;PSMC1;GNAQ;NDUFAB1;ATP5PO;BAX* |
| Protein processing in endoplasmic reticulum | 1.7461578958954392E-23 | *ERO1B;TRAM1;HSP90AB1;UBXN1;UBE2D2;UBE2D3;NGLY1;HERPUD1;SEC61A1;SEC61G;MAN1A1;CAPN1;SEC61B;SEC62;UBXN4;UBXN6;SEC63;PDIA3;HSP90AA1;SEC13;SSR4;SSR3;RAD23A;PDIA6;DDOST;PDIA4;RBX1;ERN1;DNAJC3;DNAJC1;SELENOS;DAD1;DNAJB11;CANX;ERP29;ATF4;PPP1R15A;VCP;RPN2;DERL3;RPN1;RRBP1;RNF5;HSP90B1;UBE2J1;LMAN1;OS9;BAG1;SSR1;SEC31A;TXNDC5;BCAP31;XBP1;HSPA5;EDEM1;EIF2AK4;UBE2G2;SVIP;DNAJA1;NPLOC4;STT3A;BAX;HYOU1;STUB1;CALR;P4HB* |

| | | |
|---|---|---|
| Prion disease | 1.7461578958954392E-23 | *COX7B;NDUFA13;NCF1;NDUFA11;COX4I1;NCF4;COX6A1;COX7C;PSMD9;TUBA1B;TUBB6;KIF5B;CREB3L2;PSMD3;RAC2;UQCRFS1;RAC1;COX8A;TUBB;PRKCD;CYBB;NDUFC2;CYBA;SDHC;NDUFC1;SDHB;COX6B1;PSMA6;COX7A2L;PSMA3;PSMA4;PSMA1;PSMA2;NDUFS6;CSNK2B;VDAC3;VDAC2;UQCRC2;SLC25A5;ATF4;SLC25A6;NDUFB8;NDUFB7;NDUFB10;UQCRB;NDUFB11;NDUFB4;ATP5MC2;ITPR1;ATP5MC3;ITPR2;NDUFB1;UQCR11;COX7A2;UQCR10;COX5B;KLC1;COX5A;PSMA7;ATP5MC1;PSMB6;PSMB7;PPP3R1;ATP5F1A;PSMB4;ATP5F1B;PSMB3;PSMB1;CYC1;ATP5PF;NDUFA7;HSPA5;ATP5PB;NDUFA4;NDUFA3;NDUFA2;NDUFA1;ATP5F1C;COX6C;ATP5F1D;ATP5F1E;UQCRQ;PSMC1;NDUFAB1;ATP5PO;BAX* |
| Amyotrophic lateral sclerosis | 6.97009391020417E-23 | *NDUFA13;NDUFA11;COX6A1;ACTB;ACTG1;PSMD9;TUBB6;KIF5B;PSMD3;MAP2K3;GABARAPL2;ANXA11;SDHC;SDHB;COX6B1;ERN1;COX7A2L;SRSF3;ULK1;UQCRC2;PFN1;SRSF7;ATF4;POM121;VCP;NDUFB10;UQCRB;NDUFB11;ATP5MC2;ATP5MC3;COX7A2;KLC1;GABARAP;ATP5MC1;PPP3R1;ATG101;CYC1;MAP2K6;XBP1;HSPA5;FUS;ALYREF;SETX;UQCRQ;COX7B;COX4I1;WDR41;COX7C;TUBA1B;UQCRFS1;RAC1;RAB8A;COX8A;SEC13;TUBB;NDUFC2;NDUFC1;PSMA6;PSMA3;PSMA4;PSMA1;PSMA2;NDUFS6;CAT;NDUFB8;NDUFB7;NDUFB4;DCTN3;NDUFB1;UQCR11;UQCR10;COX5B;COX5A;PSMA7;PSMB6;PSMB7;ATP5F1A;PSMB4;ATP5F1B;PSMB3;PSMB1;BID;HNRNPA1;ATP5PF;HNRNPA3;NDUFA7;ATP5PB;NDUFA4;NDUFA3;NDUFA2;NDUFA1;ATP5F1C;COX6C;ATP5F1D;ATP5F1E;CCS;PSMC1;HNRNPA2B1;NDUFAB1;ATP5PO;BAX* |

| | | |
|---|---|---|
| Pathways of neurodegeneration | 2.947126022078476E-21 | *APP;NDUFA13;NDUFA11;PARK7;COX6A1;UBE2L3;PSMD9;TUBB6;KIF5B;PSMD3;CAPN1;MAP2K3;GABARAPL2;PRKCB;CYBB;SDHC;CSNK1E;SDHB;COX6B1;ERN1;COX7A2L;CSNK2B;VDAC3;VDAC2;ULK1;UQCRC2;UBA52;ATF4;VCP;UQCRB;NDUFB10;NDUFB11;ITPR1;ATP5MC2;ITPR2;ATP5MC3;COX7A2;PSEN1;KLC1;GABARAP;UBE2J1;ATP5MC1;PPP3R1;UBB;ATG101;UBC;CYC1;MAP2K6;XBP1;HSPA5;FUS;UQCRQ;GNAQ;CALM2;COX7B;HIP1;COX4I1;LRRK2;WDR41;ATP2A3;COX7C;TUBA1B;UQCRFS1;RAC1;RAB8A;COX8A;TUBB;NDUFC2;NDUFC1;PSMA6;PSMA3;PSMA4;PSMA1;PSMA2;NDUFS6;CAT;SLC25A5;SLC25A6;NDUFB8;NDUFB7;NDUFB4;DCTN3;NDUFB1;UQCR11;UQCR10;COX5B;COX5A;PSMA7;PSMB6;PSMB7;PSMB4;ATP5F1A;ATP5F1B;PSMB3;PSMB1;BID;RPS27A;ATP5PF;NDUFA7;CSNK1A1;NDUFA4;ATP5PB;NDUFA3;NDUFA2;NDUFA1;UBE2G2;ATP5F1C;COX6C;ATP5F1D;ATP5F1E;CCS;PSMC1;NDUFAB1;ATP5PO;CTNNB1;BAX* |
| Diabetic cardiomyopathy | 4.998070094103417E-21 | *COX7B;NDUFA13;NCF1;NDUFA11;COX4I1;NCF4;ATP2A3;PTEN;SLC2A1;COX6A1;COX7C;RAC2;UQCRFS1;CD36;RAC1;CTSD;PDK2;COX8A;PRKCB;PRKCD;CYBB;NDUFC2;CYBA;SDHC;NDUFC1;SDHB;COX6B1;TGFBR2;COX7A2L;TBC1D4;NDUFS6;VDAC3;VDAC2;UQCRC2;SLC25A5;SLC25A6;NDUFB8;NDUFB7;NDUFB10;UQCRB;NDUFB11;NDUFB4;ATP5MC2;ATP5MC3;NDUFB1;UQCR11;COX7A2;UQCR10;COX5B;COX5A;ATP5MC1;ATP5F1A;ATP5F1B;CYC1;ATP5PF;NDUFA7;ATP5PB;NDUFA4;NDUFA3;NDUFA2;NDUFA1;ATP5F1C;COX6C;ATP5F1D;ATP5F1E;PPP1CA;UQCRQ;NDUFAB1;ATP5PO* |

| | | |
|---|---|---|
| Thermogenesis | 4.85769743394285E-20 | *COX7B;NDUFA13;SMARCB1;KDM1A;NDUFA11;COX4I1;COX6A1;COX7C;ACTB;ACTG1;RPS6KA3;CREB3L2;UQCRFS1;COX8A;MAP2K3;CPT1A;SMARCC1;RPS6;NDUFC2;SDHC;NDUFC1;ARID1A;SDHB;ARID1B;COX6B1;COX7A2L;NDUFS6;UQCRC2;KDM3A;NDUFB8;NDUFB7;COX16;NDUFB10;UQCRB;NDUFB11;NDUFB4;COX17;ATP5MC2;ATP5MC3;NDUFB1;UQCR11;COX7A2;UQCR10;COX5B;COX5A;ATP5MC1;ATP5F1A;ATP5F1B;COX14;CYC1;ATP5MG;ATP5MF;ATP5ME;SMARCE1;ATP5PF;NDUFA7;ATP5PB;NDUFA4;NDUFA3;NDUFA2;NDUFA1;ATP5F1C;COX6C;ATP5F1D;ATP5F1E;RHEB;UQCRQ;NDUFAB1;ATP5PO;GNAS;NDUFAF3;GRB2;PNPLA2* |
| Alzheimer disease | 1.3015825964192904E-18 | *APP;COX7B;NDUFA13;NDUFA11;COX4I1;ATP2A3;COX6A1;COX7C;PSMD9;TUBA1B;TUBB6;KIF5B;PSMD3;UQCRFS1;CAPN1;COX8A;PSENEN;TUBB;CYBB;NDUFC2;SDHC;NDUFC1;CSNK1E;SDHB;COX6B1;ERN1;PSMA6;COX7A2L;PSMA3;PSMA4;PSMA1;PSMA2;NDUFS6;CSNK2B;VDAC3;VDAC2;ULK1;UQCRC2;SLC25A5;ATF4;SLC25A6;NDUFB8;NDUFB7;NDUFB10;UQCRB;NDUFB11;NDUFB4;ATP5MC2;ITPR1;ATP5MC3;ITPR2;NDUFB1;UQCR11;COX7A2;UQCR10;PSEN1;COX5B;KLC1;COX5A;PSMA7;RTN4;ATP5MC1;APH1A;PSMB6;PSMB7;PPP3R1;ATP5F1A;PSMB4;ATP5F1B;PSMB3;ATG101;PSMB1;CYC1;BID;ATP5PF;XBP1;NDUFA7;ATP5PB;CSNK1A1;NDUFA4;NDUFA3;NDUFA2;NDUFA1;ATP5F1C;COX6C;ATP5F1D;ATP5F1E;UQCRQ;PSMC1;GNAQ;NDUFAB1;ATP5PO;CTNNB1;CALM2* |
| Spliceosome | 7.076249266804252E-13 | *SF3B5;DDX5;RBM25;SF3B2;EIF4A3;HNRNPU;SNU13;PRPF8;SNRPD2;SNRPD1;PCBP1;TRA2B;TXNL4A;HNRNPA1;SRSF10;CTNNBL1;PRPF38B;HNRNPA3;CCDC12;FUS;ALYREF;PRPF40A;THOC2;WBP11;LSM4;CDC40;LSM3;U2SURP;HNRNPM;LSM7;HNRNPK;SNW1;DDX39B;SNRPG;SRSF2;SNRNP27;SRSF3;SNRPE;SRSF4;SNRPF;SRSF5;HNRNPC;SNRPC;SRSF6;SRSF7;SNRPB;SRSF9* |

| | | |
|---|---|---|
| Phagosome | 5.090912440825359E-12 | *ATP6V1A;RAB5C;NCF1;TFRC;NCF4;CORO1A;ACTB;CTSS;ACTG1;SEC61A1;TUBB6;TUBA1B;HLA-DMA;HLA-DMB;LAMP1;SEC61G;LAMP2;ATP6V1H;CD36;RAC1;SEC61B;HLA-DOA;HLA-DQA1;ATP6V0A1;ATP6V1F;HLA-DPA1;DYNC1H1;ATP6V1G1;ATP6V0B;ATP6V0E1;ATP6AP1;TUBB;STX7;HLA-B;TAP2;M6PR;CYBB;CYBA;CANX;HLA-DPB1;HLA-DRA;CALR;ATP6V0C;HLA-DRB1;HLA-DQB1;RAB7A* |
| Non-alcoholic fatty liver disease | 1.0416045306239778E-11 | *NDUFA13;COX7B;NDUFB8;NDUFB7;NDUFA11;NDUFB10;UQCRB;NDUFB11;COX4I1;NDUFB4;NDUFB1;COX7A2;UQCR11;UQCR10;COX6A1;COX5B;COX7C;COX5A;BCL2L11;UQCRFS1;CYC1;RAC1;BID;COX8A;MLXIP;XBP1;NDUFA7;NDUFA4;NDUFA3;NDUFA2;NDUFA1;NDUFC2;SDHC;NDUFC1;COX6C;SDHB;COX6B1;ERN1;COX7A2L;ITCH;UQCRQ;NDUFS6;NDUFAB1;BAX;UQCRC2;ATF4* |
| Lysosome | 2.3078967987686187E-10 | *SCARB2;ASAH1;HEXB;CLTC;CTSZ;CLTA;GNS;CTSS;GGA2;LAPTM4A;GNPTAB;LAMP1;GM2A;LAMP2;PSAP;AP1S2;ATP6V1H;CTSH;AP1S3;AP3S1;CTSD;CTSC;ATP6V0A1;CTSB;CD164;ATP6V0B;ATP6AP1;M6PR;LAPTM5;NAGA;IGF2R;GNPTG;NPC1;NPC2;MAN2B1;TPP1;CD68;ATP6V0C;LGMN* |
| Bacterial invasion of epithelial cells | 1.6601690566365484E-9 | *SRC;ARPC1B;CLTC;ARPC5L;CLTA;ACTB;SEPTIN11;ACTG1;CDH1;CTNNA1;RAC1;WASF2;ACTR2;SEPTIN2;GAB1;SEPTIN6;ARPC4;ARPC5;SEPTIN9;RHOA;CD2AP;DNM2;MAD2L2;ARPC2;ARPC3;ELMO1;HCLS1;CTNNB1* |
| Antigen processing and presentation | 1.2212407356706635E-8 | *CIITA;HSP90AB1;CTSS;HLA-DMA;HLA-DMB;HLA-DOA;B2M;HLA-DQA1;CTSB;HLA-DPA1;PDIA3;CD74;HSP90AA1;HSPA5;HLA-B;TAP2;TAPBP;CD4;CANX;PSME1;HLA-DPB1;PSME2;HLA-DRA;CALR;HLA-DRB1;HLA-DQB1;LGMN* |

| | | |
|---|---|---|
| Salmonella infection | 4.492583969214784E-8 | *CYFIP2;ARF1;HSP90AB1;ARPC1B;ARPC5L;BRK1;ACTB;PIK3CG;ACTG1;PYCARD;TUBA1B;TUBB6;KIF5B;KPNA4;RAC1;MAP2K3;ACTR2;HSP90AA1;ANXA2;TUBB;DYNLL1;RHOA;DNM2;TLR9;ELMO1;PFN1;ARL8A;RAB7A;RAB5C;DCTN3;TXN;KLC1;MYL12A;MYL12B;HSP90B1;CYTH4;IRAK1;RPS3;NCKAP1L;SNX9;FLNB;MAP2K6;DYNC1H1;M6PR;ARPC4;ARPC5;SNX18;ARPC2;ARPC3;CTNNB1;PLEKHM2;BAX;DYNLRB1;MYD88* |
| Endocytosis | 6.674218975361407E-8 | *ARF3;WASHC4;VPS29;ARF4;ARF1;TFRC;SH3KBP1;ARPC1B;CLTC;ARPC5L;CLTA;CAPZB;KIF5B;VPS36;GIT1;RAB8A;AP2M1;SH3GLB1;USP8;ACTR2;HLA-B;ARFGAP3;RHOA;EPN1;DNM2;TGFBR2;ACAP2;RABEP1;CHMP4B;CHMP3;ARF5;RAB7A;RAB5C;SRC;SNX3;SNX4;GRK3;RAB11FIP1;SNX2;GRK2;CYTH4;AP2S1;GIT2;IQSEC1;ARPC4;SNF8;AP2B1;ARPC5;IGF2R;ITCH;DAB2;ARPC2;ARPC3;CAPZA1* |
| RNA transport | 7.397326135246779E-8 | *CYFIP2;EIF4A1;POM121;EIF4A3;FXR1;PNN;NUP42;SUMO3;SUMO2;EIF4EBP1;SAP18;EIF4EBP3;EIF4B;PAIP1;SEC13;UBE2I;EIF1AX;FUS;ALYREF;PABPC4;UPF3A;THOC2;EIF2S2;SENP2;SRRM1;EIF1;EEF1A1;EIF2S3;EIF5;DDX39B;EIF3I;EIF3G;EIF3H;RNPS1;EIF3E;PABPC1;EIF3F;EIF1B;EIF4G3;EIF3D;EIF4G2;EIF3A;RAN;EIF4G1* |
| Vibrio cholerae infection | 8.997428017240305E-8 | *ATP6V1A;SLC12A2;ARF1;ATP6V0B;ATP6V1G1;ATP6V0E1;ATP6AP1;KDELR1;ACTB;ACTG1;PDIA4;SEC61A1;SEC61G;GNAS;ATP6V1H;KDELR2;SEC61B;ATP6V0C;ATP6V0A1;ATP6V1F* |
| Protein export | 2.3215643612147268E-7 | *HSPA5;SRP14;SEC61A1;SPCS3;SPCS2;SPCS1;SRP72;SEC61G;SRPRA;SEC61B;SEC62;SEC11C;SEC63* |
| Epstein-Barr virus infection | 3.1058277879993843E-7 | *CDKN1A;HDAC2;ICAM1;HLA-DMA;HLA-DMB;BCL2L11;IRAK1;SIN3A;PSMD3;BLNK;CIR1;RAC1;BID;HLA-DOA;B2M;HLA-DQA1;HLA-DPA1;MAP2K6;LYN;IFNAR2;MAP2K3;PDIA3;USP7;GADD45B;SYK;STAT1;STAT2;HLA-B;TAP2;TAPBP;NCOR2;SNW1;OAS1;PSMC1;BTK;HLA-DPB1;IRF7;HLA-DRA;BAX;CALR;VIM;HLA-DRB1;MYD88;HLA-DQB1;IFNAR1* |

| | | |
|---|---|---|
| Proteasome | 6.769619270411246E-7 | *POMP;PSMA7;PSMB10;PSMB6;PSMB7;PSMD9;PSMA6;PSMB4;PSMA3;PSMA4;PSMA1;PSMB3;PSMA2;PSMC1;PSMB1;PSMD3;PSME1;PSME2* |
| Shigellosis | 1.1242368196699213E-6 | *ARF1;ARPC1B;UBE2D2;UBE2D3;ARPC5L;CGAS;ACTB;SEPTIN11;ACTG1;PYCARD;CAPNS1;RAC1;CAPN1;GABARAPL2;ACTR2;FNBP1;FBXW11;PRKCD;ACTN4;RHOA;RBX1;RRAGC;ELMO1;HCLS1;RBCK1;UBE2V1;PFN1;TLN1;UBA52;SRC;ITPR1;ITPR2;GABARAP;MYL12A;MYL12B;CYTH4;UBB;UBC;RPS27A;WASF2;SEPTIN2;SEPTIN6;ARPC4;ARPC5;SEPTIN9;ARPC2;TNIP1;ARPC3;BAX;MYD88* |
| Viral myocarditis | 1.2772359604600269E-5 | *HLA-B;ACTB;ACTG1;ICAM1;HLA-DMA;HLA-DMB;RAC2;HLA-DPB1;HLA-DRA;RAC1;HLA-DOA;BID;EIF4G3;HLA-DQA1;EIF4G2;HLA-DRB1;HLA-DPA1;HLA-DQB1;EIF4G1* |
| Tuberculosis | 1.3174638916066332E-5 | *CIITA;RAB5C;SRC;LSP1;CORO1A;CTSS;PPP3R1;HLA-DMA;HLA-DMB;LAMP1;IRAK1;LAMP2;ATP6V1H;HLA-DOA;BID;CTSD;HLA-DQA1;ATP6V0A1;HLA-DPA1;HSPA9;CD74;ATP6V0B;FCER1G;ATP6AP1;SYK;STAT1;IL10RA;RHOA;TLR9;HLA-DPB1;HLA-DRA;BAX;CALM2;ATP6V0C;HLA-DRB1;MYD88;HLA-DQB1;RAB7A* |
| Cardiac muscle contraction | 3.175627288272789E-5 | *COX8A;COX7B;UQCRB;TPM3;COX4I1;TPM2;ATP2A3;COX7A2;UQCR11;UQCR10;COX6A1;COX5B;COX6C;COX7C;COX5A;COX6B1;COX7A2L;ASPH;SLC9A7;UQCRQ;UQCRFS1;UQCRC2;CYC1* |
| Human immunodeficiency virus 1 infection | 2.793468502806305E-4 | *ITPR1;ITPR2;CGAS;RNF7;SAMHD1;GNAI2;GNG10;PPP3R1;IRAK1;GNG5;GNG7;CFL1;AP1S2;RAC2;ELOB;AP1S3;RAC1;BID;B2M;PAK2;MAP2K6;MAP2K3;PDIA3;PRKCB;FBXW11;HLA-B;TAP2;RBX1;TAPBP;BST2;DDB1;CD4;GNAQ;GNB2;GNB1;BAX;CALR;CALM2;MYD88* |
| Spinocerebellar ataxia | 4.4603427059352463E-4 | *ITPR1;ATP2A3;ITPR2;PSMA7;PSMB6;PSMD9;PSMB7;PSMB4;ATG101;PSMB3;PSMD3;PSMB1;NOP56;XBP1;PRKCB;PUM1;ERN1;PSMA6;PSMA3;PSMA4;PSMA1;PSMA2;GNAQ;PSMC1;VDAC3;VDAC2;ULK1;SLC25A5;SLC25A6* |

| | | |
|---|---|---|
| Influenza A | 4.4603427059352463E-4 | *CIITA;ADAR;ACTB;ACTG1;ICAM1;PYCARD;HLA-DMA;HLA-DMB;PABPN1;HLA-DOA;KPNA2;BID;HLA-DQA1;HLA-DPA1;IFNAR2;PRKCB;STAT1;STAT2;MX1;DNAJC3;HNRNPUL1;OAS1;IRF7;HLA-DPB1;HLA-DRA;BAX;TLR7;SLC25A5;HLA-DRB1;MYD88;IFNAR1;HLA-DQB1;SLC25A6* |
| Autophagy | 4.8771320491224464E-4 | *MTMR14;ITPR1;PTEN;WDR41;HMGB1;GABARAP;LAMP1;ATG101;LAMP2;CTSD;SNAP29;RAB8A;CTSB;SH3GLB1;GABARAPL2;UVRAG;RUBCN;DAPK1;PDPK1;DAPK2;PRKCD;EIF2AK4;ERN1;VAMP8;RRAGC;RHEB;ULK1;RAB7A* |
| Ferroptosis | 4.8771320491224464E-4 | *TFRC;GPX4;NCOA4;CYBB;SLC3A2;SLC7A11;SAT1;FTH1;PCBP1;PCBP2;VDAC3;VDAC2;FTL* |
| Asthma | 5.540222382991184E-4 | *HLA-DMA;HLA-DMB;FCER1G;HLA-DPB1;FCER1A;HLA-DRA;HLA-DOA;HLA-DQA1;HLA-DRB1;HLA-DPA1;HLA-DQB1* |
| Rheumatoid arthritis | 8.621878492173493E-4 | *ATP6V1A;ATP6V0B;ATP6V1G1;ATP6V0E1;ATP6AP1;TNFSF13;ICAM1;TNFSF13B;HLA-DMA;HLA-DMB;HLA-DPB1;ATP6V1H;HLA-DRA;HLA-DOA;ATP6V0C;HLA-DQA1;HLA-DRB1;HLA-DPA1;HLA-DQB1;ATP6V0A1;ATP6V1F* |
| Mitophagy | 0.0010033019779792749 | *GABARAPL2;USP8;FIS1;USP15;SRC;GABARAP;UBB;CSNK2B;UBC;TAX1BP1;TOMM7;PGAM5;ULK1;RPS27A;UBA52;ATF4;RAB7A* |
| Leishmaniasis | 0.0015707217619922616 | *NCF1;PRKCB;STAT1;NCF4;CYBB;CYBA;EEF1A1;HLA-DMA;HLA-DMB;IRAK1;HLA-DPB1;HLA-DRA;HLA-DOA;HLA-DQA1;HLA-DRB1;MYD88;HLA-DPA1;HLA-DQB1* |
| Ubiquitin mediated proteolysis | 0.0015715065218439885 | *UBE2D2;UBE2D3;UBE3A;RNF7;UBE2L3;ANAPC11;UBE2J1;UBB;UBC;ELOB;RPS27A;UBE2I;FBXW11;UBE2E3;UBE2E2;UBE2G2;PIAS1;RBX1;DDB1;ITCH;CDC16;ANAPC5;BIRC6;STUB1;TRIP12;UBA52;UBE2K* |
| Kaposi sarcoma-associated herpesvirus infection | 0.0015807922712062167 | *CDKN1A;SRC;ITPR1;ITPR2;GABARAP;PIK3CG;ICAM1;GNG10;PPP3R1;ZFP36;UBB;GNG5;GNG7;UBC;RAC1;BID;RPS27A;MAP2K6;LYN;IFNAR2;GABARAPL2;SYK;STAT1;STAT2;HLA-B;GNB2;MAPKAPK2;GNB1;IRF7;CTNNB1;BAX;UBA52;CALM2;IFNAR1* |

| | | |
|---|---|---|
| Adherens junction | 0.0015946873678581621 | *FARP2;PTPN1;SRC;ACTN4;IQGAP1;ACTB;RHOA;ACTG1;TGFBR2;CDH1;CSNK2B;RAC2;CTNNA1;CTNNB1;RAC1;WASF2;NECTIN1* |
| Human T-cell leukemia virus 1 infection | 0.0019330912712461337 | *NRP1;CDKN1A;PTEN;SLC2A1;ICAM1;ANAPC11;POLB;PPP3R1;HLA-DMA;ZFP36;HLA-DMB;CREB3L2;HLA-DOA;B2M;HLA-DQA1;HLA-DPA1;RANBP1;HLA-B;TGFBR2;CD4;CDC16;CANX;VDAC3;HLA-DPB1;VDAC2;HLA-DRA;BAX;ANAPC5;CALR;TCF3;TLN1;SLC25A5;HLA-DRB1;RAN;ATF4;HLA-DQB1;SLC25A6* |
| Yersinia infection | 0.00241004587203927 | *SRC;ARPC1B;ARPC5L;ACTB;ACTG1;PYCARD;RPS6KA3;IRAK1;RAC2;RAC1;WASF2;MAP2K6;MAP2K3;VAV3;ACTR2;GIT2;ARPC4;ARPC5;RHOA;CD4;ARPC2;ARPC3;GNAQ;ELMO1;MYD88;SKAP2* |
| Human cytomegalovirus infection | 0.00316521395364021 | *CDKN1A;SRC;ITPR1;ITPR2;CGAS;GNAI2;AKAP13;GNG10;PPP3R1;GNG5;GNG7;CREB3L2;GNA12;RAC2;EIF4EBP1;RAC1;BID;B2M;MAP2K6;PDIA3;PRKCB;IL10RA;HLA-B;TAP2;RHOA;TAPBP;RHEB;GNAQ;GNB2;GNB1;GNAS;CTNNB1;BAX;GRB2;CALR;CALM2;ATF4* |
| Allograft rejection | 0.003325844460619528 | *HLA-DMA;HLA-DMB;HLA-B;HLA-DPB1;GZMB;HLA-DRA;HLA-DOA;HLA-DQA1;HLA-DRB1;HLA-DPA1;HLA-DQB1* |
| Retrograde endocannabinoid signaling | 0.0033641146495805786 | *NDUFA13;NDUFB8;NDUFB7;NDUFA11;NDUFB10;NDUFB11;NDUFB4;ITPR1;ITPR2;NDUFB1;GNAI2;GNG10;GNG5;GNG7;NDUFA7;PRKCB;NDUFA4;NDUFA3;NDUFA2;NDUFA1;NDUFC2;NDUFC1;NDUFS6;GNAQ;GNB2;GNB1;NDUFAB1* |
| Fc gamma R-mediated phagocytosis | 0.0034260629401408845 | *LYN;VAV3;ACTR2;GSN;NCF1;SYK;PRKCB;ARPC1B;PRKCD;ARPC5L;ARPC4;ARPC5;GAB2;DNM2;ARPC2;ARPC3;CFL1;RAC2;RAC1;WASF2* |
| Viral carcinogenesis | 0.00344645097332284 | *YWHAE;HDAC5;CDKN1A;GTF2A2;HDAC2;DDX3X;YWHAB;SRC;UBE3A;HDAC9;POLB;CREB3L2;RAC1;LYN;RANBP1;USP7;GSN;SYK;HLA-B;ACTN4;YWHAZ;SND1;RHOA;DDB1;HNRNPK;SNW1;PSMC1;MAPKAPK2;VDAC3;REL;IRF7;BAX;GRB2;ATF4* |

| | | |
|---|---|---|
| Various types of N-glycan biosynthesis | 0.0039001691834723573 | *B4GALT1;RPN2;MAN2A1;DAD1;HEXB;RPN1;ALG2;STT3A;MAN1A1;MGAT1;DDOST* |
| Leukocyte transendothelial migration | 0.004526382512086873 | *VAV3;NCF1;PRKCB;NCF4;CYBB;CYBA;ACTN4;F11R;ACTB;MYL12A;RHOA;ACTG1;ICAM1;MYL12B;GNAI2;RASSF5;RAC2;PECAM1;CTNNA1;CTNNB1;RAC1;CD99* |
| Intestinal immune network for IgA production | 0.0070188118011436824 | *HLA-DMA;HLA-DMB;TNFSF13;HLA-DPB1;HLA-DRA;HLA-DOA;HLA-DQA1;ICOSLG;HLA-DRB1;TNFSF13B;HLA-DPA1;HLA-DQB1* |
| Graft-versus-host disease | 0.007235581235241142 | *HLA-DMA;HLA-DMB;HLA-B;HLA-DPB1;GZMB;HLA-DRA;HLA-DOA;HLA-DQA1;HLA-DRB1;HLA-DPA1;HLA-DQB1* |
| Pathogenic Escherichia coli infection | 0.007553507555370403 | *CYFIP2;ARF1;TMED10;SRC;ARPC1B;ARPC5L;BRK1;ACTB;ACTG1;PYCARD;TUBA1B;TUBB6;CYTH4;IRAK1;GNA12;RPS3;NCKAP1L;RAC1;PAK2;WASF2;ACTR2;TUBB;ARPC4;ARPC5;RHOA;ARPC2;ARPC3;NCL;HCLS1;BAX;TMBIM6;MYD88* |
| Type I diabetes mellitus | 0.008586205254653178 | *HLA-DMA;HLA-DMB;HLA-B;HLA-DPB1;GZMB;HLA-DRA;HLA-DOA;HLA-DQA1;HLA-DRB1;HLA-DPA1;HLA-DQB1* |
| Fluid shear stress and atherosclerosis | 0.012160252943115464 | *MEF2A;MEF2C;HSP90AA1;HSP90AB1;NCF1;SRC;GSTP1;MGST2;CYBA;TXN;ACTB;RHOA;ACTG1;ICAM1;HSP90B1;THBD;SUMO3;SUMO2;RAC2;PECAM1;CTNNB1;RAC1;CALM2;MAP2K6* |
| Platelet activation | 0.012578463045481842 | *LYN;FCER1G;SYK;SRC;ITPR1;ITPR2;ACTB;MYL12A;PIK3CG;RHOA;ACTG1;MYL12B;GNAI2;PPP1CA;APBB1IP;VAMP8;GNAQ;P2RX1;BTK;GNAS;TLN1;FERMT3* |
| B cell receptor signaling pathway | 0.01539882988524994 | *VAV3;LYN;SYK;PRKCB;LILRB1;LILRB2;LILRB4;LILRA4;PPP3R1;BLNK;RAC2;BTK;GRB2;RAC1;PIK3AP1;CARD11* |
| Apoptosis | 0.01539882988524994 | *GADD45B;PDPK1;CTSZ;ITPR1;ITPR2;GZMB;CSF2RB;ACTB;CTSS;ACTG1;ERN1;TUBA1B;BCL2L11;IL3RA;CTSH;BAX;BIRC6;CAPN1;BID;CTSD;CTSC;ATF4;MCL1;CTSB* |
| Osteoclast differentiation | 0.016216066343604896 | *IFNAR2;NCF1;SYK;STAT1;STAT2;NCF4;CYBA;LILRB1;GAB2;LILRB2;LILRB4;SIRPB1;LILRA4;TGFBR2;PPP3R1;TYROBP;BTK;BLNK;GRB2;RAC1;IFNAR1;MAP2K6* |
| Fc epsilon RI signaling pathway | 0.017697083205037815 | *MAP2K3;VAV3;LYN;FCER1G;SYK;PDPK1;GAB2;ALOX5AP;RAC2;BTK;FCER1A;GRB2;RAC1;MAP2K6* |

| | | |
|---|---|---|
| mRNA surveillance pathway | 0.01886190843797378 | *FUS;ALYREF;PABPC4;EIF4A3;UPF3A;SMG7;SRRM1;PPP1CA;PNN;DDX39B;FIP1L1;PABPN1;PCF11;PAPOLA;SAP18;RNPS1;ETF1;PABPC1* |
| Epithelial cell signaling in Helicobacter pylori infection | 0.022543289761284665 | *ATP6V1A;LYN;ATP6V0B;ATP6V1G1;ATP6V0E1;ATP6AP1;SRC;F11R;ATP6V1H;RAC1;ATP6V0C;GIT1;ATP6V0A1;ATP6V1F* |
| Synaptic vesicle cycle | 0.024291502850226902 | *ATP6V1A;NAPA;ATP6V0B;ATP6V1G1;ATP6V0E1;CLTC;CLTA;AP2B1;DNM2;AP2S1;ATP6V1H;ATP6V0C;AP2M1;ATP6V0A1;ATP6V1F* |
| NOD-like receptor signaling pathway | 0.024385325015092465 | *YWHAE;HSP90AB1;ITPR1;ITPR2;TXN;ANTXR2;GABARAP;PYCARD;TRPM2;NAMPT;CTSB;IFNAR2;GABARAPL2;HSP90AA1;STAT1;STAT2;PRKCD;ERBIN;CYBB;CYBA;RHOA;OAS1;VDAC3;IRF7;VDAC2;RBCK1;MYD88;IFNAR1* |
| RNA degradation | 0.026620089400521848 | *HSPA9;TTC37;BTG2;DIS3;PABPC4;LSM1;ENO1;LSM4;LSM3;LSM7;CNOT7;CNOT1;CNOT2;PABPC1;DCPS* |
| Regulation of actin cytoskeleton | 0.03002428181885416 | *CYFIP2;SRC;ARPC1B;ARPC5L;BRK1;IQGAP1;ITGAE;IQGAP2;ACTB;MYL12A;MYL12B;ACTG1;CFL1;GNA12;RAC2;NCKAP1L;RAC1;PAK2;WASF2;GIT1;VAV3;ACTR2;GSN;ARPC4;ACTN4;ARPC5;RHOA;PPP1CA;ABI2;ARPC2;ARPC3;PFN1* |
| Toxoplasmosis | 0.03300029708245039 | *MAP2K3;CIITA;PDPK1;STAT1;IL10RA;PIK3CG;GNAI2;HLA-DMA;HLA-DMB;IRAK1;HLA-DPB1;HLA-DRA;HLA-DOA;HLA-DQA1;HLA-DRB1;MYD88;MAP2K6;HLA-DPA1;HLA-DQB1* |
| Thyroid hormone signaling pathway | 0.035834993690714106 | *NOTCH2;PFKFB2;NCOA1;HDAC2;PRKCB;PDPK1;SRC;STAT1;NOTCH4;NCOA3;ATP2A3;SLC2A1;ACTB;ACTG1;MED13L;MED30;TBC1D4;RHEB;SIN3A;CTNNB1* |
| Autoimmune thyroid disease | 0.037644994715885755 | *HLA-DMA;HLA-DMB;HLA-B;HLA-DPB1;GZMB;HLA-DRA;HLA-DOA;HLA-DQA1;HLA-DRB1;HLA-DPA1;HLA-DQB1* |
| Cellular senescence | 0.04209442079268754 | *MAP2K3;CDKN1A;GADD45B;FBXW11;HLA-B;ITPR1;PTEN;ITPR2;TGFBR2;PPP1CA;PPP3R1;RBBP4;RHEB;RASSF5;MAPKAPK2;VDAC3;EIF4EBP1;VDAC2;MYBL2;CAPN1;SLC25A5;CALM2;MAP2K6;SLC25A6* |

| | | |
|---|---|---|
| Cell adhesion molecules | 0.04251197858639292 | *SELPLG;HLA-B;F11R;GLG1;ICAM1;HLA-DMA;CD4;ALCAM;HLA-DMB;SELL;CDH1;PECAM1;HLA-DPB1;HLA-DRA;HLA-DOA;ICOSLG;CD99;HLA-DQA1;HLA-DRB1;MPZL1;HLA-DPA1;HLA-DQB1;NECTIN1* |
| Cluster 2 | | |
| Ribosome | 3.2519522397822318E-53 | *RPL4;RPL5;RPL30;RPL3;RPL32;RPL31;RPL34;RPLP1;RPLP0;MRPS10;MRPL36;RPL10A;RPL8;RPL9;RPL6;RPL7;RPS15;RPS4X;RPS14;RPL7A;RPS17;RPS16;RPL18A;RPS19;RPL36AL;RPS18;RPL35;RPLP2;RPL38;RPL37;RPS11;RPL39;RPS10;RPS13;RPS12;RPS9;RPL21;RPS7;RPS8;RPL23;RPS5;RPL22;RPS6;MRPS21;RPL13A;RPSA;RPS3A;RPL37A;RPL24;RPL27;RPL26;RPL29;RPL28;UBA52;RPL10;RPL12;RPL11;RPL36A;RPS15A;RPL14;RPS3;RPL13;RPL15;RPS2;RPL18;RPS27A;RPL17;RPL19;RPL41;RPL35A;RPL23A;RPS26;RPS25;RPS28;RPS27;RPS29;RPL27A;RPS20;RPL22L1;FAU;RPS21;RPS24;RPS23* |
| Coronavirus disease | 4.6937108782575656E-45 | *RPL4;RPL5;NRP1;RPL30;RPL3;RPL32;RPL31;RPL34;RPLP1;RPLP0;RPL8;RPL10A;RPL9;RPL6;RPL7;RPS15;RPS4X;RPS14;RPL7A;RPS17;RPS16;RPL18A;RPS19;RPL36AL;RPS18;RPL35;RPLP2;RPL38;RPL37;RPS11;RPL39;RPS10;RPS13;RPS12;IFNAR2;RPS9;RPL21;SYK;RPS7;PRKCB;RPS8;RPL23;RPS5;RPL22;RPS6;CYBB;RPL13A;RPSA;RPS3A;OAS1;RPL37A;RPL24;RPL27;TLR7;RPL26;RPL29;RPL28;UBA52;IFNAR1;RPL10;RPL12;RPL11;RPL36A;RPS15A;RPL14;RPS3;RPL13;RPL15;RPS2;RPL18;RPS27A;RPL17;RPL19;RPL41;STAT2;MX1;RPL35A;RPL23A;RPS26;RPS25;RPS28;RPS27;RPS29;RPL27A;RPS20;RPL22L1;FAU;RPS21;RPS24;MYD88;RPS23* |

| | | |
|---|---|---|
| Parkinson disease | 5.609408004748859E-25 | *COX7B;NDUFA13;NDUFA11;COX4I1;PARK7;COX6A1;COX7C;UBE2L3;TUBA1B;TUBB6;KIF5B;COX8A;TUBB;NDUFC2;SDHC;NDUFC1;COX6B1;ERN1;PSMA6;COX7A2L;PSMA3;PSMA4;PSMA2;NDUFS6;SLC25A5;UBA52;SLC25A6;NDUFB8;NDUFB7;NDUFB10;UQCRB;NDUFB11;NDUFB4;ATP5MC2;ITPR1;ATP5MC3;ITPR2;NDUFB1;UQCR11;COX7A2;TXN;UQCR10;KLC1;COX5A;PSMA7;UBE2J1;GNAI2;ATP5MC1;PSMB6;ATP5F1A;ATP5F1B;UBB;PSMB3;PSMB1;UBC;CYC1;RPS27A;ATP5PF;XBP1;NDUFA7;ATP5PB;NDUFA4;NDUFA1;UBE2G2;ATP5F1C;COX6C;ATP5F1D;ATP5F1E;UQCRQ;GNAS;CALM2* |
| Oxidative phosphorylation | 1.0756552419327067E-21 | *NDUFA13;COX7B;NDUFB8;NDUFB7;NDUFA11;NDUFB10;UQCRB;NDUFB11;COX4I1;NDUFB4;ATP5MC2;ATP5MC3;NDUFB1;COX7A2;UQCR11;UQCR10;COX6A1;COX7C;COX5A;ATP5MC1;ATP5F1A;ATP5F1B;CYC1;ATP5MG;ATP5MF;ATP5ME;ATP6V1F;COX8A;ATP5PF;ATP6V1G1;ATP6V0B;NDUFA7;ATP6V0E1;ATP5PB;NDUFA4;NDUFA1;NDUFC2;SDHC;NDUFC1;ATP5F1C;COX6C;ATP5F1D;ATP5F1E;COX6B1;COX7A2L;UQCRQ;NDUFS6;ATP6V0C* |
| Huntington disease | 1.8939987575879193E-19 | *COX7B;NDUFA13;HIP1;NDUFA11;COX4I1;CLTC;CLTB;CLTA;COX6A1;COX7C;TUBA1B;TUBB6;KIF5B;CREB3L2;AP2M1;COX8A;TUBB;GPX7;NDUFC2;SDHC;NDUFC1;COX6B1;ERN1;PSMA6;COX7A2L;PSMA3;PSMA4;PSMA2;NDUFS6;ULK1;SLC25A5;SLC25A6;NDUFB8;NDUFB7;NDUFB10;UQCRB;NDUFB11;NDUFB4;ATP5MC2;ITPR1;ATP5MC3;NDUFB1;UQCR11;COX7A2;UQCR10;KLC1;COX5A;PSMA7;ATP5MC1;PSMB6;ATP5F1A;ATP5F1B;PSMB3;ATG101;PSMB1;AP2S1;CYC1;POLR2I;POLR2L;POLR2J3;ATP5PF;NDUFA7;ATP5PB;NDUFA4;NDUFA1;ATP5F1C;COX6C;ATP5F1D;ATP5F1E;UQCRQ;GNAQ* |

| | | |
|---|---|---|
| Prion disease | 3.8398037243565913E-19 | *COX7B;NDUFA13;NCF1;NDUFA11;COX4I1;COX6A1;COX7C;TUBA1B;TUBB6;KIF5B;CREB3L2;RAC2;RAC1;COX8A;TUBB;PRKCD;CYBB;NDUFC2;CYBA;SDHC;NDUFC1;COX6B1;PSMA6;COX7A2L;PSMA3;PSMA4;PSMA2;NDUFS6;CSNK2B;SLC25A5;SLC25A6;NDUFB8;NDUFB7;NDUFB10;UQCRB;NDUFB11;NDUFB4;ATP5MC2;ITPR1;ATP5MC3;ITPR2;NDUFB1;UQCR11;COX7A2;UQCR10;KLC1;COX5A;PSMA7;ATP5MC1;PSMB6;PPP3R1;ATP5F1A;ATP5F1B;PSMB3;PSMB1;CYC1;ATP5PF;NDUFA7;ATP5PB;NDUFA4;NDUFA1;ATP5F1C;COX6C;ATP5F1D;ATP5F1E;UQCRQ* |
| Amyotrophic lateral sclerosis | 1.0771289762735564E-18 | *COX7B;NDUFA13;NDUFA11;COX4I1;COX6A1;COX7C;ACTB;ACTG1;TUBA1B;TUBB6;KIF5B;RAC1;COX8A;MAP2K3;GABARAPL2;TUBB;ANXA11;GPX7;NDUFC2;SDHC;NDUFC1;COX6B1;ERN1;PSMA6;COX7A2L;PSMA3;PSMA4;PSMA2;NDUFS6;CAT;ULK1;PFN1;SRSF7;NDUFB8;NDUFB7;NDUFB10;UQCRB;NDUFB11;NDUFB4;ATP5MC2;ATP5MC3;NDUFB1;UQCR11;COX7A2;UQCR10;KLC1;COX5A;GABARAP;PSMA7;ATP5MC1;PSMB6;PPP3R1;ATP5F1A;ATP5F1B;PSMB3;ATG101;PSMB1;CYC1;BID;HNRNPA1;MAP2K6;ATP5PF;XBP1;HNRNPA3;NDUFA7;ATP5PB;FUS;NDUFA4;NDUFA1;CHCHD10;ATP5F1C;COX6C;ATP5F1D;ATP5F1E;CCS;UQCRQ;HNRNPA2B1* |
| Diabetic cardiomyopathy | 1.8421552486129345E-18 | *COX7B;NDUFA13;NCF1;NDUFA11;COX4I1;ATP2A3;SLC2A1;COX6A1;COX7C;RAC2;CD36;RAC1;PDK2;COX8A;PRKCB;PRKCD;CYBB;NDUFC2;CYBA;SDHC;NDUFC1;COX6B1;TGFBR2;COX7A2L;TBC1D4;NDUFS6;SLC25A5;SLC25A6;NDUFB8;NDUFB7;NDUFB10;UQCRB;NDUFB11;NDUFB4;ATP5MC2;ATP5MC3;NDUFB1;UQCR11;COX7A2;UQCR10;COX5A;ATP5MC1;ATP5F1A;ATP5F1B;CYC1;ATP5PF;NDUFA7;ATP5PB;NDUFA4;NDUFA1;ATP5F1C;COX6C;ATP5F1D;ATP5F1E;UQCRQ* |

| | | |
|---|---|---|
| Protein processing in endoplasmic reticulum | 2.2617910089225946E-18 | *TRAM1;HSP90AB1;UBE2D2;NGLY1;HERPUD1;SEC61A1;SEC61G;MAN1A1;SEC61B;SEC62;UBXN4;SEC63;PDIA3;HSP90AA1;SSR4;SSR3;RAD23A;PDIA6;DDOST;PDIA4;RBX1;ERN1;DNAJC3;SELENOS;DAD1;CANX;ERP29;RPN2;DERL3;RPN1;RRBP1;RNF5;HSP90B1;UBE2J1;LMAN1;OS9;BAG1;SSR1;SEC31A;TXNDC5;BCAP31;XBP1;EDEM1;EIF2AK4;UBE2G2;DNAJA1;HYOU1;STUB1;CALR;P4HB* |
| Pathways of neurodegeneration | 3.982605144139458E-18 | *APP;COX7B;NDUFA13;HIP1;NDUFA11;COX4I1;ATP2A3;PARK7;COX6A1;COX7C;UBE2L3;TUBA1B;TUBB6;KIF5B;RAC1;MAP2K3;COX8A;GABARAPL2;PRKCB;TUBB;GPX7;NDUFC2;CYBB;NDUFC1;SDHC;CSNK1E;COX6B1;ERN1;PSMA6;PSMA3;COX7A2L;PSMA4;PSMA2;NDUFS6;CSNK2B;CAT;ULK1;SLC25A5;UBA52;SLC25A6;NDUFB8;NDUFB7;UQCRB;NDUFB10;NDUFB11;NDUFB4;ITPR1;ATP5MC2;ITPR2;NDUFB1;ATP5MC3;UQCR11;COX7A2;UQCR10;KLC1;COX5A;GABARAP;PSMA7;UBE2J1;ATP5MC1;PSMB6;PPP3R1;ATP5F1A;ATP5F1B;UBB;PSMB3;ATG101;PSMB1;UBC;CYC1;BID;RPS27A;MAP2K6;ATP5PF;NDUFA7;XBP1;FUS;NDUFA4;ATP5PB;NDUFA1;UBE2G2;ATP5F1C;COX6C;ATP5F1D;ATP5F1E;CCS;UQCRQ;GNAQ;CALM2* |
| Thermogenesis | 4.325148716619073E-17 | *COX7B;NDUFA13;SMARCB1;KDM1A;NDUFA11;COX4I1;COX6A1;COX7C;ACTB;ACTG1;CREB3L2;COX8A;MAP2K3;RPS6;NDUFC2;SDHC;NDUFC1;ARID1B;COX6B1;COX7A2L;NDUFS6;NDUFB8;NDUFB7;COX16;NDUFB10;UQCRB;NDUFB11;NDUFB4;ATP5MC2;ATP5MC3;NDUFB1;UQCR11;COX7A2;UQCR10;COX5A;ATP5MC1;ATP5F1A;ATP5F1B;COX14;CYC1;ATP5MG;ATP5MF;ATP5ME;SMARCE1;ATP5PF;NDUFA7;ATP5PB;NDUFA4;NDUFA1;ATP5F1C;COX6C;ATP5F1D;ATP5F1E;UQCRQ;GNAS;NDUFAF3;GRB2* |

| | | |
|---|---|---|
| Alzheimer disease | 3.154714734930592E-16 | *APP;COX7B;NDUFA13;NDUFA11;COX4I1;ATP2A3;COX6A1;COX7C;TUBA1B;TUBB6;KIF5B;COX8A;PSENEN;TUBB;CYBB;NDUFC2;SDHC;NDUFC1;CSNK1E;COX6B1;ERN1;PSMA6;COX7A2L;PSMA3;PSMA4;PSMA2;NDUFS6;CSNK2B;ULK1;SLC25A5;SLC25A6;NDUFB8;NDUFB7;NDUFB10;UQCRB;NDUFB11;NDUFB4;ATP5MC2;ITPR1;ATP5MC3;ITPR2;NDUFB1;UQCR11;COX7A2;UQCR10;KLC1;COX5A;PSMA7;RTN4;ATP5MC1;APH1A;PSMB6;PPP3R1;ATP5F1A;ATP5F1B;PSMB3;ATG101;PSMB1;CYC1;BID;ATP5PF;XBP1;NDUFA7;ATP5PB;NDUFA4;NDUFA1;ATP5F1C;COX6C;ATP5F1D;ATP5F1E;UQCRQ;GNAQ;CALM2* |
| Antigen processing and presentation | 2.91277589334995E-10 | *CIITA;HSP90AB1;CTSS;HLA-DMA;HLA-DMB;HLA-DOA;B2M;HLA-DQA1;CTSB;HLA-DPA1;PDIA3;CD74;HSP90AA1;HLA-B;TAPBP;CD4;CANX;PSME1;HLA-DPB1;PSME2;HLA-DRA;CALR;HLA-DRB1;HLA-DQB1;LGMN* |
| Phagosome | 9.96252292141832E-10 | *RAB5C;NCF1;CORO1A;ACTB;CTSS;ACTG1;SEC61A1;TUBB6;TUBA1B;HLA-DMA;HLA-DMB;SEC61G;CD36;RAC1;SEC61B;HLA-DOA;HLA-DQA1;ATP6V1F;HLA-DPA1;ATP6V1G1;ATP6V0B;ATP6V0E1;TUBB;STX7;HLA-B;M6PR;CYBB;CYBA;CANX;HLA-DPB1;HLA-DRA;CALR;ATP6V0C;HLA-DRB1;HLA-DQB1* |
| Non-alcoholic fatty liver disease | 1.6618469435748856E-9 | *NDUFA13;COX7B;NDUFB8;NDUFB7;NDUFA11;NDUFB10;UQCRB;NDUFB11;COX4I1;NDUFB4;NDUFB1;COX7A2;UQCR11;UQCR10;COX6A1;COX7C;COX5A;CYC1;RAC1;BID;COX8A;XBP1;NDUFA7;NDUFA4;NDUFA1;NDUFC2;SDHC;NDUFC1;COX6C;COX6B1;ERN1;COX7A2L;ITCH;UQCRQ;NDUFS6* |
| Bacterial invasion of epithelial cells | 7.620108292859876E-9 | *ACTR2;SEPTIN2;ARPC1B;CLTC;ARPC5L;CLTB;GAB1;CLTA;SEPTIN6;ARPC4;ARPC5;SEPTIN9;ACTB;SEPTIN11;RHOA;ACTG1;CD2AP;ARPC2;CDH1;ARPC3;ELMO1;RAC1;WASF2* |

| | | |
|---|---|---|
| Salmonella infection | 8.910557902528503E-8 | *ARF1;RAB5C;HSP90AB1;ARPC1B;ARPC5L;BRK1;TXN;KLC1;ACTB;MYL12A;PIK3CG;MYL12B;ACTG1;HSP90B1;PYCARD;TUBA1B;TUBB6;CYTH4;KIF5B;RPS3;NCKAP1L;SNX9;FLNB;RAC1;MAP2K6;MAP2K3;ACTR2;HSP90AA1;TUBB;M6PR;ARPC4;ARPC5;DYNLL1;RHOA;SNX18;ARPC2;ARPC3;TLR9;ELMO1;PLEKHM2;DYNLRB1;PFN1;MYD88* |
| Protein export | 9.90901017560801E-7 | *SEC61A1;SPCS3;SPCS2;SPCS1;SRP72;SEC61G;SEC61B;SRP14;SEC62;SEC11C;SEC63* |
| Vibrio cholerae infection | 9.90901017560801E-7 | *SLC12A2;ARF1;ATP6V0B;ATP6V1G1;ATP6V0E1;KDELR1;ACTB;ACTG1;PDIA4;SEC61A1;SEC61G;GNAS;KDELR2;SEC61B;ATP6V0C;ATP6V1F* |
| Shigellosis | 1.4158994233245686E-6 | *ARF1;ARPC1B;UBE2D2;ARPC5L;ITPR1;ITPR2;ACTB;GABARAP;MYL12A;SEPTIN11;MYL12B;ACTG1;PYCARD;CYTH4;UBB;UBC;RAC1;RPS27A;WASF2;GABARAPL2;ACTR2;SEPTIN2;FBXW11;PRKCD;SEPTIN6;ARPC4;ARPC5;SEPTIN9;RHOA;RBX1;RRAGC;ARPC2;ARPC3;ELMO1;RBCK1;UBE2V1;PFN1;TLN1;UBA52;MYD88* |
| Viral myocarditis | 2.5855651761357855E-6 | *HLA-B;ACTB;ACTG1;ICAM1;HLA-DMA;HLA-DMB;RAC2;HLA-DPB1;HLA-DRA;RAC1;HLA-DOA;BID;HLA-DQA1;EIF4G2;HLA-DRB1;HLA-DPA1;HLA-DQB1* |
| Lysosome | 2.821358707411542E-6 | *SCARB2;HEXB;CLTC;CTSZ;CLTB;CLTA;CTSS;GGA2;LAPTM4A;PSAP;AP1S2;AP3S1;CTSC;CTSB;CD164;ATP6V0B;M6PR;LAPTM5;IGF2R;NPC1;NPC2;MAN2B1;TPP1;CD68;ATP6V0C;LGMN* |
| Spliceosome | 5.9023985775696845E-6 | *SF3B5;DDX5;RBM25;EIF4A3;HNRNPU;SNU13;SNRPD2;SNRPD1;TXNL4A;HNRNPA1;HNRNPA3;FUS;LSM4;CDC40;LSM3;HNRNPM;LSM8;LSM7;HNRNPK;SNRNP27;SNRPE;SNRPF;SRSF5;HNRNPC;SNRPC;SRSF7;SNRPB;SRSF9* |
| Cardiac muscle contraction | 8.152824636018537E-6 | *COX8A;COX7B;UQCRB;TPM3;COX4I1;TPM2;ATP2A3;COX7A2;UQCR11;UQCR10;COX6A1;COX6C;COX7C;COX5A;COX6B1;COX7A2L;ASPH;SLC9A7;UQCRQ;CYC1* |

| | | |
|---|---|---|
| Influenza A | 9.752368691613747E-6 | *CIITA;ACTB;ACTG1;ICAM1;PYCARD;HLA-DMA;HLA-DMB;PABPN1;HLA-DOA;KPNA2;BID;HLA-DQA1;HLA-DPA1;IFNAR2;PRKCB;STAT2;MX1;DNAJC3;HNRNPUL1;OAS1;IRF7;HLA-DPB1;HLA-DRA;TLR7;SLC25A5;HLA-DRB1;MYD88;IFNAR1;HLA-DQB1;SLC25A6* |
| Asthma | 2.6320811735228854E-5 | *HLA-DMA;HLA-DMB;FCER1G;HLA-DPB1;FCER1A;HLA-DRA;HLA-DOA;HLA-DQA1;HLA-DRB1;HLA-DPA1;HLA-DQB1* |
| Human immunodeficiency virus 1 infection | 9.517977549057058E-5 | *ITPR1;ITPR2;SAMHD1;GNAI2;PPP3R1;GNG5;GNG7;CFL1;AP1S2;RAC2;ELOB;RAC1;BID;B2M;PAK2;MAP2K6;MAP2K3;PDIA3;PRKCB;FBXW11;HLA-B;RBX1;TAPBP;BST2;DDB1;CD4;GNAQ;GNB2;GNB1;CALR;CALM2;MYD88* |
| Endocytosis | 9.517977549057058E-5 | *VPS29;ARF1;RAB5C;SH3KBP1;ARPC1B;CLTC;ARPC5L;CLTB;CLTA;SNX3;GRK3;RAB11FIP1;SNX2;CYTH4;KIF5B;AP2S1;VPS36;GIT1;AP2M1;GIT2;ACTR2;IQSEC1;HLA-B;ARPC4;SNF8;ARPC5;ARFGAP3;RHOA;IGF2R;TGFBR2;ITCH;DAB2;ARPC2;ARPC3;CHMP4B;ARF5* |
| Allograft rejection | 2.187244253813071E-4 | *HLA-DMA;HLA-DMB;HLA-B;HLA-DPB1;GZMB;HLA-DRA;HLA-DOA;HLA-DQA1;HLA-DRB1;HLA-DPA1;HLA-DQB1* |
| Epstein-Barr virus infection | 2.2701963172561742E-4 | *ICAM1;HLA-DMA;HLA-DMB;BLNK;RAC1;BID;HLA-DOA;B2M;HLA-DQA1;HLA-DPA1;MAP2K6;LYN;IFNAR2;MAP2K3;PDIA3;USP7;SYK;STAT2;HLA-B;TAPBP;OAS1;HLA-DPB1;IRF7;HLA-DRA;CALR;VIM;HLA-DRB1;MYD88;HLA-DQB1;IFNAR1* |
| Yersinia infection | 2.6899112203834536E-4 | *MAP2K3;VAV3;ACTR2;GIT2;ARPC1B;ARPC5L;ARPC4;ARPC5;ACTB;RHOA;ACTG1;PYCARD;CD4;ARPC2;ARPC3;GNAQ;ELMO1;RAC2;RAC1;WASF2;MYD88;SKAP2;MAP2K6* |
| Proteasome | 2.6899112203834536E-4 | *PSMB6;PSMA6;PSMA3;PSMA4;PSMB3;PSMA2;POMP;PSMB1;PSME1;PSME2;PSMA7;PSMB10* |
| Fc gamma R-mediated phagocytosis | 4.795360813972455E-4 | *LYN;VAV3;ACTR2;GSN;NCF1;SYK;PRKCB;ARPC1B;PRKCD;ARPC5L;ARPC4;ARPC5;ARPC2;ARPC3;CFL1;RAC2;RAC1;WASF2* |
| Graft-versus-host disease | 5.20203721116499E-4 | *HLA-DMA;HLA-DMB;HLA-B;HLA-DPB1;GZMB;HLA-DRA;HLA-DOA;HLA-DQA1;HLA-DRB1;HLA-DPA1;HLA-DQB1* |

| | | |
|---|---|---|
| Type I diabetes mellitus | 6.389965769165125E-4 | *HLA-DMA;HLA-DMB;HLA-B;HLA-DPB1;GZMB;HLA-DRA;HLA-DOA;HLA-DQA1;HLA-DRB1;HLA-DPA1;HLA-DQB1* |
| Retrograde endocannabinoid signaling | 8.091804521922659E-4 | *NDUFA13;NDUFB8;NDUFA7;NDUFB7;NDUFA11;NDUFB10;PRKCB;NDUFB11;NDUFA4;NDUFB4;ITPR1;NDUFA1;ITPR2;NDUFC2;NDUFB1;NDUFC1;GNAI2;GNG5;NDUFS6;GNAQ;GNB2;GNG7;GNB1* |
| Fluid shear stress and atherosclerosis | 8.390081645821232E-4 | *MEF2A;MEF2C;HSP90AA1;HSP90AB1;NCF1;GSTP1;MGST2;CYBA;TXN;ACTB;RHOA;ACTG1;ICAM1;HSP90B1;THBD;SUMO3;SUMO2;RAC2;PECAM1;RAC1;CALM2;MAP2K6* |
| Leishmaniasis | 9.479295836070322E-4 | *NCF1;PRKCB;CYBB;CYBA;EEF1A1;HLA-DMA;HLA-DMB;HLA-DPB1;HLA-DRA;HLA-DOA;HLA-DQA1;HLA-DRB1;MYD88;HLA-DPA1;HLA-DQB1* |
| Tuberculosis | 9.479295836070322E-4 | *CIITA;RAB5C;LSP1;CORO1A;CTSS;PPP3R1;HLA-DMA;HLA-DMB;HLA-DOA;BID;HLA-DQA1;HLA-DPA1;CD74;ATP6V0B;FCER1G;SYK;IL10RA;RHOA;TLR9;HLA-DPB1;HLA-DRA;CALM2;ATP6V0C;HLA-DRB1;MYD88;HLA-DQB1* |
| Kaposi sarcoma-associated herpesvirus infection | 0.0011923487022006776 | *ITPR1;ITPR2;GABARAP;PIK3CG;ICAM1;PPP3R1;UBB;GNG5;GNG7;UBC;RAC1;BID;RPS27A;MAP2K6;LYN;IFNAR2;GABARAPL2;SYK;STAT2;HLA-B;GNB2;MAPKAPK2;GNB1;IRF7;UBA52;CALM2;IFNAR1* |
| RNA transport | 0.001542983817987833 | *EIF4A1;EIF4A3;FXR1;PNN;NUP42;SUMO3;SUMO2;EIF4EBP1;SAP18;EIF4EBP3;EIF4B;PAIP1;UBE2I;EIF1AX;FUS;UPF3A;EIF2S2;EIF1;EEF1A1;EIF3I;EIF3H;RNPS1;PABPC1;EIF3F;EIF3D;EIF4G2* |
| Pathogenic Escherichia coli infection | 0.0015973569901083086 | *ARF1;TMED10;ARPC1B;ARPC5L;BRK1;ACTB;ACTG1;PYCARD;TUBA1B;TUBB6;CYTH4;GNA12;RPS3;NCKAP1L;RAC1;PAK2;WASF2;ACTR2;TUBB;ARPC4;ARPC5;RHOA;ARPC2;ARPC3;NCL;TMBIM6;MYD88* |
| Rheumatoid arthritis | 0.0023192909061958617 | *ATP6V0B;ATP6V1G1;ATP6V0E1;ICAM1;TNFSF13B;HLA-DMA;HLA-DMB;HLA-DPB1;HLA-DRA;HLA-DOA;ATP6V0C;HLA-DQA1;HLA-DRB1;HLA-DPA1;HLA-DQB1;ATP6V1F* |

| | | |
|---|---|---|
| Autoimmune thyroid disease | 0.003674325841018449 | *HLA-DMA;HLA-DMB;HLA-B;HLA-DPB1;GZMB;HLA-DRA;HLA-DOA;HLA-DQA1;HLA-DRB1;HLA-DPA1;HLA-DQB1* |
| Ubiquitin mediated proteolysis | 0.0054220626246490045 | *UBE2I;FBXW11;UBE2D2;UBE2E3;UBE2E2;UBE2G2;UBE2L3;RBX1;ANAPC11;UBE2J1;DDB1;ITCH;UBB;UBC;ANAPC5;ELOB;STUB1;RPS27A;UBA52;UBE2K* |
| Intestinal immune network for IgA production | 0.006012403644259154 | *HLA-DMA;HLA-DMB;HLA-DPB1;HLA-DRA;HLA-DOA;HLA-DQA1;HLA-DRB1;TNFSF13B;HLA-DPA1;HLA-DQB1* |
| Toxoplasmosis | 0.006246569632068153 | *MAP2K3;CIITA;PDPK1;IL10RA;PIK3CG;GNAI2;HLA-DMA;HLA-DMB;HLA-DPB1;HLA-DRA;HLA-DOA;HLA-DQA1;HLA-DRB1;MYD88;MAP2K6;HLA-DPA1;HLA-DQB1* |
| Spinocerebellar ataxia | 0.0066472964295361805 | *NOP56;XBP1;PRKCB;ITPR1;ATP2A3;ITPR2;PSMA7;ERN1;PSMB6;PSMA6;PSMA3;PSMA4;ATG101;PSMA2;PSMB3;GNAQ;PSMB1;ULK1;SLC25A5;SLC25A6* |
| Human T-cell leukemia virus 1 infection | 0.007052278643489939 | *NRP1;SLC2A1;ICAM1;ANAPC11;POLB;PPP3R1;HLA-DMA;HLA-DMB;CREB3L2;HLA-DOA;B2M;HLA-DQA1;HLA-DPA1;HLA-B;TGFBR2;CD4;CANX;HLA-DPB1;HLA-DRA;ANAPC5;CALR;TCF3;TLN1;SLC25A5;HLA-DRB1;HLA-DQB1;SLC25A6* |
| Leukocyte transendothelial migration | 0.007052278643489939 | *VAV3;NCF1;PRKCB;CYBB;CYBA;ACTB;MYL12A;RHOA;ACTG1;ICAM1;MYL12B;GNAI2;RASSF5;RAC2;PECAM1;RAC1;CD99* |
| Platelet activation | 0.007052278643489939 | *LYN;FCER1G;SYK;ITPR1;ITPR2;ACTB;MYL12A;PIK3CG;RHOA;ACTG1;MYL12B;GNAI2;APBB1IP;VAMP8;GNAQ;GNAS;TLN1;FERMT3* |
| Fc epsilon RI signaling pathway | 0.00810840110918624 | *MAP2K3;VAV3;LYN;FCER1G;SYK;PDPK1;ALOX5AP;RAC2;FCER1A;GRB2;RAC1;MAP2K6* |
| Human cytomegalovirus infection | 0.009954284833833064 | *ITPR1;ITPR2;GNAI2;PPP3R1;GNG5;GNG7;CREB3L2;GNA12;RAC2;EIF4EBP1;RAC1;BID;B2M;MAP2K6;PDIA3;PRKCB;IL10RA;HLA-B;RHOA;TAPBP;GNAQ;GNB2;GNB1;GNAS;GRB2;CALR;CALM2* |
| Hematopoietic cell lineage | 0.010677469388855089 | *FLT3;CSF2RA;HLA-DMA;CD4;HLA-DMB;IL3RA;HLA-DPB1;HLA-DRA;CD37;CD36;HLA-DOA;HLA-DQA1;HLA-DRB1;HLA-DPA1;HLA-DQB1* |

| | | |
|---|---|---|
| B cell receptor signaling pathway | 0.012103532365716478 | *VAV3;LYN;SYK;PRKCB;LILRB1;LILRB4;LILRA4;PPP3R1;BLNK;RAC2;GRB2;RAC1;PIK3AP1* |
| Regulation of actin cytoskeleton | 0.012332643158392055 | *ARPC1B;ARPC5L;BRK1;ITGAE;ACTB;MYL12A;MYL12B;ACTG1;CFL1;GNA12;RAC2;NCKAP1L;RAC1;PAK2;WASF2;GIT1;VAV3;ACTR2;GSN;ARPC4;ARPC5;RHOA;ABI2;ARPC2;ARPC3;PFN1* |
| Various types of N-glycan biosynthesis | 0.017045234088994044 | *B4GALT1;RPN2;DAD1;HEXB;RPN1;MAN1A1;MGAT1;DDOST* |
| Cell adhesion molecules | 0.019405835177293097 | *SELPLG;HLA-B;ICAM1;HLA-DMA;CD4;ALCAM;HLA-DMB;SELL;CDH1;PECAM1;HLA-DPB1;HLA-DRA;HLA-DOA;CD99;HLA-DQA1;HLA-DRB1;HLA-DPA1;HLA-DQB1;NECTIN1* |
| Osteoclast differentiation | 0.019561993366523048 | *IFNAR2;NCF1;SYK;STAT2;CYBA;LILRB1;LILRB4;SIRPB1;LILRA4;TGFBR2;PPP3R1;TYROBP;BLNK;GRB2;RAC1;IFNAR1;MAP2K6* |
| Th17 cell differentiation | 0.020621367480175998 | *HSP90AA1;HSP90AB1;TGFBR2;PPP3R1;HLA-DMA;CD4;HLA-DMB;IRF4;HLA-DPB1;HLA-DRA;HLA-DOA;HLA-DQA1;HLA-DRB1;HLA-DPA1;HLA-DQB1* |
| Mitophagy | 0.021630066714005965 | *GABARAPL2;FIS1;USP15;UBB;CSNK2B;UBC;TOMM7;ULK1;RPS27A;UBA52;GABARAP* |
| Lipid and atherosclerosis | 0.03611819917196944 | *LYN;VAV3;MAP2K3;XBP1;HSP90AA1;HSP90AB1;NCF1;PDPK1;ITPR1;CYBB;CYBA;RHOA;ICAM1;HSP90B1;PYCARD;ERN1;PPP3R1;IRF7;CD36;RAC1;BID;CALM2;MYD88;MAP2K6* |
| Chemokine signaling pathway | 0.03611819917196944 | *LYN;VAV3;NCF1;PRKCB;STAT2;PRKCD;PIK3CG;RHOA;GNAI2;GRK3;GNG5;GNAQ;GNB2;CXCR3;GNG7;GNB1;ELMO1;RAC2;GRB2;RAC1;DOCK2;CCR2* |
| NOD-like receptor signaling pathway | 0.03650091844795935 | *IFNAR2;YWHAE;GABARAPL2;HSP90AA1;HSP90AB1;STAT2;PRKCD;ITPR1;CYBB;ITPR2;CYBA;TXN;GABARAP;RHOA;PYCARD;OAS1;IRF7;RBCK1;MYD88;IFNAR1;CTSB* |
| Apelin signaling pathway | 0.037954802356315244 | *MEF2A;GABARAPL2;MEF2C;RPS6;ITPR1;ITPR2;GABARAP;PIK3CG;GNAI2;GNG5;CDH1;GNAQ;GNB2;GNG7;GNB1;CALM2;MEF2D* |
| Parathyroid hormone synthesis, secretion and action | 0.04176884401924933 | *MEF2A;MEF2C;PRKCB;ITPR1;ITPR2;RUNX2;RHOA;GNAI2;NACA;GNAQ;CREB3L2;GNA12;GNAS;MEF2D* |
| Glutathione metabolism | 0.04962527092491372 | *GPX4;GSTP1;ODC1;GPX7;MGST2;LAP3;PGD;PRDX6;SRM* |
| Cluster 3 | | |

| | | |
|---|---|---|
| Ribosome | 1.256183083138035E-59 | *RPL4;RPL5;RPL3;RPL32;RPL31;RPLP1;RPLP0;RPL10A;RPL8;RPL6;RPL7;RPS15;RPS4X;RPL7A;RPL18A;RPS18;RPL35;RPL37;RPS11;RPL39;RPS13;RPS12;RPS9;RPS7;RPS8;RPS5;RPL22;RPS6;RPSA;RPS3A;RPL37A;RPL24;RPL26;RPL29;RPL10;RPL12;RPL11;RPL36A;RPS15A;RPL14;RPS3;RPL15;RPS2;RPL18;RPL17;RPL19;RPL35A;RPL22L1;RPS24;RPS23* |
| Coronavirus disease | 1.6118472874656225E-50 | *RPL4;RPL5;RPL3;RPL32;RPL31;RPLP1;RPLP0;RPL8;RPL10A;RPL6;RPL7;RPS15;RPS4X;RPL7A;RPL18A;RPS18;RPL35;RPL37;RPS11;RPL39;RPS13;RPS12;RPS9;RPS7;RPS8;RPS5;RPL22;RPS6;RPSA;RPS3A;RPL37A;RPL24;RPL26;RPL29;RPL10;RPL12;RPL11;RPL36A;RPS15A;RPS3;RPL14;RPL15;RPS2;RPL18;RPL17;RPL19;RPL35A;RPL22L1;RPS24;RPS23* |
| Parkinson disease | 7.398098103080486E-11 | *NDUFA11;UQCRB;NDUFA4;TUBB;ATP5MC2;ATP5MC3;NDUFB1;TXN;ATP5F1C;COX5A;UQCRH;COX6B1;TUBA1B;ATP5F1A;ATP5F1B;NDUFS5;GNAS;SLC25A5;PRKACB;SLC25A6* |
| Thermogenesis | 1.2217058794702704E-9 | *NDUFA11;UQCRB;NDUFA4;RPS6;ATP5MC2;ATP5MC3;NDUFB1;ATP5F1C;COX5A;UQCRH;ACTG1;COX6B1;ATP5F1A;ATP5F1B;NDUFS5;GNAS;PRKACB;ATP5MG* |
| Diabetic cardiomyopathy | 1.2217058794702704E-9 | *PARP1;NDUFA11;UQCRB;NDUFA4;ATP5MC2;ATP5MC3;NDUFB1;ATP5F1C;COX5A;UQCRH;COX6B1;ATP5F1A;ATP5F1B;NDUFS5;SLC25A5;GAPDH;SLC25A6* |
| Oxidative phosphorylation | 2.8325076699019745E-9 | *NDUFA11;UQCRB;NDUFA4;ATP5MC2;ATP5MC3;NDUFB1;ATP5F1C;COX5A;UQCRH;COX6B1;ATP5F1A;ATP5F1B;NDUFS5;ATP5MG* |
| Prion disease | 1.1602761040773818E-8 | *NDUFA11;UQCRB;NDUFA4;TUBB;ATP5MC2;ATP5MC3;NDUFB1;ATP5F1C;COX5A;UQCRH;COX6B1;TUBA1B;ATP5F1A;ATP5F1B;NDUFS5;SLC25A5;PRKACB;SLC25A6* |
| Huntington disease | 6.296109594471024E-8 | *NDUFA11;UQCRB;NDUFA4;TUBB;ATP5MC2;CLTA;ATP5MC3;NDUFB1;ATP5F1C;COX5A;UQCRH;COX6B1;TUBA1B;ATP5F1A;ATP5F1B;NDUFS5;SLC25A5;SLC25A6* |
| Alzheimer disease | 1.7663105433427665E-7 | *NDUFA11;UQCRB;NDUFA4;TUBB;ATP5MC2;ATP5MC3;NDUFB1;ATP5F1C;COX5A;UQCRH;RTN4;COX6B1;TUBA1B;ATP5F1A;ATP5F1B;NDUFS5;SLC25A5;GAPDH;SLC25A6* |

| | | |
|---|---|---|
| Amyotrophic lateral sclerosis | 7.44122180590117E-7 | *NDUFA11;UQCRB;NDUFA4;TUBB;ATP5MC2;ATP5MC3;NDUFB1;ATP5F1C;COX5A;UQCRH;ACTG1;COX6B1;TUBA1B;ATP5F1A;ATP5F1B;NDUFS5;CAT;HNRNPA1* |
| Pathways of neurodegeneration | 3.3662976040637246E-5 | *NDUFA11;UQCRB;NDUFA4;TUBB;ATP5MC2;ATP5MC3;NDUFB1;ATP5F1C;COX5A;UQCRH;COX6B1;TUBA1B;ATP5F1A;ATP5F1B;NDUFS5;CAT;SLC25A5;SLC25A6* |
| Necroptosis | 6.232564481219237E-4 | *HSP90AB1;PARP1;FTH1;HMGB1;SLC25A5;GLUL;PPIA;FTL;SLC25A6* |
| Ferroptosis | 8.959311390102675E-4 | *GPX4;FTH1;PCBP2;SLC40A1;FTL* |
| Non-alcoholic fatty liver disease | 0.0027517846340294944 | *NDUFA11;UQCRB;NDUFA4;NDUFS5;NDUFB1;COX5A;UQCRH;COX6B1* |
| Legionellosis | 0.03636567798554285 | *EEF1A1;EEF1G;IL18;HSPD1* |
| Salmonella infection | 0.048761102206712904 | *TUBA1B;HSP90AB1;TUBB;IL18;RPS3;TXN;GAPDH;ACTG1* |
| Retrograde endocannabinoid signaling | 0.048761102206712904 | *NDUFA11;GNG5;NDUFA4;NDUFS5;NDUFB1;PRKACB* |
| Base excision repair | 0.05091192214013534 | *PARP1;APEX1;HMGB1* |
| Glycolysis / Gluconeogenesis | 0.051502973175651626 | *LDHB;TPI1;ENO1;GAPDH* |
| Influenza A | 0.08494996713176824 | *CDK6;IL18;HLA-DRA;SLC25A5;ACTG1;SLC25A6* |
| Cardiac muscle contraction | 0.10986456539738385 | *UQCRB;COX5A;UQCRH;COX6B1* |
| RNA transport | 0.10986456539738385 | *EEF1A1;EIF4A1;SUMO2;EIF3H;EIF3E;EIF3D* |
| Gap junction | 0.10986456539738385 | *TUBA1B;TUBB;GNAS;PRKACB* |
| Vibrio cholerae infection | 0.12097521965749299 | *GNAS;PRKACB;ACTG1* |
| Pathogenic Escherichia coli infection | 0.1257204884953342 | *TUBA1B;TUBB;IL18;RPS3;GAPDH;ACTG1* |
| Dilated cardiomyopathy | 0.1257204884953342 | *ITGA4;GNAS;PRKACB;ACTG1* |
| Endocrine and other factor-regulated calcium reabsorption | 0.1257204884953342 | *GNAS;CLTA;PRKACB* |
| Spliceosome | 0.13548068328813853 | *SNRPD2;SNRPF;HNRNPA1;CTNNBL1;SRSF9* |
| Pyrimidine metabolism | 0.13548838200340124 | *DUT;NME2;NME4* |
| Mineral absorption | 0.15701313881390394 | *FTH1;SLC40A1;FTL* |
| Drug metabolism | 0.16009712004979224 | *DUT;IMPDH2;NME2;NME4* |
| HIF-1 signaling pathway | 0.16009712004979224 | *LDHB;RPS6;ENO1;GAPDH* |
| Cortisol synthesis and secretion | 0.16561644622640156 | *GNAS;PRKACB;PBX1* |
| PI3K-Akt signaling pathway | 0.16561644622640156 | *YWHAE;CDK6;HSP90AB1;ITGA4;GNG5;MYB;RPS6;EREG* |
| Glutamatergic synapse | 0.16561644622640156 | *GNG5;GNAS;GLUL;PRKACB* |

| | | |
|---|---|---|
| Tight junction | 0.16561644622640156 | *TUBA1B;RAB13;PRKACB;YBX3;ACTG1* |
| Glyoxylate and dicarboxylate metabolism | 0.19874241761021852 | *CAT;GLUL* |
| Pentose phosphate pathway | 0.19874241761021852 | *TALDO1;TKT* |
| Asthma | 0.20564632799463026 | *FCER1A;HLA-DRA* |
| Gastric acid secretion | 0.21181001077833705 | *GNAS;PRKACB;ACTG1* |
| Bacterial invasion of epithelial cells | 0.21181001077833705 | *CLTA;SEPTIN6;ACTG1* |
| Leishmaniasis | 0.21181001077833705 | *EEF1A1;ITGA4;HLA-DRA* |
| Propanoate metabolism | 0.2206403296562778 | *LDHB;HADHA* |
