## Supplementary Table 22 for "Heterogeneity of circulating epithelial cells in breast cancer at single-cell resolution: identifying tumor and hybrid cells"

Supplementary Table 22. The association of the number of CECs with breast tumor grade

| Grade | $CD45^+$ CECs | $CD45^+$ CECs (cluster 1) | $CD45^+$ CECs (cluster 2) | Diploid $CD45^+$ CECs | Aneuploid $CD45^+$ CECs (cluster 1) | Aneuploid $CD45^+$ CECs (cluster 2) | Diploid $CD45^+$ CECs (cluster 2) |
|---|---|---|---|---|---|---|---|
| 1 | 1.46* (0.74-2.12) | 0.58* (0.11-1.04) | 0.76* (0.40-1.04) | 0.61* (0.27-1.06) | 0.37 (0.11-0.70) | 0.13*/** (0.05-0.32) | 0.42* (0.25-0.60) |
| 2 | 0.57* (0.42-3.99) | 0.04* (0.00-0.34) | 0.29* (0.01-2.35) | 0.45 (0.08-3.99) | 0.00* (0.00-0.25) | 0.00 | 0.29* (0.00-2.35) |
| 3 | 12.64 (6.56-14.89) | 5.75 (3.17-6.77) | 5.98 (3.88-8.35) | 9.34 (4.40-14.67) | 0.80 (0.22-2.26) | 0.00 | 5.98 (3.88-8.35) |

Data are presented as Median (Quartile1-Quartile3); *p<0.05 compared to grade 3; **p<0.05 compared to grade 2.
