## Supplementary figures and images for "Heterogeneity of circulating epithelial cells in breast cancer at single-cell resolution: identifying tumor and hybrid cells"

### Supplementary Figure 1

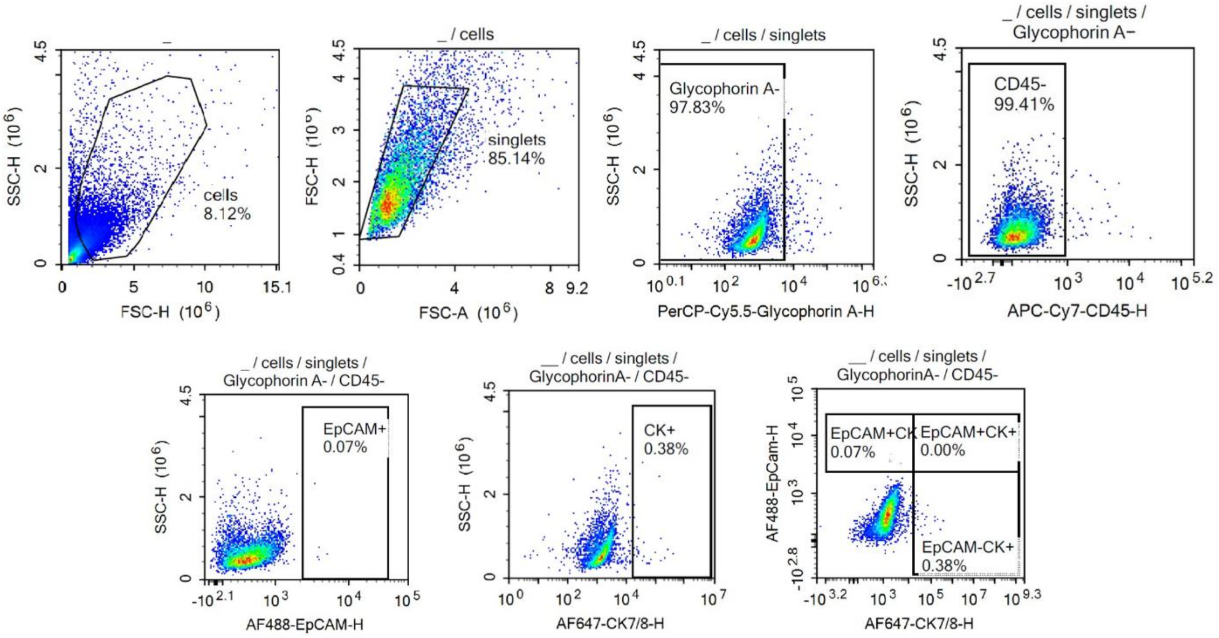


Supplementary Figure 1. Flow cytometry gating strategy for the detection of CD45¯EpCAM$^{+}$/KRT7/8$^{+}$ cells.
